# Systems-based approach for optimization of a scalable bacterial ST mapping assembly-free algorithm

**DOI:** 10.1101/2021.10.28.466354

**Authors:** Natasha Pavlovikj, Joao Carlos Gomes-Neto, Jitender S. Deogun, Andrew K. Benson

## Abstract

Epidemiological surveillance of bacterial pathogens requires real-time data analysis with a fast turn-around, while aiming at generating two main outcomes: 1) Species level identification; and 2) Variant mapping at different levels of genotypic resolution for population-based tracking, in addition to predicting traits such as antimicrobial resistance (AMR). With the recent advances and continual dissemination of whole-genome sequencing technologies, large-scale population-based genotyping of bacterial pathogens has become possible. Since bacterial populations often present a high degree of clonality in the genomic backbone (i.e., low genetic diversity), the choice of genotyping scheme can even facilitate the understanding of ancestral relationships and can be used for prediction of co-inherited traits such as AMR. Multi-locus sequence typing (MLST) fits that purpose and can identify sequence types (ST) based on seven ubiquitous genome-scattered loci that aid in genotyping isolates beneath the species level. ST-based mapping also standardizes genotyping across laboratories and is used by laboratories worldwide. However, algorithms for inferring ST from Illumina paired-end sequencing data typically rely on genome assembly prior to classification. Genome assembly is computationally intensive and is a bottleneck for speed and scalability, which are important aspects of genomic epidemiology. The stringMLST program uses an assembly-free, kmer-based algorithm for inferring STs, which can overcome the speed and scalability bottlenecks. Here we have systematically studied the accuracy and scalability of stringMLST relative to the standard MLST program across a wide array of phylogenetically divergent Public Health-relevant bacterial pathogens. Our data shows that optimal kmer length for stringMLST is species-specific and that genome-intrinsic and -extrinsic features can affect performance and accuracy of the program. While suitable parameters could be identified for most organisms, there were a few instances where this program may not be directly deployable in its current format. More importantly, we integrated stringMLST into our freely available and scalable hierarchical-based population genomics platform, ProkEvo, and further demonstrated how the implementation facilitates automated, reproducible bacterial population analysis. The ProkEvo implementation provides a rapidly deployable genomic epidemiology tool for ST mapping along with other pan-genomic data mining strategies, while providing specific guidance on how to optimize stringMLST performance for a wide variety of bacterial pathogens.

## Introduction

Modern epidemiological investigation of bacterial pathogens relies on rapid genomic characterization of new isolates routinely received by Public Health laboratories, along with bioinformatics programs for classification/comparison of genomic data from new isolates to existing data from many thousands of isolates [1][2]. While surveillance, source-tracking, and attribution are primary goals for use of whole-genome sequencing (WGS) data by Public Health agencies, there is growing interest in using the WGS data from thousands to hundreds of thousands of isolates of a given pathogen to study hierarchical genotypic classification at different levels of resolution, and prediction of population-specific traits contributing to virulence, ecological fitness, or antimicrobial resistance (AMR) [3].

Evolutionarily-related bacterial populations (e.g., species and subtypes of a species) share common genotypic backbones and the degree of clonality of a population provides a context for studying co-inheritance of core-and accessory-genes [4][5][6][7]. Core loci are shared by at least 99% of the genomes, whereas accessory loci represent a sparse ensemble that are not shared by all variants of a population, but jointly, the core and accessory genomic content form a given species forms its pan-genome [66]. Population genetic analysis of bacterial species using multi-locus methods have shown that multiple types of genotyping schemes can provide sufficient resolution for classifying populations based on genotypes beneath the species level [8][22][79][80][81], and the genetic relationships between different populations reveal the pattern of population diversification and structuring. This optimal level of genotypic resolution can be considered an informative genotypic unit that facilitates both ecological and epidemiological inquiries [57][82][83].

Multi-locus sequence typing (MLST) is a well-established and widely used genotyping technique that classifies bacterial genomes into sequence types (ST) [8][81]. ST classification based on traditional MLST is usually inferred from seven loci that are found ubiquitously in the species and are scattered around the genome. Highly curated, species-specific databases of allelic variation at MLST loci and distributions of STs are publicly available [27][30][67]. Historically, sequences for MLST loci were generated from locus-specific polymerase chain reaction (PCR) assays, but now are typically inferred from assembled WGS data [8][9][44].

ST-based classification provides useful and relevant genotypic units for epidemiological surveillance, population genetic analysis, and evolutionary inference. Relative to surveillance, ST-based genotyping standardizes the nomenclature for intra-and inter-laboratorial diagnostics and epidemiological inquiries worldwide [9][10][11]. With respect to evolutionary inference, isolates sharing alleles at five or more loci are more likely to be ancestrally related and are commonly classified as members of a clonal complex (i.e., group of STs that have shared a common ancestor very recently) [8][9][28][44]. The ST and clonal complexes provide a point of reference for genetic analysis (e.g., Linkage Disequilibrium -LD) to track inheritance/co-variance of accessory loci among different STs or clonal complexes. For example, LD analyses of pan-genomic content can inform phenotypic predictions for inheritance of unique traits in an ST related to virulence, environmental stability, or AMR [12][13][14].

Recent studies have shown that LD between specific sets of accessory genes such as AMR genes is an intrinsic property of some STs of specific bacterial pathogens, and rapid identification of a given ST from WGS data can quickly provide accurate prediction of AMR profiles [14][15]. There are important applications of this innovative strategy for treatment of patients and there is growing interest in defining LD relationships of AMR genes and virulence-associated genes with STs. A limiting factor in ST-based classification from WGS data, however, is the dependence of on genome assembly, most often from Illumina-based short-read data [27][30][67]. This bottleneck can also hinder ST-based surveillance efforts when many thousands of genomes are involved in a real-time analysis [16][21]. One approach to overcome the computational bottleneck is to use kmer-based ST classification directly from Illumina paired-end raw reads (assembly-free). The stringMLST algorithm was recently developed for this purpose and shown to provide computationally efficient, assembly-free ST classifications [17]. However, stringMLST has not been broadly evaluated across data from multiple Public Health-related bacterial pathogenic species. Given the remarkable variation in the intrinsic features of genomes from different species (e.g., compositional features, abundances of repetitive sequences, mutation rates, and rates of recombination); it is likely that parameters for assembly-free algorithms, such as stringMLST, will need to be empirically optimized for genetically diverse species and even subtypes (e.g., serotypes) of pathogens.

To test this hypothesis, we systematically examined the performance of stringMLST across a phylogenetically diverse set of bacterial pathogens that are of primary interest to Public Health. Our systematic approach compared accuracy of classifications using the standard MLST program vs. stringMLST at varying kmer lengths across many thousands of genomes from 15 different pathogenic species from three highly divergent phyla. Performance was first evaluated across a broad spectrum of phylogenetic diversity using genomes from isolates of *Acinetobacter baumannii (*Phylum: Proteobacteria)*, Clostridium difficile (*Phylum: Firmicutes)*, Enterococcus faecium (*Phylum: Firmicutes)*, Escherichia coli (*Phylum: Proteobacteria)*, Haemophilus influenzae (*Phylum: Proteobacteria)*, Helicobacter pylori (*Phylum: Proteobacteria)*, Klebsiella pneumoniae (*Phylum: Proteobacteria)*, Mycobacterium tuberculosis (*Phylum: Actinobacteria)*, Neisseria gonorrhoeae (*Phylum: Proteobacteria)*, Pseudomonas aeruginosa (*Phylum: Proteobacteria)*, Streptococcus pneumoniae (*Phylum: Firmicutes)*, Campylobacter jejuni* (Phylum: Proteobacteria), *Listeria monocytogenes (*Phylum: Firmicutes), *Salmonella enterica* (Phylum: Proteobacteria), and *Staphylococcus aureus* (Phylum: Firmicutes). To evaluate performance within diverse populations of a single species, we also measured performance of different kmer lengths across 23 of the most relevant serovars of *Salmonella enterica* subsp. *enterica* lineage I (*S*. *enterica*). Our results show that optimal performance of stringMLST can, for some species, be achieved with a single kmer length. However, several different bacterial species, and even different serovars of *S. enterica*, require species or serovar-specific kmer lengths for accurate ST classifications. Based on these findings, we further implemented stringMLST into the scalable ProkEvo platform [21], which produces hierarchical genotypic classifications from raw WGS data, and demonstrated how use of species-specific kmer settings for stringMLST enhances computational performance of the platform. [4][5][6][7][66][8][27][30][67][8][9][44] [14][15][27][30][67][16][21][17] [21]

## Materials and methods

This systems-based comparison between mlst and stringMLST was centered at capturing their differences in computational and statistical performances, and was accomplished through the following steps: 1) Narrow-scope comparative analysis across four phylogenetic distinct pathogens species; 2) Further examination of algorithmic performance within a single ecologically diverse bacterial species; and 3) Wide-scope comparison between phylogenetic divergent pathogenic species with Public Health relevance and with databases available on pubMLST (https://pubmlst.org/) for direct contrast between stringMLST and mlst.

### Datasets used for narrow-scope analysis

WGS data from four major bacterial pathogens, including *Campylobacter jejuni*, *Listeria monocytogenes*, *Salmonella enterica* subps. *enterica* lineage I (*S*. *enterica*) and *Staphylococcus aureus*, were selected to be used in this first part of the study. Our basis for that choice was due to three *a priori* defined criteria: 1) Select bacterial species from two main phylogenetic divergent Phyla: Firmicutes (*L*. *monocytogenes* and *S*. *aureus*) and Proteobacteria (*C*. *jejuni* and *S*. *enterica*); 1) Select zoonotic pathogens that continually cause human illnesses worldwide [18]; and 3) Consider their epidemiological relevance according to the Centers for Disease Control and Prevention (CDC) [19]. Specifically for *S*. *enterica*, 20 of the CDC most investigated serovars were represented in the dataset, which includes: *S.* Agona, *S.* Anatum, *S.* Braenderup, *S.* Derby, *S.* Dublin, *S.* Enteritidis, *S.* Hadar, *S.* Heidelberg, *S.* Infantis, *S.* Javiana, *S.* Johannesburg, *S.* Kentucky, *S.* Mbandaka, *S.* Montevideo, *S.* Muenchen, *S.* Newport, *S.* Schwarzengrund, *S.* Senftenberg, *S.* Thompson, and *S.* Typhimurium [20]. All publicly available raw paired-end Illumina reads for these organisms were downloaded from NCBI using parallel-fastq-dump [58]. Genomes used for all analyses were randomly selected from a previously downloaded samples of isolates containing *C*. *jejuni* (n = 21,919 genomes), *L*. *monocytogenes* (n = 19,633 genomes), *S*. *enterica* (n = 25,284 genomes), and *S*. *aureus* (n = 11,990 genomes) that were processed through the computational platform ProkEvo [21]. Specifically, our study design was comprised of random sampling of 600 genomes from each species, except for *S*. *enterica* for which 600 genomes were randomly drawn per serovar (list of all 20 serovars is shown in S1 Table). For each species and all *S. enterica* serovars, all ∼600 genomes were randomly split into three independent batches, with ∼200 genomes each. The batches were created to measure the degree of variation in classification accuracy when comparing the two ST-based genotyping programs. While for the majority of *S*. *enterica* serovars there were a total of 600 genomes available, the total number of raw reads publicly available on NCBI and ultimately used for the analyses for *S*. Agona, *S*. Derby, *S*. Johannesburg, *S*. Mbandaka and *S*. Senftenberg was 565, 590, 534, 535 and 563 respectively. The final total number of genomes used per species was *C*. *jejuni* (n = 600), *L*. *monocytogenes* (n = 600), *S*. *enterica* (n = 11,787), and *S*. *aureus* (n = 600). Text file containing all genome NCBI-SRA identifications is available here, https://figshare.com/articles/dataset/_/16735411.

### Software tools

#### mlst

mlst is a standard approach for scanning genome assemblies against traditional PubMLST typing schemes [22]. The genome assemblies can be in FASTA/GenBank/EMBL formats [22]. mlst (version 2.16.2) was installed using Anaconda, a package and environment manager that supports maintaining and installing various open-source conda packages [26]. mlst uses genome assemblies as an input. In order to generate assemblies from the raw Illumina paired-end reads, multiple pre-processing steps were performed. Quality trimming and adapter clipping were performed using Trimmomatic [50], while FastQC was used to check and verify the quality of the trimmed reads [51]. The paired-end reads were assembled *de novo* into contigs using SPAdes with the default parameters [52]. The quality of the assemblies was evaluated using QUAST [53]. The information obtained from QUAST was used to discard assemblies with 0 or more than 300 contigs, or assemblies with N50 value of less than 25,000 [21]. Finally, the assemblies that passed the quality control were used with mlst, where they are categorized into specific variants based on the allele combinations from seven ubiquitous, house-keeping genes [22]. A list of the exact versions of the bioinformatics tools used for generating assemblies for mlst are shown on S2 Table. We used mlst with the default options (e.g., *mlst --legacy --scheme <scheme> --csv <assembly.fasta> > <output>*) and the following schemes: “senterica” (for *S*. *enterica*), “campylobacter” (for *C*. *jejuni)*, “lmonocytogenes” (for *L*. *monocytogenes*), “saureus” (for *S*. *aureus*). The distribution of mlst comes with set of pre-downloaded ST schemes. More details about these MLST schemes, such as the number of alleles in the seven genes and the number of ST classifications available are shown on S3 Table. To obtain the ST classifications of all datasets, mlst was run as part of the computational platform ProkEvo [21]. Additionally, a separate run of the mlst program was used to conduct a pairwise comparison between the computational performance (runtime and memory usage) of mlst and stringMLST. The used mlst script can be found here, https://github.com/npavlovikj/MLST_stringMLST_analyses/blob/main/scripts/mlst.submit.

#### stringMLST

stringMLST is an assembly-and alignment-free rapid tool for ST-based classification of Illumina paired-end raw reads based on kmers [17]. For the analyses performed in this paper, we used stringMLST version 0.6.3. stringMLST was installed using Anaconda [26]. The first step of using stringMLST was to download the respective MLST scheme from PubMLST. In order to do this, a species name and a kmer length were needed. The default kmer length used and suggested by the developers of stringMLST for reads with lengths between 55 and 150 base pairs or nucleotides is 35 (common read length for Illumina paired-end reads) [17]. We used stringMLST with the default options (e.g., *stringMLST.py --getMLST --species=<species_name> -P <output_prefix> -k <kmer>*) and the following species names, “Salmonella enterica”, “Campylobacter jejuni”, “Listeria monocytogenes”, “Staphylococcus aureus”, and kmer lengths of 10, 20, 30, 35, 45, 55, 65, 70, 80, 90 independently (https://github.com/npavlovikj/MLST_stringMLST_analyses/blob/main/scripts/stringMLST_dbs.submit). More details about the downloaded MLST schemes, such as the number of alleles in the seven genes and the number of ST classifications available are shown on S3 Table. After the MLST scheme was downloaded and prepared, the final step was to run “stringMLT.py –predict” for the ST classification. For this, we ran stringMLST with the databases *a priori* created and the respective paired-end raw reads and kmer lengths of 10, 20, 30, 35, 45, 55, 65, 70, 80, 90 independently (e.g., *stringMLST.py --predict -d <directory_raw_reads> -p -r -t -x -P <database_prefix> -k <kmer> -o <output>*) (https://github.com/npavlovikj/MLST_stringMLST_analyses/blob/main/scripts/stringMLST.submit). Our choice of using an increasing gradient of kmer lengths was to evaluate whether the kmer length parameter could be optimized to enhance ST-based classification accuracy across bacterial species. Lastly, stringMLST was also integrated as part of the computational platform ProkEvo for a rapid ST-based genotyping as part of a hierarchical genotypic scheme [21][57]. This implementation can be found here, https://github.com/npavlovikj/MLST_stringMLST_analyses/tree/main/Prokevo_stringMLST.

#### ProkEvo-based MLST classifications

In order to compare the ST-based classification accuracy and conduct other statistical analysis (e.g., identifying major contributing factors influencing ST-based classifications) between mlst version 2.16.2 (assembly-dependent) and stringMLST version 0.6.3 (assembly-independent), all initial ST calls for all selected genomes, across all four species (*C*. *jejuni*, *L*. *monocytogenes*, *S*. *aureus* and *S*. Typhimurium), were done using mlst [22] through the computational platform ProkEvo [21]. In brief, ProkEvo uses bacterial Illumina raw paired-end sequences as an input, and the following steps are sequentially done prior to ST-based genotyping using mlst: Trimmomatic for sequence trimming [50], FastQC for quality control of the trimmed reads [51], SPAdes for *de novo* genome assembly [52], and QUAST for quality assessment of the genome assemblies [53]. More information on how to install and use ProkEvo for hierarchical bacterial population genomic analyses can be found here, https://github.com/npavlovikj/prokevo.

### Genome-intrinsic and –extrinsic factors that can influence algorithmic performance

Both genome-intrinsic and –extrinsic factors were considered to determine their contribution on the accuracy of ST classifications when comparing mlst vs. stringMLST. The genome-intrinsic variables considered in these analyses were: number of contigs per genome, total number of nucleotides per genome (genome length), GC% content per genome, and dinucleotide composition of genomes. The number of contigs per genome, as well as the genome length, were calculated using the assembled contigs from SPAdes [52]. The number of contigs was calculated for each genome using the Linux “grep” utility (e.g., *grep “>” assembly.fasta | wc -l*). The total number of nucleotides per genome was calculated using the “getlengths” function from the AMOS package [54]. For this analysis, we used AMOS v3.1. “getlengths” provides the length for each contig, and a custom Bash script was used to summarize these values per genome. The GC% content was calculated using the program FastQC [51]. FastQC is used to check and verify the quality of the raw Illumina paired-end raw reads. With each pair of raw reads from all datasets, FastQC v0.11 was used. One of the statistics checked for read quality is GC% and this value was extracted with custom Bash script from the file “fastqc_data.txt” once the FastQC output was generated. Since FastQC outputs the GC% per read, the average of both reads was calculated as the final read GC%. The dinucleotide composition of the genomes was calculated with the function “compseq” from the EMBOSS package [55]. “compseq” calculates the frequency of words of a specific length (e.g., length is 2 in the case of dinucleotides) from given input genome sequences. For these analyses we used EMBOSS v6.6 with the command “*compseq -word 2 -outfile <output> assembly.fasta*” for all datasets and genomes *a priori* assembled with SPAdes [52]. Next, a customized Bash script was used to count the total number of occurrences of each dinucleotide for each genome across all bacterial species. Finally, all these outputs were merged per genome using custom Python script to facilitate statistical analyses and data visualization. The used scripts can be found here, https://github.com/npavlovikj/MLST_stringMLST_analyses/tree/main/scripts.

The genome-extrinsic variables used in the analyses presented in this paper were the total count of unique STs per database and the total count of unique alleles across all seven loci used for ST classification across all bacterial species. These genome-extrinsic variables were extracted from the PubMLST databases for both stringMLST and mlst using custom Bash scripts. While the first step of stringMLST is to download the most up-to-date available MLST scheme from PubMLST, the distributed version of mlst comes with set of pre-downloaded ST and allelic schemes. For each MLST scheme, the mlst distribution has a separate directory with 8 files - seven are “.tfa” files with the fasta sequences of the alleles for each locus, and one file (e.g., senterica.txt) contains the ST information (i.e., the total number of STs mapped including their specific allelic composition across all seven loci for that given species). To calculate the total number of unique STs, we used the Linux utility “wc” with the text file with ST information (e.g., *wc -l senterica.txt*). To calculate the total count of unique alleles across the seven loci, the “grep” Linux utility was used with the seven “.tfa” files (e.g., *grep “>” *.tfa | wc -l*). All calculations were done per bacterial species. The downloaded MLST scheme with stringMLST is in a separate directory for each organism and used kmer length. This directory had 12 files - seven are “.tfa” files with fasta sequences for all alleles across all seven loci, and one file has the ST profiles (e.g., Salmonella_enterica_profile.txt), while the remaining files contained information about the extracted kmers and additional config and log information. Similarly, the total number of unique STs for stringMLST was counted using the Linux utility “wc” with the text file with ST profile information (e.g., *wc -l Salmonella_enterica_profile.txt*), and the total count of unique alleles per loci was extracted using the “grep” Linux utility with the seven “.tfa” files (e.g., *grep “>” *.tfa | wc -l*). Similarly, all ST and allelic counts were carried out per bacterial species. With stringMLST, the MLST schemes are downloaded and prepared separately for each different kmer length used. However, the kmer length did not affect the number of STs and unique alleles per organism. Thus, these values remained the same across organisms and kmer lengths for stringMLST.

### Kmer-based distribution across ST programs

In order to assess the potential impact of random mapping or occurrence of kmers of different lengths across different bacterial species, we randomly chose 100 raw Illumina paired-end reads from the initial *C*. *jejuni*, *L*. *monocytogenes*, *S*. *aureus* and *S*. Typhimurium (major representative zoonotic serovar of *S*. *enterica*) isolates. For each read, we extracted all unique kmers of length 10, 20, 30, 35, 45, 55, 65, 70, 80 and 90 respectively, and counted their occurrence in the corresponding raw reads. This was done using DSK v2.2.0 [56] (https://github.com/npavlovikj/MLST_stringMLST_analyses/blob/main/scripts/dsk.submit). Next, the total number of kmer frequency was summarized per organism and kmer length, and the mean value was calculated to examine the distribution of different kmers across the raw reads. For each database created with stringMLST, a file with the kmer frequency for the used ST scheme was generated. Using the kmers generated from the raw reads and the stringMLST database, a relative frequency of the common kmers was calculated (calculated as a ratio between the common kmers and the unique kmers from all the kmers generated between the raw reads and the stringMLST database, e.g., *(common_kmers/unique_total_observations)*100*). The code used for this can be found in our GitHub repository (https://github.com/npavlovikj/MLST_stringMLST_analyses/tree/main/figures_code).

### Agreement in ST classification between programs

In order to assess the overall accuracy of stringMLST compared to the standard mlst approach for ST calls, a percentage of agreement in ST classification was calculated. For this, the initial dataset composed of 600 genomes from either *C. jejuni*, or *L. monocytogenes*, or *S. aureus* was selected, in addition to a total of 11,787 genomes across twenty zoonotic serovars of *S. enterica* (∼600 genomes per serovar, S1 Table). The program stringMLST was run with increasing kmer lengths ranging from 10 to 90 nucleotides. If both stringMLST and mlst produced identical ST calls, either “good” or “bad” ones, the call was a match. A “good” and “bad” call represent ST with a number or a missing/blank value, respectively. The remaining combinations were classified as a mismatch. Next, the percentage of agreement (concordance) was calculated with custom R base script (https://github.com/npavlovikj/MLST_stringMLST_analyses/tree/main/figures_code).

### Computational platforms

All computational analyses performed for this paper were done on Crane - one of the high-performance computing clusters at the University of Nebraska-Lincoln Holland Computing Center [23]. Crane is Linux cluster, having 548 Intel Xeon nodes with RAM ranging from 64 GB to 1.5 TB. The scalability of ProkEvo with stringMLST was tested on the Open Science Grid (OSG), a distributed, high-throughput computational platform for large-scale scientific research [24][25]. OSG is a national consortium of more than 100 academic institutions and laboratories that provide storage and tens of thousands of resources to OSG users. These sites share their idle resources via OSG for opportunistic usage. The OSG resources are Linux-based, and due to the different sites involved, the hardware specifications of the resources are different and vary.

### Computational performance

To evaluate the computational performance of stringMLST in comparison to the mlst program, we assessed the runtime and memory usage of both programs. For this, we chose four different datasets, *C*. *jejuni*, *L*. *monocytogenes*, *S*. *aureus* and *S.* Typhimurium (major representative zoonotic serovar of *S*. *enterica*), with three different batches of 200 genomes each, with a total of 600 genomes each. We ran mlst with all required steps, such as quality trimming and adapter clipping, *de novo* assembly, and assembly discarding on each dataset (see Section Software tools: mlst for more detailed description). Separately, we ran stringMLST with a range of 10 different kmer lengths (10, 20, 30, 35, 45, 55, 65, 70, 80, 90) on each dataset. For each organism, the runtime was calculated as an average of all 200 genomes per batch. In general, the runtime depends on multiple factors, such as the specification and capabilities of the used computational platform. Since the runtime can vary depending on these various factors, average statistics were used to show the central tendency of the runtime when comparing stringMLST vs. mlst. The runtime was calculated using the “date” command integrated in the Unix operating systems (e.g., *t=‘date +%s’; mlst --legacy --scheme senterica -- csv assembly.fasta > <output>; tt=‘date +%s’; total_time=$((tt-t))*). For each organism, the memory was calculated as the maximum memory recorded from all 200 genomes per batch, since all genomes were analyzed separately and concurrently. In the case of mlst, the recorded memory was the maximum memory of all the steps ran prior to mlst, such as trimming, *de novo* assembly, quality checking, filtering, and ST typing. The memory used for these steps considerably varies from a few MBs to a few GBs (e.g., filtering vs. *de novo* assembly), and since the memory is a physical limitation of the computational platform, the maximum used memory was calculated for each organism and batch. The memory used was calculated using the “cgget” command that tracks various parameters from the Linux Control Groups (cgroups) per running job (e.g., *mlst --legacy --scheme senterica --csv assembly.fasta > <output>; r=‘cgget -r memory.usage_in_bytes /slurm/uid_${UID}/job_${SLURM_JOBID}/’; mem=‘echo $r | awk -F: ‘{print $3}’*’).

### Incorporating stringMLST in ProkEvo

ProkEvo is a freely available and scalable computational platform capable of facilitating bacterial population genomics analyses while combining various independent algorithms in a portable pipeline [21]. One of the advantages of ProkEvo is its ability to facilitate the addition and removal of new steps and programs without disrupting its workflow. For instance, more details about adding new programs to ProkEvo are given here https://github.com/npavlovikj/ProkEvo/wiki/4.1.-Add-new-bioinformatics-tool-to-ProkEvo. By following these instructions, we were able to successfully add stringMLST to the current ProkEvo platform. The ultimate description of how stringMLST was integrated into ProkEvo can be found here, https://github.com/npavlovikj/MLST_stringMLST_analyses/tree/main/Prokevo_stringMLST.

### Comparison between mlst and stringMLST performance using ProkEvo

In order to compare the performance/accuracy of MLST and stringMLST as part of the ProkEvo platform, two subsets of the *C. jejuni*, *L. monocytogenes*, *S.* Typhimurium and *S. aureus* datasets used in this paper were selected. One subset was composed of 100 randomly selected genomes, while the second one contained 1,000. The subsets were randomly selected from the original datasets used in this paper. As part of ProkEvo, stringMLST was run with the default kmer length of 35. The ProkEvo workflows with mlst and stringMLST and the two datasets were individually run on Crane - one of the high-performance computing clusters at the Holland Computing Center. Once the four workflows finished, the performance of ProkEvo with mlst and stringMLST and the datasets with 100 and 1,000 genomes, respectively, was compared using: i) the total running time; ii) the percentage of non-classified STs; and iii) the percentage of agreement between programs. Since ProkEvo is an automated platform, a list of NCBI-SRA identifications was provided with the ProkEvo implementations with both mlst and stringMLST. In brief, ProkEvo manages all the dependencies and intermediate steps, and produces the final ST classification as an output.

### stringMLST-based kmer length optimization across phylogenetic divergent bacterial pathogens

To identify the optimal species-specific kmer length that minimizes the frequency of ST miscalls, we ran stringMLST with a range of different kmer lengths across phylogenetic divergent pathogenic species. First, we chose twenty-three *S. enterica* serovars (*S*. Agona, *S*. Anatum, *S*. Braenderup, *S*. Derby, *S*. Dublin, *S*. Enteritidis, *S*. Hadar, *S*. Heidelberg, *S*. Infantis, *S*. Javiana, *S*. Johannesburg, *S*. Kentucky, *S*. Mbandaka, *S*. Montevideo, *S*. Muenchen, *S*. Newport, *S*. Oranienburg, *S*. Poona, *S*. Saintpaul, *S*. Schwarzengrund, *S*. Senftenberg, *S*. Thompson, *S*. Typhimurium), and for each dataset we randomly selected 100 paired-end Illumina reads from NCBI-SRA. Second, for each dataset we ran mlst and stringMLST with kmer lengths ranging from 20, 30, 35, 40, 45, 50, 55, 60, 65, 70, 80, 90. The kmer length of 10 was excluded due to its poor performance in previous analyses. Additionally, we use data from fourteen pathogens with Public Health relevance and with available MLST schemes, to widen the scope of the analysis and assess the necessity of fine-tunning the kmer length on a more broadly selected collection of species. In particular, we chose the following pathogens: *Acinetobacter baumannii, Clostridioides difficile, Enterococcus faecium, Escherichia coli, Haemophilus influenzae, Helicobacter pylori, Klebsiella pneumoniae, Mycobacterium tuberculosis, Neisseria gonorrhoeae, Pseudomonas aeruginosa, Streptococcus pneumoniae, Campylobacter jejuni, Listeria monocytogenes*, and *Staphylococcus aureus*. For each pathogen, we randomly selected and downloaded 1,000 paired-end reads from NCBI-SRA and processed these reads separately with mlst and stringMLST. stringMLST was run with kmer lengths ranging from 20, 30, 35, 45, 55, 65, 70, 80, 90 and different schemes for the different pathogens. Similar to the *S*. *enterica* datasets, the kmer length of 10 was excluded from the analysis.

Across all datasets, the percentage of ST miscalls was calculated for stringMLST for each kmer length, whereby miscalls were defined as “bad” ST calls - calls with a missing or blank values. Next, for each dataset, the kmer length that equated with the lowest percentage of ST miscalls was recorded. For some datasets, multiple kmer lengths generated an identical lowest percentage for ST miscalls. In this case, we applied a two-folded approach to select the most optimal kmer length: 1) if kmer of length 35 was part of the array of kmer lengths that showed the most optimal results, we recorded kmer 35 as the optimal kmer length since that is the default and recommended value for stringMLST (parsimonious approach); or 2) if kmer of length 35 was not part of the kmer lengths that showed the most optimal results, we recorded the kmer with the highest value as the most optimal one, since in general our analysis showed that longer kmers consumed less computational resources and speed up the entire analysis. Ultimately, the optimal kmer length and the percentage of ST miscalls were visualized onto a core-genome phylogeny generated for all twenty-three *S. enterica* serovars, as well as for all fourteen pathogens including all twenty-three *S*. *enterica* serovars which jointly totaled fifteen pathogens (total of 37 genomes, one per species including one per serovar of *S*. *enterica*, were used to construct the 16S rRNA-based phylogeny for visualization purposes). The core-genome alignment for the twenty-three *S. enterica* serovars was generated using Roary with this set of parameters, “*roary -s -e --mafft -p 8 -cd 99 -i 95 ./prokka_output/*.gff -f roary_output*” and the phylogenetic tree was produced using FastTree [76]. The phylogenetic tree for the fifteen pathogens was generated using CLUSTALW (https://www.genome.jp/tools-bin/clustalw) with the 16S rRNA sequences of the selected 37 genomes generated by Prokka [78]. All phylogeny-based visualizations were done using iTOL [77], and the recorded and overlaid statistics were extracted with custom R scripts (https://github.com/npavlovikj/MLST_stringMLST_analyses/blob/main/figures_code/figures_code.Rmd).

In addition to calculating the percentage of ST miscalls for different kmer lengths with stringMLST, for each dataset we calculated the percentage of agreement (concordance) between mlst and stringMLST on ST calls (“good” or “bad”), as previously described here. Of note, when the stringMLST and mlst results were combined, the number of returned ST calls was not always 1,000 (the original size of the used datasets). If 1,000 reads are used with stringMLST, stringMLST generates ST calls for all 1,000 reads. On the other hand, when using mlst, a set of steps are used before mlst, including filtering, and a fraction of assemblies were disregarded due to poor quality. Thus, only genome sequences that passed through the mlst program and yielded a “good” or “bad” call were ultimately used to compare with stringMLST. The number of raw reads for each dataset, as well as the number of final reads from mlst used for these analyses are shown on S4 Table.

### Statistical analyses

To compare the overall performance and accuracy of mlst vs. stringMLST on ST-based classifications, the following statistics were used across all bacterial species datasets: 1) ST richness; 2) Simpson’s D index (1 – *D*) of diversity using ST counts as input data; 3) Proportion of non-classified STs (missing values or blank calls); and 4) Standard deviation of the proportion of non-classified STs. These statistics were calculated to evaluate the algorithmic performance on ST-based classification accuracy within and between bacterial species selected to be used in the narrow scope analysis (*C. jejuni*, *S. aureus*, *L. monocytogenes*, and *S. enterica*). ST richness was calculated by identifying the number of distinct STs present in each species. The Simpson’s D index of diversity (1-*D*) was used to calculate the degree of genotypic diversity across species, using the diversity() function available in the vegan (version 2.5-6) R library [29]. The proportion of non-classified STs was calculated using the counts of isolates or genomes that were not assigned a ST number after each run of either mlst or stringMLST. The standard deviation of the proportion of non-classified STs was calculated using the sd() function which is derived from an unbiased estimate of the sample variance corrected by *n* – 1 (*n* for number of observations). The frequency of genomes used for all analyses was calculated per batch and program across all species, including across serovars for *S*. *enterica*. The relative frequency of the most dominant ST lineages was also assessed across bacterial species.

PERMANOVA univariate or multivariate models were used to assess the significance (*p* < 0.05) and the degree of association between the genome-intrinsic and –extrinsic factors with the following dependent variables: ST richness, Simpson’s D index of diversity, or proportion of non-classified STs. Statistical models were built for each of the dependent variables separately. Multivariate models included either the combination of bacterial species and program, or serovars in the case of *S*. *enterica* and program. These multivariate models were stated to calculate the main and synergistic effects of the explanatory variables (e.g., species*program or serovar*program). Univariate models were also assessed for each of the dependent variables, using one of the following independent/explanatory variables: 1) Genome-intrinsic variables: median number of contigs, mean of the total count of nucleotides per genome, mean of the average GC% content per genome, standard deviation of the number of contigs, standard deviation of the total count of nucleotides per genome, and standard deviation of the average GC% content per genome; 2) Genome-extrinsic variables: species, serovar of *S*. *enterica*, program (mlst vs. stringMLST with kmer lengths of 10, 20, 30, 35, 45, 55, 65, 70, 80, 90), mean of the total count of unique STs per program, mean of the total count of unique alleles across all genes per program, and the Simpson’s D index of diversity per species. Statistical significance and strength of association between the dependent and independent variables were measured with *p*-values (*p* < 0.05) and *R*-squared, respectively. In the case of contig size (median), total number of nucleotides per genome (mean), and GC% content per genome, summary statistic values (median or mean) were calculated grouped by species and batch (there was a total of three batches per bacterial species or serovar). For the total count of STs and total number of alleles in the database, summary statistic values (mean) were calculated grouped by species, batch, and program. Lastly, the standard deviation of number of contigs, total count of nucleotides per genome, or GC% content per genome were calculated grouped by species. PERMANOVA models were run using the adonis() function with 1,000 permutations using the vegan (version 2.5-6) R library [29]. Principal component analysis (PCA) was used to analyze the dinucleotide distribution across species and across serovars for *S*. *enterica* with two dimensions using the prcomp() function. The PCA calculations and the selection of the number of PCs were done using the factoextra (version 1.0.7) library. Bar-plots, box-and-whiskers plots, and bivariate/trivariate scatter plots were used to assess and visualize the distribution and associations within and between dependent and independent/explanatory variables. The R software (version 4.0.3) and R libraries such as Tidyverse (version 1.3.0) were used to conduct all statistical analyses, and all R scripts are available here (https://github.com/npavlovikj/MLST_stringMLST_analyses/tree/main/figures_code). Data quality control was achieved with R base functions, in addition to the following packages: skimr (version 2.1.3) and visdat (version 0.5.3). Graphical visualizations were achieved using ggplot2 (version 3.3.2), GGally (version 2.1.2), and plotly (version 4.9.4.1). R code integrity was checked using the assertive (version 0.3-6) package.

## Results

The computational and analytical approaches used in this paper are shown on Fig 1. Our analytical approach was sub-divided into a narrow- and wide-scope analysis aiming at accomplishing two goals: 1) Comparing the computational performance and accuracy of MLST vs. stringMLST; 2) Optimizing the use of stringMLST for a wide range of bacterial species; and 3) Implementing stringMLST as part of the ProkEvo computational genomics platform. We initially used publicly available raw Illumina paired-end sequence data from *C. jejuni*, *L. monocytogenes*, *S. enterica* and *S. aureus*, to run stringMLST and mlst independently in order to compare the accuracy in ST-based classifications and assess the computational needs and performance in the overall analysis (narrow-scope step). For this narrow-scope analysis, we performed a detailed comparative analysis between these two programs including: i) analyses of computational performance and resources needed (e.g., average runtime per genome and maximum memory needed to analyze all genomes), and ii) statistical analyses to determine the accuracy of classifications (e.g., ST richness, Simpson’s D index of ST-based diversity, proportion of miscalls, and percentage of agreement or concordance between programs). For the wide-scope step of the analysis, we systematically evaluated the accuracy and concordance between mlst and stringMLST across a broader array of phylogenetic divergent pathogens with direct implication for Public Health (*A. baumannii, C. difficile, E. faecium, E. coli, H. influenzae, H. pylori, K. pneumoniae, M. tuberculosis, N. gonorrhoeae, P. aeruginosa, S. pneumoniae, C. jejuni, L. monocytogenes, S. enterica* and *S. aureus*). Combined with the intra-species analysis done across 23 serovars of *S. enterica*, our assessment aimed at revealing the optimized kmer length to be used with stringMLST in order to: i) minimize the percentage of ST miscalls, and ii) maximize the use of computational resources by speeding up the analysis. Lastly, we provided an implementation of stringMLST within ProkEvo - a freely available and scalable computational platform that facilitates hierarchical genotyping of bacterial populations including pan-genomic mapping [21].

**Fig 1.**
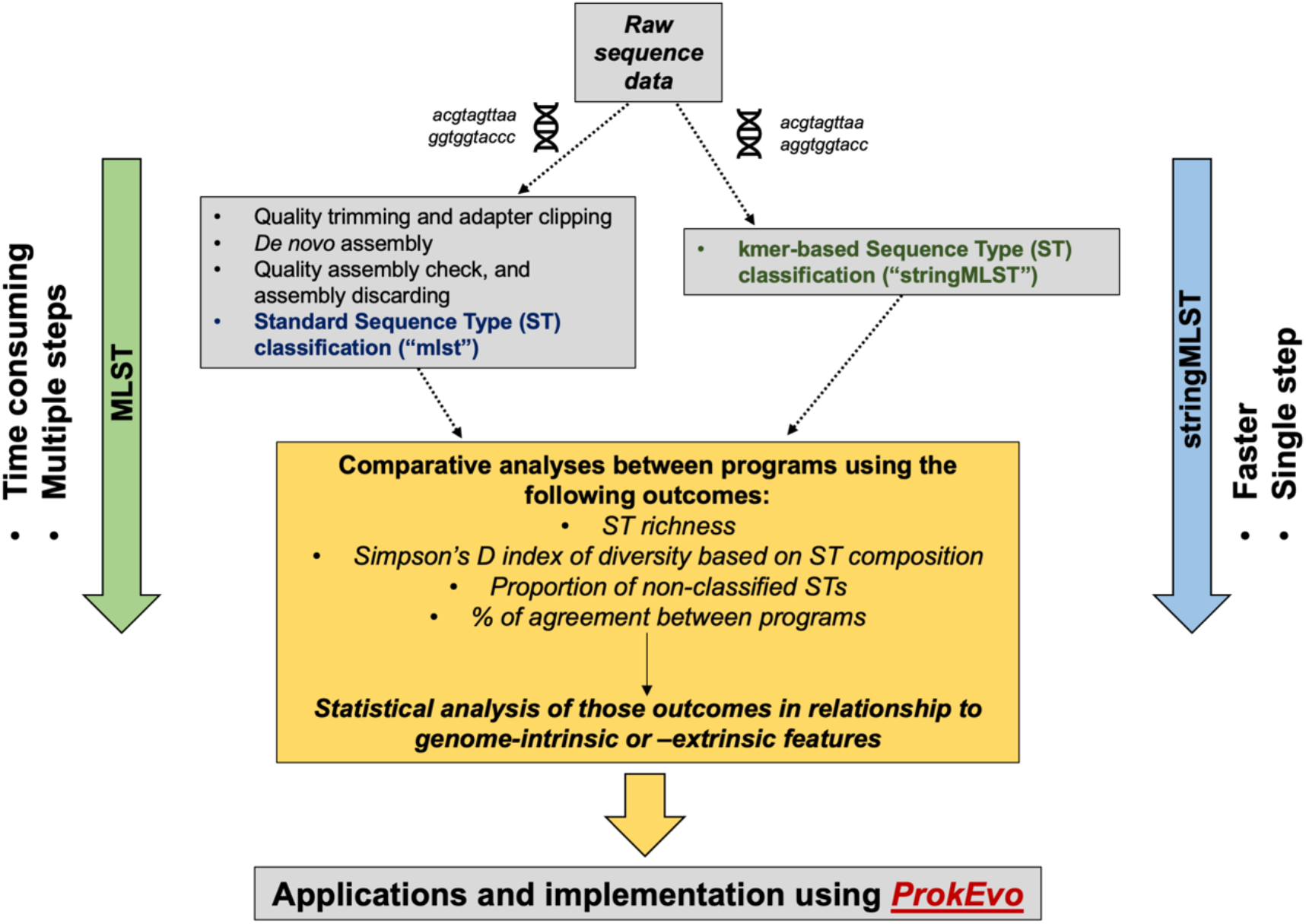
Computational workflow describing the analytical steps for a comparative analysis of two algorithms used for ST-based classification. From top-down, the first step (narrow-scope) of the analytical approach entailed the acquisition and processing of Illumina paired-end raw reads from four distinct pathogens (*C. jejuni*, *L. monocytogenes*, *S. enterica* and *S. aureus*), through an assembly-dependent (mlst) or assembly-free (stringMLST) approach for ST-based classification. Next, a set of comparative analyses encompassing measuring the computational performance, statistical metrics, and modeling were used to assess the accuracy and efficiency of mlst vs. stringMLST. Additionally, the contribution of genome-intrinsic and –extrinsic variables were used to identify explanatory factors that could impact the algorithmic efficiency across phylogenetic divergent species. Upon identification of inter-species differences in the performance of stringMLST, a wide-scope analysis was done to assess its accuracy across an array of fifteen phylogenetic divergent pathogenic species of bacteria with Public Health relevance. Ultimately, stringMLST was added to the computational platform ProkEvo to facilitate ST-based classification at scale, as part of a hierarchical-based approach for population genomic analyses.

### Computational performance

The computational performance between stringMLST and mlst was measured using two metrics: 1) The average computational runtime per genome; and 2) The maximum memory used per dataset. The average runtime in minutes per genome per batch between mlst and stringMLST with different kmer lengths, for *C. jejuni*, *L. monocytogenes*, *S. aureus*, and *S.* Typhimurium (major representative of *S*. *enterica*), is shown on Fig 2. While the runtime of mlst varies between 20 and 80 minutes per genome depending on the dataset used, all stringMLST runs with different kmers finished within a few minutes (ranging from ∼1-16 minutes when kmer 10 was included and ∼1-5 minutes when kmer 10 was excluded). Apart from stringMLST with kmer 10, all other kmer lengths showed a uniform runtime. The longer runtime observed with kmer 10 can be partially explained by the higher number of kmers that were generated and used for mapping (S1 Fig). The obtained results show that ST-based classifications are accomplished considerably more rapidly when carried out using stringMLST compared to the standard MLST program.

**Fig 2.**
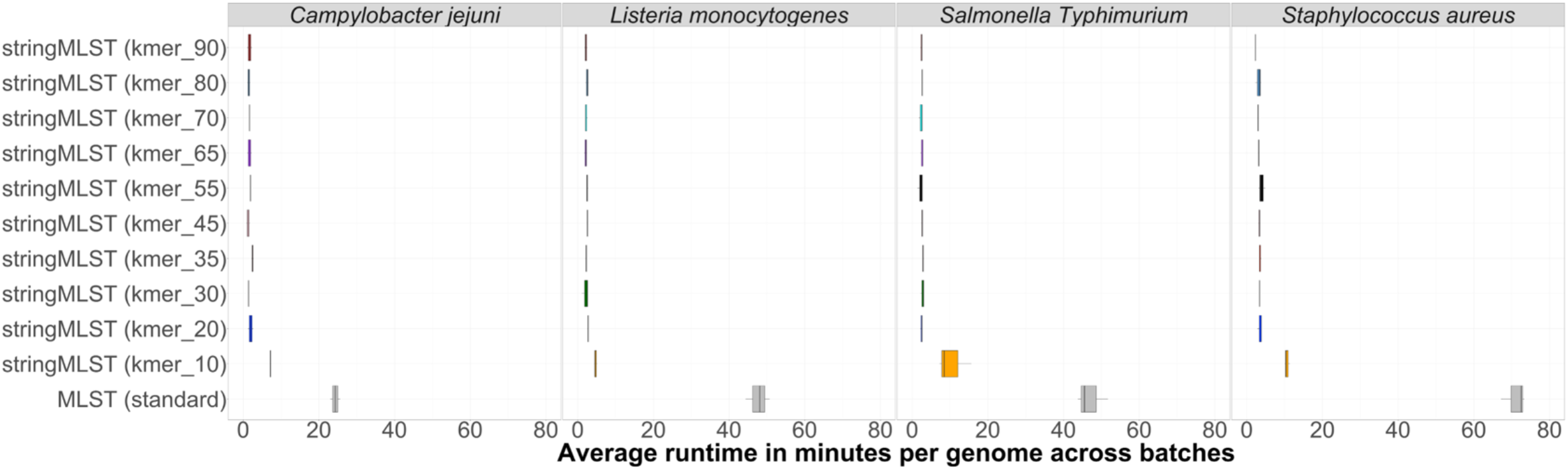
Box-and-whiskers plot showing the comparison of the average runtime per genome per batch (in minutes) needed by mlst and stringMLST for ST classification of genomes across four distinct bacterial species. In order to compare the average runtime used by mlst and stringMLST with different kmer lenghts, we chose four different datasets, including four phylogenetic divergent bacterial pathogenic species: *C. jejuni*, *L. monocytogenes*, one major serovar of *S. enterica* (*S*. Typhimurium) and *S. aureus -* using 600 randomly selected genomes for each species. These 600 genomes were randomly split into three batches with 200 genomes each. We then ran mlst with all required steps, such as quality trimming and adapter clipping, *de novo* assembly and assembly discarding, on each batch and dataset. Separately, we ran stringMLST with a range of 10 different kmer values (10, 20, 30, 35, 45, 55, 65, 70, 80, 90) on each dataset, including the default length of 35 (y-axis). For each organism, the runtime was calculated as an average of 200 genomes per batch - since there were three batches, three datapoints were used to depict the distribution of runtime in minutes (x-axis).

Additionally, a comparison of maximum memory used when both stringMLST and mlst were run for *C. jejuni*, *L. monocytogenes*, *S. aureus*, and *S*. Typhimurium (major representative of *S*. *enterica*) is shown on S2 Fig. Across all species, the range of maximum memory usage for mlst and stringMLST (across all kmers) was ∼2-16GBs and ∼3-30GBs respectively. Although the memory used across datasets is variable, none of the analyses we ran exceeded 30GBs of RAM. Since most high-performance computers can consistently provide resources from 32GBs to a few TBs of RAM, the memory available should not be considered a bottleneck for running either program.

### Factors that can influence ST-based classification

First, we describe the characteristics and composition of the data utilized for comparison between programs regarding ST-based classification in the narrow-scope approach (utilization of fewer phylogenetic diverse pathogen datasets). The frequency of genomes utilized per species and across programs is shown on S3A-D Fig. The frequency of *S*. *enterica* genomes was higher than other species because an equal sample of ∼600 genomes was taken from 20 representative zoonotic serovars (S4A-D Fig). An assessment of the proportion of the most dominant STs across species (proportion ≥ 2% - S5A-D Fig) or serovars of *S*. *enterica* (proportion ≥ 15% - S4N Fig) initially revealed a similar ST-based distribution across programs. Furthermore, genome-intrinsic and -extrinsic factors that could potentially impact the mlst vs. stringMLST algorithmic comparison and performance were *a priori* determined in the analysis. Among the genome-intrinsic factors considered across species were the number of contigs per genome (Fig 3A), the total number of nucleotides per genome (Fig 3B), GC% content per genome (Fig 3C), and the distribution and composition of dinucleotides per species (Fig 3D and S3E-F Fig). Similarly, the distribution of the genome-intrinsic factors was analyzed across all twenty serovars of *S*. *enterica* (S4G-L Fig). A correlogram (pairwise correlation analysis) was also used to assess the bivariate correlation (Pearson’s correlation coefficient) across genome-intrinsic variables, for either all four bacterial species (S3G Fig) or serovars across *S*. *enterica* (S4M Fig). At large, the differences observed in the distribution of genomic-intrinsic variables were species driven, with a strong uniformity found across serovars of *S*. *enterica*.

**Fig 3.**
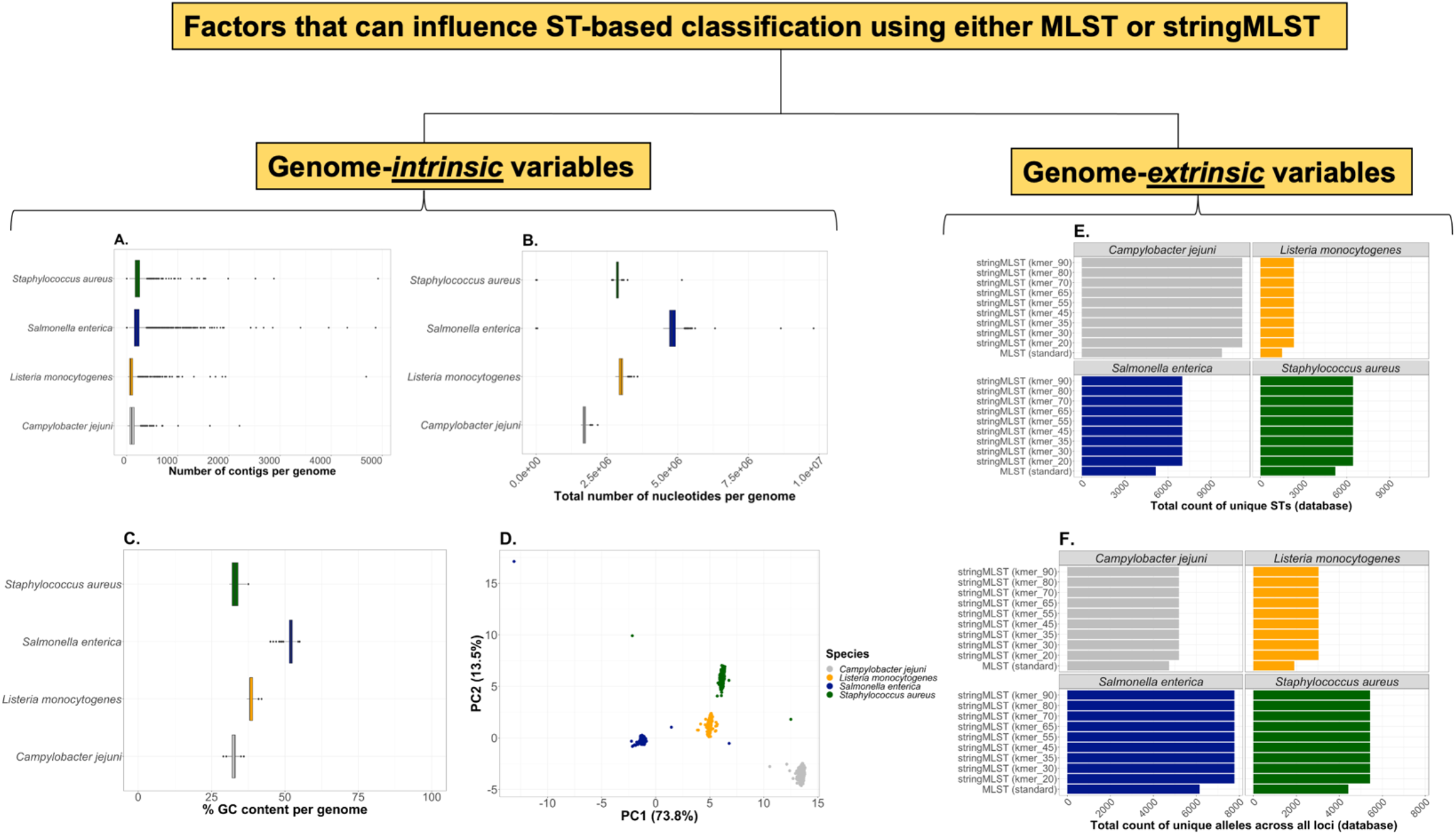
Genome-intrinsic and –extrinsic variables that can impact the accuracy of ST-based classification using either mlst (MLST-based genotyping) or stringMLST algorithmic approaches. Box-and-whiskers plot showing genome-intrinsic variables, varying in distribution according to the bacterial species (A-C as y-axis), that may affect ST-based classification, include: (A) Number of contigs per genome (x-axis); (B) Total number of nucleotides per genome (x-axis); (C) GC% content per genome (x-axis); and (D) Dinucleotide composition of genomes. (D) Inter-species PCA using the relative frequency of all pairs of dinucleotides (16 pairs) present in the genome as input data. Only two PCs are shown, and the percentage of variance explained by either PC is depicted in parenthesis. Bar plots showing genome-extrinsic variables that may influence the performance of mlst vs. stringMLST across species include but are not limited to: (E) Total count of unique STs per database (ST richness in the database used for mapping of raw reads or assemblies) (x-axis); and (F) Total count of unique alleles across all seven loci used for ST classification (x-axis). Specifically, the differences in ST richness and allelic composition in the databases reflect difference between mlst vs. stringMLST, and were not impacted by the kmer length (E-F, y-axis).

As for the genome-extrinsic variables, the total count of unique STs (for species - Fig 3E) and unique number of alleles across all seven loci (for species - Fig 3F), and across all batches were selected as factors that could influence the comparative analysis between mlst and stringMLST. Similarly, the genome-extrinsic variables were analyzed across all twenty serovars of *S*. *enterica* (S4E-F Fig). Of note, database size differences (number of STs and alleles) may directly influence the number of miscalls since it is expected that the larger the database is, the more likely STs are to be classified, or to find a match, and not be miscalled [30]. Considering the differences in genome-intrinsic and -extrinsic variable distribution across species, such factors were further utilized for assessing their statistical contribution in the accuracy of ST-based classification between mlst and stringMLST.

### Assessing the contribution of genome-intrinsic and – extrinsic variables

In order to assess the statistical association and contribution of each genomic-intrinsic and -extrinsic variable onto the accuracy of mlst vs. stringMLST on ST calls (narrow-scope analysis since it only included four bacterial species, *C. jejuni*, *S. aureus*, *L. monocytogenes*, and *S. enterica*), the following dependent variables (outcomes) were used in the PERMANOVA models: 1) ST richness (Fig 4A); 2) Simpson’s D index of ST diversity (Fig 4B); and 3) Proportion of non-classified STs (Fig 4C). Additionally, the standard deviation of the proportion of non-classified STs was measured as an auxiliary metric for accuracy (Fig 4D). At the species level, a multivariate model approach was used to examine the interaction of species and program (mlst vs. stringMLST), whereas all remaining analyses were done using univariate models containing each genome-intrinsic and -extrinsic variable for all three outcomes (S6A-L Fig, S7A-K Fig, S8A-L Fig).

**Fig 4.**
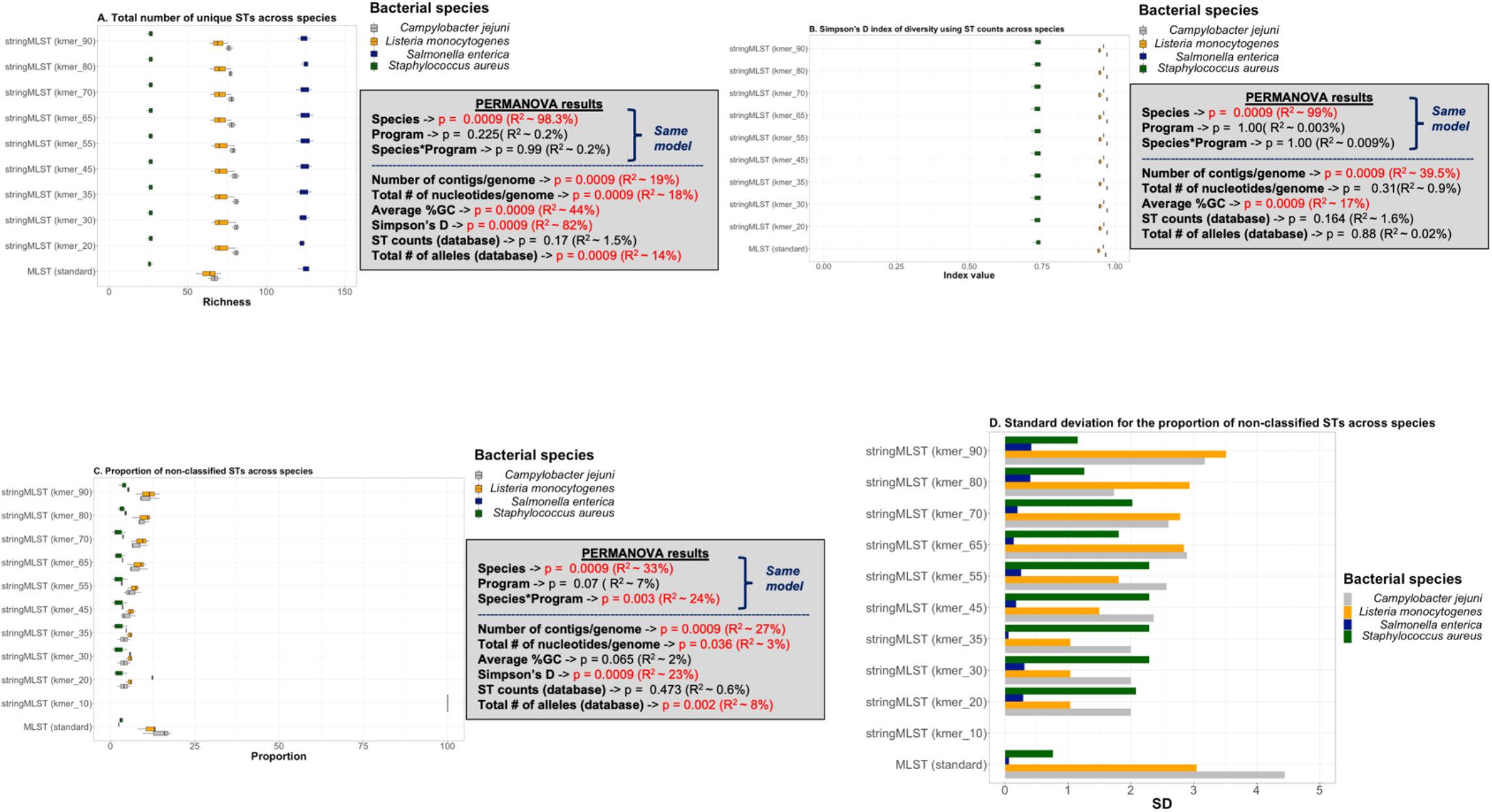
Statistical analysis of ST-based classification outcomes for comparison between mlst and stringMLST performance across bacterial species. (A-C) Box-and-whiskers plots A-C demonstrate the relationship between ST richness (x-axis), Simpson’s index of diversity (1 – *D*) based on ST composition (x-axis), or the proportion of non-classified STs (x-axis) across bacterial species (color-coded differently) and programs (y-axis), respectively. Along with plots A-C are depicted all PERMANOVA results including *p*-values (*p* < 0.05) and the univariate or synergistic contribution of factors measured by *R*-squared. PERMANOVA modeling was done in two specific ways: 1) A model including species, program, and their interaction, considering that those were the main variables of interest; and 2) All other results were calculated using univariate models and included modeling using genome-intrinsic (number of contigs per genome, total number of nucleotides per genome, and average GC% content) and – extrinsic (Simpson’s D index of diversity, ST and allelic counts per database) variables. (D) Bar plot depicting the distribution of the standard deviation (SD, y-axis) for the proportion of non-classified STs based on species (color-coded differently) and programs (y-axis).

For each variable, the significance and strength of association were assessed by jointly examining the *p*-value (*p* < 0.05) and *R*-squared, respectively. For both ST richness (Fig 4A) and the Simpson’s D index of diversity (Fig 4B), the difference between species explained most of the variation with ∼98.3% and ∼99%, respectively. As expected, based on the phylogenetic divergence of the four chosen pathogens, differences across species could largely be explained by genome-intrinsic variables associated with genome composition, such as: GC% content (*p* ∼0.0009, *R-*squared ∼44%) for ST richness, and the number of contigs per genome (*p* ∼0.0009, *R-*squared ∼39.5%) for the Simpson’s D index of diversity (Fig 4A-B). Notably, for both ST richness and the Simpson’s D index of diversity most of the differences between species could be explained by variation in genome composition (Fig 4A-B). Not surprisingly, co-linearity was observed between ST richness and the Simpson’s D index of diversity across species (Fig 4A). In the case of the proportion of non-classified STs (ST miscalls) (Fig 4C), most of the variation was explained by inter-species differences (*p* ∼0.0009, *R-* squared ∼33%), with the number of contigs per genome being the most important genome-intrinsic contributing factor (*p* ∼0.0009, *R-* squared ∼27%). As for the kmer length parameter used by stringMLST, results for ST richness and the Simpson’s D index of diversity were uniform across all lengths (Fig 4A-B). However, when examining the proportion of miscalls (Fig 4C) and the standard deviation of that proportion (Fig 4D), the data pointed toward the optimal kmer length being between 35 and 65 across all four species due to the intrinsic variance within the *S*. *enterica* data (narrow scope analysis). Specifically, this kmer length range was defined based on two criteria: i) minimization of the proportion of miscalls; and ii) less variation (standard deviation) around the average of ST-based miscalls. Of note, mlst demonstrated the highest proportion of miscalls and standard deviation of that proportion for both *L*. *monocytogenes* and *C*. *jejuni* (Fig 4C-D), and the kmer length 10 for stringMLST yielded very low accuracy and null results for ST richness and Simpson’s D index of diversity (Fig 4A-D). Differences between species across ST richness, Simpson’s D index of diversity, and proportion of ST miscalls along with all genome-intrinsic and - extrinsic variables across programs (mlst vs. stringMLST) were further examined here (Fig 5A-D, S9A-O Fig). In general, differences in ST-based calls across programs were largely influenced by the bacterial species dataset.

**Fig 5.**
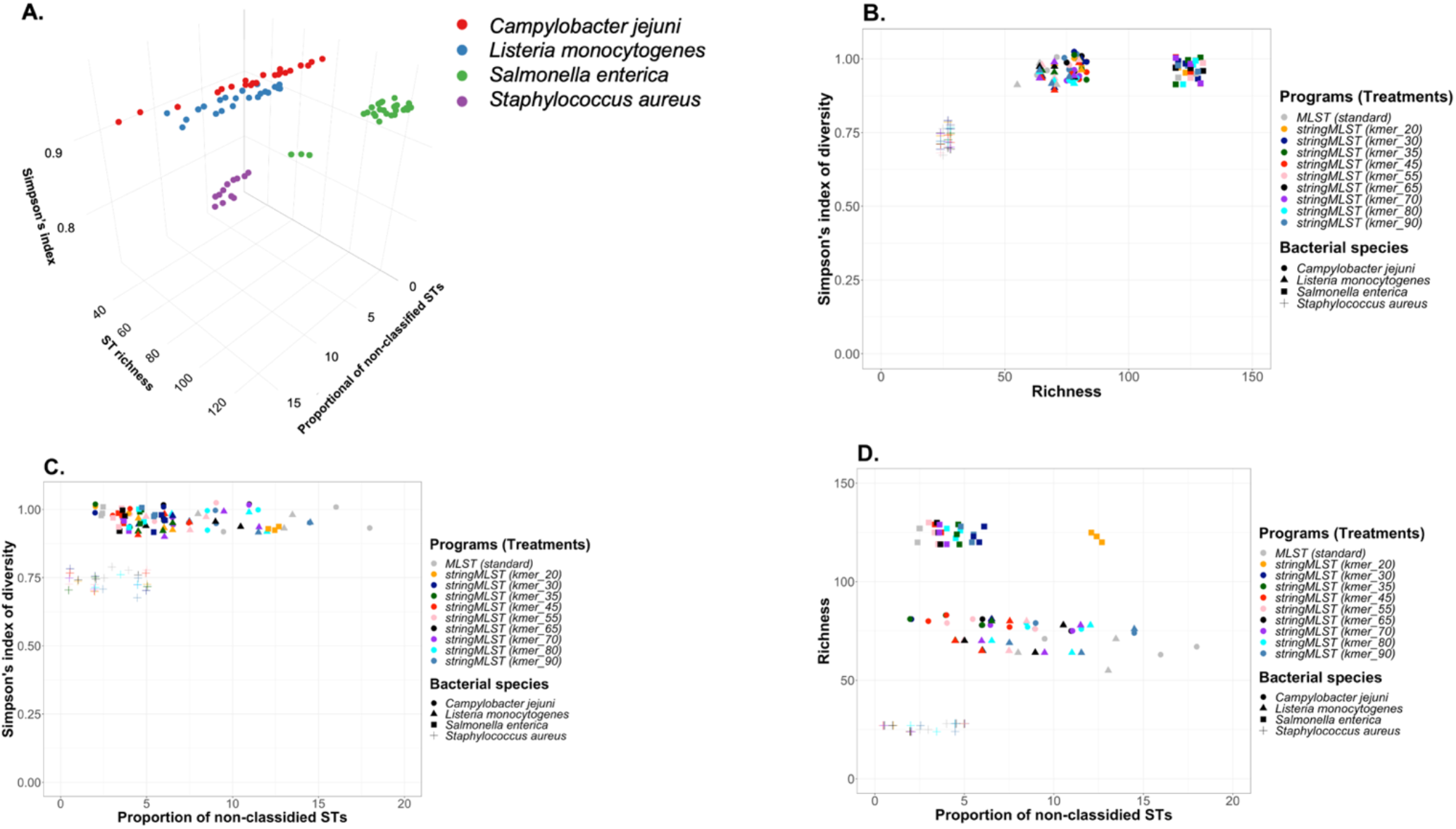
Multi-dimensional analysis of ST-based classification outcomes across different species using mlst vs. stringMLST. (A) Tri-dimensional scatter plot demonstrating species grouping based on the outcomes calculated using the ST classification across programs, including: 1) Simpson’s index of diversity (1 – *D*, Simpson’s index); 2) ST richness; and 3) proportion of non-classified STs. (B-D) Biplots demonstrating groupings formed across species and programs based on the same outcomes. Scatter plot B depicts groupings produced based on the relationship between Simpson’s index of diversity vs. ST richness (Richness); whereas scatter plot C shows the relationship between the Simpson’s index of diversity and the proportion of non-classified STs; and lastly, scatter plot D depicts the relationship between ST richness (Richness) and the proportion of non-classified STs.

Given the complexity and diversity of the *S*. *enterica* population structure [12], the stringMLST performance was analyzed across twenty zoonotic serovars (S4O-R Fig), and resulted in a significant and predominant contribution of the “serovar groupings” across all outcomes and PERMANOVA models (S10A-L Fig, S11A-K Fig, S12A-L Fig): ST richness (*p* ∼0.0009, *R-*squared ∼75.4%), Simpson’s D index of diversity (*p* ∼0.0009, *R-*squared ∼88%), and proportion of ST miscalls (*p* ∼0.0009, *R-* squared ∼35.4%). By assessing the distribution of the model outcomes, along with PERMANOVA model results and bivariate association between dependent and explanatory variables (S13A-R Fig), the results recapitulated the species-level results with the optimal kmer length for stringMLST being around 35 and 65, but also revealed the need to consider difference across *S*. *enterica* serovars prior to implementation. Combined, these accuracy-based results suggest that: i) stringMLST minimizes the ST miscalls compared to mlst in a species-specific fashion, and by consequence the optimal kmer length for stringMLST ranged from 35 to 65 overall; ii) the performance and accuracy of stringMLST can vary across species and serovars of *S*. *enterica* allowing for data-driven fine-tunning of the kmer length; and iii) the use of sequence platform with longer reads, which would maximize the number of contigs per genome, could directly alter both mlst and stringMLST accuracy in ST calls across species.

### Concordance between programs

Concordance between programs was calculated as the percentage of cases in which outputs from both mlst vs. stringmlst agreed in the call (“good” or “bad”). Results demonstrating the percentage agreement in ST calls between mlst and stringMLST with different kmer lengths are shown on Fig 6. Apart from kmer 10, across all species, the percentage of agreement between mlst and stringMLST varies between ∼81% and 97%. In the case of *L. monocytogenes*, *C. jejuni*, and *S. aureus*, the kmer length of 35 appears to be the optimal value to reach the same accuracy as mlst, which matches the original default and recommended parameter value for stringMLST [17]. However, for *S. enterica*, a higher percentage of agreement with MLST was achieved for kmer lengths of 55 and 65 (Fig 6). This *S*. *enterica*-related observation recapitulated the initial findings of decreased proportion of ST miscalls with higher kmer lengths (Fig 4C). Of note, our finding collectively showed that the kmer length of 10 yielded low accuracy when compared to mlst and other stringMLST kmer lengths. The most likely explanation for lower accuracy generated by kmer 10 is that shorter kmers are more likely to map unambiguously onto a genome when compared to other lengths. That high frequency of kmer length 10 on a given dataset reflects their higher likelihood of mapping to multiple regions of a genome (S1 Fig, S14A-B Fig). Overall, stringMLST is a rapidly deployable and optimizable ST-based genotyping algorithm that in this narrow-scope analysis proved to be applicable to four phylogenetic distinct pathogens.

**Fig 6.**
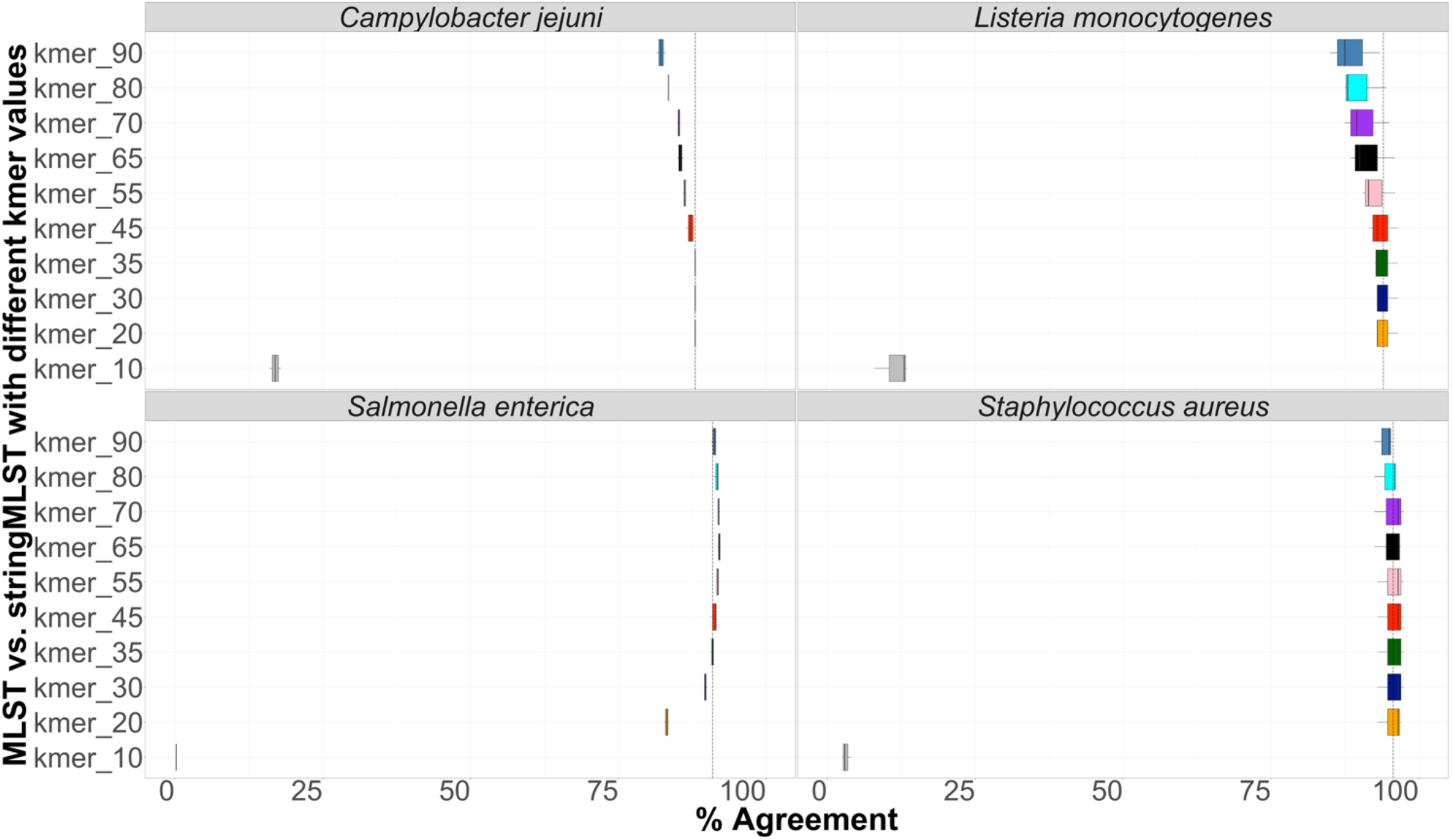
Box-and-whiskers plot depicting the concordance between mlst and stringMLST in ST calls. Four different datasets belonging to four phylogenetic distinct bacterial pathogens, including *C. jejuni* (600 genomes), *L. monocytogenes* (600 genomes), *S. enterica* (11,787 genomes from 20 different serovars) and *S. aureus* (600 genomes) were run with mlst and stringMLST for ST-based classification. In the case of stringMLST, kmer lengths varied from 10 to 90 to identify the optimal value (highest percentage of agreement with the standard MLST approach), across all four species (y-axis). If both programs outputted identical ST calls (either number of missing/blank value), the call was defined as a match; otherwise, it was identified as a mismatch, and the percentage of agreement (x-axis, concordance) was calculated accordingly. The dashed line on the x-axis represents the percentage agreement for the kmer value of 35 which is used as a default parameter by stringMLST.

### Optimization of stringMLST kmer length across phylogenetic divergent species

Here, we systematically investigated what kmer length would give the fewest ST miscalls (optimized length) with stringMLST across a broader array of phylogenetic divergent pathogens. Given our previous results, we first fine-tunned our investigation into the *S. enterica* population given the genetic and ecological diversity across serovars. For that, we selected data from twenty-three *S. enterica* zoonotic serovars and ran stringMLST with wide range of kmer lengths (20, 30, 35, 40, 45, 50, 55, 60, 65, 70, 80, 90). Fig 7A shows the core-genome phylogeny mapping of the optimized kmer length across all twenty-three serovars along with their corresponding percentage of ST miscalls. More detailed information on the distribution of the percentage of ST miscalls for all used kmer lengths (20, 30, 35, 40, 45, 50, 55, 60, 65, 70, 80, 90) is shown on S15A Fig. As it can be seen on Fig 7A, many serovars (*S*. Anatum, *S*. Braenderup, *S*. Javiana, *S*. Mbandaka, *S*. Montevideo, *S*. Oranienburg, *S*. Poona, *S*. Schwarzengrund, *S*. Senftenberg, *S*. Typhimurium) have 0% of miscalls when the default kmer length 35 was used. *S*. Infantis and *S*. Derby show the lowest percentage of ST miscalls (3% and 2% respectively) with higher value of kmer, e.g., 90. Interestingly, *S*. Saintpaul showed the highest percentage of ST miscalls when only considering the range of kmer lengths used for the initial analyses (10-90). To investigate this further, we ran stringMLST for *S*. Saintpaul with kmer lengths up to 240 (240 was chosen because the maximum read length for the *S*. Saintpaul dataset is 250 base pairs or nucleotides) (S15C-D Fig). As it can be seen on S15C Fig, the fewest ST miscalls for *S*. Saintpaul were produced when kmer of length 140 was used (22%). When comparing the percentage of ST miscalls between mlst and stringMLST, mlst outperformed stringMLST for the used datasets and range of kmer lengths (the percentage of miscalls for mlst ranged from ∼0-1% across all serovars, while the percentage of miscalls for stringMLST ranged from ∼0-85% depending on the serovar and kmer length used, with 2% being the median across all combinations of serovar and kmer length for stringMLST). In addition to the percentage of ST miscalls, we calculated the percentage of ST agreement between mlst and stringMLST with the range of kmer lengths (S15B Fig). While for some serovars this percentage is the highest when kmer with length 35 is used (e.g., *S*. Anatum, *S*. Braenderup, *S*. Javiana, *S*. Mbandaka, *S*. Montevideo, *S*. Oranienburg, *S*. Poona, *S*. Schwarzengrund, *S*. Senftenberg, *S*. Typhimurium), for other serovars (e.g., *S*. Derby, *S*. Dublin, *S*. Enteritidis, *S*. Hadar, *S*. Heidelberg, *S*. Infantis, *S*. Kentucky, *S*. Saintpaul) the percentage of ST agreement between the two programs was higher with higher kmer lengths (the percentage of ST agreement varied from ∼14-100% depending on the serovar and kmer length used, with 97% being the median across all combinations of serovar and kmer length).

**Fig 7.**
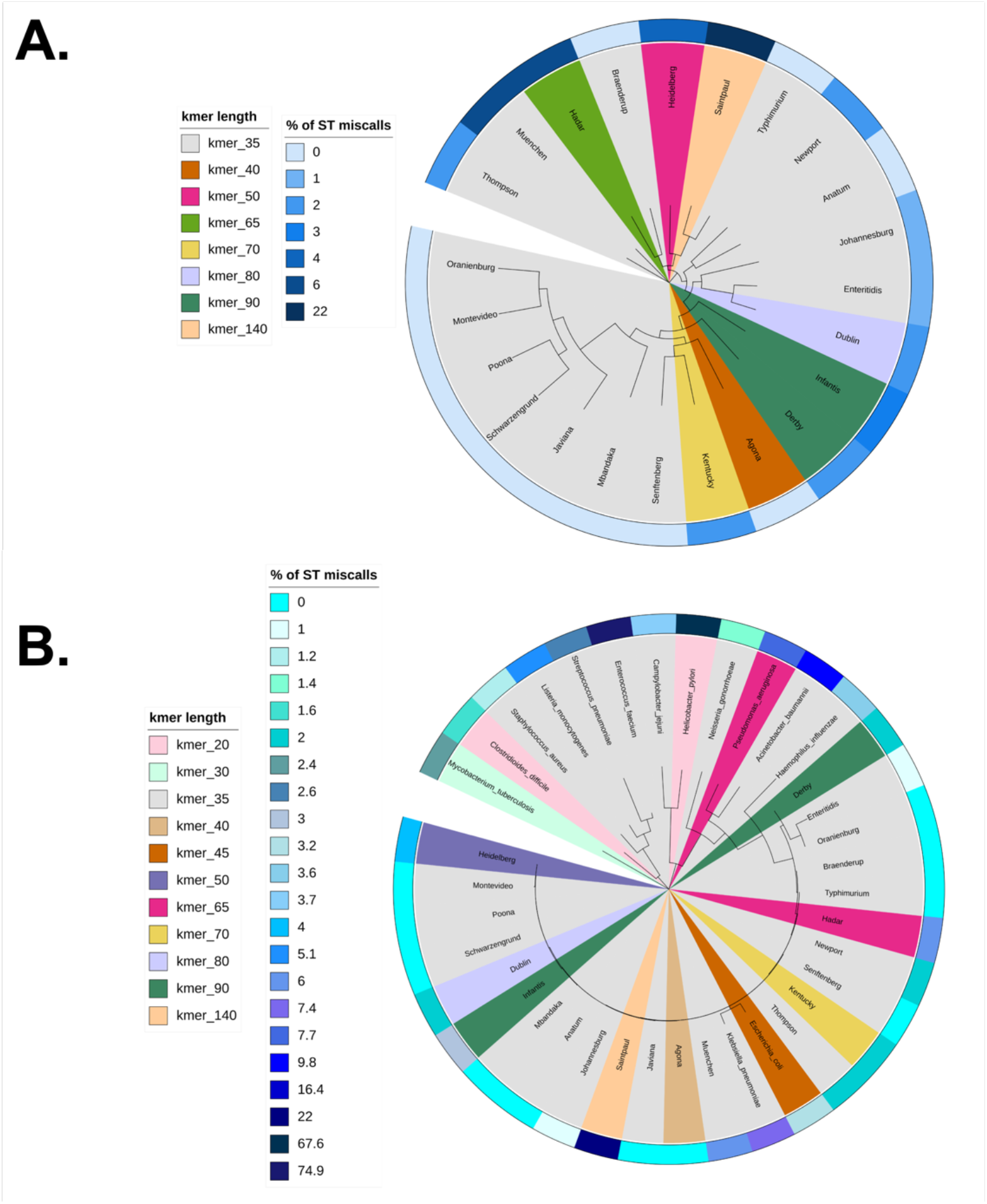
Phylogeny-guided display of optimal kmer length and algorithmic performance when using stringMLST for ST mapping across bacterial species. (A) Core-genome based phylogenetic display of stringMLST results across the twenty-three zoonotic serovars of *Salmonella enterica* subsp*. enterica* lineage I (*S*. *enterica*). The branches are colored based on the optimal kmer length which gives the lowest percentage of ST miscalls (ST calls that returned missing/blank values for stringMLST). The outer ring present in the phylogeny is colored based on the corresponding ST miscall percentages associated with each optimal kmer length. The dataset used to identify the optimal kmer length and percentage of ST miscalls was composed of 2,300 genomes (100 genomes per serovar) and the phylogenetic tree was generated using twenty-three core-genome sequences (one of each serovar to facilitate data visualization); (B) Single locus-based (i.e., 16S rRNA gene) phylogenetic display of stringMLST results across fourteen divergent bacterial pathogens, including twenty-three representative genomes across each zoonotic serovar of the *S*. *enterica* species. The tree branches are colored based on the optimal kmer length which minimizes the percentage of ST miscalls (ST calls that returned missing/blank values for stringMLST). The outer ring present in the phylogeny corresponds to ST miscall percentage associated with each optimal kmer length. The dataset used to identify the optimal kmer length and percentage of ST miscalls was composed of 14,000 genomes (1,000 genomes for each bacterial pathogen) and 2,300 *Salmonella* genomes (100 genomes per serovar). The phylogeny (B) was generated using the 16S rRNA representative sequences across 37 bacterial species (one for each species was used to facilitate visualization). All phylogeny-based visualization were generated using iTOL version 6.4.

In order to widen the scope of our phylogenetic-based analysis, we assessed the percentage of ST miscalls across varying kmer lengths for divergent bacterial pathogens with Public Health relevance. We selected 14 distinct organisms and ran stringMLST with wide range of kmer lengths (20, 30, 35, 45, 55, 65, 70, 80, 90). Fig 7B depicts the 16S rRNA-based phylogeny overlaid with the optimal kmer length that minimized the percentage of ST miscalls. Of note, the phylogeny contained fourteen distinct pathogens and twenty-three genomes across each serovar of *S. enterica.* The distribution of the percentage of ST miscalls for all used kmer lengths (20, 30, 35, 40, 45, 50, 55, 60, 65, 70, 80, 90) is shown on S15E Fig. While the percentage of ST miscalls varied between 0% and 22% across the *S. enterica* serovars as shown in Fig 7A, the percentage of miscalls was expectedly more variable for the fourteen bacterial pathogens, ranging from 1.2% to 74.9%. The datasets for *A. baumannii*, *C. jejuni*, *H. influenzae*, *K. pneumoniae*, *L. monocytogenes*, *N. gonorrhoeae*, *S. aureus* and *S. pneumoniae* showed the lowest percentage of ST calls with the default kmer length of 35. *C. difficile* and *M. tuberculosis* had minimized ST miscalls with kmer lengths of 20 and 30 respectively, while *P. aeruginosa* with kmer length of 65. Interestingly, for *E. faecium* and *H. pylori*, the optimal kmer lengths were 35 and 20, even though the percentage of miscalls was high (74.9% and 67.6%). To further investigate this, we ran stringMLST for *E. faecium* and *H. pylori* with kmer lengths up to 140 (140 was chosen because the maximum read length for the two datasets is 150 base pairs or nucleotides) (S15G-H Fig, S15K-L Fig). As can be seen on the Figures, the percentage of miscalls was higher with higher kmer lengths, and the lower kmer lengths yielded fewer miscalls, albeit this number was still considerably high. Additionally, we ran stringMLST on another set of randomly selected 100 paired-end reads for *E. faecium* (S15M-N Fig), *H. pylori* (S15I-J Fig) and *Enterococcus faecalis* (S15O-P Fig). These 100 reads were not part of the initial datasets and were chosen to validate that the initial random data selection was not completely biased. We also added *E. faecalis* here due to its close phylogenetic association with *E. faecium*. For *E. faecium* and *H. pylori* we observed the same pattern with 100 reads as with 1,000 reads. On the other hand, the pattern for *E. faecalis* was quite opposite with lowest percentage of ST miscalls of 5.43% for kmer 35. When comparing the percentage of ST miscalls between mlst and stringMLST, for some datasets, such as *C. jejuni*, *H. pylori*, *L. monocytogenes*, *M. tuberculosis*, *N. gonorrhoeae*, *S. aureus*, mlst performed worse than stringMLST (with increase in percentage of miscalls by ∼48%, 13%, 11%, 50%, 9%, 45% respectively). In addition to the percentage of ST miscalls, we calculated the percentage of ST agreement between mlst and stringMLST with the range of kmer lengths (S15F Fig). Of note, in the case of stringMLST, when the optimal kmer length was above the default parameter of 35, the ultimately selected kmer length was picked based on our empirical evidence for longer kmers being capable of speeding up the computational analysis.

In summary, while the default kmer length of 35 used by stringMLST performs accurately across many organisms, our systems-based approach encompassed the analysis of a broad array of phylogenetic divergent organisms revealed: i) intra- and inter-species variation in the percentage of ST miscalls requires fine-tunning of the kmer length parameter; ii) lack of association between taxonomy or phylogenetic placement of organisms and the optimal kmer length; and iii) unique species behave as outliers for which stringMLST cannot be directly applied with the default settings.

### Incorporating stringMLST in ProkEvo

ProkEvo was recently developed as an automated and scalable computational platform for bacterial population genomics analyses that uses the Pegasus Workflow Management System (WMS) [31]. In particular, ProkEvo facilitates the use of a hierarchical approach for population stratification with different layers of genotypic resolution. MLST-based classification of genomes into STs is part of this hierarchical approach that has been proven to be predictive of ecological traits such as AMR in *S*. *enterica* lineages [57]. However, ProkEvo currently only uses the standard mlst algorithm for ST calls [21]. As part of this paper, the stringMLST program was incorporated into ProkEvo without any disruption in its workflow. The workflow design of ProkEvo with both mlst and stringMLST is shown on S16 Fig.

In order to compare the performance of ProkEvo with mlst and stringMLST, randomly shuffled subsets derived from the original datasets used for *C. jejuni*, *L. monocytogenes*, *S.* Typhimurium, and *S. aureus* were used. One random subset contained 100 genomes, while the second one had 1,000 genomes. ProkEvo was run using either mlst or stringMLST on Crane, one of the high-performance computing clusters at the Holland Computing Center [23]. For mlst, the pipeline used was previously established and included a few required steps, such as quality trimming and adapter clipping, *de novo* assembly and assembly discarding prior to the ST mapping [21]. Based on the inter-species results shown here (Fig 6), the default kmer length of 35 was used with stringMLST for this comparison. The outcomes measured for this analysis were: i) total running time (Fig 8A); ii) the percentage of non-classified STs (Fig 8B); and iii) the percentage of agreement between programs (Fig 8C).

**Fig 8.**
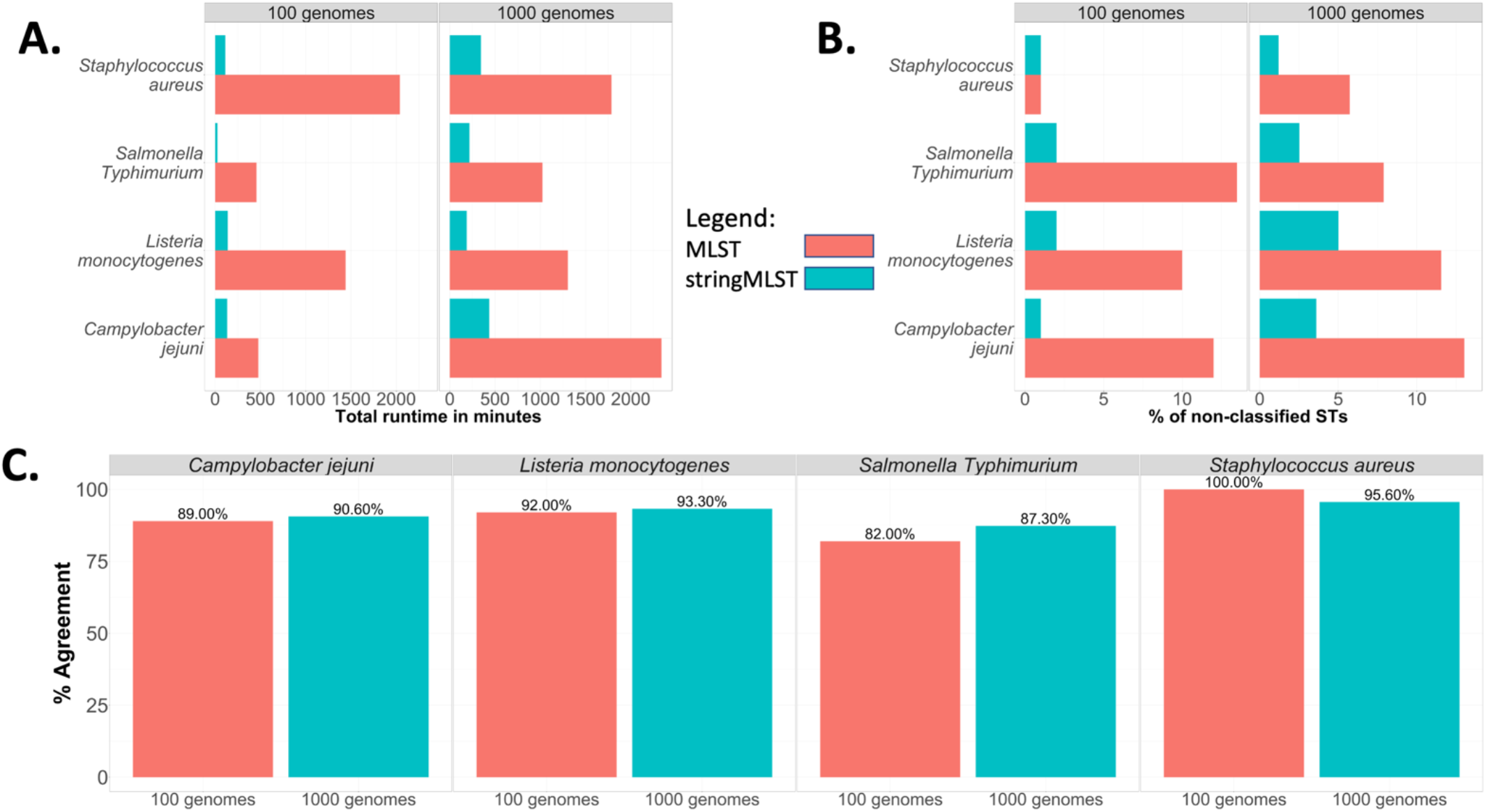
Comparison between the computational and statistical performance of mlst and stringMLST when using ProkEvo to run both programs. Two subsets, one with 100 and the second one with 1,000 randomly chosen genomes, were selected from *C. jejuni*, *L. monocytogenes*, one major serovar of *S. enterica* (*S*. Typhimurium) and *S. aureus* to compare the performance of running mlst or stringMLST through ProkEvo. The performance and statistical metrics used for comparison were: (A) Total runtime of individual workflow in minutes; (B) Percentage of non-classified STs (ST calls that returned missing/blank values); and (C) Percentage of agreement (concordance) between programs (“good” or “bad” ST calls that matched between mlst and stringMLST).

While the runtime of using ProkEvo with mlst varied from ∼8 to 34 hours for the subset containing 100 genomes, the runtime of ProkEvo with stringMLST varied from ∼25 minutes to 3 hours (Fig 8A). Similarly, for the larger datasets containing 1,000 genomes, the runtime of ProkEvo with mlst varied from ∼17 to 39 hours, while the runtime of ProkEvo with stringMLST varied from ∼4 to 8 hours. Regardless of the pathogen species tested, stringMLST speeded up the analyses ∼4 times when utilizing 1,000 genomes across species.

In terms of accuracy in ST classifications, the use of stringMLST considerably decreased the number of non-classified STs, regardless of the dataset size (100 or 1,000 genomes) and bacterial species (Fig 8B). In accordance, stringMLST resulted in a higher frequency of genomes classified as novel STs (ST numbers that were not classified by mlst) (S17 Fig). Additionally, the overall concordance between mlst and stringMLST varies from 82% to 100% across all datasets. The percentage of agreement is the lowest for *S.* Typhimurium, while it is the highest for *S. aureus* (Fig 8C). The lower proportion of ST miscalls and high percentage of agreement between programs for *S*. *aureus*, compared to other species, is associated with its higher degree of genetic homogeneity (fewer dominant STs) (S5 Fig). This difference in miscalls and concordance between programs may be further explained by the variation in database sizes, since the PubMLST schemes used for mlst have fewer alleles across all seven loci which results in fewer STs compared to stringMLST as shown on S3 Table.

Previously, the scalability of ProkEvo was assessed by a comparative analysis of its computational performance on Crane and OSG, using two datasets with 2,392 and 23,045 genomes each (10 X difference), and the standard mlst approach for ST calling [21]. To further demonstrate the gain in computational runtime obtained with the use of stringMLST within ProkEvo, the complete *S*. Typhimurium dataset containing 23,045 genomes was run on OSG. While ProkEvo with mlst finished all ST calls in 26 days and 6 hours when OSG was used as a computational platform [21], ProkEvo with stringMLST completed the task in 3 days and 6 hours. Altogether, stringMLST provides fine-tunable and rapid alternative to mlst for scalable ST genotyping that is portable to be implemented in any high-performance and high-throughput platform, with its use being further facilitated by its implementation in ProkEvo.

## Discussion

The incorporation of WGS technology has advanced the study of bacterial populations, enabling new strategies of inferring traits associated with populations at different scales of resolution that may be contributing to important ecological characteristics and/or epidemiological patterns [21][32][33][34][35][36][37][38]. In particular, the use of a hierarchical population structure analysis permits inference into ancestral vs. recent evolutionary relationships and consequently, the patterns of genomic and phenotypic diversification among genotypic units can be studied within these evolutionary contexts [21][39][40][41][12][42][43].

ST-based classification is an integral part of the hierarchical genotyping approach [21][27]. ST lineages are inferred from the patterns of allelic variation at a relatively small (typically seven) set of conserved loci [8][9][44] and remain a very useful context in which to study bacterial populations. However, as the field transitions to WGS data, the ability to infer STs is bottlenecked by the computationally intensive step of genome assembly. Thus, there is a great need for algorithms that can efficiently infer STs from large WGS-based datasets. [21][27][8][9][44]There are multiple tools available for MLST classification, such as mlst [22], ARIBA **Error! Reference source not found.**, stringMLST [17], MentaLiST **Error! Reference source not found.**, STing **Error! Reference source not found.**. In general, the available tools can be categorized based on the input data they use - some tools use raw Illumina paired-end sequence data, while others use *de novo* assemblies [16]. Using raw sequence data for ST-based classification has a tremendous advantage especially in pathogen surveillance, since all the computationally demanding steps prior to the *de novo* assembly are bypassed and the STs calls are made as the sequence reads are generated. Specifically, mlst uses *de novo* genome assemblies as an input and performs mapping in order to align sequences to pre-downloaded allelic files across all target loci. ARIBA identifies AMR-associated genes, single nucleotide polymorphisms, and ST calls using Illumina paired-end raw sequencing reads. ARIBA clusters the raw reads by mapping them to genes, and then performs local assembly within clusters to identify AMR genes and ST calls. On the other hand, stringMLST and MentaLiST rely on kmer matching between raw sequence reads and available ST schemes that allows for fast mapping and ST-based typing. In a narrow setting with few bacterial species, both tools are shown to be accurate and fast for standard MLST classification, while providing comparable accuracy with MentaLiST albeit using less computational resources **Error! Reference source not found.**. STing is the successor of stringMLST - it uses the same algorithmic approach with additional computational applications for large MLST schemes such as ribosomal MLST (rMLST) and core-genome MLST (cgMLST) **Error! Reference source not found.**. All these tools have integrated ST schemes and/or provide utilities for downloading the available PubMLST databases. There are a few available comparisons of assembly-free tools for ST classification, and the published studies are primarily focused on the computational resources used and the percentage of correctly classified STs in relatively small data sets [16][63][64]. When tools were tested with real outbreak datasets (*L. monocytogenes*, *E. coli*, *C. jejuni*, *S. enterica*) comprising 85 samples, stringMLST showed the fastest running time of 80.8 minutes and high accuracy in ST calls (100%) [16]. While MentaLiST does not scale well when reads with high coverage are used, it performs well on MLST schemes with up to a few thousand genes and alleles, such as cgMLST (∼3,000 genes) [63]. While most ST tools perform satisfactorily, there are some relevant bottlenecks to be considered. For example, some tools use out-of-date MLST databases that require manual curation, and can directly affect the accuracy of ST calls, especially when mixed and low coverage samples are used [16]. ST tools that are assembly and alignment free, such as stringMLST, STing and MentaLiST, show quite a few advantages in term of accuracy and efficiency that make them applicable for real-time molecular epidemiology and surveillance. Thus, we chose stringMLST as a representative of the kmer-based ST tools to perform a systems-based comparative analysis that assess the computational and statistical efficacy of ST calls across divergent pathogens in contrast to the legacy MLST approach.

As shown in our studies, the accuracy of stringMLST is affected by the species being tested without any specific phylogenetic patterns. In particular, the choice of kmer length used directly impacts the proportion of ST miscalls across species, and in certain cases it may not be applied as designed even after parameter tunning. It is likely that the varying accuracy reflects the different levels of population structuring, patterns of genome diversification, degree of repetitive sequences, degree of horizontal gene transfer (HGT), and relative abundances of mobile elements such as prophages and insertion sequences, etc. [12][41][68][69][70][71][72][73]. A clear example in our study was *S*. *enterica*, for which the accuracy of stringMLST at different kmer lengths varied across ecologically distinct serovars that are known to have unique pan-genomic composition such as prophage content [12][71][74][75]. To the best of our knowledge, the currently available comparisons between ST tools have not considered any systematic approach for parameter tunning across phylogenetic divergent species known to vary in population structure [21][27][67].

In evaluating genome-intrinsic and -extrinsic variables that could contribute to differences in accuracy between mlst and stringMLST, we found that species level variation in accuracy was mostly explained by the uniqueness of their genomic composition and number of contigs per genome. As genomic composition is an inheritable property of the bacterial species, subtype, and evolutionary history of specific populations, the association with algorithmic performance was somewhat expected [46][47]. However, the contribution of the number of contigs to performance implies that data from long-read sequencing platforms such as PacBio and Oxford Nanopore Technologies (ONT) may have a considerable effect on accuracy of programs such as stringMLST because these technologies produce reads with lower accuracy (∼80-90%) that may inflate the number of false allelic calls at MLST loci, with the effect of artificially splitting major STs into multiple sub-populations [48]**Error! Reference source not found.Error! Reference source not found.**. Therefore, while more work is needed in this field, current studies using hybrid assembly approaches of both Illumina short reads and ONT long reads **Error! Reference source not found.**, as well as only polished ONT reads **Error! Reference source not found.** for performing ST-based classification may lead to promising, cost-effective results from these types of platforms. Hence, we expect that a combination of hybrid sequencing strategies with species and even subtype-specific tuning of programs such as stringMLST and STing, will facilitate real-time surveillance, prediction of STs and prediction of traits such as AMR that are found to associated with specific STs [14][49].

While the kmer length of 35 is currently recommended as a default value of stringMLST, our systems-based approach demonstrated that for specific bacterial species it will result in increasing the frequency of ST miscalls which in turn may hinder epidemiological investigations. Across phylogenetic divergent pathogenic bacterial species, the optimal kmer length ranged from 20 to 140, regardless of their ancestral relationship or speciation pattern. The varying population structure, the pattern of genome diversification and architecture (e.g., impact of HGT), as well as sequence coverage may be some of the reasons underlying the observed statistics [12][41][68][69][70][71][72][73]. Although we hypothesize that longer sequence reads will help overcome this limitation, there is still a context-dependent consideration for parameter tunning and overall algorithmic implementation. Therefore, in the case of stringMLST, we suggest the following actionable strategies to maximize its utilization, including: i) developers to consider implementing a pre-step that heuristically searches for the optimal kmer length (minimizes ST miscalls) in dataset-dependent fashion (sampling from the testing data), perhaps even by comparing with the standard MLST program as positive controls; and/or ii) researchers to run wide range of kmer lengths on a subset of the dataset in order to select the optimal kmer length that minimizes the percentage of ST miscalls. Given the speed and scalability of stringMLST, using multiple kmer lengths is not likely to add much overhead to the analyses, and this provides an empirical statistical approach for kmer selection and optimization of ST classifications. With this data-driven fine-tunning of the kmer length, stringMLST is a powerful program that can be efficiently and effectively used in microbiological and epidemiological laboratories.

We recently developed ProkEvo, a freely available scalable platform for performing hierarchical-based bacterial population genomics analyses [21]. ProkEvo: 1) uses the Pegasus Workflow Management System to ensure reproducibility, scalability, and modularity; 2) uses high-performance and high-throughput computational platforms; 3) automates and scales multitude of computational analyses of a few to tens of thousands of bacterial genomes; 4) can run many thousands of analyses concurrently if the computational resources are available; 5) is easily modifiable and expandable platform that can incorporate additional algorithmic steps and custom scripts. The initial implementation of ST-based classifications through ProkEvo, as part of a hierarchical genotyping strategy to map and track populations, was done using the assembly-dependent MLST program [21][22]. Running mlst inside ProkEvo allows for parallelization of the genome assemblies (run per isolate or genome) which enhances scalability and facilitates the optimal use of computation resources. Theoretically, if there are *n* isolates and *n* cores available on the computational platform, ProkEvo can linearly utilize all resources and run all *n* independent tasks simultaneously. Typically, ST-based classifications are time consuming because the mapping process is run sequentially in a set of genomes instead of running them independently. Thus, using modular and distributed platforms such as ProkEvo for performing ST-based genotyping provides great benefit, especially if additional features such as other hierarchical genotypes and pan-genomic mapping tools are part of the same platform [21]. As part of this work, we modified ProkEvo to not only offer the standard assembly-dependent MLST mapping approach, but it now contains stringMLST, and our tests showed a significant speed-up in runtime for datasets ranging from a few hundreds to tens of thousands of genomes. Moreover, we additionally added STing to ProkEvo, as an efficient successor of stringMLST (https://github.com/npavlovikj/MLST_stringMLST_analyses/tree/main/Prokevo_STing). To use ProkEvo with stringMLST or STing, the researcher only needs to provide a list of SRA identifications and run the submit script without any advanced experience in high-performance or high-throughput computing. Depending on the configuration set, ProkEvo can use locally downloaded sequence data or download the data from NCBI directly. The Pegasus Workflow Managements System that is used by ProkEvo automatically handles the dependencies, as well as all the intermediate and final files. Thus, using platforms such as ProkEvo with fast tool for hierarchical genotyping, such as stringMLST, allows for robust and efficient population-based genomics analyses that facilitate: i) mapping and tracking of variants or lineages for epidemiological inquiries; ii) population structure analysis; and iii) ecological trait prediction using pan-genomic mapping to specific genotypes.

## Conclusion

In conclusion, stringMLST largely proved to be an accurate, rapid, and scalable tool for ST-based classifications that could be deployed in microbiological laboratories and epidemiological agencies. However, our work clearly illustrates the need to optimize stringMLST across phylogenetic divergent species and populations of bacterial pathogens. Based on our results, we propose that the kmer length should be optimized in two ways on a case-by-case basis: 1) intrinsically by implementing a pre-step inside the algorithm to sample from the target data and select the optimal kmer length; or 2) by the user through a heuristic data mining approach to select the optimal kmer length prior to finalizing the ST calls. Also, by assessing genome-intrinsic and -extrinsic factors that could affect the stringMLST performance, our work suggests that longer sequence reads have the potential to improve its accuracy for specific bacterial species. Furthermore, the integration of stringMLST into ProkEvo allows users to take advantage of other hierarchical genotyping strategies, including pan-genomic mapping, which reproducibly facilitates ecological and epidemiological inquiries at scale. Ultimately, this work emphasizes the importance of developing robust algorithmic tools for mining WGS data that can have direct implications for mapping and tracking of bacterial populations.

## Acknowledgements

This work was completed by utilizing the Holland Computing Center of the University of Nebraska, which receives support from the Nebraska Research Initiative, and using resources provided by the Open Science Grid, which is supported by the National Science Foundation and the U.S. Department of Energy’s Office of Science. This research used the Pegasus Workflow Management Software funded by the National Science Foundation under grant #1664162. This publication made use of the PubMLST website (https://pubmlst.org/) developed by Keith Jolley (Jolley & Maiden 2010, BMC Bioinformatics, 11:595) and sited at the University of Oxford. The development of that website was funded by the Wellcome Trust. We would like to greatly thank Mats Rynge for his extensive assistance and valuable suggestions while setting up and running ProkEvo on the Open Science Grid. We also thank Dr. Derek Weitzel and Karan Vahi for their technical support.

