## Supplementary materials for "Systems-based approach for optimization of a scalable bacterial ST mapping assembly-free algorithm"

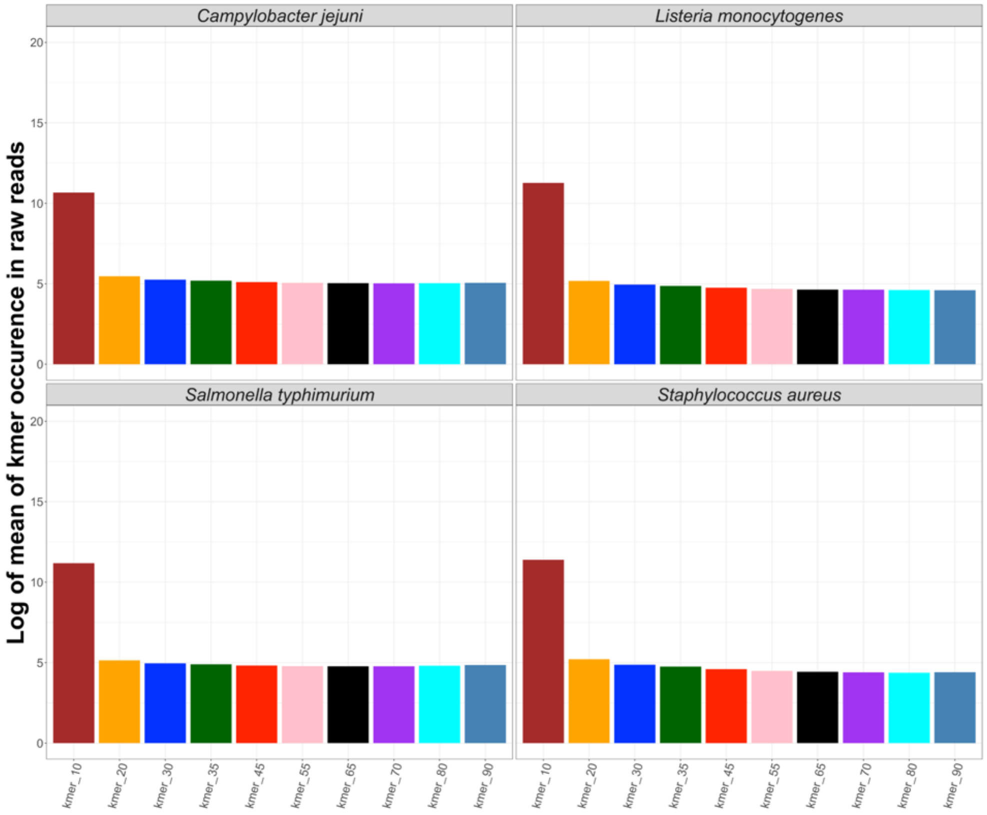

Figure S1. Distribution of kmers across raw reads per bacterial species.
Random 100 raw paired-end reads from the initial *C. jejuni*, *L. monocytogenes*, *S. aureus* and *S.* Typhimurium (major representative zoonotic serovar of *S. enterica*) datasets were selected, and DSK was used to count the occurrence of kmers of lengths 10, 20, 30, 35, 45, 55, 65, 70, 80 and 90, respectively, in the raw reads. The mean for kmer occurrence per organism and kmer length was calculated, and all values were ultimately transformed to natural logarithm (base *e*) for the final visualization.

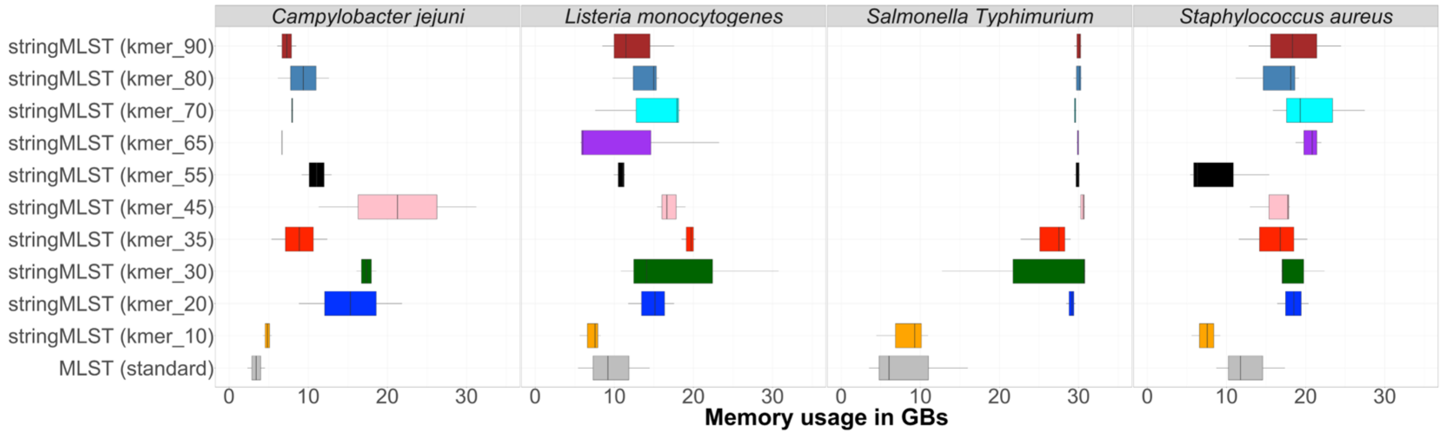

Figure S2. Comparison of the maximum computational memory used in GBs by mlst and stringMLST for ST-based classification of genomes across four bacterial species.
In order to compare the memory used between mlst and stringMLST with different kmer lengths, we chose four different datasets, including four phylogenetic divergent bacterial species (pathogens): *C. jejuni*, *L. monocytogenes*, one major serovar of *S. enterica* (*S*. Typhimurium) and *S. aureus*, with 600 genomes each. These randomly selected 600 genomes were randomly split into three batches with 200 genomes each. We then ran mlst with all required steps, such as quality trimming and adapter clipping, *de novo* assembly and assembly discarding, on each batch and dataset (three datapoints per bacterial species). Separately, we ran stringMLST with a range of 10 different kmer values (10, 20, 30, 35, 45, 55, 65, 70, 80, 90) on each dataset. For each organism, the memory was calculated as the maximum of all 200 genomes per batch. In the case of mlst, the recorded memory was the maximum memory of all the steps ran prior to mlst, such as trimming, *de novo* assembly, quality checking, filtering, and ST typing. The memory was calculated using the "cgget" command (*cgget -r memory.usage_in_bytes/slurm/uid_${UID}/job_${SLURM_JOBID}/*) part of the Linux Control Groups (cgroup).

A.

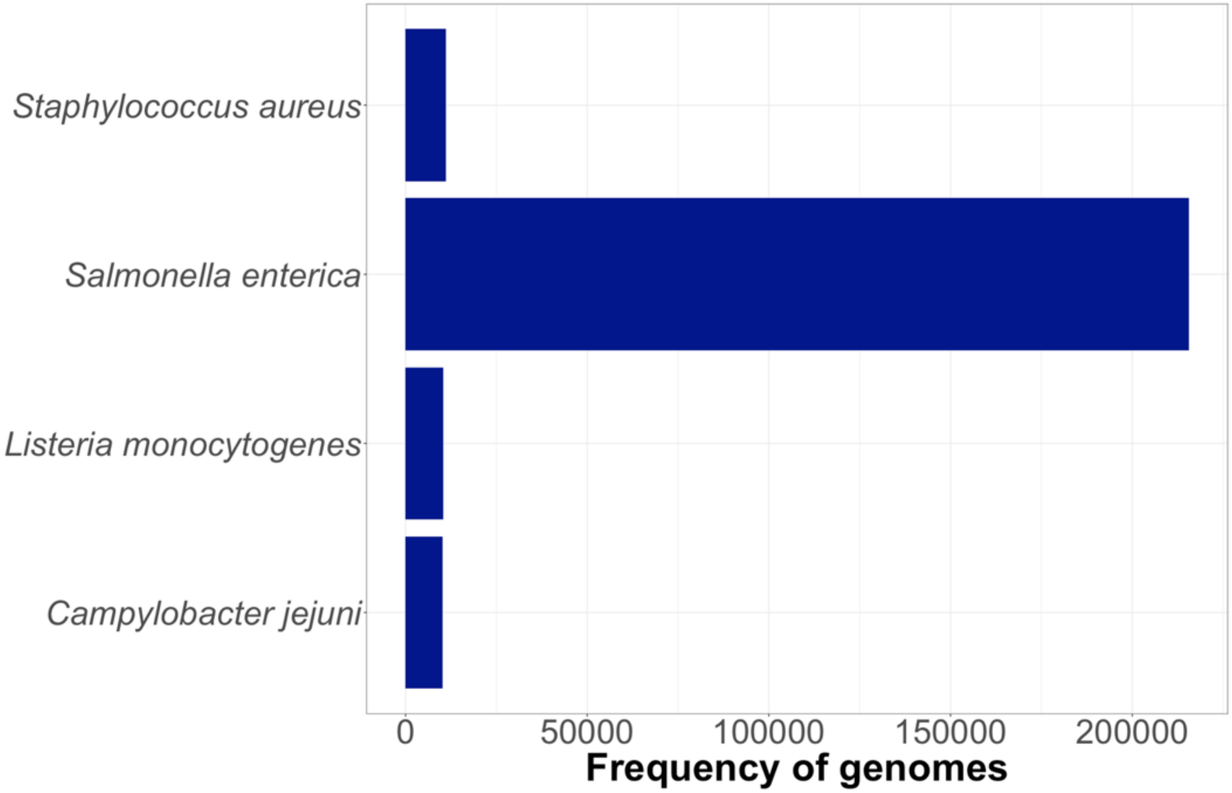
Staphylococcus aureus
Salmonella enterica
Listeria monocytogenes
Campylobacter jejuni
0
50000
100000
150000
200000
Frequency of genomes

**B.**

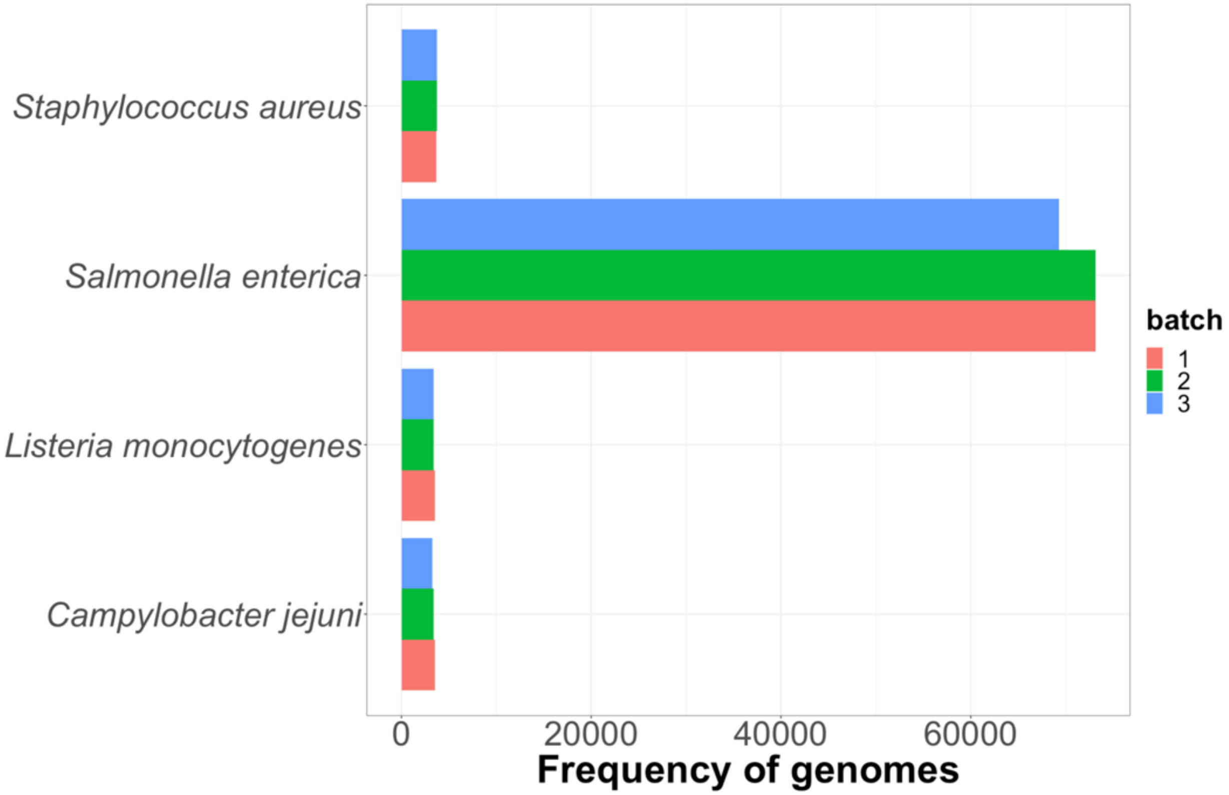
Staphylococcus aureus
Salmonella enterica
Listeria monocytogenes
Campylobacter jejuni
0
20000
40000
60000
Frequency of genomes
batch
1
2
3

C.

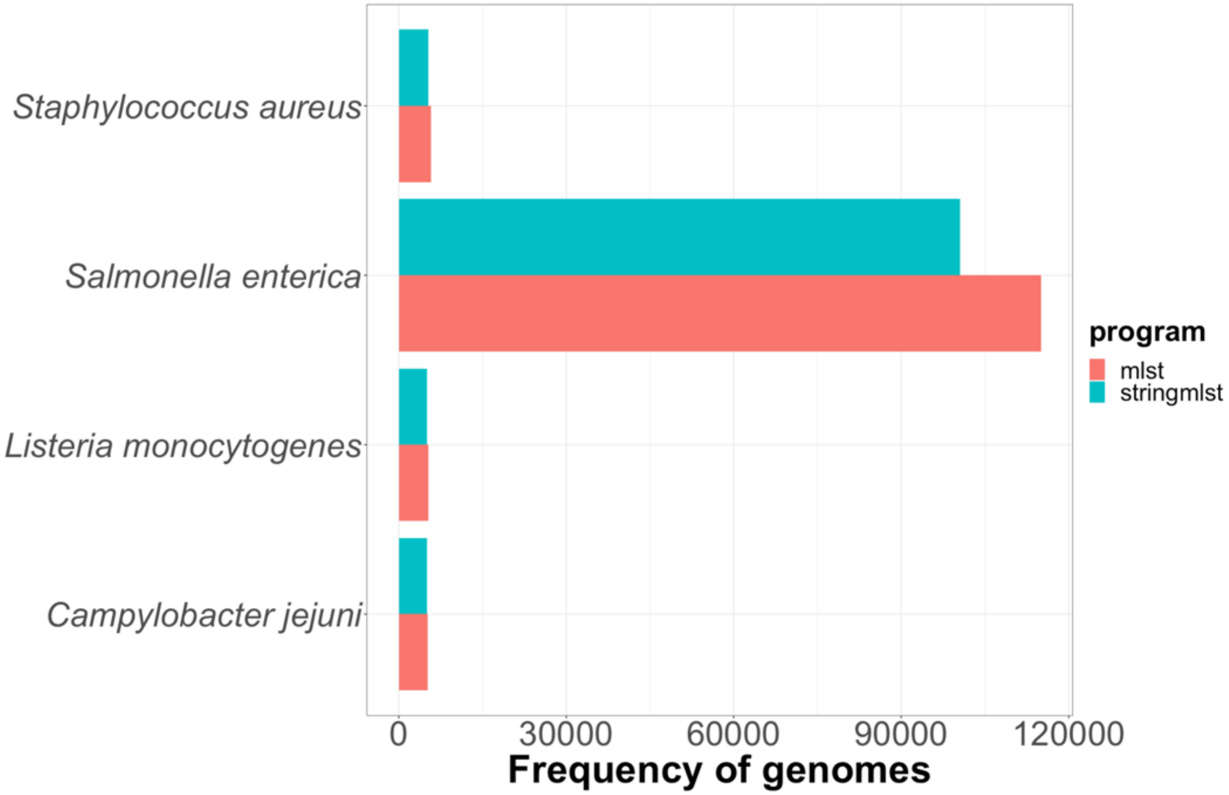
Staphylococcus aureus
Salmonella enterica
Listeria monocytogenes
Campylobacter jejuni
0
30000
60000
90000
120000
Frequency of genomes
program
mlst
stringmlst

D.

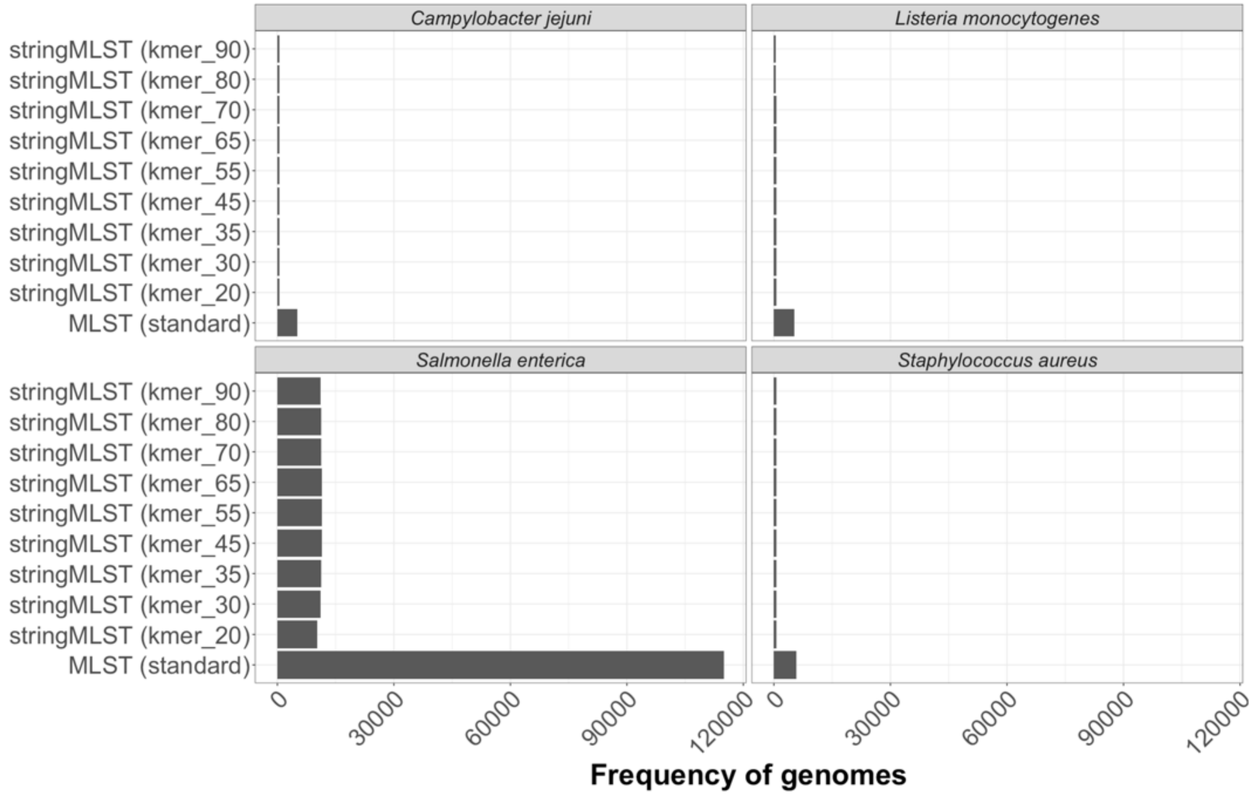

Campylobacter jejuni
Listeria monocytogenes
Salmonella enterica
Staphylococcus aureus
stringMLST (kmer_90)
stringMLST (kmer_80)
stringMLST (kmer_70)
stringMLST (kmer_65)
stringMLST (kmer_55)
stringMLST (kmer_45)
stringMLST (kmer_35)
stringMLST (kmer_30)
stringMLST (kmer_20)
MLST (standard)
0
30000
60000
90000
120000
Frequency of genomes

**E.**

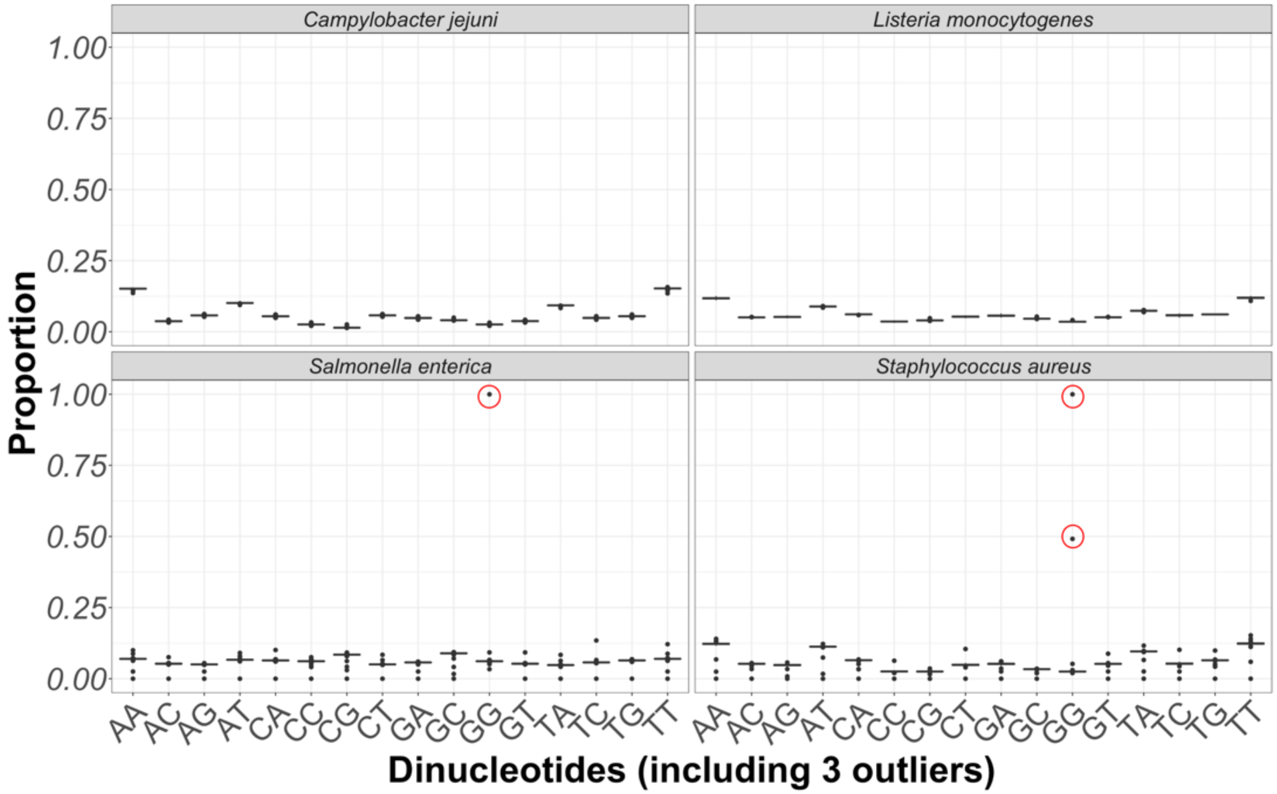

Campylobacter jejuni
Listeria monocytogenes
Salmonella enterica
Staphylococcus aureus
Proportion
1.00
0.75
0.50
0.25
0.00
AA AC AG AT CA CC CG CT GA GC GG GT TA TC TG TT
Dinucleotides (including 3 outliers)

F.

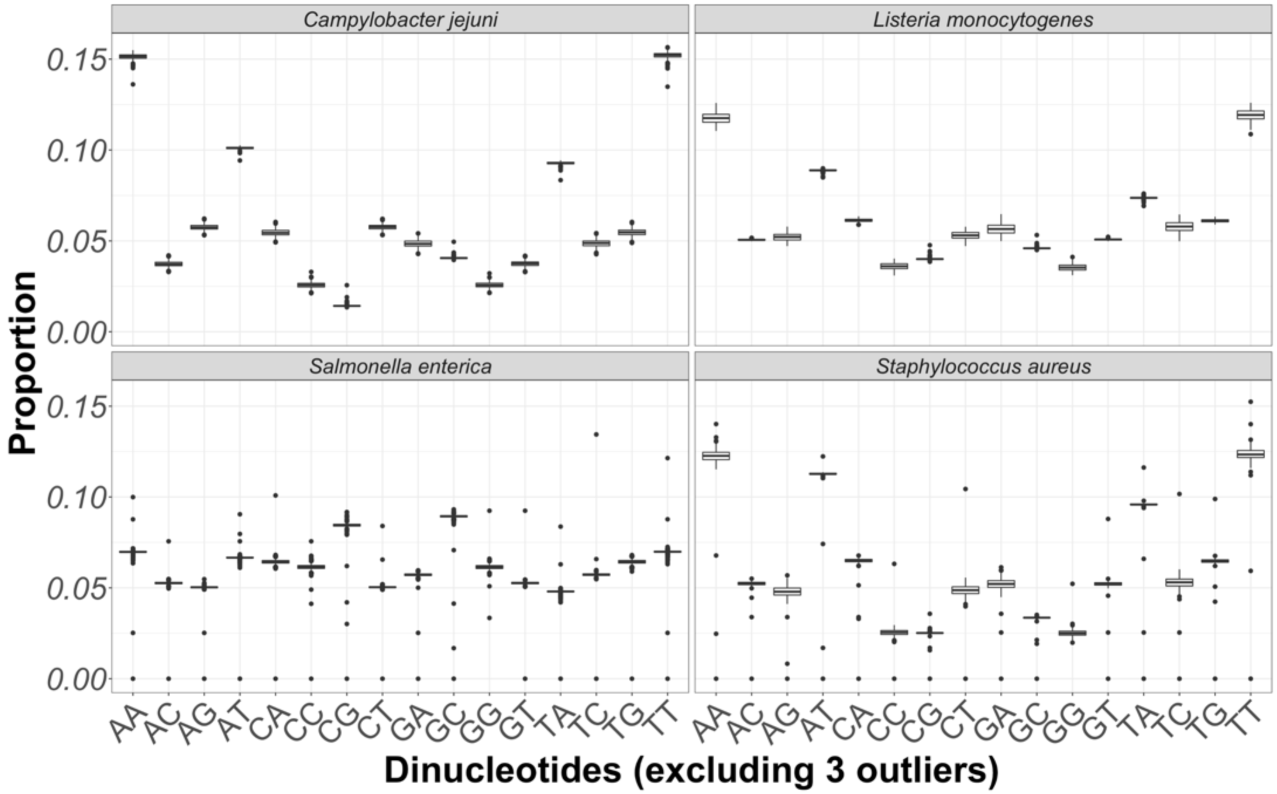
Campylobacter jejuni
Listeria monocytogenes
Salmonella enterica
Staphylococcus aureus
Proportion
0.15
0.10
0.05
0.00
AA AC AG AT CA CC CG CT GA GC GG GT TA TC TG TT
Dinucleotides (excluding 3 outliers)

G.

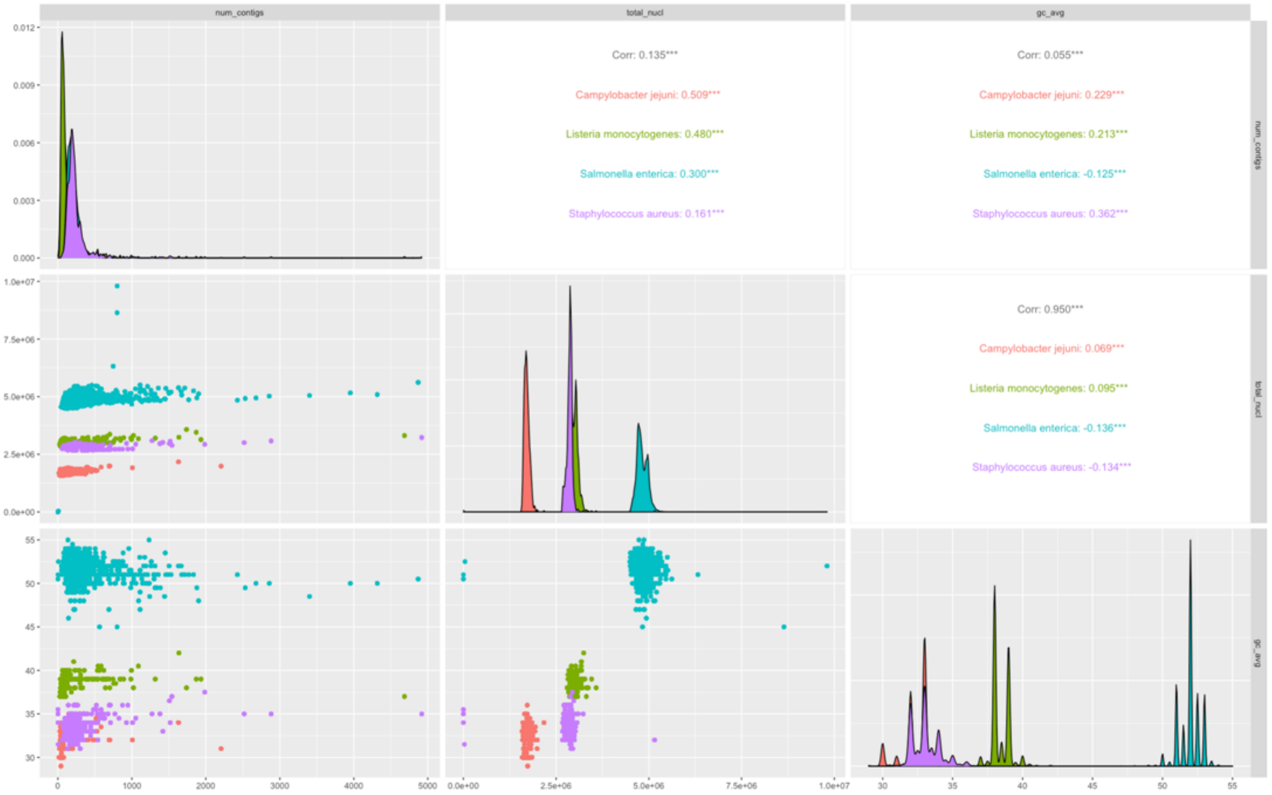

Figure S3. Summary statistics of the frequency of genomes, including the distribution of dinucleotides and bivariate associations between genome-intrinsic variables across all four bacterial species.
Frequency-based distribution of randomly selected genomes across bacterial species (A), including a stratification by batch (B), program (C), and further differentiation by kmer length used by stringMLST (D). Proportion of all sixteen pairs of dinucleotides present in a bacterial genome, across species, with (E) or without outliers (red-circled data points) (F). (G) Bivariate association between genome-intrinsic variables across species with statistical significance measured by the Pearson correlation coefficient (Corr). Genome-intrinsic variables used were the total number of contigs (num_contigs), the total number of nucleotides per genome (assembly), and the GC% content per genome (gc_avg). (G) Asterisks refer to the degree of significance for the correlation coefficient, with *p*-value

thresholds being: *$p < 0.05$, **$p \leq 0.01$, ***$p \leq 0.001$, ****$p \leq 0.0001$ and NS = not significant at $p \geq 0.05$. Across all figures A-G, data for *S. enterica* Subspecies *enterica* (*S. enterica*) contained an even proportion of genomes across twenty serovars.

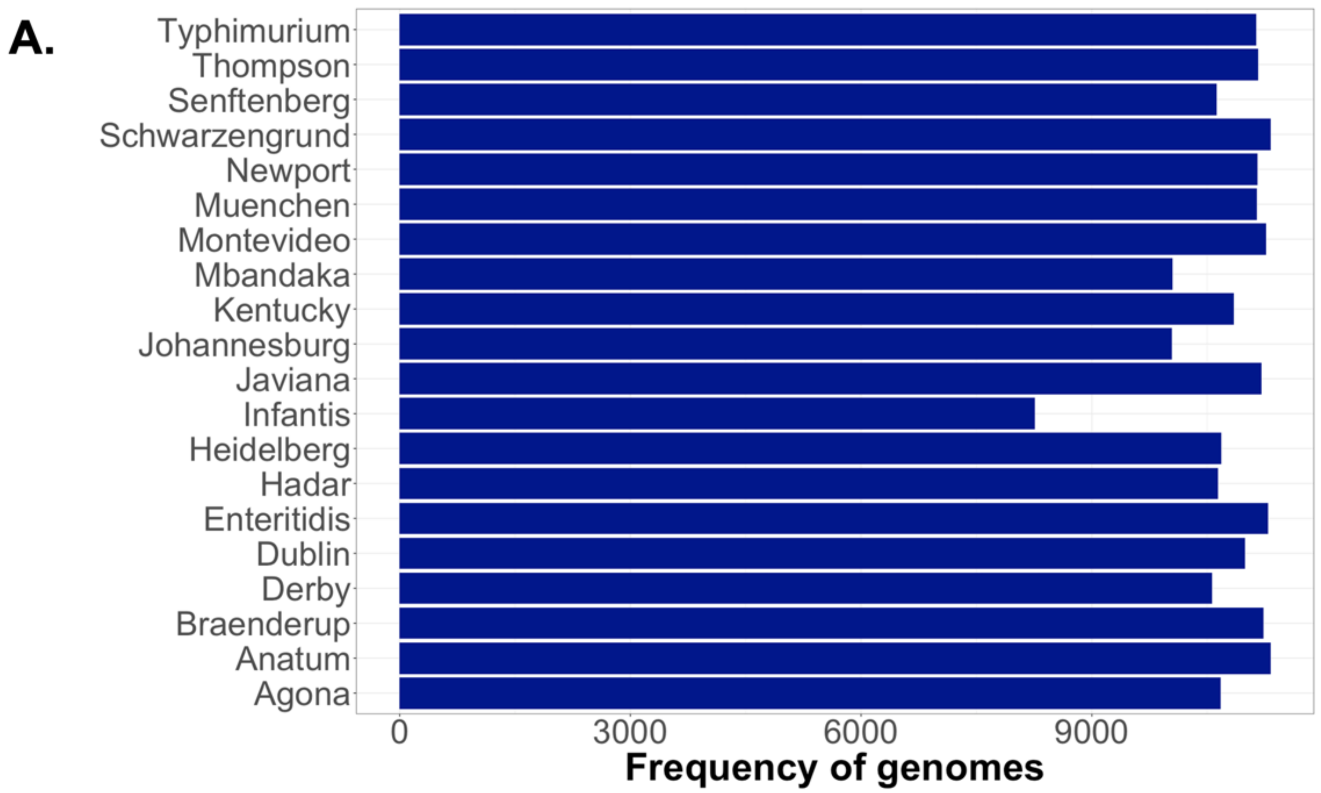
A.
Typhimurium
Thompson
Senftenberg
Schwarzengrund
Newport
Muenchen
Montevideo
Mbandaka
Kentucky
Johannesburg
Javiana
Infantis
Heidelberg
Hadar
Enteritidis
Dublin
Derby
Braenderup
Anatum
Agona
0
3000
6000
9000
Frequency of genomes

B.

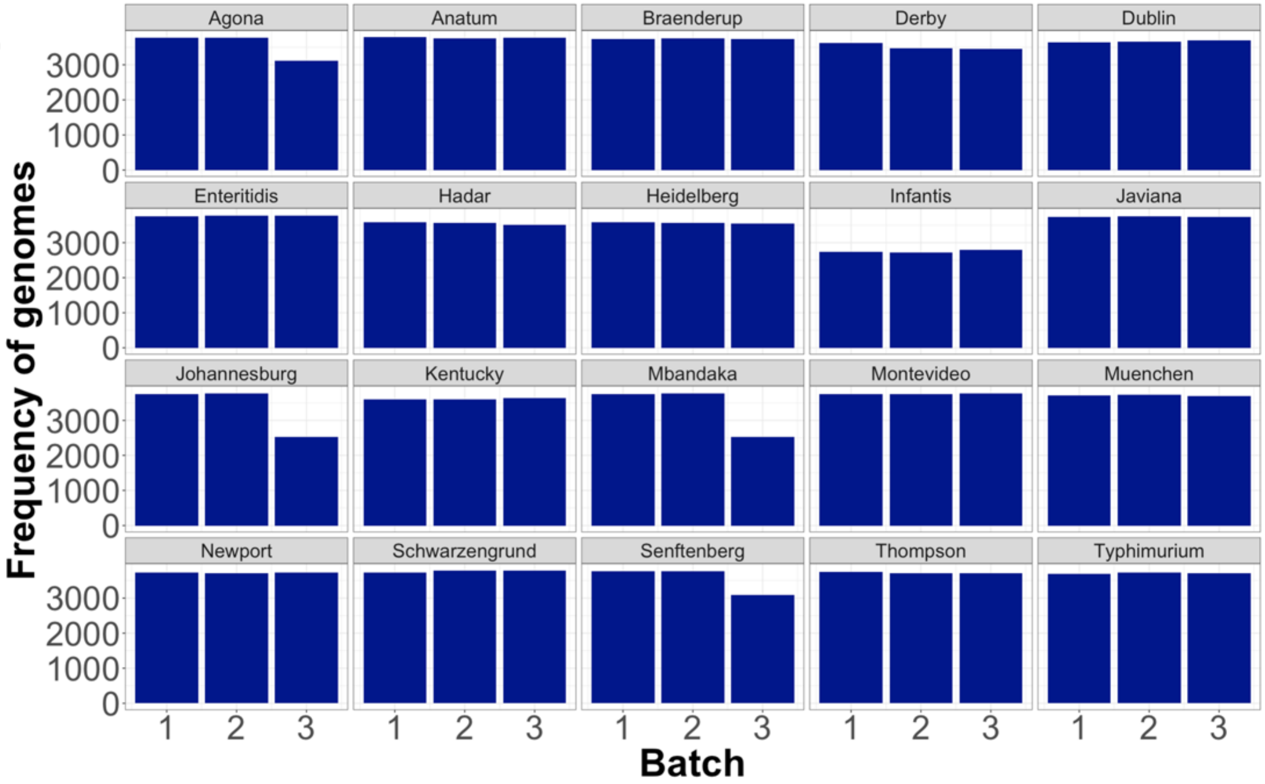
Agona
Anatum
Braenderup
Derby
Dublin
Enteritidis
Hadar
Heidelberg
Infantis
Javiana
Johannesburg
Kentucky
Mbandaka
Montevideo
Muenchen
Newport
Schwarzengrund
Senftenberg
Thompson
Typhimurium
Frequency of genomes
0
1000
2000
3000
1
2
3
Batch

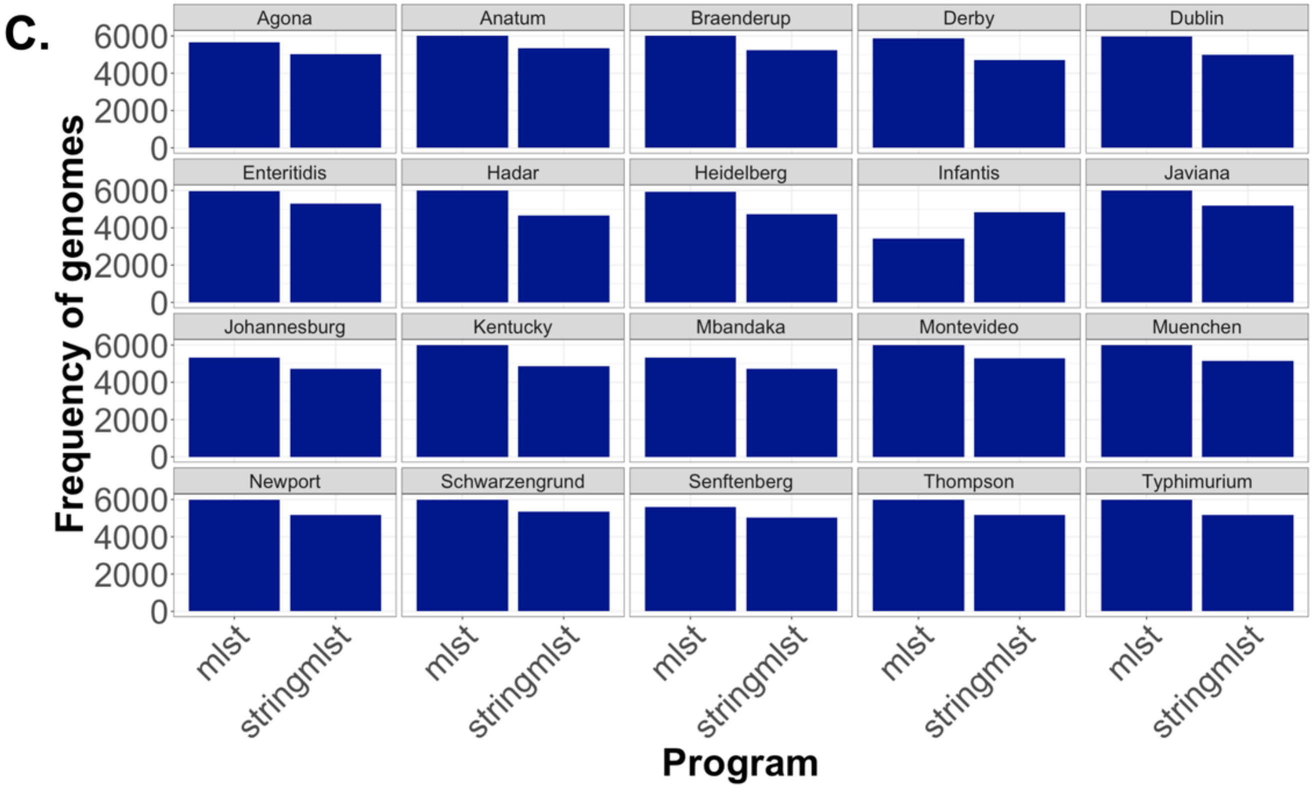
C.
Frequency of genomes
Program
6000
4000
2000
0
Agona
Anatum
Braenderup
Derby
Dublin
Enteritidis
Hadar
Heidelberg
Infantis
Javiana
Johannesburg
Kentucky
Mbandaka
Montevideo
Muenchen
Newport
Schwarzengrund
Senftenberg
Thompson
Typhimurium
mlst
stringmlst

D.

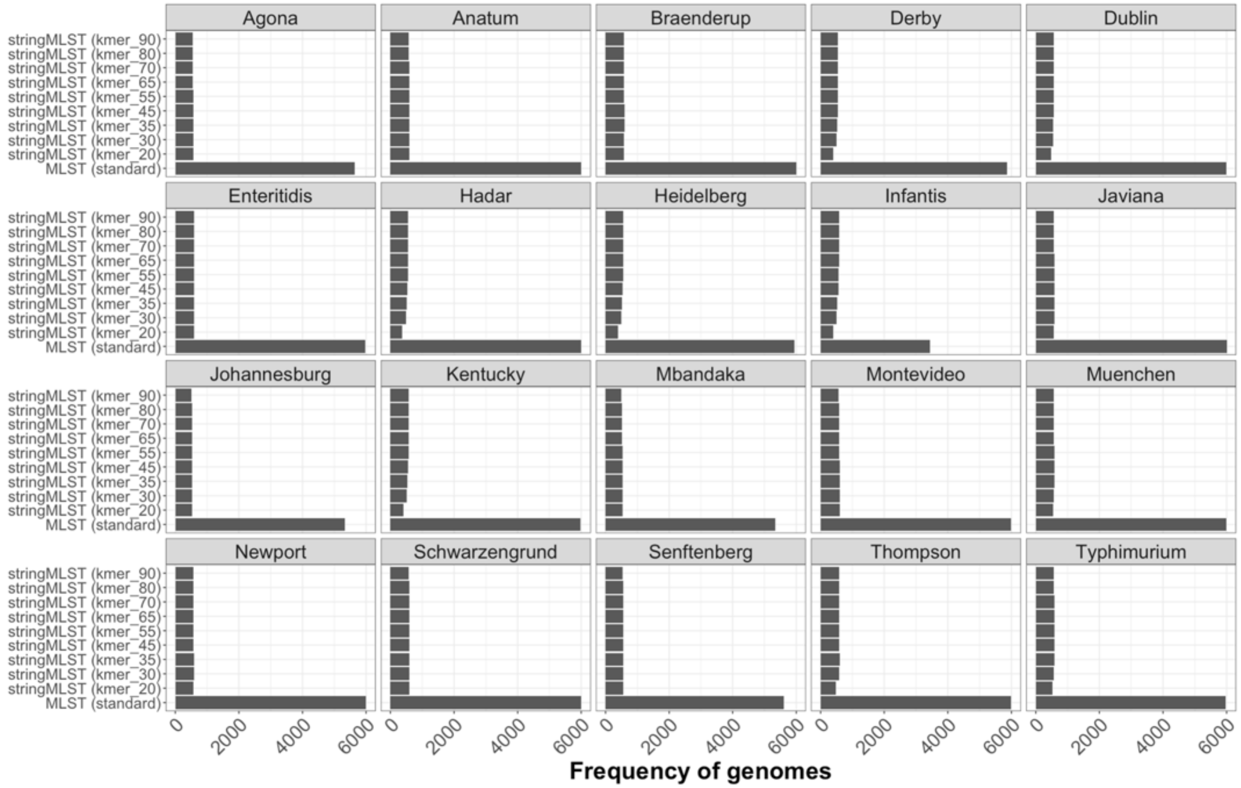

Agona
Anatum
Braenderup
Derby
Dublin
Enteritidis
Hadar
Heidelberg
Infantis
Javiana
Johannesburg
Kentucky
Mbandaka
Montevideo
Muenchen
Newport
Schwarzengrund
Senftenberg
Thompson
Typhimurium
stringMLST (kmer_90)
stringMLST (kmer_80)
stringMLST (kmer_70)
stringMLST (kmer_65)
stringMLST (kmer_55)
stringMLST (kmer_45)
stringMLST (kmer_35)
stringMLST (kmer_30)
stringMLST (kmer_20)
MLST (standard)
0
2000
4000
6000
Frequency of genomes

E.

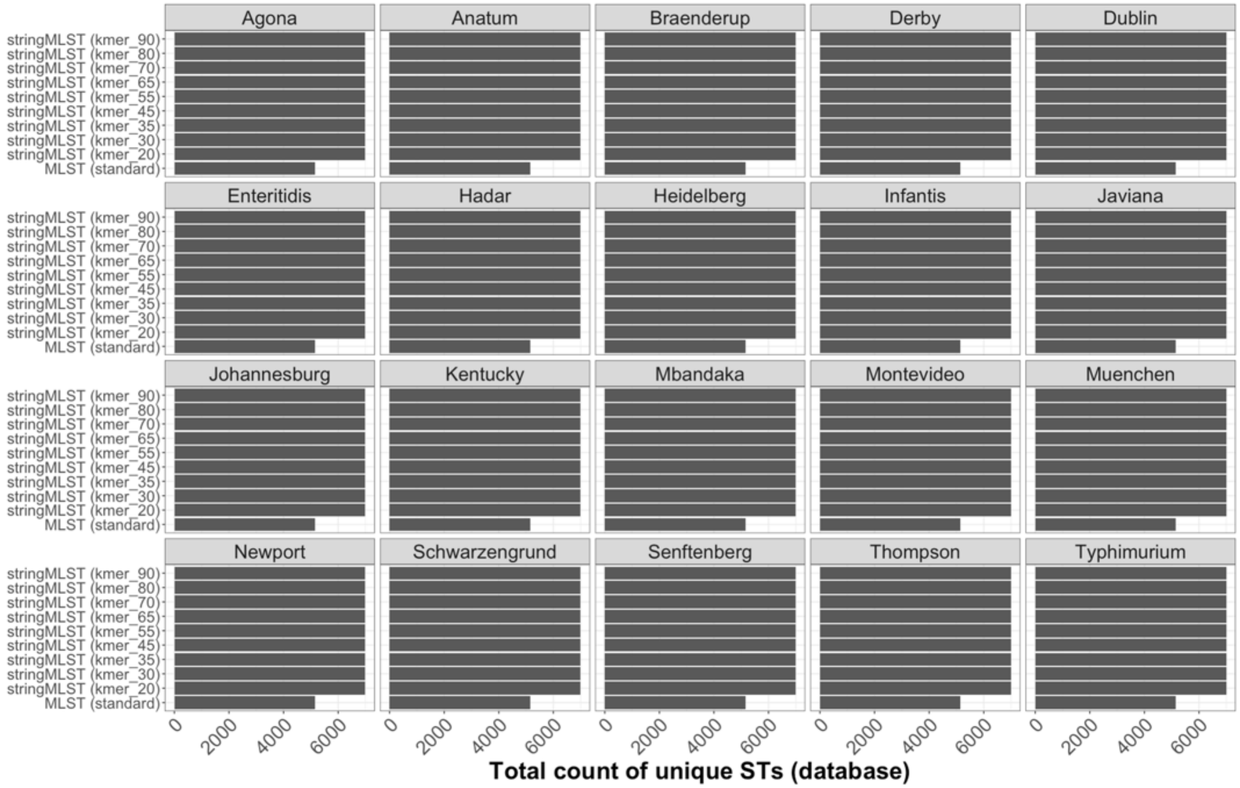

Agona
Anatum
Braenderup
Derby
Dublin
Enteritidis
Hadar
Heidelberg
Infantis
Javiana
Johannesburg
Kentucky
Mbandaka
Montevideo
Muenchen
Newport
Schwarzengrund
Senftenberg
Thompson
Typhimurium
stringMLST (kmer_90)
stringMLST (kmer_80)
stringMLST (kmer_70)
stringMLST (kmer_65)
stringMLST (kmer_55)
stringMLST (kmer_45)
stringMLST (kmer_35)
stringMLST (kmer_30)
stringMLST (kmer_20)
MLST (standard)
0
2000
4000
6000
Total count of unique STs (database)

F.

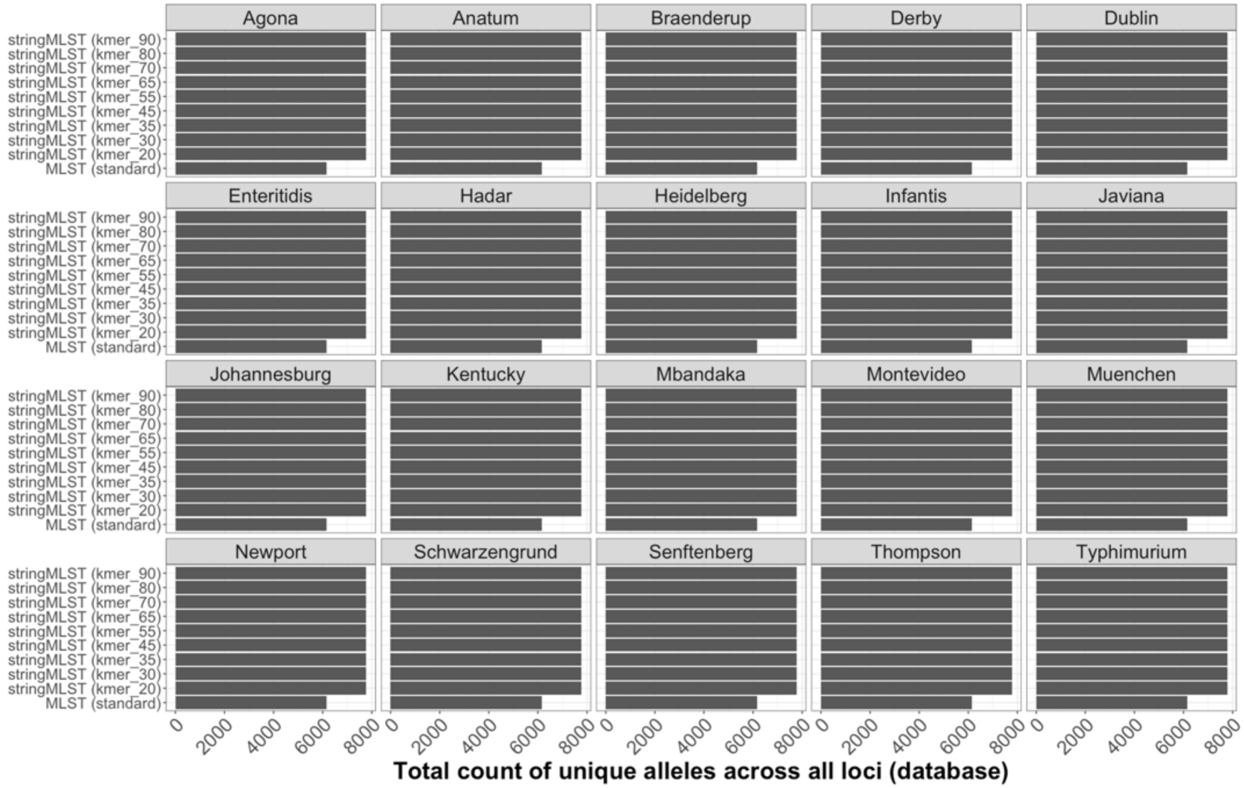
Agona
Anatum
Braenderup
Derby
Dublin
Enteritidis
Hadar
Heidelberg
Infantis
Javiana
Johannesburg
Kentucky
Mbandaka
Montevideo
Muenchen
Newport
Schwarzengrund
Senftenberg
Thompson
Typhimurium
stringMLST (kmer_90)
stringMLST (kmer_80)
stringMLST (kmer_70)
stringMLST (kmer_65)
stringMLST (kmer_55)
stringMLST (kmer_45)
stringMLST (kmer_35)
stringMLST (kmer_30)
stringMLST (kmer_20)
MLST (standard)
0
2000
4000
6000
8000
Total count of unique alleles across all loci (database)

G.

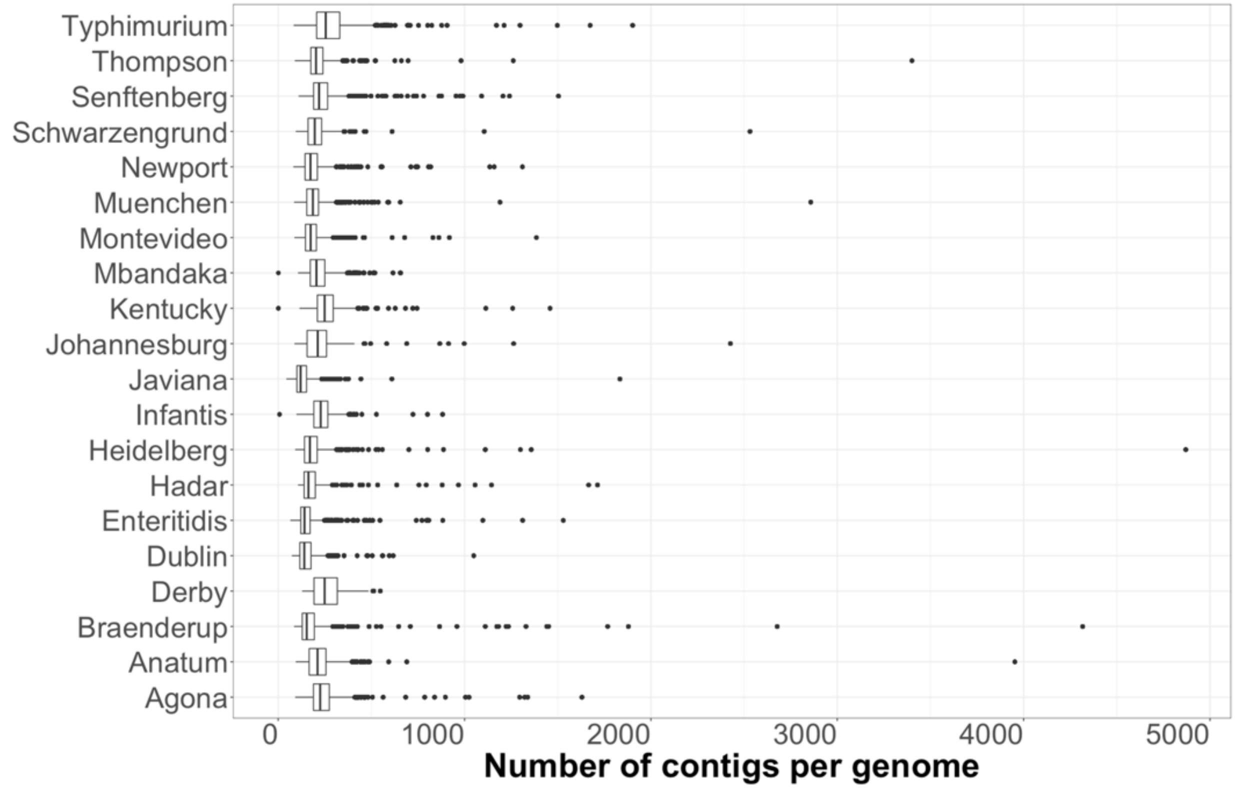
Typhimurium
Thompson
Senftenberg
Schwarzengrund
Newport
Muenchen
Montevideo
Mbandaka
Kentucky
Johannesburg
Javiana
Infantis
Heidelberg
Hadar
Enteritidis
Dublin
Derby
Braenderup
Anatum
Agona
0
1000
2000
3000
4000
5000
Number of contigs per genome

H.

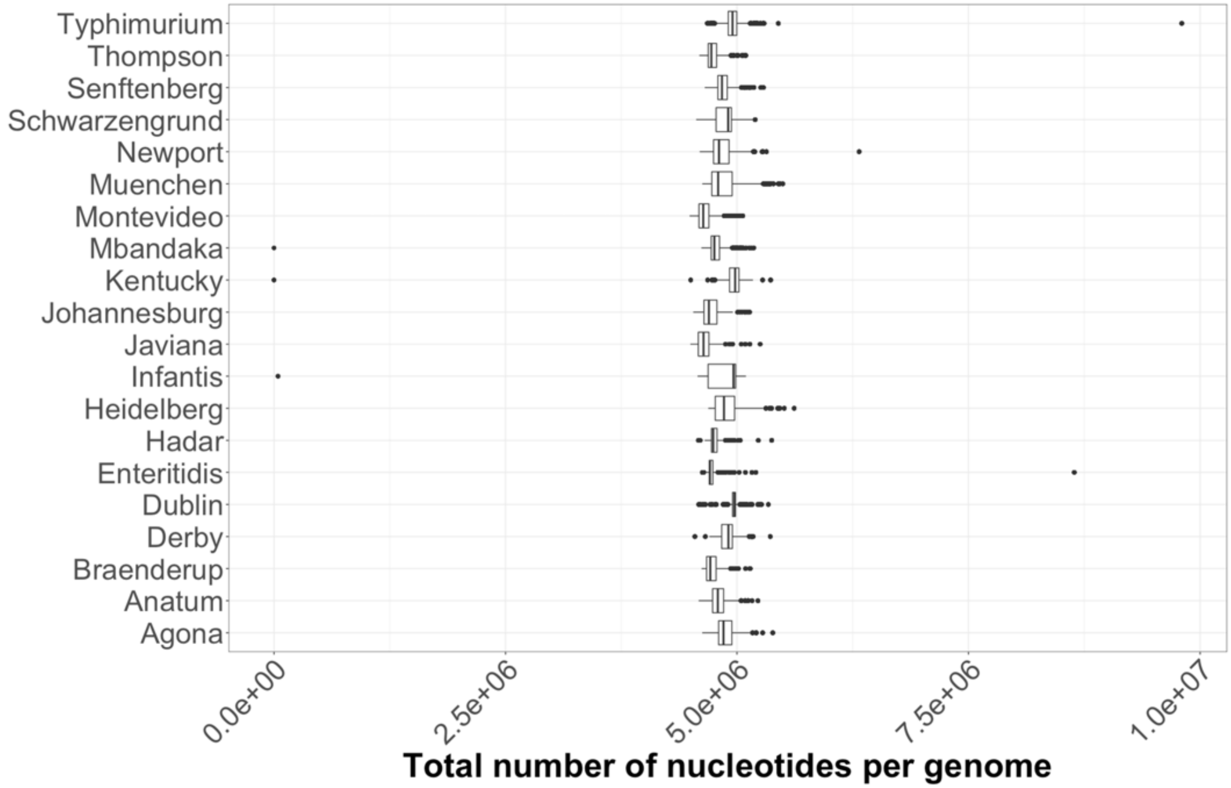
Typhimurium
Thompson
Senftenberg
Schwarzengrund
Newport
Muenchen
Montevideo
Mbandaka
Kentucky
Johannesburg
Javiana
Infantis
Heidelberg
Hadar
Enteritidis
Dublin
Derby
Braenderup
Anatum
Agona
0.0e+00
2.5e+06
5.0e+06
7.5e+06
1.0e+07
Total number of nucleotides per genome

**I.**

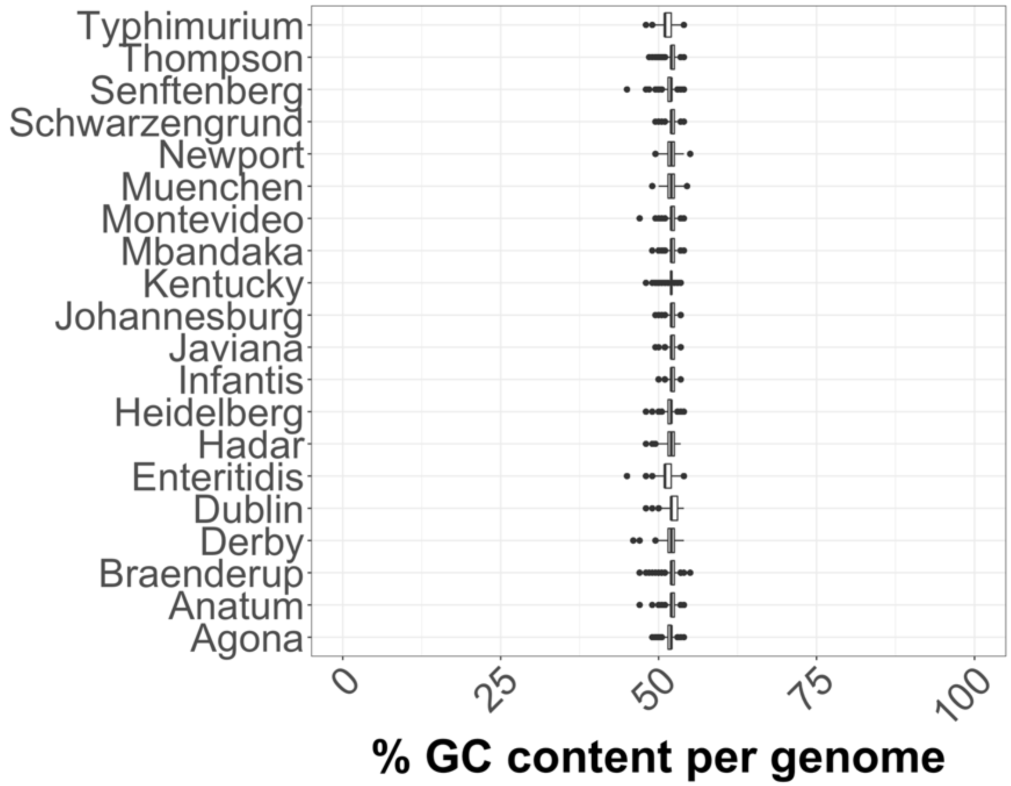
Typhimurium
Thompson
Senftenberg
Schwarzengrund
Newport
Muenchen
Montevideo
Mbandaka
Kentucky
Johannesburg
Javiana
Infantis
Heidelberg
Hadar
Enteritidis
Dublin
Derby
Braenderup
Anatum
Agona
0
25
50
75
100
% GC content per genome

J.

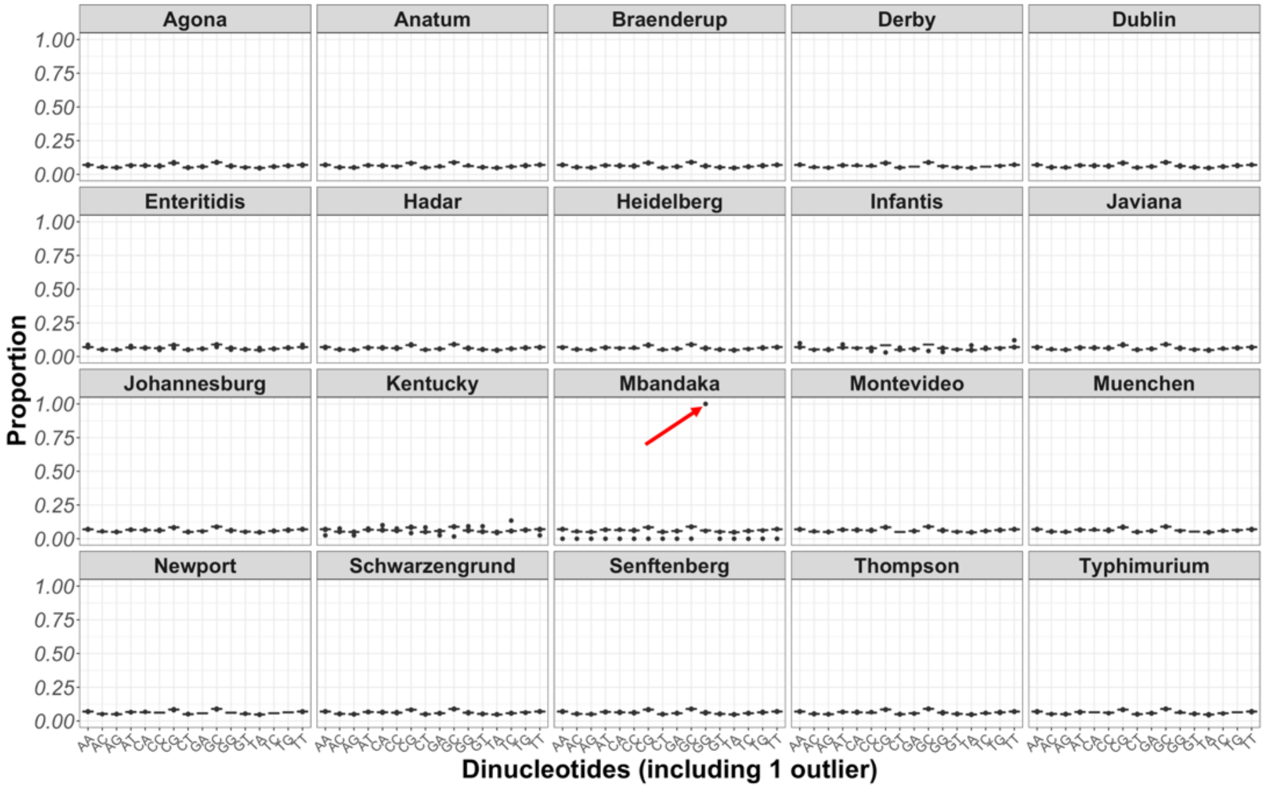
Agona
Anatum
Braenderup
Derby
Dublin
Enteritidis
Hadar
Heidelberg
Infantis
Javiana
Johannesburg
Kentucky
Mbandaka
Montevideo
Muenchen
Newport
Schwarzengrund
Senftenberg
Thompson
Typhimurium
1.00
0.75
0.50
0.25
0.00
Proportion
AA AC AG AT CA CC CG CT GA GC GG GT TA TC TG TT
Dinucleotides (including 1 outlier)

K.

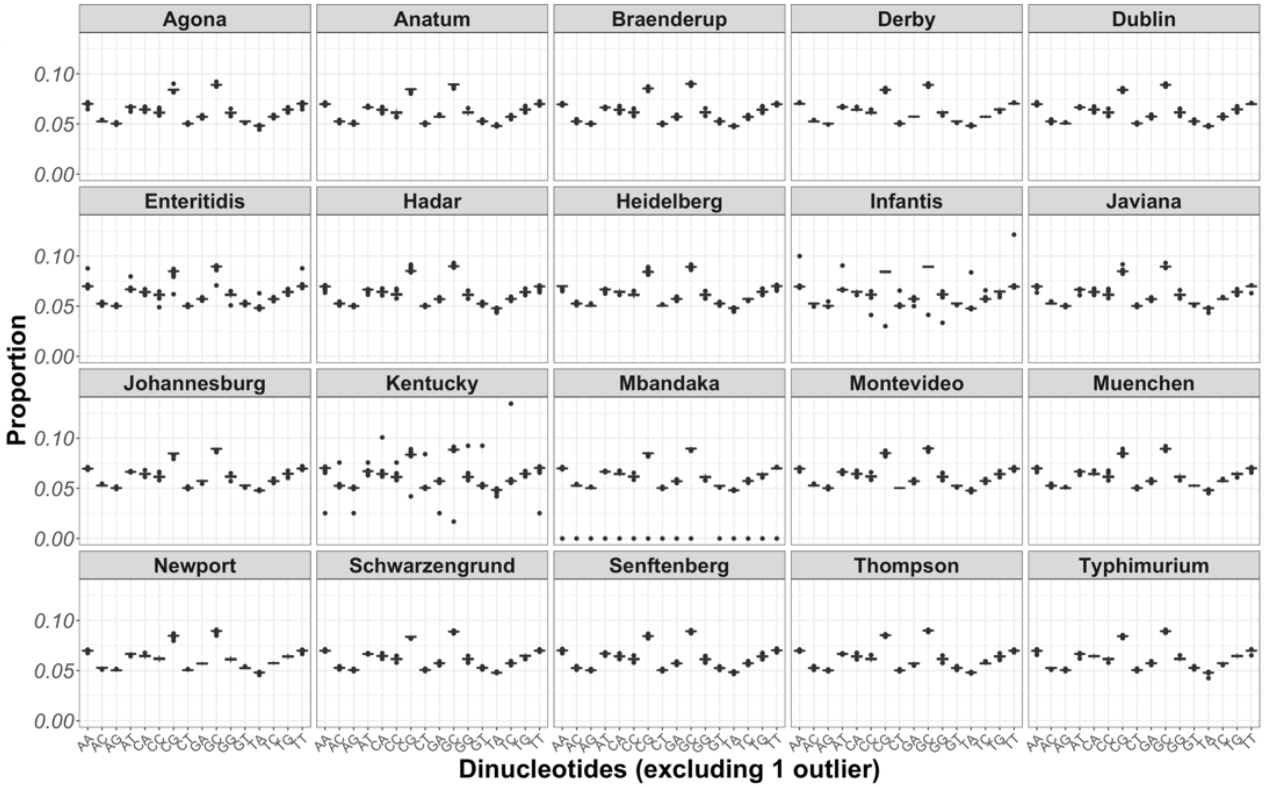
Agona
Anatum
Braenderup
Derby
Dublin
Enteritidis
Hadar
Heidelberg
Infantis
Javiana
Johannesburg
Kentucky
Mbandaka
Montevideo
Muenchen
Newport
Schwarzengrund
Senftenberg
Thompson
Typhimurium
Proportion
0.10
0.05
0.00
Dinucleotides (excluding 1 outlier)

L.

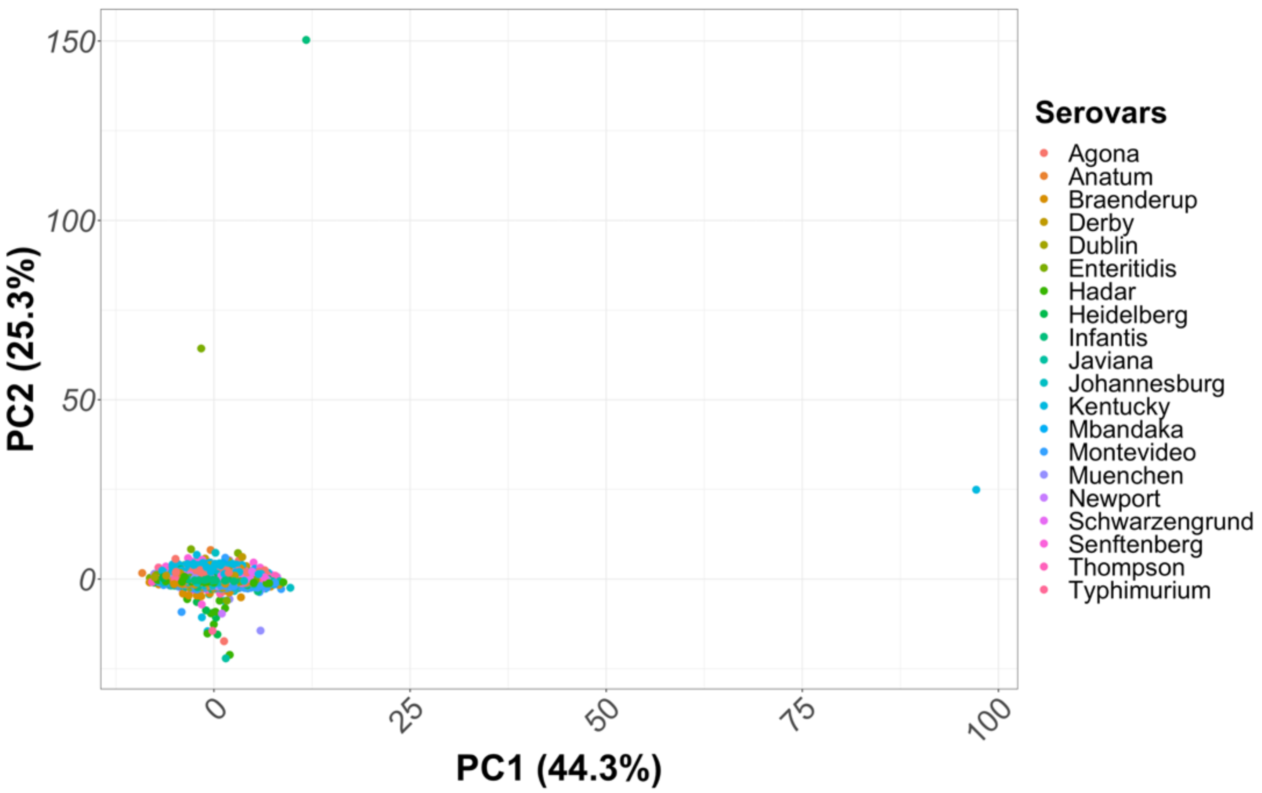

150
100
50
0
PC2 (25.3%)
0
25
50
75
100
PC1 (44.3%)
Serovars
Agona
Anatum
Braenderup
Derby
Dublin
Enteritidis
Hadar
Heidelberg
Infantis
Javiana
Johannesburg
Kentucky
Mbandaka
Montevideo
Muenchen
Newport
Schwarzengrund
Senftenberg
Thompson
Typhimurium

**M.**

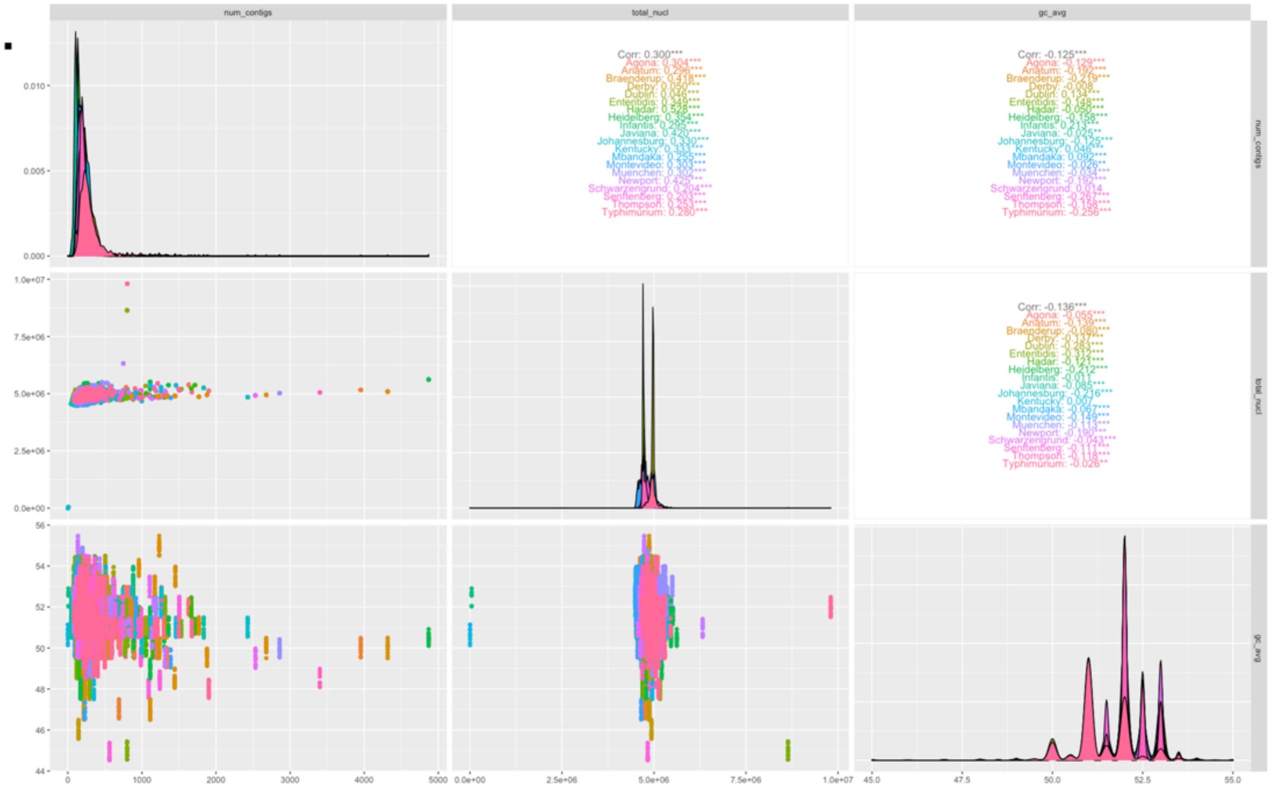
num_contigs
total_nucl
gc_avg
Corr: 0.300***
Agona: 0.304***
Anatum: 0.296***
Braenderup: 0.418***
Derby: 0.050***
Dublin: 0.046***
Enteritidis: 0.349***
Hadar: 0.528***
Heidelberg: 0.354***
Infantis: 0.295***
Javiana: 0.420***
Johannesburg: 0.330***
Kentucky: 0.333***
Mbandaka: 0.255***
Montevideo: 0.303***
Muenchen: 0.302***
Newport: 0.425***
Schwarzengrund: 0.204***
Senftenberg: 0.203***
Thompson: 0.253***
Typhimurium: 0.280***
Corr: -0.125***
Agona: -0.129***
Anatum: -0.192***
Braenderup: -0.219***
Derby: -0.008
Dublin: 0.134***
Enteritidis: -0.148***
Hadar: -0.050***
Heidelberg: -0.158***
Infantis: 0.213***
Javiana: -0.025**
Johannesburg: -0.125***
Kentucky: 0.046***
Mbandaka: 0.092***
Montevideo: -0.026**
Muenchen: -0.034***
Newport: -0.192***
Schwarzengrund: 0.014
Senftenberg: -0.267***
Thompson: -0.158***
Typhimurium: -0.256***
Corr: -0.136***
Agona: -0.055***
Anatum: -0.139***
Braenderup: -0.080***
Derby: -0.137***
Dublin: -0.283***
Enteritidis: -0.312***
Hadar: -0.121***
Heidelberg: -0.212***
Infantis: -0.011
Javiana: -0.085***
Johannesburg: -0.216***
Kentucky: 0.007
Mbandaka: -0.067***
Montevideo: -0.149***
Muenchen: -0.113***
Newport: -0.190***
Schwarzengrund: -0.043***
Senftenberg: -0.111***
Thompson: -0.118***
Typhimurium: -0.026**

**N.**

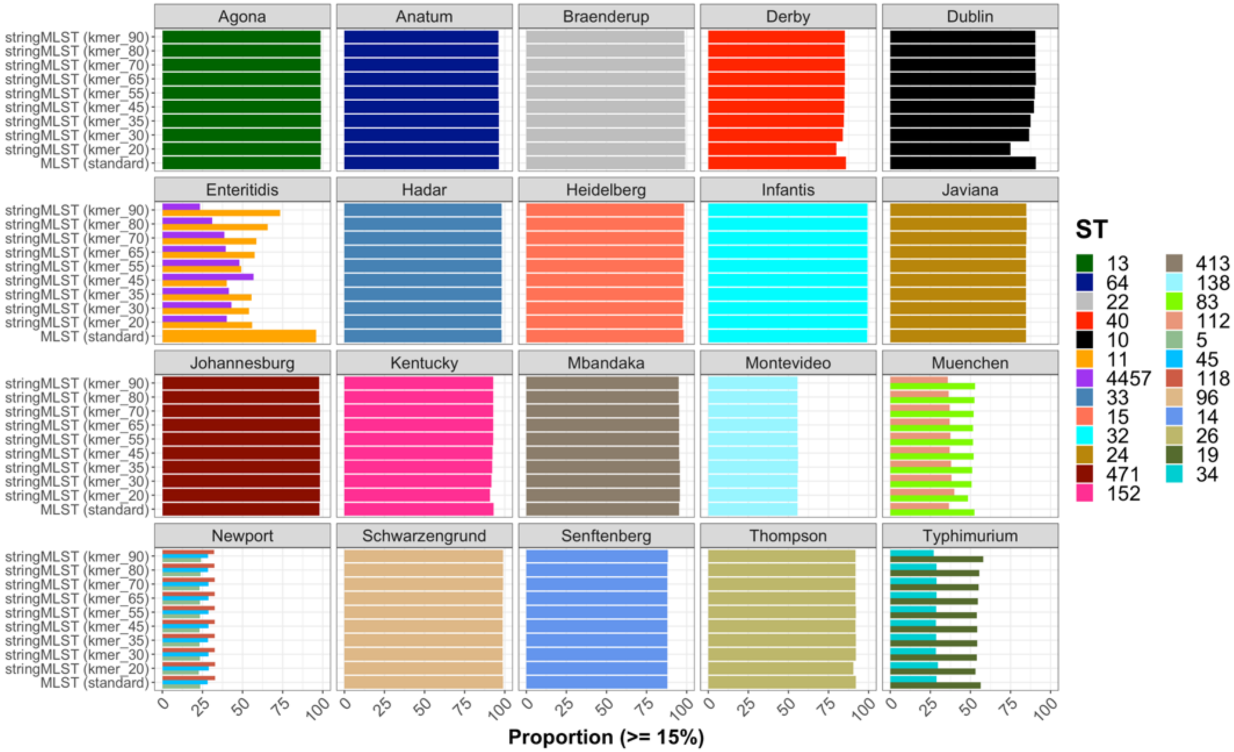

Agona
Anatum
Braenderup
Derby
Dublin
Enteritidis
Hadar
Heidelberg
Infantis
Javiana
Johannesburg
Kentucky
Mbandaka
Montevideo
Muenchen
Newport
Schwarzengrund
Senftenberg
Thompson
Typhimurium
stringMLST (kmer_90)
stringMLST (kmer_80)
stringMLST (kmer_70)
stringMLST (kmer_65)
stringMLST (kmer_55)
stringMLST (kmer_45)
stringMLST (kmer_35)
stringMLST (kmer_30)
stringMLST (kmer_20)
MLST (standard)
0 25 50 75 100
Proportion (>= 15%)
ST
13 64 22 40 10 11 4457 33 15 32 24 471 152
413 138 83 112 5 45 118 96 14 26 19 34

O.

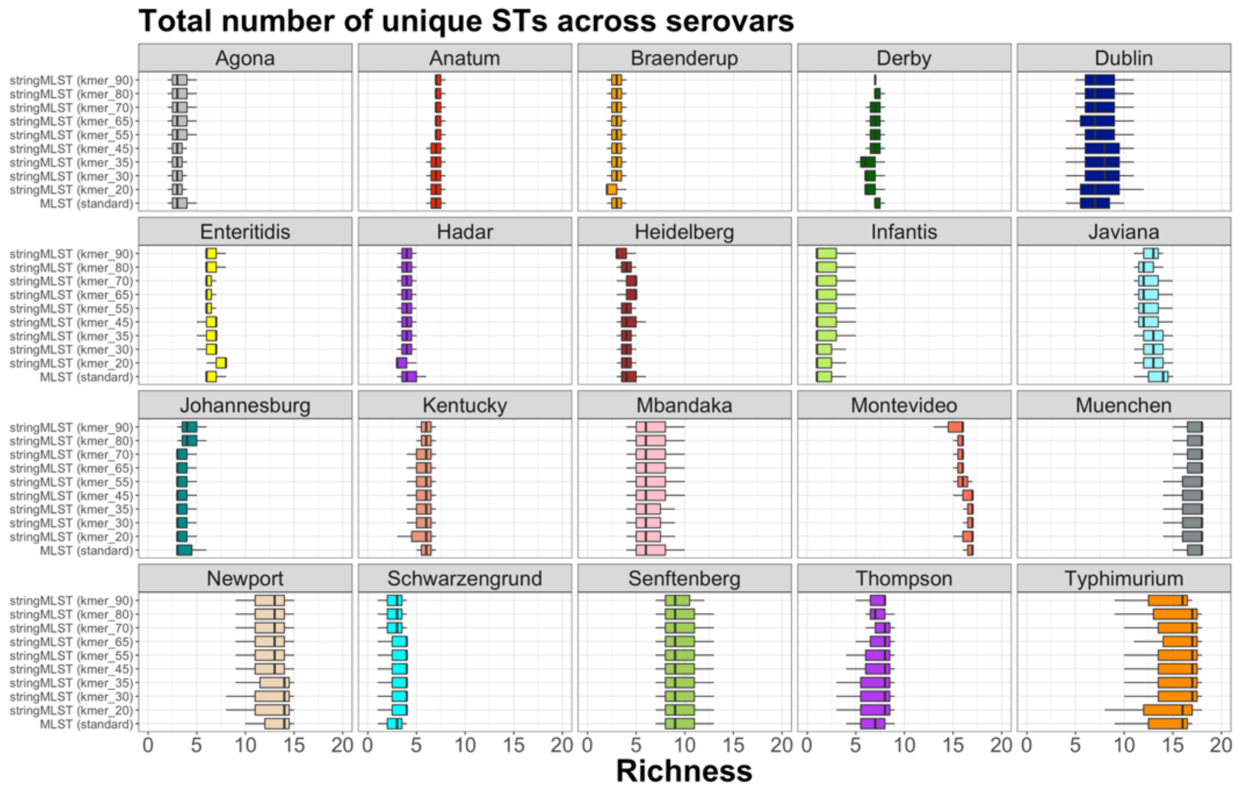
Total number of unique STs across serovars
Agona
Anatum
Braenderup
Derby
Dublin
Enteritidis
Hadar
Heidelberg
Infantis
Javiana
Johannesburg
Kentucky
Mbandaka
Montevideo
Muenchen
Newport
Schwarzengrund
Senftenberg
Thompson
Typhimurium
stringMLST (kmer_90)
stringMLST (kmer_80)
stringMLST (kmer_70)
stringMLST (kmer_65)
stringMLST (kmer_55)
stringMLST (kmer_45)
stringMLST (kmer_35)
stringMLST (kmer_30)
stringMLST (kmer_20)
MLST (standard)
0 5 10 15 20
Richness

P.

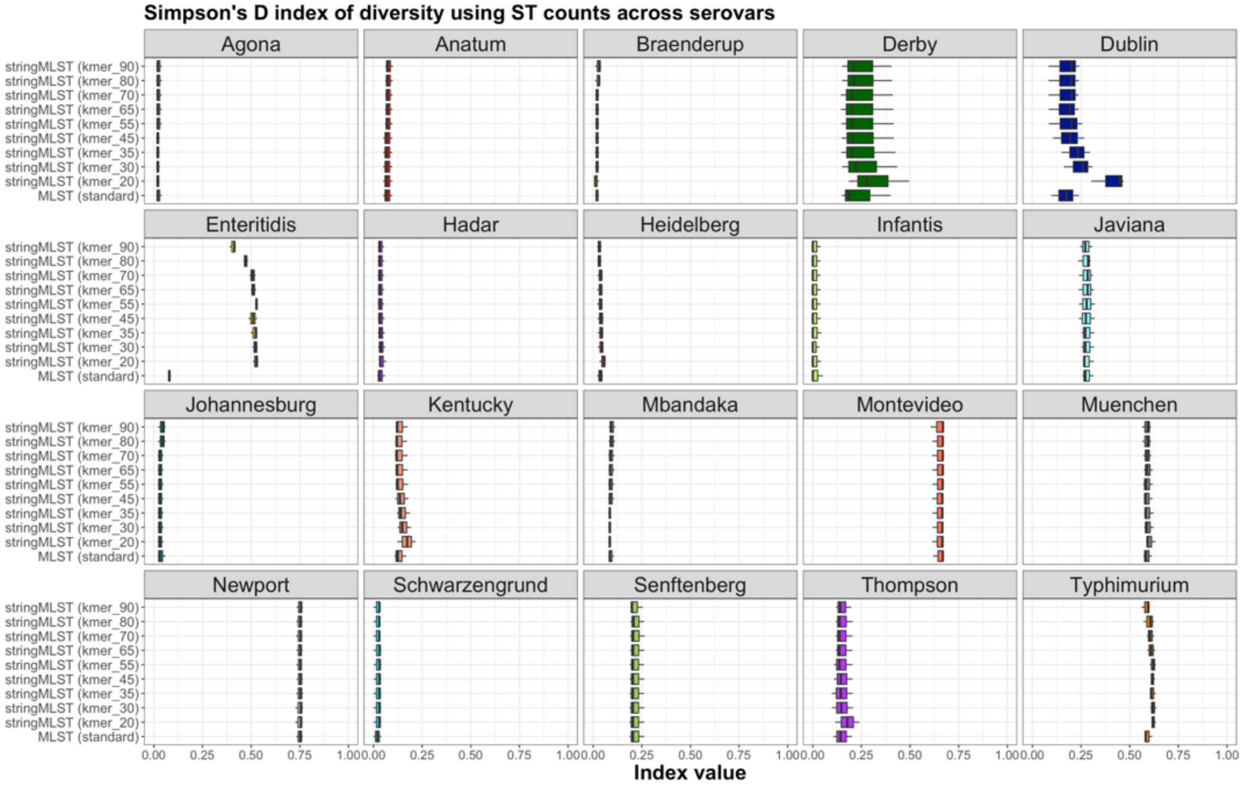

Simpson's D index of diversity using ST counts across serovars
Agona
Anatum
Braenderup
Derby
Dublin
Enteritidis
Hadar
Heidelberg
Infantis
Javiana
Johannesburg
Kentucky
Mbandaka
Montevideo
Muenchen
Newport
Schwarzengrund
Senftenberg
Thompson
Typhimurium
stringMLST (kmer_90)
stringMLST (kmer_80)
stringMLST (kmer_70)
stringMLST (kmer_65)
stringMLST (kmer_55)
stringMLST (kmer_45)
stringMLST (kmer_35)
stringMLST (kmer_30)
stringMLST (kmer_20)
MLST (standard)
0.00
0.25
0.50
0.75
1.00
Index value

Q.

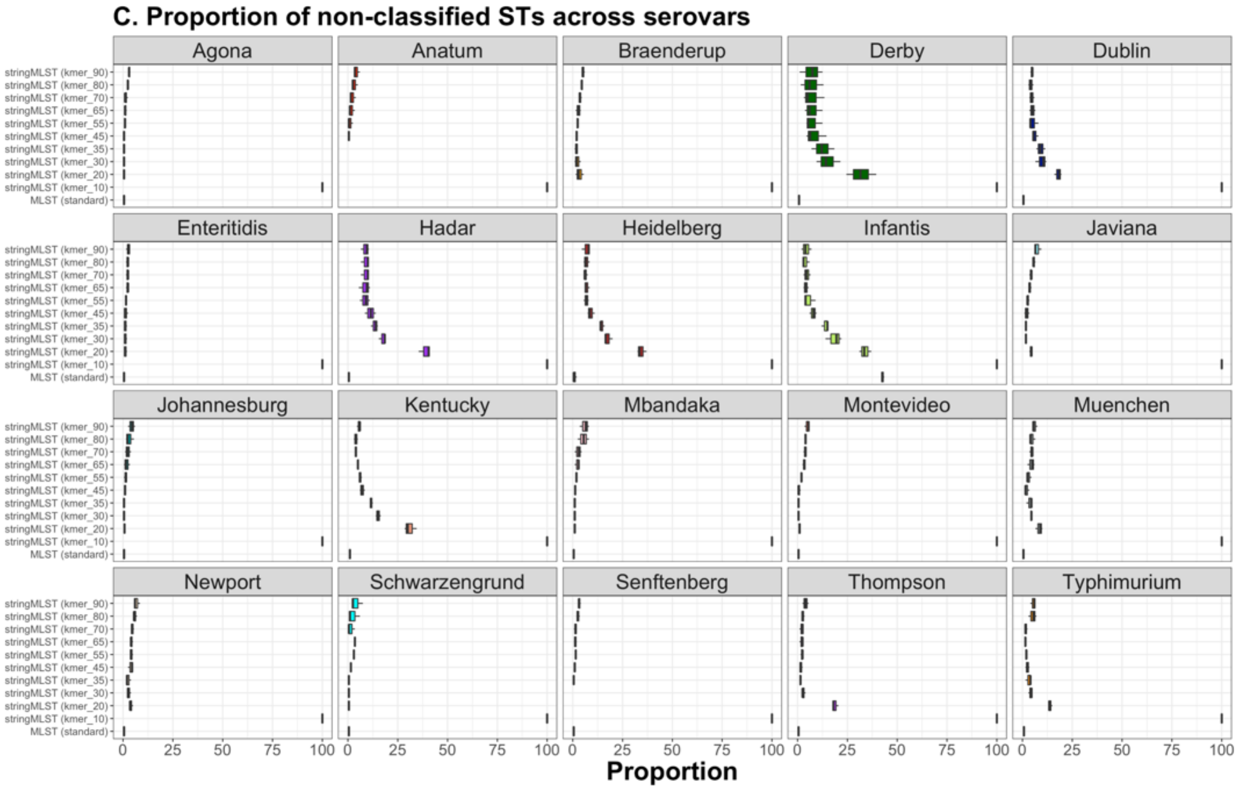
C. Proportion of non-classified STs across serovars
Agona
Anatum
Braenderup
Derby
Dublin
Enteritidis
Hadar
Heidelberg
Infantis
Javiana
Johannesburg
Kentucky
Mbandaka
Montevideo
Muenchen
Newport
Schwarzengrund
Senftenberg
Thompson
Typhimurium
stringMLST (kmer_90)
stringMLST (kmer_80)
stringMLST (kmer_70)
stringMLST (kmer_65)
stringMLST (kmer_55)
stringMLST (kmer_45)
stringMLST (kmer_35)
stringMLST (kmer_30)
stringMLST (kmer_20)
stringMLST (kmer_10)
MLST (standard)
0
25
50
75
100
Proportion

**R.**

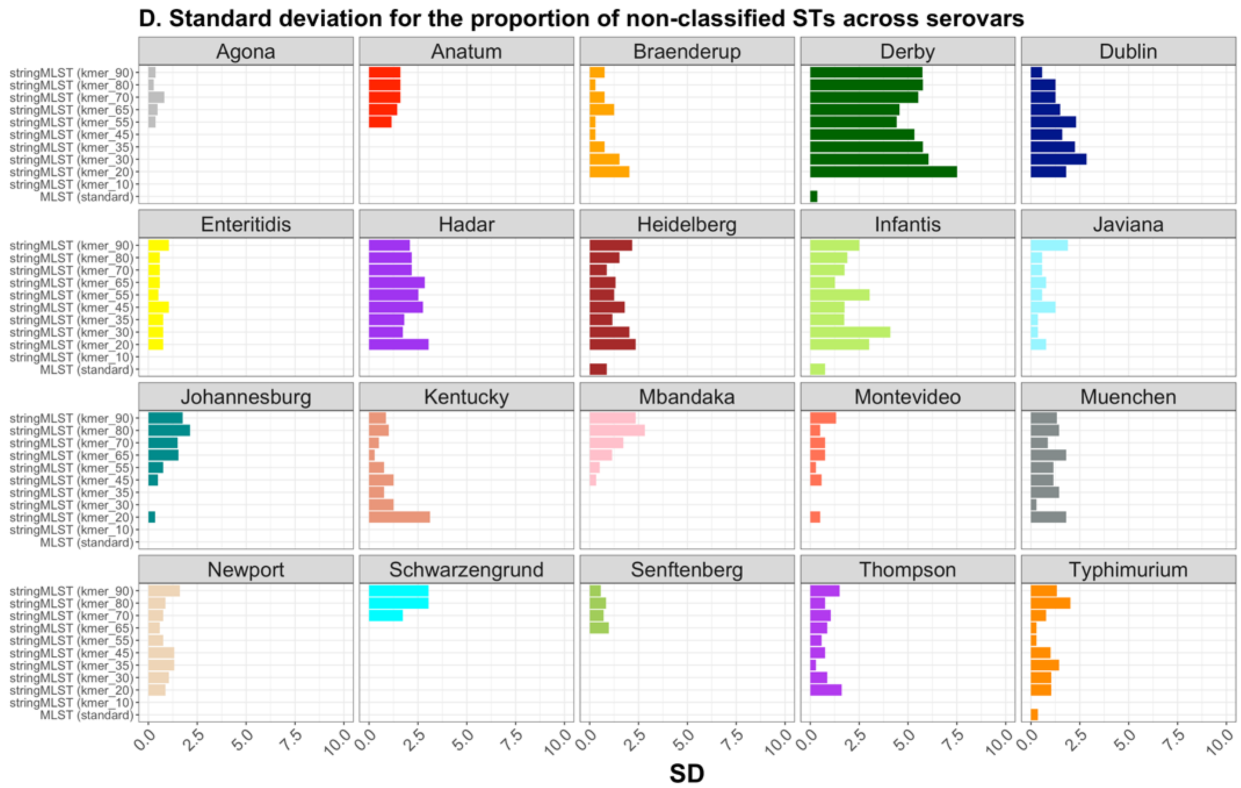

Figure S4. Summary statistics of the frequency of genomes, including the distribution of dinucleotides and bivariate associations between genome-intrinsic variables across all twenty *S. enterica* serovars (serovars). Frequency-based distribution of randomly selected genomes across serovars (A), including a stratification by batch (B), program (C), and further discrimination by kmer length used by stringMLST (D). Total count of unique STs (E), and alleles across all loci (F) by serovar and program. (G) Total number of contigs per genome across all serovars. (H) Total number of nucleotides per genome across all serovars. (I) Percent of GC% content per genome across all serovars. Proportion of all sixteen pairs of dinucleotides present, across serovars, with (J) or without outliers (red-circled data points) (K). (L) Principal component analysis plot depicting two PCs, along with variance explained, for the distribution of serovars using the dinucleotide data as input. (M) Bivariate association between genome-intrinsic variables across species with statistical significance measured by the Pearson correlation coefficient (Corr). Genome-intrinsic variables used were the total number of contigs (num_contigs), the total number of nucleotides per genome (assembly), and the GC% content per genome (gc_avg). (M) Asterisks refer to the degree of significance for the correlation coefficient, with $p$-value thresholds being: *$p < 0.05$, **$p \leq 0.01$, ***$p \leq 0.001$, ****$p \leq 0.0001$ and NS = not significant at $p \geq 0.05$. (N) Proportion of dominant STs (proportion ≥ 15%) across programs and serovars. Distribution of ST richness (O), Simpson's D index of diversity ($1 - D$) (P), proportion of non-classified STs (Q), and the standard deviation of the proportion of non-classified STs (R) across programs and serovars.

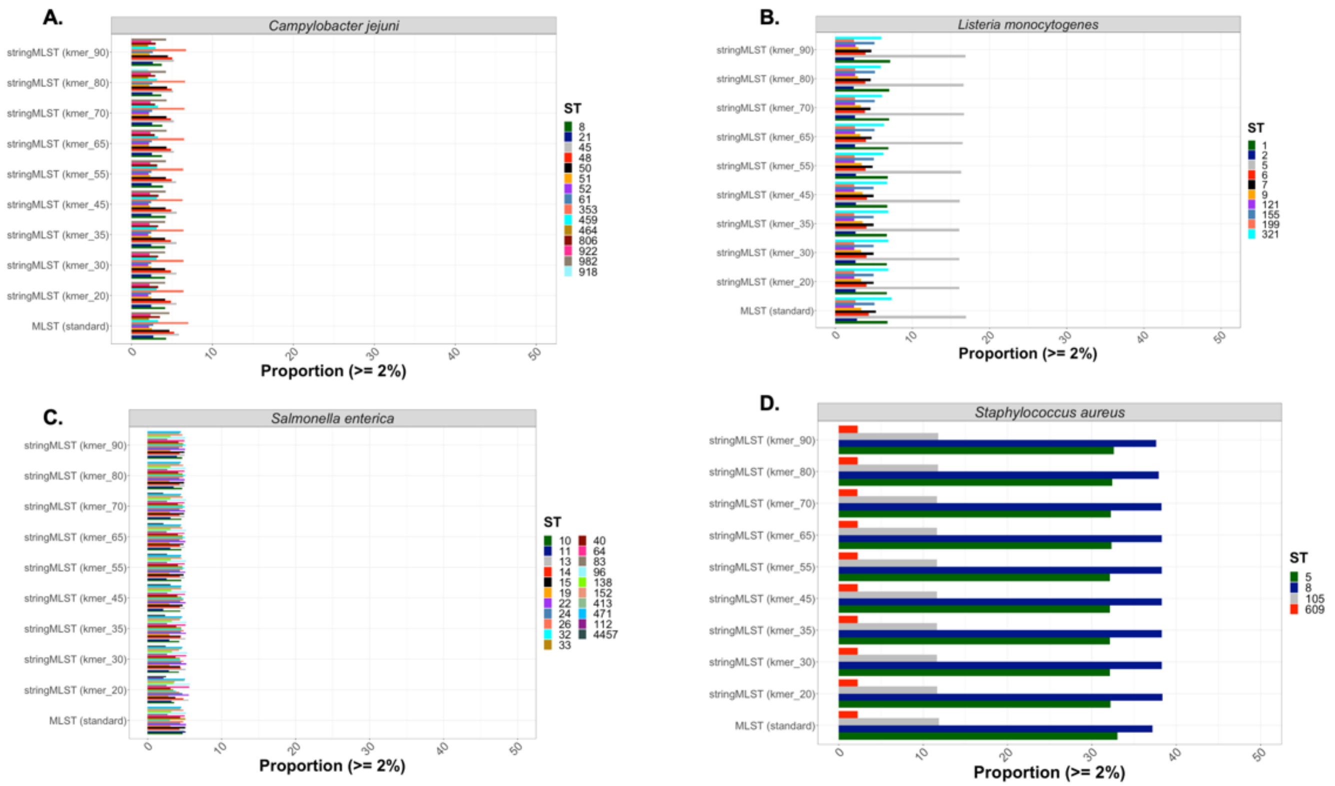

Figure S5. Relative frequency of dominant STs across programs and bacterial species. Plots A-D depict the proportion of STs across all four bacterial species including *C. jejuni*, *L. monocytogenes*, *S. enterica* (all twenty serovars), and *S. aureus*, respectively. Only STs with a proportion equal or above 2% were shown across all plots.

A.

**PERMANOVA model using ST richness as *dependent variable* and the following independent variables: bacterial species (species) and program as MLST or stringMLST (program)**

Model = adonis(formula = n ~ species * program, data = d14, permutations = 1000)

| | Df | SumsOfSqs | MeanSqs | F.Model | $R^2$ | Pr(>F) |
|---|---|---|---|---|---|---|
| **species** | 3 | 7.51793789980392 | 2.50597929993464 | 2011.1886976523 | 0.982921621030805 | 0.000999000999000999 |
| **program** | 9 | 0.0146871548379872 | 0.00163190609310969 | 1.30969600993022 | 0.00192025023804218 | 0.224775224775225 |
| **species:program** | 27 | 0.0162563830939995 | 0.000602088262740721 | 0.483209541692874 | 0.00212541665491391 | 0.99000999000999 |
| **Residuals** | 80 | 0.099681518809644 | 0.00124601898512055 | | 0.0130327120762388 | |
| **Total** | 119 | 7.64856295654555 | | | 1 | |

**B.**

**PERMANOVA model using ST richness as *dependent variable* and the following independent variable: bacterial species (species)**

Model = adonis(formula = n ~ species, data = d14, permutations = 1000)

| | Df | SumsOfSqs | MeanSqs | F.Model | $R^2$ | Pr(>F) |
|---|---|---|---|---|---|---|
| **species** | 3 | 7.51793789980392 | 2.50597929993464 | 2225.40457431069 | 0.982921621030805 | 0.000999000999000999 |
| **Residuals** | 116 | 0.130625056741631 | 0.00112607807535889 | | 0.0170783789691949 | |
| **Total** | 119 | 7.64856295654555 | | | 1 | |

**C.**

**PERMANOVA model using ST richness as *dependent variable* and the following independent variable: program as MLST or stringMLST (program)**

Model = adonis(formula = n ~ program, data = d14, permutations = 1000)

| | Df | SumsOfSqs | MeanSqs | F.Model | $R^2$ | Pr(>F) |
|---|---|---|---|---|---|---|
| **program** | 9 | 0.0146871548379877 | 0.00163190609310974 | 0.0235148795847475 | 0.00192025023804224 | 1 |
| **Residuals** | 110 | 7.63387580170756 | 0.0693988709246142 | | 0.998079749761958 | |
| **Total** | 119 | 7.64856295654555 | | | 1 | |

## D.

**PERMANOVA model using ST richness as *dependent variable* and the following independent variable: median of the number of contigs (num_contigs_median)**

Model = adonis(formula = n ~ num_contigs_median, data = d14, permutations = 1000)

| | Df | SumsOfSqs | MeanSqs | F.Model | $R^2$ | Pr(>F) |
|---|---|---|---|---|---|---|
| **num_contigs_median** | 1 | 1.44641113449941 | 1.44641113449941 | 27.5189190410083 | 0.189108874793478 | 0.000999000999000999 |
| **Residuals** | 118 | 6.20215182204614 | 0.052560608661408 | | 0.810891125206522 | |
| **Total** | 119 | 7.64856295654555 | | | 1 | |

E.

**PERMANOVA model using ST richness as *dependent variable* and the following independent variable: mean of the total counts for nucleotides per genomes (total_nucl_mean)**

Model = adonis(formula = n ~ total_nucl_mean, data = d14, permutations = 1000)

| | Df | SumsOfSqs | MeanSqs | F.Model | $R^2$ | Pr(>F) |
|---|---|---|---|---|---|---|
| **total_nucl_mean** | 1 | 1.3737610065047 | 1.3737610065047 | 25.834090072994 | 0.179610341747798 | 0.000999000999000999 |
| **Residuals** | 118 | 6.27480195004085 | 0.0531762877122106 | | 0.820389658252202 | |
| **Total** | 119 | 7.64856295654555 | | | 1 | |

**F.**

**PERMANOVA model using ST richness as *dependent variable* and the following independent variable:**
**mean of the average GC% per genome (gc_avg_mean)**

Model = adonis(formula = n ~ gc_avg_mean, data = d14, permutations = 1000)

| | Df | SumsOfSqs | MeanSqs | F.Model | $R^2$ | Pr(>F) |
|---|---|---|---|---|---|---|
| **gc_avg_mean** | 1 | 3.35589095505983 | 3.35589095505983 | 92.2491009236216 | 0.438760976947689 | 0.000999000999000999 |
| **Residuals** | 118 | 4.29267200148572 | 0.0363785762837772 | | 0.56123902305231 | |
| **Total** | 119 | 7.64856295654555 | | | 1 | |

**G.**

**PERMANOVA model using ST richness as *dependent variable* and the following independent variable:**
**mean of the total counts of unique STs per program (st_count_mean)**

Model = adonis(formula = n ~ st_count_mean, data = d14, permutations = 1000)

| | Df | SumsOfSqs | MeanSqs | F.Model | $R^2$ | Pr(>F) |
|---|---|---|---|---|---|---|
| **st_count_mean** | 1 | 0.118289690970522 | 0.118289690970522 | 1.85360916426925 | 0.0154656098985615 | 0.166833166833167 |
| **Residuals** | 118 | 7.53027326557503 | 0.0638158751319917 | | 0.984534390101439 | |
| **Total** | 119 | 7.64856295654555 | | | 1 | |

## H.

**PERMANOVA model using ST richness as *dependent variable* and the following independent variable: mean of the total counts of unique alleles across all genes per program (total_alleles_genes_mean)**

Model = adonis(formula = n ~ total_alleles_genes_mean, data = d14, permutations = 1000)

| | Df | SumsOfSqs | MeanSqs | F.Model | $R^2$ | Pr(>F) |
|---|---|---|---|---|---|---|
| **total_alleles_genes_mean** | 1 | 1.06185600851364 | 1.06185600851364 | 19.0230125604796 | 0.138830786194277 | 0.000999000999000999 |
| **Residuals** | 118 | 6.58670694803191 | 0.0558195504070501 | | 0.861169213805723 | |
| **Total** | 119 | 7.64856295654555 | | | 1 | |

I.

**PERMANOVA model using ST richness as *dependent variable* and the following independent variable:**
**Simpson's D index of diversity per species (simpson)**

Model = adonis(formula = n ~ simpson, data = d14, permutations = 1000)

| | Df | SumsOfSqs | MeanSqs | F.Model | $R^2$ | Pr(>F) |
|---|---|---|---|---|---|---|
| **simpson** | 1 | 6.2561804696614 | 6.2561804696614 | 530.191454125543 | 0.817955020466614 | 0.000999000999000999 |
| **Residuals** | 118 | 1.39238248688415 | 0.011799851583764 | | 0.182044979533386 | |
| **Total** | 119 | 7.64856295654555 | | | 1 | |

**J.**

**PERMANOVA model using ST richness as *dependent variable* and the following independent variable:**
**Standard deviation of the number of contigs (num_contigs_sd)**

Model = adonis(formula = n ~ num_contigs_sd, data = d16, permutations = 1000)

| | Df | SumsOfSqs | MeanSqs | F.Model | $R^2$ | Pr(>F) |
|---|---|---|---|---|---|---|
| **num_contigs_sd** | 1 | 3.8179928916089 | 3.8179928916089 | 117.612562509622 | 0.499177808079818 | 0.000999000999000999 |
| **Residuals** | 118 | 3.83057006493665 | 0.0324624581774292 | | 0.500822191920182 | |
| **Total** | 119 | 7.64856295654555 | | | 1 | |

**K.**

**PERMANOVA model using ST richness as *dependent variable* and the following independent variable:**
**Standard deviation of the total counts for nucleotides per genomes (total_nucl_sd)**

Model = adonis(formula = n ~ total_nucl_sd, data = d16, permutations = 1000)

| | Df | SumsOfSqs | MeanSqs | F.Model | $R^2$ | Pr(>F) |
|---|---|---|---|---|---|---|
| **total_nucl_sd** | 1 | 2.05375408880878 | 2.05375408880878 | 43.3156857023196 | 0.268515027002713 | 0.000999000999000999 |
| **Residuals** | 118 | 5.59480886773677 | 0.0474136344723455 | | 0.731484972997287 | |
| **Total** | 119 | 7.64856295654555 | | | 1 | |

**L.**

**PERMANOVA model using ST richness as *dependent variable* and the following independent variable: Standard deviation of the average GC% per genome (gc_avg_sd)**

Model = adonis(formula = n ~ gc_avg_sd, data = d16, permutations = 1000)

| | Df | SumsOfSqs | MeanSqs | F.Model | $R^2$ | Pr(>F) |
|---|---|---|---|---|---|---|
| **gc_avg_sd** | 1 | 1.58113983642368 | 1.58113983642368 | 30.7502043296177 | 0.20672377875514 | 0.000999000999000999 |
| **Residuals** | 118 | 6.06742312012187 | 0.0514188400010328 | | 0.79327622124486 | |
| **Total** | 119 | 7.64856295654555 | | | 1 | |

Figure S6. PERMANOVA results measuring the association between species, program, or other genome-intrinsic and –extrinsic variables and ST richness.

PERMANOVA results demonstrating the association (*R*-squared and *p*-values) between ST richness (n) and: (A) bacterial species and program (mlst vs. stringMLST with all kmer lengths); (B) bacterial species; (C) program (mlst vs. stringMLST with all kmer lengths); (D) the median number of contigs (num_contigs_median); (E) the mean total number of nucleotides (total_nucl_mean); (F) the mean GC% content originally calculated per genome (gc_avg_mean); (G) the mean total count of STs present in each generated database (st_count_mean); (H) the mean total count of unique alleles (across all 7 loci) present in each generated database (total_alleles_genes_mean); (I) the Simpson's D index of diversity (simpson); (J) the standard deviation (SD) of the number of contigs (num_contigs_sd); (K) the SD of the total number of nucleotides (total_nucl_sd); (L) the SD of the GC% content per genome (gc_avg_sd). The median number of contigs, mean total number of nucleotides, and mean GC% content were grouped by species and batch (experimental replicate). The SD of the number of contigs, SD of the total number of nucleotides, and SD of GC% content were calculated by species only. The mean total count of STs and mean total count of unique alleles (across all 7 loci) present in each generated database were calculated after grouping by species, batch (three experimental replicates), and program. The Simpson's D index of diversity was calculated after grouping by program, species, and batch (three experimental replicates). All PERMANOVA models were run with 1,000 permutations.

**A.**

**PERMANOVA model using the Simpson's D index of diversity as *dependent variable* and the following independent variables:**
**bacterial species (species) and program as MLST or stringMLST (program)**

Model = adonis(formula = simpson ~ species * program, data = d13, permutations = 1000)

| | Df | SumsOfSqs | MeanSqs | F.Model | $R^2$ | Pr(>F) |
|---|---|---|---|---|---|---|
| **species** | 3 | 0.410477977217033 | 0.136825992405678 | 2872.17128004194 | 0.990689005884547 | 0.000999000999000999 |
| **program** | 9 | 1.11486912828251E-05 | 1.23874347586946E-06 | 0.0260029792013696 | 2.690737748899E-05 | 1 |
| **species:program** | 27 | 3.56483293753596E-05 | 1.32030849538369E-06 | 0.0277151444295248 | 8.60372783693751E-05 | 1 |
| **Residuals** | 80 | 0.00381108169575958 | 4.76385211969947E-05 | | 0.00919804945959517 | |
| **Total** | 119 | 0.41433585593345 | | | 1 | |

**B.**

**PERMANOVA model using the Simpson's D index of diversity as *dependent variable* and the following independent variable:**
**bacterial species (species)**

Model = adonis(formula = simpson ~ species, data = d13, permutations = 1000)

| | Df | SumsOfSqs | MeanSqs | F.Model | $R^2$ | Pr(>F) |
|---|---|---|---|---|---|---|
| **species** | 3 | 0.410477977217033 | 0.136825992405678 | 4114.13014398661 | 0.990689005884547 | 0.000999000999000999 |
| **Residuals** | 116 | 0.00385787871641774 | 3.32575751415323E-05 | | 0.0093109941154535 | |
| **Total** | 119 | 0.41433585593345 | | | 1 | |

**C.**

**PERMANOVA model using the Simpson's D index of diversity as *dependent variable* and the following independent variable:**
**program as MLST or stringMLST (program)**

Model = adonis(formula = simpson ~ program, data = d13, permutations = 1000)

| | Df | SumsOfSqs | MeanSqs | F.Model | $R^2$ | Pr(>F) |
|---|---|---|---|---|---|---|
| **program** | 9 | 1.11486912828301E-05 | 1.23874347587002E-06 | 0.000328876796299909 | 2.69073774890021E-05 | 1 |
| **Residuals** | 110 | 0.414324707242168 | 0.00376658824765607 | | 0.999973092622511 | |
| **Total** | 119 | 0.41433585593345 | | | 1 | |

**D.**

**PERMANOVA model using the Simpson's D index of diversity as *dependent variable* and the following independent variable:**
**median of the number of contigs (num_contigs_median)**

Model = adonis(formula = simpson ~ num_contigs_median, data = d13, permutations = 1000)

| | Df | SumsOfSqs | MeanSqs | F.Model | $R^2$ | Pr(>F) |
|---|---|---|---|---|---|---|
| **num_contigs_median** | 1 | 0.163614770286817 | 0.163614770286817 | 77.004065470006 | 0.394884410662967 | 0.000999000999000999 |
| **Residuals** | 118 | 0.250721085646634 | 0.00212475496310707 | | 0.605115589337033 | |
| **Total** | 119 | 0.41433585593345 | | | 1 | |

E.

**PERMANOVA model using the Simpson's D index of diversity as *dependent variable* and the following independent variable:**
**mean of the total counts for nucleotides per genomes (total_nucl_mean)**

Model = adonis(formula = simpson ~ total_nucl_mean, data = d13, permutations = 1000)

| | Df | SumsOfSqs | MeanSqs | F.Model | $R^2$ | Pr(>F) |
|---|---|---|---|---|---|---|
| **total_nucl_mean** | 1 | 0.00373280185890127 | 0.00373280185890127 | 1.07274072849536 | 0.00900912099555493 | 0.310689310689311 |
| **Residuals** | 118 | 0.410603054074549 | 0.00347968689893686 | | 0.990990879004445 | |
| **Total** | 119 | 0.41433585593345 | | | 1 | |

**F.**

**PERMANOVA model using the Simpson's D index of diversity as *dependent variable* and the following independent variable:**
**mean of the average GC% per genome (gc_avg_mean)**

Model = adonis(formula = simpson ~ gc_avg_mean, data = d13, permutations = 1000)

| | Df | SumsOfSqs | MeanSqs | F.Model | $R^2$ | Pr(>F) |
|---|---|---|---|---|---|---|
| **gc_avg_mean** | 1 | 0.0717695925601677 | 0.0717695925601677 | 24.7216752715419 | 0.173215982957785 | 0.000999000999000999 |
| **Residuals** | 118 | 0.342566263373283 | 0.00290310392689223 | | 0.826784017042215 | |
| **Total** | 119 | 0.41433585593345 | | | 1 | |

**G.**

**PERMANOVA model using the Simpson's D index of diversity as *dependent variable* and the following independent variable:**
**mean of the total counts of unique STs per program (st_count_mean)**

Model = adonis(formula = simpson ~ st_count_mean, data = d13, permutations = 1000)

| | Df | SumsOfSqs | MeanSqs | F.Model | $R^2$ | Pr(>F) |
|---|---|---|---|---|---|---|
| **st_count_mean** | 1 | 0.00642494016874258 | 0.00642494016874258 | 1.85859929364807 | 0.0155065994813988 | 0.163836163836164 |
| **Residuals** | 118 | 0.407910915764708 | 0.00345687216749752 | | 0.984493400518601 | |
| **Total** | 119 | 0.41433585593345 | | | 1 | |

## H.

**PERMANOVA model using the Simpson's D index of diversity as *dependent variable* and the following independent variable:**
**mean of the total counts of unique alleles across all genes per program (total_alleles_genes_mean)**

Model = adonis(formula = simpson ~ total_alleles_genes_mean, data = d13, permutations = 1000)

| | Df | SumsOfSqs | MeanSqs | F.Model | $R^2$ | Pr(>F) |
|---|---|---|---|---|---|---|
| **total_alleles_genes_mean** | 1 | 6.56768036007165E-05 | 6.56768036007165E-05 | 0.0187072669366709 | 0.000158511030750053 | 0.879120879120879 |
| **Residuals** | 118 | 0.41427017912985 | 0.00351076422991398 | | 0.99984148896925 | |
| **Total** | 119 | 0.41433585593345 | | | 1 | |

**I.**

**PERMANOVA model using the Simpson's D index of diversity as *dependent variable* and the following independent variable:**
**Standard deviation of the number of contigs (num_contigs_sd)**

Model = adonis(formula = simpson ~ num_contigs_sd, data = d14, permutations = 1000)

| | Df | SumsOfSqs | MeanSqs | F.Model | $R^2$ | Pr(>F) |
|---|---|---|---|---|---|---|
| **num_contigs_sd** | 1 | 0.184650012452613 | 0.184650012452613 | 94.8630579020699 | 0.445652988531776 | 0.000999000999000999 |
| **Residuals** | 118 | 0.229685843480837 | 0.00194649019899014 | | 0.554347011468224 | |
| **Total** | 119 | 0.41433585593345 | | | 1 | |

J.

**PERMANOVA model using the Simpson's D index of diversity as *dependent variable* and the following independent variable:**
**Standard deviation of the total counts for nucleotides per genomes (total_nucl_sd)**

Model = adonis(formula = simpson ~ total_nucl_sd, data = d14, permutations = 1000)

| | Df | SumsOfSqs | MeanSqs | F.Model | $R^2$ | Pr(>F) |
|---|---|---|---|---|---|---|
| **total_nucl_sd** | 1 | 0.220300024808541 | 0.220300024808541 | 133.972178111131 | 0.531694328776423 | 0.000999000999000999 |
| **Residuals** | 118 | 0.19403583112491 | 0.0016443714502111 | | 0.468305671223577 | |
| **Total** | 119 | 0.41433585593345 | | | 1 | |

**K.**

**PERMANOVA model using the Simpson's D index of diversity as *dependent variable* and the following independent variable:**
**Standard deviation of the average GC% per genome (gc_avg_sd)**

Model = adonis(formula = simpson ~ gc_avg_sd, data = d14, permutations = 1000)

| | Df | SumsOfSqs | MeanSqs | F.Model | $R^2$ | Pr(>F) |
|---|---|---|---|---|---|---|
| **gc_avg_sd** | 1 | 0.0712010372963118 | 0.0712010372963118 | 24.4851934127079 | 0.171843774263524 | 0.000999000999000999 |
| **Residuals** | 118 | 0.343134818637139 | 0.00290792219184016 | | 0.828156225736476 | |
| **Total** | 119 | 0.41433585593345 | | | 1 | |

Figure S7. PERMANOVA results measuring the association between species, program, or other genome-intrinsic and –extrinsic variables and the Simpson's D index of diversity.
PERMANOVA results demonstrating the association (*R*-squared and *p*-values) between the Simpson's D index of diversity (simpson) and: (A) bacterial species and program (mlst vs. stringMLST with all kmer lengths); (B) bacterial species; (C) program (mlst vs. stringMLST with all kmer lengths); (D) the median number of contigs (num_contigs_median); (E) the mean total number of nucleotides (total_nucl_mean); (F) the mean GC% content originally calculated per genome (gc_avg_mean); (G) the mean total count of STs present in each generated database (st_count_mean); (H) the mean total count of unique alleles (across all 7 loci) present in each generated database (total_alleles_genes_mean); (I) the standard deviation (SD) of the number of contigs (num_contigs_sd); (J) the SD of the total number of nucleotides (total_nucl_sd); (K) the SD of the GC% content per genome (gc_avg_sd). The median number of contigs, mean total number of nucleotides, and mean GC% content were grouped by species and batch (experimental replicate). The SD of the number of contigs, SD of the total number of

nucleotides, and SD of GC% content were calculated by species only. The mean total count of STs and mean total count of unique alleles (across all 7 loci) present in each generated database were calculated after grouping by species, batch (three experimental replicates), and program. The Simpson's D index of diversity was calculated after grouping by program, species, and batch (three experimental replicates). All PERMANOVA models were run with 1,000 permutations.

**A.**

**PERMANOVA model using the proportion of non-classified STs as *dependent variable* and the following independent variables:**
**bacterial species (species) and program as MLST or stringMLST (program)**

Model = adonis(formula = prop ~ species * program, data = d18, permutations = 1000)

| | Df | SumsOfSqs | MeanSqs | F.Model | $R^2$ | Pr(>F) |
|---|---|---|---|---|---|---|
| **species** | 3 | 3.34480401068386 | 1.11493467022795 | 23.3264912660022 | 0.324922836900245 | 0.000999000999000999 |
| **program** | 9 | 0.681352798076 | 0.0757058664528889 | 1.58390646533342 | 0.0661883576357917 | 0.0719280719280719 |
| **species:program** | 27 | 2.44423752682071 | 0.0905273158081743 | 1.89399854352207 | 0.237439499813996 | 0.002997002997003 |
| **Residuals** | 80 | 3.82375440014142 | 0.0477969300017678 | | 0.371449305649967 | |
| **Total** | 119 | 10.294148735722 | | | 1 | |

**B.**

**PERMANOVA model using the proportion of non-classified STs as *dependent variable* and the following independent variable:**
**bacterial species (species)**

Model = adonis(formula = prop ~ species, data = d18, permutations = 1000)

| | Df | SumsOfSqs | MeanSqs | F.Model | $R^2$ | Pr(>F) |
|---|---|---|---|---|---|---|
| **species** | 3 | 3.34480401068386 | 1.11493467022795 | 18.610736244018 | 0.324922836900245 | 0.000999000999000999 |
| **Residuals** | 116 | 6.94934472503813 | 0.0599081441813632 | | 0.675077163099755 | |
| **Total** | 119 | 10.294148735722 | | | 1 | |

**C.**

**PERMANOVA model using the proportion of non-classified STs as *dependent variable* and the following independent variable:**
**program as MLST or stringMLST (program)**

Model = adonis(formula = prop ~ program, data = d18, permutations = 1000)

| | Df | SumsOfSqs | MeanSqs | F.Model | $R^2$ | Pr(>F) |
|---|---|---|---|---|---|---|
| **program** | 9 | 0.681352798076 | 0.0757058664528889 | 0.866308341905474 | 0.0661883576357917 | 0.631368631368631 |
| **Residuals** | 110 | 9.61279593764599 | 0.0873890539785999 | | 0.933811642364208 | |
| **Total** | 119 | 10.294148735722 | | | 1 | |

**D.**

**PERMANOVA model using the proportion of non-classified STs as *dependent variable* and the following independent variable:**
**median of the number of contigs (num_contigs_median)**

Model = adonis(formula = prop ~ num_contigs_median, data = d18, permutations = 1000)

| | Df | SumsOfSqs | MeanSqs | F.Model | $R^2$ | Pr(>F) |
|---|---|---|---|---|---|---|
| **num_contigs_median** | 1 | 2.78200760329276 | 2.78200760329276 | 43.6995114177773 | 0.270251351006698 | 0.000999000999000999 |
| **Residuals** | 118 | 7.51214113242923 | 0.0636622129866884 | | 0.729748648993302 | |
| **Total** | 119 | 10.294148735722 | | | 1 | |

E.

**PERMANOVA model using the proportion of non-classified STs as *dependent variable* and the following independent variable:**
**mean of the total counts for nucleotides per genomes (total_nucl_mean)**

Model = adonis(formula = prop ~ total_nucl_mean, data = d18, permutations = 1000)

| | Df | SumsOfSqs | MeanSqs | F.Model | $R^2$ | Pr(>F) |
|---|---|---|---|---|---|---|
| **total_nucl_mean** | 1 | 0.312873903121628 | 0.312873903121628 | 3.69883819327053 | 0.0303933730854224 | 0.035964035964036 |
| **Residuals** | 118 | 9.98127483260036 | 0.0845870748525455 | | 0.969606626914578 | |
| **Total** | 119 | 10.294148735722 | | | 1 | |

**F.**

**PERMANOVA model using the proportion of non-classified STs as *dependent variable* and the following independent variable:**
**mean of the average GC% per genome (gc_avg_mean)**

Model = adonis(formula = prop ~ gc_avg_mean, data = d18, permutations = 1000)

| | Df | SumsOfSqs | MeanSqs | F.Model | $R^2$ | Pr(>F) |
|---|---|---|---|---|---|---|
| **gc_avg_mean** | 1 | 0.235317166826526 | 0.235317166826526 | 2.7605021015954 | 0.022859312884216 | 0.0659340659340659 |
| **Residuals** | 118 | 10.0588315688955 | 0.0852443353296226 | | 0.977140687115784 | |
| **Total** | 119 | 10.294148735722 | | | 1 | |

**G.**

**PERMANOVA model using the proportion of non-classified STs as *dependent variable* and the following independent variable:**
**mean of the total counts of unique STs per program (st_count_mean)**

Model = adonis(formula = prop ~ st_count_mean, data = d18, permutations = 1000)

| | Df | SumsOfSqs | MeanSqs | F.Model | $R^2$ | Pr(>F) |
|---|---|---|---|---|---|---|
| **st_count_mean** | 1 | 0.0631382909852366 | 0.0631382909852366 | 0.728209435079862 | 0.00613341545825336 | 0.472527472527473 |
| **Residuals** | 118 | 10.2310104447368 | 0.0867034783452267 | | 0.993866584541747 | |
| **Total** | 119 | 10.294148735722 | | | 1 | |

## H.

**PERMANOVA model using the proportion of non-classified STs as *dependent variable* and the following independent variable:**
**mean of the total counts of unique alleles across all genes per program (total_alleles_genes_mean)**

Model = adonis(formula = prop ~ total_alleles_genes_mean, data = d18, permutations = 1000)

| | Df | SumsOfSqs | MeanSqs | F.Model | $R^2$ | Pr(>F) |
|---|---|---|---|---|---|---|
| **total_alleles_genes_mean** | 1 | 0.835211515433743 | 0.835211515433743 | 10.4192423023798 | 0.0811345878980215 | 0.001998001998002 |
| **Residuals** | 118 | 9.45893722028825 | 0.080160484917697 | | 0.918865412101978 | |
| **Total** | 119 | 10.294148735722 | | | 1 | |

# I.

**PERMANOVA model using the proportion of non-classified STs as *dependent variable* and the following independent variable:**
**Simpson's D index of diversity per species (simpson)**

Model = adonis(formula = prop ~ simpson, data = d18, permutations = 1000)

| | Df | SumsOfSqs | MeanSqs | F.Model | $R^2$ | Pr(>F) |
|---|---|---|---|---|---|---|
| **simpson** | 1 | 2.37335228285581 | 2.37335228285581 | 35.3569961106178 | 0.230553525481906 | 0.000999000999000999 |
| **Residuals** | 118 | 7.92079645286619 | 0.0671253936683575 | | 0.769446474518094 | |
| **Total** | 119 | 10.294148735722 | | | 1 | |

**J.**

**PPERMANOVA model using the proportion of non-classified STs as *dependent variable* and the following independent variable:**
**Standard deviation of the number of contigs (num_contigs_sd)**

Model = adonis(formula = prop ~ num_contigs_sd, data = d18b, permutations = 1000)

| | Df | SumsOfSqs | MeanSqs | F.Model | $R^2$ | Pr(>F) |
|---|---|---|---|---|---|---|
| **num_contigs_sd** | 1 | 0.343617214899233 | 0.343617214899233 | 4.07484075330651 | 0.0333798572102265 | 0.022977022977023 |
| **Residuals** | 118 | 9.95053152082276 | 0.0843265383120573 | | 0.966620142789774 | |
| **Total** | 119 | 10.294148735722 | | | 1 | |

**K.**

**PERMANOVA model using the proportion of non-classified STs as *dependent variable* and the following independent variable:**
**Standard deviation of the total counts for nucleotides per genomes (total_nucl_sd)**

Model = adonis(formula = prop ~ total_nucl_sd, data = d18b, permutations = 1000)

| | Df | SumsOfSqs | MeanSqs | F.Model | $R^2$ | Pr(>F) |
|---|---|---|---|---|---|---|
| **total_nucl_sd** | 1 | 2.51052137919844 | 2.51052137919844 | 38.0595716079769 | 0.243878483170407 | 0.000999000999000999 |
| **Residuals** | 118 | 7.78362735652355 | 0.0659629436993521 | | 0.756121516829593 | |
| **Total** | 119 | 10.294148735722 | | | 1 | |

**L.**

**PERMANOVA model using the proportion of non-classified STs as *dependent variable* and the following independent variable:**
**Standard deviation of the average GC% per genome (gc_avg_sd)**

Model = adonis(formula = prop ~ gc_avg_sd, data = d18b, permutations = 1000)

| | Df | SumsOfSqs | MeanSqs | F.Model | $R^2$ | Pr(>F) |
|---|---|---|---|---|---|---|
| **gc_avg_sd** | 1 | 0.67622148287334 | 0.67622148287334 | 8.29639618613466 | 0.0656898885215022 | 0.002997002997003 |
| **Residuals** | 118 | 9.61792725284865 | 0.0815078580749886 | | 0.934310111478498 | |
| **Total** | 119 | 10.294148735722 | | | 1 | |

Figure S8. PERMANOVA results measuring the association between species, program, or other genome-intrinsic and –extrinsic variables and the proportion of non-classified STs.
PERMANOVA results demonstrating the association (*R*-squared and *p*-values) between non-classified STs (prop) and: (A) bacterial species and program (mlst vs. stringMLST with all kmer lengths); (B) bacterial species; (C) program (mlst vs. stringMLST with all kmer lengths); (D) the median number of contigs (num_contigs_median); (E) the mean total number of nucleotides (total_nucl_mean); (F) the mean GC% content originally calculated per genome (gc_avg_mean); (G) the mean total count of STs present in each generated database (st_count_mean); (H) the mean total count of unique alleles (across all 7 loci) present in each generated database (total_alleles_genes_mean); (I) the Simpson's D index of diversity (simpson); (J) the standard deviation (SD) of the number of contigs (num_contigs_sd); (K) the SD of the total number of nucleotides (total_nucl_sd); (L) the SD of the GC% content per genome (gc_avg_sd). The median number of contigs, mean total number of nucleotides, and mean GC% content were grouped by species and batch (experimental replicate). The SD of the number of contigs, SD

of the total number of nucleotides, and SD of GC% content were calculated by species only. The mean total count of STs and mean total count of unique alleles (across all 7 loci) present in each generated database were calculated after grouping by species, batch (three experimental replicates), and program. The Simpson's D index of diversity was calculated after grouping by program, species, and batch (three experimental replicates). All PERMANOVA models were run with 1,000 permutations.

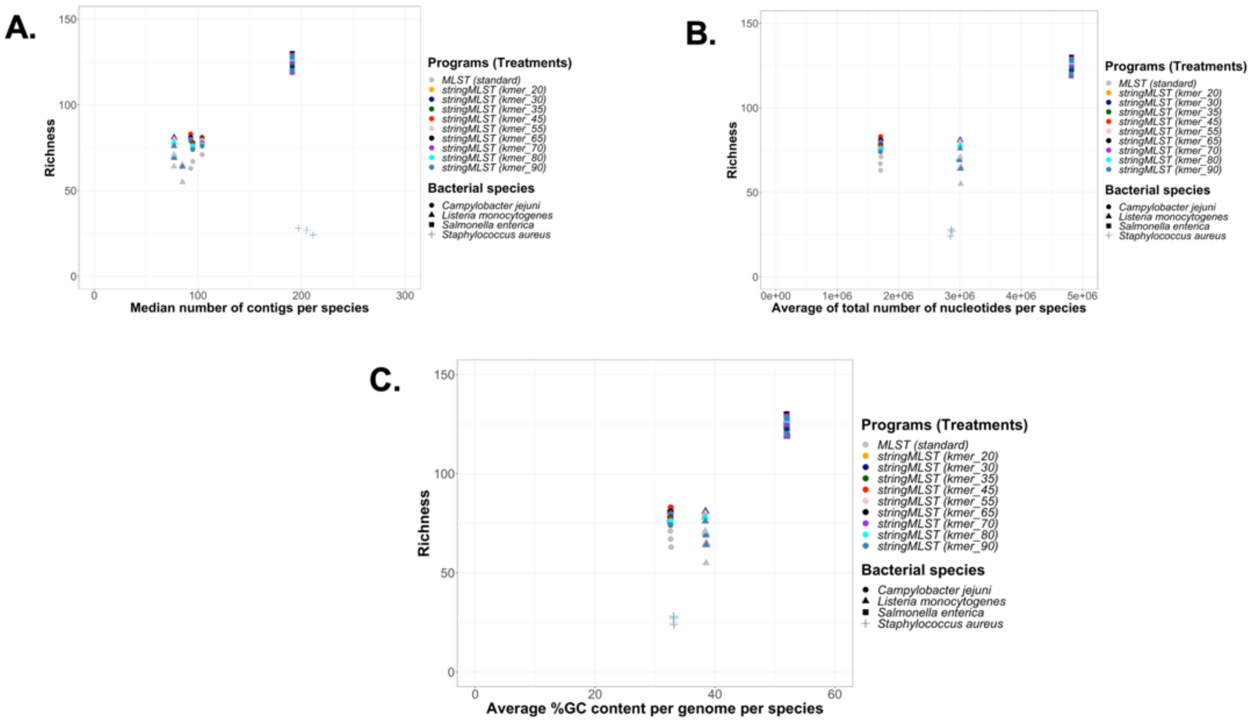
A.
B.
C.
Richness
Median number of contigs per species
Average of total number of nucleotides per species
Average %GC content per genome per species
Programs (Treatments)
MLST (standard)
stringMLST (kmer_20)
stringMLST (kmer_30)
stringMLST (kmer_35)
stringMLST (kmer_45)
stringMLST (kmer_55)
stringMLST (kmer_65)
stringMLST (kmer_70)
stringMLST (kmer_80)
stringMLST (kmer_90)
Bacterial species
Campylobacter jejuni
Listeria monocytogenes
Salmonella enterica
Staphylococcus aureus

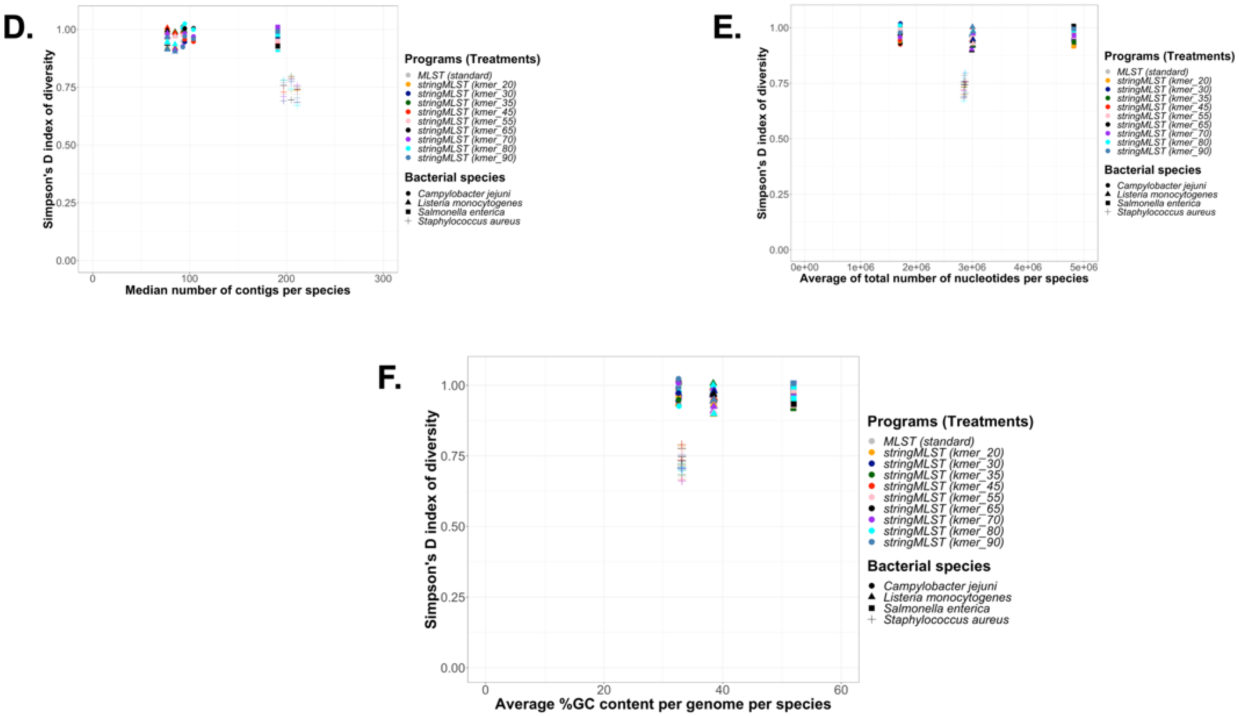
D.
Simpson's D index of diversity
Median number of contigs per species
Programs (Treatments)
MLST (standard)
stringMLST (kmer_20)
stringMLST (kmer_30)
stringMLST (kmer_35)
stringMLST (kmer_45)
stringMLST (kmer_55)
stringMLST (kmer_65)
stringMLST (kmer_70)
stringMLST (kmer_80)
stringMLST (kmer_90)
Bacterial species
Campylobacter jejuni
Listeria monocytogenes
Salmonella enterica
Staphylococcus aureus
E.
Simpson's D index of diversity
Average of total number of nucleotides per species
Programs (Treatments)
MLST (standard)
stringMLST (kmer_20)
stringMLST (kmer_30)
stringMLST (kmer_35)
stringMLST (kmer_45)
stringMLST (kmer_55)
stringMLST (kmer_65)
stringMLST (kmer_70)
stringMLST (kmer_80)
stringMLST (kmer_90)
Bacterial species
Campylobacter jejuni
Listeria monocytogenes
Salmonella enterica
Staphylococcus aureus
F.
Simpson's D index of diversity
Average %GC content per genome per species
Programs (Treatments)
MLST (standard)
stringMLST (kmer_20)
stringMLST (kmer_30)
stringMLST (kmer_35)
stringMLST (kmer_45)
stringMLST (kmer_55)
stringMLST (kmer_65)
stringMLST (kmer_70)
stringMLST (kmer_80)
stringMLST (kmer_90)
Bacterial species
Campylobacter jejuni
Listeria monocytogenes
Salmonella enterica
Staphylococcus aureus

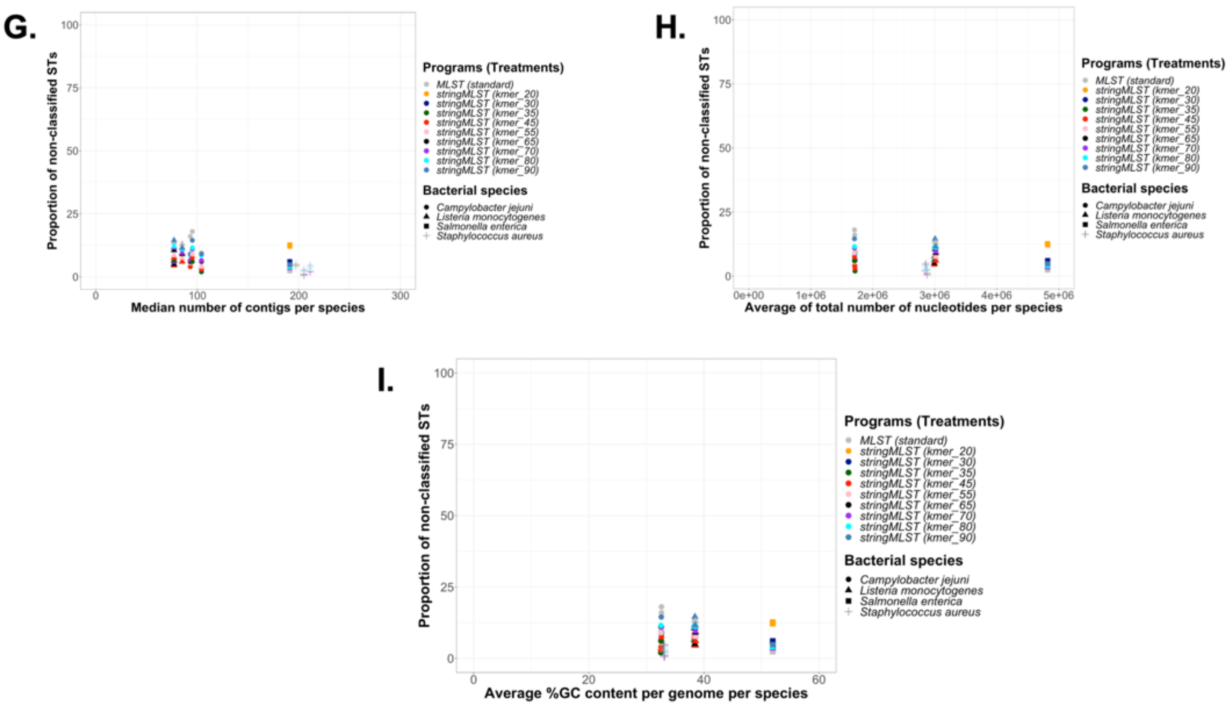

G.
Proportion of non-classified STs
Median number of contigs per species
Programs (Treatments)
MLST (standard)
stringMLST (kmer_20)
stringMLST (kmer_30)
stringMLST (kmer_35)
stringMLST (kmer_45)
stringMLST (kmer_55)
stringMLST (kmer_65)
stringMLST (kmer_70)
stringMLST (kmer_80)
stringMLST (kmer_90)
Bacterial species
Campylobacter jejuni
Listeria monocytogenes
Salmonella enterica
Staphylococcus aureus
H.
Proportion of non-classified STs
Average of total number of nucleotides per species
I.
Proportion of non-classified STs
Average %GC content per genome per species

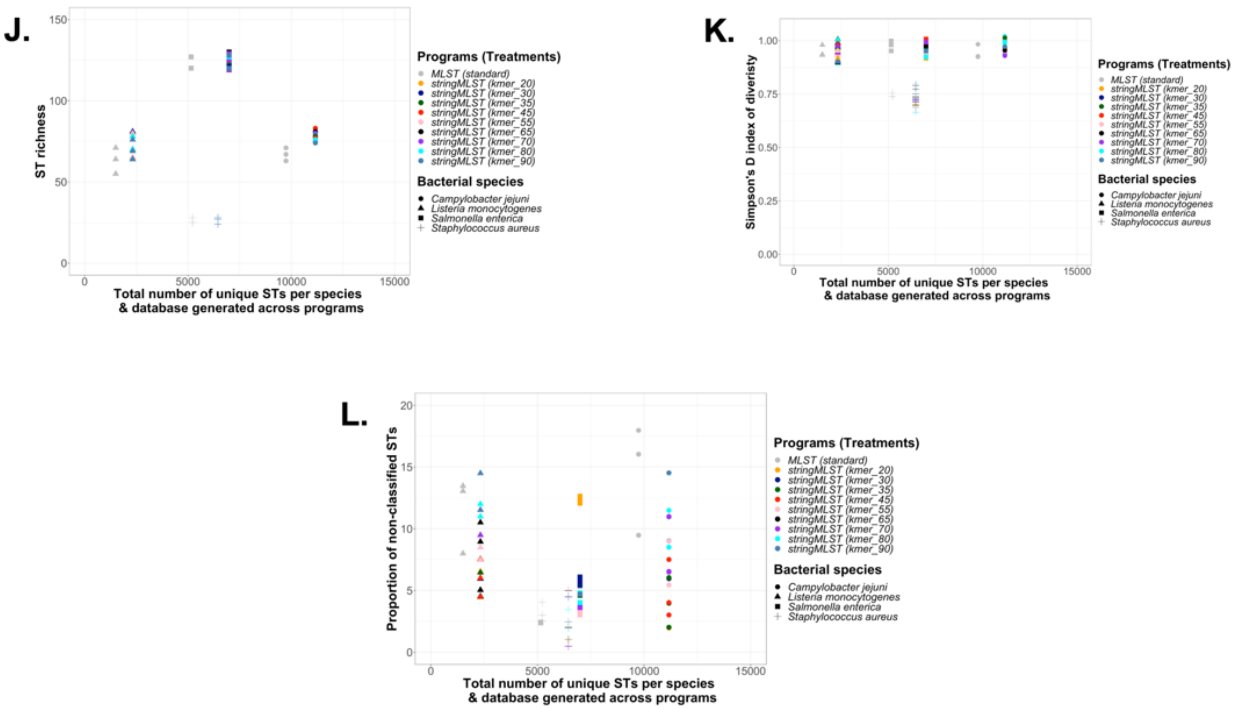
J.
ST richness
Total number of unique STs per species
& database generated across programs
K.
Simpson's D index of diveristy
Total number of unique STs per species
& database generated across programs
L.
Proportion of non-classified STs
Total number of unique STs per species
& database generated across programs
Programs (Treatments)
MLST (standard)
stringMLST (kmer_20)
stringMLST (kmer_30)
stringMLST (kmer_35)
stringMLST (kmer_45)
stringMLST (kmer_55)
stringMLST (kmer_65)
stringMLST (kmer_70)
stringMLST (kmer_80)
stringMLST (kmer_90)
Bacterial species
Campylobacter jejuni
Listeria monocytogenes
Salmonella enterica
Staphylococcus aureus

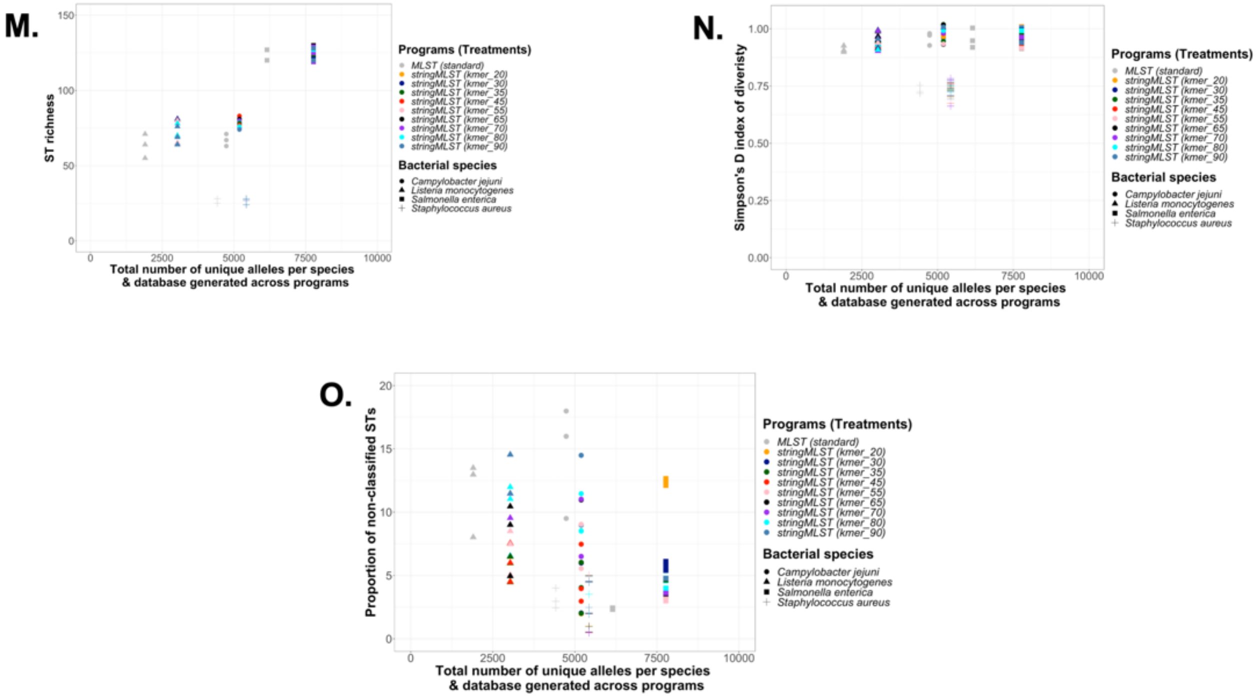

Figure S9. Bivariate association between accuracy-based classification metrics for ST calls and genome-intrinsic and –extrinsic variables.

Plot A-C depict groupings generated based on the relationship between ST richness (Richness) vs. the median number of contigs per species, or the average of total number of nucleotides per species, or the average GC% content per genome per species, respectively. Plot D-F depict groupings generated based on the relationship between the Simpson's D index of diversity vs. the median number of contigs per species, or the average of total number of nucleotides per species, or the average GC% content per genome per species, respectively. Plot G-I depict groupings generated based on the relationship between the proportion of non-classified STs vs. the median number of contigs per species, or the average of total number of nucleotides per species, or the average GC% content per genome per species, respectively. Plot J-L depict groupings generated based on the relationship between the total number of unique STs per species (generated across databases per program) vs. ST richness, or the Simpson's D index of diversity, or the proportion of non-classified STs, respectively. Plot M-O depict groupings generated based on the relationship between the total number of unique alleles per species (per database generated across programs) vs. ST richness, or the Simpson's D index of diversity, or the proportion of non-classified STs, respectively.

**A.**

**PERMANOVA model using ST richness as *dependent variable* and the following independent variables:**
***S. enterica* serovars (serovar) and program as MLST or stringMLST (program)**

Model = adonis(formula = n ~ serovar * program, data = d14, permutations = 1000)

| | Df | SumsOfSqs | MeanSqs | F.Model | $R^2$ | Pr(>F) |
|---|---|---|---|---|---|---|
| **serovar** | 19 | 39.5743056229991 | 2.08285819068416 | 65.7453972769742 | 0.753954496075907 | 0.000999000999000999 |
| **program** | 9 | 0.0302818094505213 | 0.00336464549450236 | 0.106205000283566 | 0.000576917422178744 | 1 |
| **serovar:program** | 171 | 0.212129026087382 | 0.00124052062039405 | 0.0391570205705148 | 0.00404140086475292 | 1 |
| **Residuals** | 400 | 12.6722677294621 | 0.0316806693236552 | | 0.241427185637162 | |
| **Total** | 599 | 52.4889841879991 | | | 1 | |

**B.**

**PERMANOVA model using ST richness as *dependent variable* and the following independent variable:**
***S. enterica* serovars (serovar)**

Model = adonis(formula = n ~ serovar, data = d14, permutations = 1000)

| | Df | SumsOfSqs | MeanSqs | F.Model | $R^2$ | Pr(>F) |
|---|---|---|---|---|---|---|
| **serovar** | 19 | 39.5743056229991 | 2.08285819068416 | 93.5414493296616 | 0.753954496075907 | 0.000999000999000999 |
| **Residuals** | 580 | 12.914678565 | 0.0222666871810345 | | 0.246045503924093 | |
| **Total** | 599 | 52.4889841879991 | | | 1 | |

C.

**PERMANOVA model using ST richness as *dependent variable* and the following independent variable: program as MLST or stringMLST (program)**

Model = adonis(formula = n ~ program, data = d14, permutations = 1000)

| | Df | SumsOfSqs | MeanSqs | F.Model | $R^2$ | Pr(>F) |
|---|---|---|---|---|---|---|
| **program** | 9 | 0.0302818094505365 | 0.00336464549450406 | 0.0378419738146088 | 0.000576917422179034 | 1 |
| **Residuals** | 590 | 52.4587023785486 | 0.0889130548788959 | | 0.999423082577821 | |
| **Total** | 599 | 52.4889841879991 | | | 1 | |

**D.**

**PERMANOVA model using ST richness as *dependent variable* and the following independent variable:**
**median of the number of contigs (num_contigs_median)**

Model = adonis(formula = n ~ num_contigs_median, data = d14, permutations = 1000)

| | Df | SumsOfSqs | MeanSqs | F.Model | $R^2$ | Pr(>F) |
|---|---|---|---|---|---|---|
| **num_contigs_median** | 1 | 0.44768112521022 | 0.44768112521022 | 5.1442469177359 | 0.00852904913546748 | 0.011988011988012 |
| **Residuals** | 598 | 52.0413030627889 | 0.0870255904060015 | | 0.991470950864533 | |
| **Total** | 599 | 52.4889841879991 | | | 1 | |

**E.**

**PERMANOVA model using ST richness as *dependent variable* and the following independent variable:**
**mean of the total counts for nucleotides per genomes (total_nucl_mean)**

Model = adonis(formula = n ~ total_nucl_mean, data = d14, permutations = 1000)

| | Df | SumsOfSqs | MeanSqs | F.Model | $R^2$ | Pr(>F) |
|---|---|---|---|---|---|---|
| **total_nucl_mean** | 1 | 0.238098824297164 | 0.238098824297164 | 2.72498917364979 | 0.00453616750220138 | 0.0729270729270729 |
| **Residuals** | 598 | 52.2508853637019 | 0.0873760624811069 | | 0.995463832497799 | |
| **Total** | 599 | 52.4889841879991 | | | 1 | |

**F.**

**PERMANOVA model using ST richness as *dependent variable* and the following independent variable: mean of the average GC% per genome (gc_avg_mean)**

Model = adonis(formula = n ~ gc_avg_mean, data = d14, permutations = 1000)

| | Df | SumsOfSqs | MeanSqs | F.Model | $R^2$ | Pr(>F) |
|---|---|---|---|---|---|---|
| **gc_avg_mean** | 1 | 2.54799906279066 | 2.54799906279066 | 30.5100797617166 | 0.0485435011213881 | 0.000999000999000999 |
| **Residuals** | 598 | 49.9409851252084 | 0.083513353052188 | | 0.951456498878612 | |
| **Total** | 599 | 52.4889841879991 | | | 1 | |

G.

**PERMANOVA model using ST richness as *dependent variable* and the following independent variable: mean of the total counts of unique STs per program (st_count_mean)**

Model = adonis(formula = n ~ st_count_mean, data = d14, permutations = 1000)

| | Df | SumsOfSqs | MeanSqs | F.Model | $R^2$ | Pr(>F) |
|---|---|---|---|---|---|---|
| **st_count_mean** | 1 | 0.00089060831664228 | 0.00089060831664228 | 0.0101467539975245 | 1.69675281474757E-05 | 0.996003996003996 |
| **Residuals** | 598 | 52.4880935796825 | 0.0877727317385995 | | 0.999983032471852 | |
| **Total** | 599 | 52.4889841879991 | | | 1 | |

## H.

**PERMANOVA model using ST richness as *dependent variable* and the following independent variable: mean of the total counts of unique alleles across all genes per program (total_alleles_genes_mean)**

Model = adonis(formula = n ~ total_alleles_genes_mean, data = d14, permutations = 1000)

| | Df | SumsOfSqs | MeanSqs | F.Model | $R^2$ | Pr(>F) |
|---|---|---|---|---|---|---|
| **total_alleles_genes_mean** | 1 | 0.000890608316637105 | 0.000890608316637105 | 0.0101467539974656 | 1.69675281473771E-05 | 0.996003996003996 |
| **Residuals** | 598 | 52.4880935796825 | 0.0877727317385994 | | 0.999983032471853 | |
| **Total** | 599 | 52.4889841879991 | | | 1 | |

I.

**PERMANOVA model using ST richness as *dependent variable* and the following independent variable:**
**Simpson's D index of diversity per *S. enterica* serovar (simpson)**

Model = adonis(formula = n ~ simpson, data = d14, permutations = 1000)

| | Df | SumsOfSqs | MeanSqs | F.Model | $R^2$ | Pr(>F) |
|---|---|---|---|---|---|---|
| **simpson** | 1 | 24.4898921577139 | 24.4898921577139 | 523.051086602101 | 0.466572034810176 | 0.000999000999000999 |
| **Residuals** | 598 | 27.9990920302852 | 0.0468212241309117 | | 0.533427965189824 | |
| **Total** | 599 | 52.4889841879991 | | | 1 | |

**J.**

**PERMANOVA model using ST richness as *dependent variable* and the following independent variable:**
**Standard deviation of the number of contigs (num_contigs_sd)**

Model = adonis(formula = n ~ num_contigs_sd, data = d16, permutations = 1000)

| | Df | SumsOfSqs | MeanSqs | F.Model | $R^2$ | Pr(>F) |
|---|---|---|---|---|---|---|
| **num_contigs_sd** | 1 | 2.36289028502892 | 2.36289028502892 | 28.1890783906377 | 0.0450168796668989 | 0.000999000999000999 |
| **Residuals** | 598 | 50.1260939029702 | 0.0838228995032946 | | 0.954983120333101 | |
| **Total** | 599 | 52.4889841879991 | | | 1 | |

K.

**PERMANOVA model using ST richness as *dependent variable* and the following independent variable: Standard deviation of the total counts for nucleotides per genomes (total_nucl_sd)**

Model = adonis(formula = n ~ total_nucl_sd, data = d16, permutations = 1000)

| | Df | SumsOfSqs | MeanSqs | F.Model | $R^2$ | Pr(>F) |
|---|---|---|---|---|---|---|
| **total_nucl_sd** | 1 | 1.704263803706 | 1.704263803706 | 20.0680391051517 | 0.0324689804931615 | 0.000999000999000999 |
| **Residuals** | 598 | 50.7847203842931 | 0.0849242815790854 | | 0.967531019506839 | |
| **Total** | 599 | 52.4889841879991 | | | 1 | |

L.

PERMANOVA model using ST richness as ***dependent variable*** and the following **independent variable**:
**Standard deviation of the average GC% per genome (gc_avg_sd)**

Model = adonis(formula = n ~ gc_avg_sd, data = d16, permutations = 1000)

| | Df | SumsOfSqs | MeanSqs | F.Model | $R^2$ | Pr(>F) |
|---|---|---|---|---|---|---|
| **gc_avg_sd** | 1 | 0.884904844087155 | 0.884904844087155 | 10.2544818838348 | 0.0168588677753355 | 0.002997002997003 |
| **Residuals** | 598 | 51.6040793439119 | 0.0862944470633979 | | 0.983141132224664 | |
| **Total** | 599 | 52.4889841879991 | | | 1 | |

Figure S10. PERMANOVA results measuring the association between *S. enterica* serovars (serovar), program, or genome-intrinsic and –extrinsic variables and ST richness.
PERMANOVA results demonstrating the association (*R*-squared and *p*-values) between ST richness (n) and: (A) serovar and program (mlst vs. stringMLST with all kmer lengths); (B) serovar; (C) program (mlst vs. stringMLST with all kmer lengths); (D) the median number of contigs (num_contigs_median); (E) the mean total number of nucleotides (total_nucl_mean); (F) the mean GC% content originally calculated per genome (gc_avg_mean); (G) the mean total count of STs present in each generated database (st_count_mean); (H) the mean total count of unique alleles (across all 7 loci) present in each generated database (total_alleles_genes_mean); (I) the Simpson's D index of diversity (simpson); (J) the standard deviation (SD) of the number of contigs (num_contigs_sd); (K) the SD of the total number of nucleotides (total_nucl_sd); (L) the SD of the GC% content per genome (gc_avg_sd). The median number of contigs, mean total number of nucleotides, and mean GC% content were grouped by serovar and batch (experimental replicate). The SD of the number of contigs, SD of the total number of nucleotides, and SD of

GC% content were calculated by serovar only. The mean total count of STs and mean total count of unique alleles (across all 7 loci) present in each generated database were calculated after grouping by serovar, batch (three experimental replicates), and program. The Simpson's D index of diversity was calculated after grouping by program, serovar, and batch (three experimental replicates). All PERMANOVA models were run with 1,000 permutations.

**A.**

**PERMANOVA model using the Simpson's D index of diversity as *dependent variable* and the following independent variables:**
***S. enterica* serovars (serovar) and program as MLST or stringMLST (program)**

Model = adonis(formula = simpson ~ serovar * program, data = d14, permutations = 1000)

| | Df | SumsOfSqs | MeanSqs | F.Model | $R^2$ | Pr(>F) |
|---|---|---|---|---|---|---|
| **serovar** | 17 | 85.7297109948326 | 5.04292417616662 | 194.682446036456 | 0.880851447565824 | 0.000999000999000999 |
| **program** | 9 | 0.192616168185751 | 0.0214017964650834 | 0.826217872774765 | 0.00197908319767026 | 0.659340659340659 |
| **serovar:program** | 153 | 2.07843106001404 | 0.0135845167321179 | 0.524431233396458 | 0.0213553619466825 | 1 |
| **Residuals** | 360 | 9.32519978241913 | 0.025903332728942 | | 0.0958141072898229 | |
| **Total** | 539 | 97.3259580054515 | | | 1 | |

**B.**

**PERMANOVA model using the Simpson's D index of diversity as *dependent variable* and the following independent variable:**
***S. enterica* serovars (serovar)**

Model = adonis(formula = simpson ~ serovar, data = d14, permutations = 1000)

| | Df | SumsOfSqs | MeanSqs | F.Model | $R^2$ | Pr(>F) |
|---|---|---|---|---|---|---|
| **serovar** | 17 | 85.7297109948326 | 5.04292417616662 | 227.005031675199 | 0.880851447565824 | 0.000999000999000999 |
| **Residuals** | 522 | 11.5962470106189 | 0.0222150325873926 | | 0.119148552434176 | |
| **Total** | 539 | 97.3259580054515 | | | 1 | |

C.

**PERMANOVA model using the Simpson's D index of diversity as *dependent variable* and the following independent variable:**
**program as MLST or stringMLST (program)**

Model = adonis(formula = simpson ~ program, data = d14, permutations = 1000)

| | Df | SumsOfSqs | MeanSqs | F.Model | $R^2$ | Pr(>F) |
|---|---|---|---|---|---|---|
| **program** | 9 | 0.192616168185761 | 0.0214017964650846 | 0.116777122169836 | 0.00197908319767037 | 1 |
| **Residuals** | 530 | 97.1333418372657 | 0.183270456296728 | | 0.99802091680233 | |
| **Total** | 539 | 97.3259580054515 | | | 1 | |

**D.**

**PERMANOVA model using the Simpson's D index of diversity as *dependent variable* and the following independent variable:**
**median of the number of contigs (num_contigs_median)**

Model = adonis(formula = simpson ~ num_contigs_median, data = d14, permutations = 1000)

| | Df | SumsOfSqs | MeanSqs | F.Model | $R^2$ | Pr(>F) |
|---|---|---|---|---|---|---|
| **num_contigs_median** | 1 | 1.06238403124153 | 1.06238403124153 | 5.93747546669179 | 0.0109157315582963 | 0.003996003996004 |
| **Residuals** | 538 | 96.26357397421 | 0.178928576160242 | | 0.989084268441704 | |
| **Total** | 539 | 97.3259580054515 | | | 1 | |

E.

**PERMANOVA model using the Simpson's D index of diversity as *dependent variable* and the following independent variable:**
**mean of the total counts for nucleotides per genomes (total_nucl_mean)**

Model = adonis(formula = simpson ~ total_nucl_mean, data = d14, permutations = 1000)

| | Df | SumsOfSqs | MeanSqs | F.Model | $R^2$ | Pr(>F) |
|---|---|---|---|---|---|---|
| **total_nucl_mean** | 1 | 0.70701945670127 | 0.70701945670127 | 3.93687276447733 | 0.00726444898350415 | 0.01998001998002 |
| **Residuals** | 538 | 96.6189385487502 | 0.179589105109201 | | 0.992735551016496 | |
| **Total** | 539 | 97.3259580054515 | | | 1 | |

**F.**

**PERMANOVA model using the Simpson's D index of diversity as *dependent variable* and the following independent variable:**
**mean of the average GC% per genome (gc_avg_mean)**

Model = adonis(formula = simpson ~ gc_avg_mean, data = d14, permutations = 1000)

| | Df | SumsOfSqs | MeanSqs | F.Model | $R^2$ | Pr(>F) |
|---|---|---|---|---|---|---|
| **gc_avg_mean** | 1 | 7.47341510145232 | 7.47341510145232 | 44.7477299432379 | 0.0767874804893646 | 0.000999000999000999 |
| **Residuals** | 538 | 89.8525429039992 | 0.167012161531597 | | 0.923212519510635 | |
| **Total** | 539 | 97.3259580054515 | | | 1 | |

G.

**PERMANOVA model using the Simpson's D index of diversity as *dependent variable* and the following independent variable: mean of the total counts of unique STs per program (st_count_mean)**

Model = adonis(formula = simpson ~ st_count_mean, data = d14, permutations = 1000)

| | Df | SumsOfSqs | MeanSqs | F.Model | $R^2$ | Pr(>F) |
|---|---|---|---|---|---|---|
| **st_count_mean** | 1 | 0.110218487813067 | 0.110218487813067 | 0.609958292120707 | 0.00113246753560745 | 0.533466533466533 |
| **Residuals** | 538 | 97.2157395176384 | 0.180698400590406 | | 0.998867532464393 | |
| **Total** | 539 | 97.3259580054515 | | | 1 | |

**H.**

**PERMANOVA model using the Simpson's D index of diversity as *dependent variable* and the following independent variable:**
**mean of the total counts of unique alleles across all genes per program (total_alleles_genes_mean)**

Model = adonis(formula = simpson ~ total_alleles_genes_mean, data = d14, permutations = 1000)

| | Df | SumsOfSqs | MeanSqs | F.Model | $R^2$ | Pr(>F) |
|---|---|---|---|---|---|---|
| **total_alleles_genes_mean** | 1 | 0.110218487813107 | 0.110218487813107 | 0.609958292120928 | 0.00113246753560786 | 0.533466533466533 |
| **Residuals** | 538 | 97.2157395176384 | 0.180698400590406 | | 0.998867532464392 | |
| **Total** | 539 | 97.3259580054515 | | | 1 | |

**I.**

**PERMANOVA model using the Simpson's D index of diversity as *dependent variable* and the following independent variable:**
**Standard deviation of the number of contigs (num_contigs_sd)**

Model = adonis(formula = simpson ~ num_contigs_sd, data = d16, permutations = 1000)

| | Df | SumsOfSqs | MeanSqs | F.Model | $R^2$ | Pr(>F) |
|---|---|---|---|---|---|---|
| **num_contigs_sd** | 1 | 18.1227489554589 | 18.1227489554589 | 123.101564380841 | 0.186206735868388 | 0.000999000999000999 |
| **Residuals** | 538 | 79.2032090499926 | 0.147217860687719 | | 0.813793264131612 | |
| **Total** | 539 | 97.3259580054515 | | | 1 | |

J.

**PERMANOVA model using the Simpson's D index of diversity as *dependent variable* and the following independent variable:**
**Standard deviation of the total counts for nucleotides per genomes (total_nucl_sd)**

Model = adonis(formula = simpson ~ total_nucl_sd, data = d16, permutations = 1000)

| | Df | SumsOfSqs | MeanSqs | F.Model | $R^2$ | Pr(>F) |
|---|---|---|---|---|---|---|
| **total_nucl_sd** | 1 | 12.5127094628339 | 12.5127094628339 | 79.3724778460993 | 0.128564976078972 | 0.000999000999000999 |
| **Residuals** | 538 | 84.8132485426176 | 0.157645443387765 | | 0.871435023921028 | |
| **Total** | 539 | 97.3259580054515 | | | 1 | |

**K.**

**PERMANOVA model using the Simpson's D index of diversity as *dependent variable* and the following independent variable: Standard deviation of the average GC% per genome (gc_avg_sd)**

Model = adonis(formula = simpson ~ gc_avg_sd, data = d16, permutations = 1000)

| | Df | SumsOfSqs | MeanSqs | F.Model | $R^2$ | Pr(>F) |
|---|---|---|---|---|---|---|
| **gc_avg_sd** | 1 | 1.35976083775894 | 1.35976083775894 | 7.62301052146506 | 0.0139712042462791 | 0.001998001998002 |
| **Residuals** | 538 | 95.9661971676926 | 0.178375831166715 | | 0.986028795753721 | |
| **Total** | 539 | 97.3259580054515 | | | 1 | |

Figure S11. PERMANOVA results measuring the association between *S. enterica* serovars (serovar), program, or genome-intrinsic and –extrinsic variables and the Simpson's D index of diversity.
PERMANOVA results demonstrating the association (*R*-squared and *p*-values) between the Simpson's D index of diversity (simpson) and: (A) bacterial serovar and program (mlst vs. stringMLST with all kmer lengths); (B) bacterial serovar; (C) program (mlst vs. stringMLST with all kmer lengths); (D) the median number of contigs (num_contigs_median); (E) the mean total number of nucleotides (total_nucl_mean); (F) the mean GC% content originally calculated per genome (gc_avg_mean); (G) the mean total count of STs present in each generated database (st_count_mean); (H) the mean total count of unique alleles (across all 7 loci) present in each generated database (total_alleles_genes_mean); (I) the standard deviation (SD) of the number of contigs (num_contigs_sd); (J) the SD of the total number of nucleotides (total_nucl_sd); (K) the SD of the GC% content per genome (gc_avg_sd). The median number of contigs, mean total number of nucleotides, and mean GC% content were grouped by serovar and batch (experimental replicate). The SD of the number of contigs, SD of the total number of

nucleotides, and SD of GC% content were calculated by serovar only. The mean total count of STs and mean total count of unique alleles (across all 7 loci) present in each generated database were calculated after grouping by serovar, batch (three experimental replicates), and program. The Simpson's D index of diversity was calculated after grouping by program, serovar, and batch (three experimental replicates). All PERMANOVA models were run with 1,000 permutations.

**A.**

**PERMANOVA model using the proportion of non-classified STs as *dependent variable* and the following independent variables:**
***S. enterica* serovars (serovar) and program as MLST or stringMLST (program)**

Model = adonis(formula = prop ~ serovar * program, data = d18b, permutations = 1000)

| | Df | SumsOfSqs | MeanSqs | F.Model | $R^2$ | Pr(>F) |
|---|---|---|---|---|---|---|
| **serovar** | 19 | 26.8401628518863 | 1.41264015009928 | 37.1490819014527 | 0.353830505059659 | 0.000999000999000999 |
| **program** | 9 | 16.0133235978993 | 1.77925817754437 | 46.7902655582739 | 0.211101639270802 | 0.000999000999000999 |
| **serovar:program** | 163 | 20.8721297433185 | 0.12804987572588 | 3.36740770144276 | 0.275154672104759 | 0.000999000999000999 |
| **Residuals** | 319 | 12.1303726718492 | 0.038026246620217 | | 0.15991318356478 | |
| **Total** | 510 | 75.8559888649533 | | | 1 | |

**B.**

**PERMANOVA model using the proportion of non-classified STs as *dependent variable* and the following independent variable:**
***S. enterica* serovars (serovar)**

Model = adonis(formula = prop ~ serovar, data = d18b, permutations = 1000)

| | Df | SumsOfSqs | MeanSqs | F.Model | $R^2$ | Pr(>F) |
|---|---|---|---|---|---|---|
| **serovar** | 19 | 26.8401628518863 | 1.41264015009928 | 14.1506605134807 | 0.353830505059659 | 0.000999000999000999 |
| **Residuals** | 491 | 49.015826013067 | 0.0998285662180591 | | 0.646169494940341 | |
| **Total** | 510 | 75.8559888649533 | | | 1 | |

C.

**PERMANOVA model using the proportion of non-classified STs as *dependent variable* and the following independent variable:**
**program as MLST or stringMLST (program)**

Model = adonis(formula = prop ~ program, data = d18b, permutations = 1000)

| | Df | SumsOfSqs | MeanSqs | F.Model | $R^2$ | Pr(>F) |
|---|---|---|---|---|---|---|
| **program** | 9 | 15.6969648506825 | 1.74410720563139 | 14.5247986372593 | 0.206931121531194 | 0.000999000999000999 |
| **Residuals** | 501 | 60.1590240142707 | 0.120077892244053 | | 0.793068878468806 | |
| **Total** | 510 | 75.8559888649532 | | | 1 | |

**D.**

**PERMANOVA model using the proportion of non-classified STs as *dependent variable* and the following independent variable:**
**median of the number of contigs (num_contigs_median)**

Model = adonis(formula = prop ~ num_contigs_median, data = d18b, permutations = 1000)

| | Df | SumsOfSqs | MeanSqs | F.Model | $R^2$ | Pr(>F) |
|---|---|---|---|---|---|---|
| **num_contigs_median** | 1 | 0.391781222321214 | 0.391781222321214 | 2.64253277667546 | 0.00516480278200188 | 0.0609390609390609 |
| **Residuals** | 509 | 75.4642076426321 | 0.148259739965878 | | 0.994835197217998 | |
| **Total** | 510 | 75.8559888649533 | | | 1 | |

E.

**PERMANOVA model using the proportion of non-classified STs as *dependent variable* and the following independent variable:**
**mean of the total counts for nucleotides per genomes (total_nucl_mean)**

Model = adonis(formula = prop ~ total_nucl_mean, data = d18b, permutations = 1000)

| | Df | SumsOfSqs | MeanSqs | F.Model | $R^2$ | Pr(>F) |
|---|---|---|---|---|---|---|
| **total_nucl_mean** | 1 | 4.48323273411096 | 4.48323273411096 | 31.9725002279458 | 0.0591018955944596 | 0.000999000999000999 |
| **Residuals** | 509 | 71.3727561308423 | 0.140221524815014 | | 0.94089810440554 | |
| **Total** | 510 | 75.8559888649533 | | | 1 | |

**F.**

**PERMANOVA model using the proportion of non-classified STs as *dependent variable* and the following independent variable:**
**mean of the average GC% per genome (gc_avg_mean)**

Model = adonis(formula = prop ~ gc_avg_mean, data = d18b, permutations = 1000)

| | Df | SumsOfSqs | MeanSqs | F.Model | $R^2$ | Pr(>F) |
|---|---|---|---|---|---|---|
| **gc_avg_mean** | 1 | 0.291982834228857 | 0.291982834228857 | 1.96679967658226 | 0.00384917313184956 | 0.130869130869131 |
| **Residuals** | 509 | 75.5640060307244 | 0.148455807525981 | | 0.99615082686815 | |
| **Total** | 510 | 75.8559888649533 | | | 1 | |

**G.**

**PERMANOVA model using the proportion of non-classified STs as *dependent variable* and the following independent variable:**
**mean of the total counts of unique STs per program (st_count_mean)**

Model = adonis(formula = prop ~ st_count_mean, data = d18b, permutations = 1000)

| | Df | SumsOfSqs | MeanSqs | F.Model | $R^2$ | Pr(>F) |
|---|---|---|---|---|---|---|
| **st_count_mean** | 1 | 6.6959987945063 | 6.6959987945063 | 49.2808541894239 | 0.0882725134125825 | 0.000999000999000999 |
| **Residuals** | 509 | 69.159990070447 | 0.135874243753334 | | 0.911727486587418 | |
| **Total** | 510 | 75.8559888649533 | | | 1 | |

## H.

**PERMANOVA model using the proportion of non-classified STs as *dependent variable* and the following independent variable:**
**mean of the total counts of unique alleles across all genes per program (total_alleles_genes_mean)**

Model = adonis(formula = prop ~ total_alleles_genes_mean, data = d18b, permutations = 1000)

| | Df | SumsOfSqs | MeanSqs | F.Model | $R^2$ | Pr(>F) |
|---|---|---|---|---|---|---|
| **total_alleles_genes_mean** | 1 | 6.69599879450637 | 6.69599879450637 | 49.2808541894246 | 0.0882725134125835 | 0.000999000999000999 |
| **Residuals** | 509 | 69.1599900704469 | 0.135874243753334 | | 0.911727486587416 | |
| **Total** | 510 | 75.8559888649533 | | | 1 | |

I.

**PERMANOVA model using the proportion of non-classified STs as *dependent variable* and the following independent variable:**
**Simpson's D index of diversity per *S. enterica* serovar (simpson)**

Model = adonis(formula = prop ~ simpson, data = d18b, permutations = 1000)

| | Df | SumsOfSqs | MeanSqs | F.Model | $R^2$ | Pr(>F) |
|---|---|---|---|---|---|---|
| **simpson** | 1 | 1.1819423928987 | 1.1819423928987 | 8.05646280613147 | 0.0155813985234959 | 0.000999000999000999 |
| **Residuals** | 509 | 74.6740464720546 | 0.146707360455903 | | 0.984418601476504 | |
| **Total** | 510 | 75.8559888649533 | | | 1 | |

**J.**

**PPERMANOVA model using the proportion of non-classified STs as *dependent variable* and the following independent variable:**
**Standard deviation of the number of contigs (num_contigs_sd)**

Model = adonis(formula = prop ~ num_contigs_sd, data = d18c, permutations = 1000)

| | Df | SumsOfSqs | MeanSqs | F.Model | $R^2$ | Pr(>F) |
|---|---|---|---|---|---|---|
| **num_contigs_sd** | 1 | 0.320546129525937 | 0.320546129525937 | 2.16001884704885 | 0.00422571947610632 | 0.115884115884116 |
| **Residuals** | 509 | 75.5354427354273 | 0.148399691032274 | | 0.995774280523894 | |
| **Total** | 510 | 75.8559888649533 | | | 1 | |

**K.**

**PERMANOVA model using the proportion of non-classified STs as *dependent variable* and the following independent variable:**
**Standard deviation of the total counts for nucleotides per genomes (total_nucl_sd)**

Model = adonis(formula = prop ~ total_nucl_sd, data = d18c, permutations = 1000)

| | Df | SumsOfSqs | MeanSqs | F.Model | $R^2$ | Pr(>F) |
|---|---|---|---|---|---|---|
| **total_nucl_sd** | 1 | 0.54987638138005 | 0.54987638138005 | 3.71665816879736 | 0.00724895146194715 | 0.043956043956044 |
| **Residuals** | 509 | 75.3061124835732 | 0.14794914043924 | | 0.992751048538053 | |
| **Total** | 510 | 75.8559888649533 | | | 1 | |

**L.**

**PermPERMANOVA model using the proportion of non-classified STs as *dependent variable* and the following independent variable:**
**Standard deviation of the average GC% per genome (gc_avg_sd)**

Model = adonis(formula = prop ~ gc_avg_sd, data = d18c, permutations = 1000)

| | Df | SumsOfSqs | MeanSqs | F.Model | $R^2$ | Pr(>F) |
|---|---|---|---|---|---|---|
| **gc_avg_sd** | 1 | 0.332391331936203 | 0.332391331936203 | 2.24018973515612 | 0.00438187329583113 | 0.0929070929070929 |
| **Residuals** | 509 | 75.5235975330171 | 0.148376419514768 | | 0.995618126704169 | |
| **Total** | 510 | 75.8559888649533 | | | 1 | |

Figure S12. PERMANOVA results measuring the association between *S. enterica* serovars (serovar), program, or genome-intrinsic and –extrinsic variables and the proportion of non-classified STs.
PERMANOVA results demonstrating the association (*R*-squared and *p*-values) between the proportion of non-classified STs (prop) with: (A) bacterial serovar and program (mlst vs. stringMLST with all kmer lengths); (B) bacterial serovar; (C) program (mlst vs. stringMLST with all kmer lengths); (D) the median number of contigs (num_contigs_median); (E) the mean total number of nucleotides (total_nucl_mean); (F) the mean GC% content originally calculated per genome (gc_avg_mean); (G) the mean total count of STs present in each generated database (st_count_mean); (H) the mean total count of unique alleles (across all 7 loci) present in each generated database (total_alleles_genes_mean); (I) the Simpson's D index of diversity (simpson); (J) the standard deviation (SD) of the number of contigs (num_contigs_sd); (K) the SD of the total number of nucleotides (total_nucl_sd); (L) the SD of the GC% content per genome (gc_avg_sd). The median number of contigs, mean total number of nucleotides, and mean GC% content were grouped by serovar and batch (experimental replicate). The SD of the

number of contigs, SD of the total number of nucleotides, and SD of GC% content were calculated by serovar only. The mean total count of STs and mean total count of unique alleles (across all 7 loci) present in each generated database were calculated after grouping by serovar, batch (three experimental replicates), and program. The Simpson's D index of diversity was calculated after grouping by program, serovar, and batch (three experimental replicates). All PERMANOVA models were run with 1,000 permutations.

A.

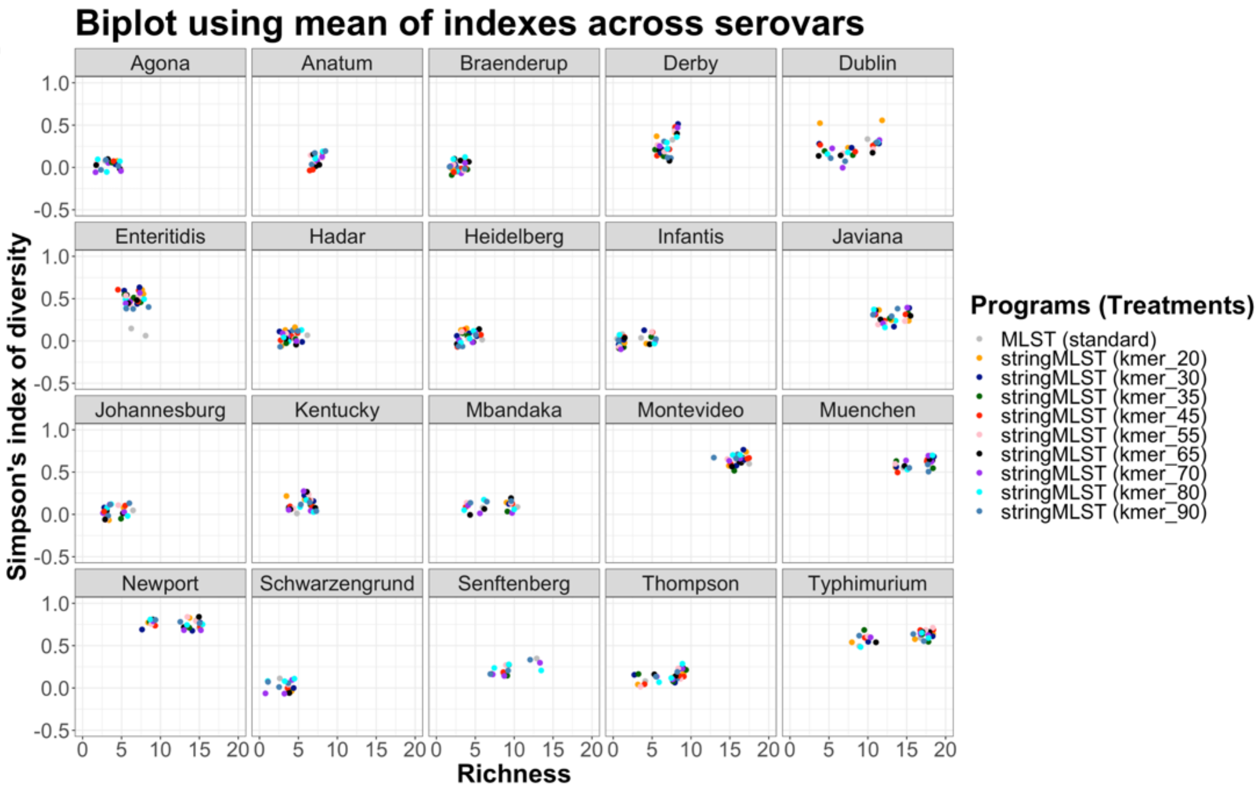
Biplot using mean of indexes across serovars
Agona
Anatum
Braenderup
Derby
Dublin
Enteritidis
Hadar
Heidelberg
Infantis
Javiana
Johannesburg
Kentucky
Mbandaka
Montevideo
Muenchen
Newport
Schwarzengrund
Senftenberg
Thompson
Typhimurium
Simpson's index of diversity
Richness
1.0
0.5
0.0
-0.5
0
5
10
15
20
Programs (Treatments)
MLST (standard)
stringMLST (kmer_20)
stringMLST (kmer_30)
stringMLST (kmer_35)
stringMLST (kmer_45)
stringMLST (kmer_55)
stringMLST (kmer_65)
stringMLST (kmer_70)
stringMLST (kmer_80)
stringMLST (kmer_90)

B.

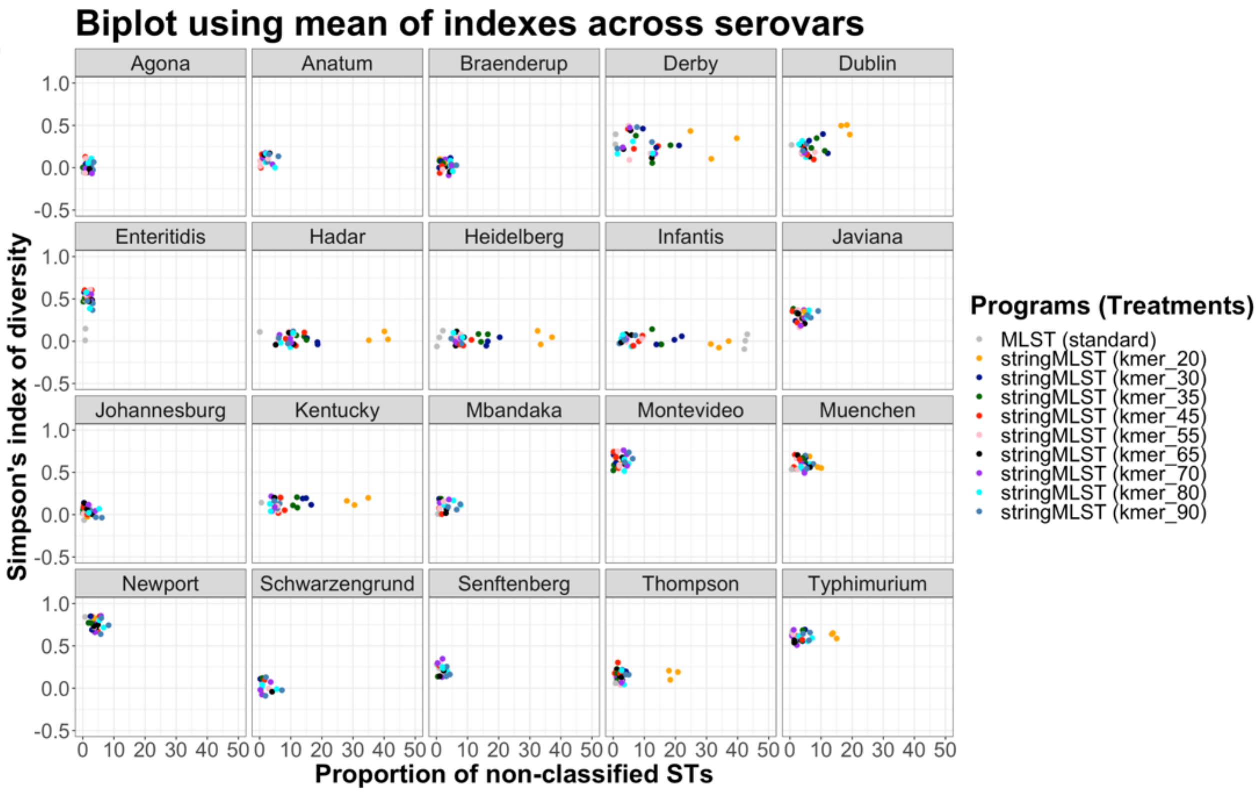

Biplot using mean of indexes across serovars
Agona
Anatum
Braenderup
Derby
Dublin
Enteritidis
Hadar
Heidelberg
Infantis
Javiana
Johannesburg
Kentucky
Mbandaka
Montevideo
Muenchen
Newport
Schwarzengrund
Senftenberg
Thompson
Typhimurium
1.0
0.5
0.0
-0.5
0 10 20 30 40 50
Simpson's index of diversity
Proportion of non-classified STs
Programs (Treatments)
MLST (standard)
stringMLST (kmer_20)
stringMLST (kmer_30)
stringMLST (kmer_35)
stringMLST (kmer_45)
stringMLST (kmer_55)
stringMLST (kmer_65)
stringMLST (kmer_70)
stringMLST (kmer_80)
stringMLST (kmer_90)

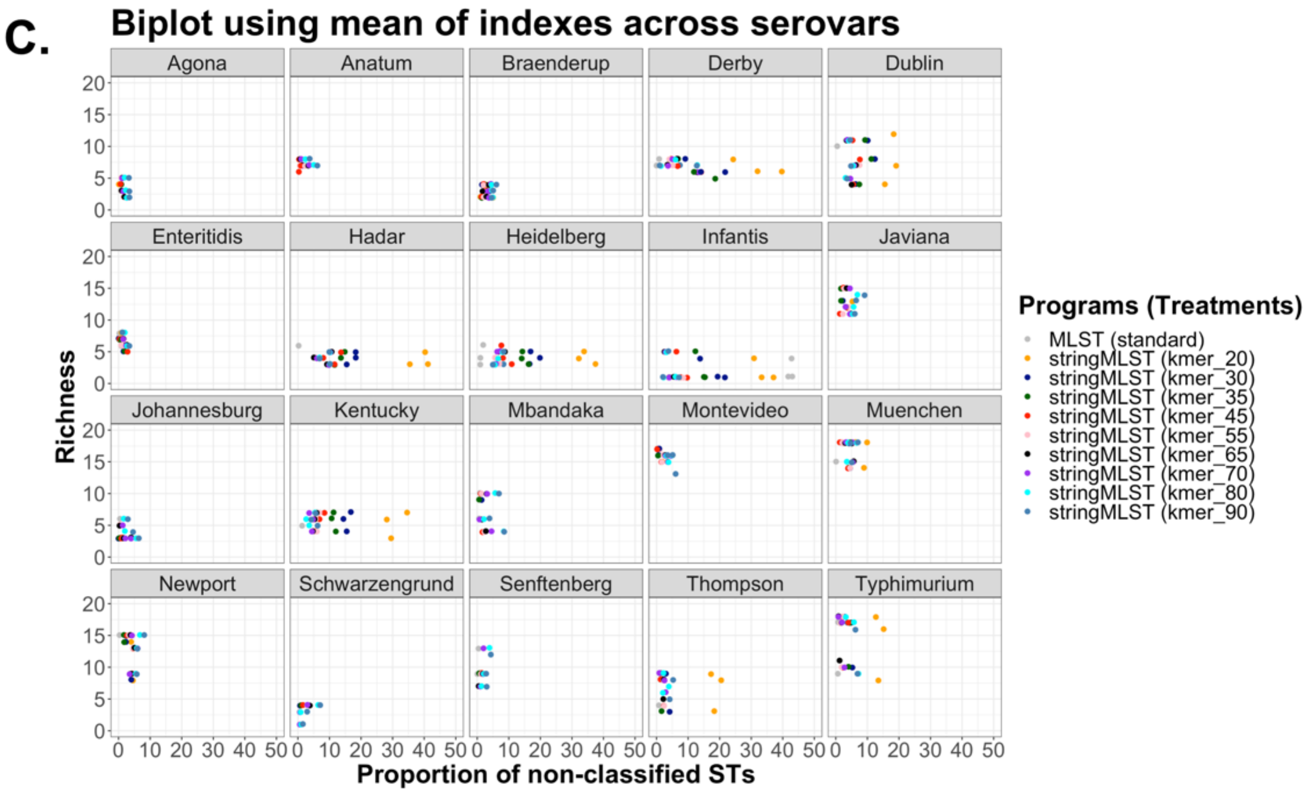
C.
Biplot using mean of indexes across serovars
Agona
Anatum
Braenderup
Derby
Dublin
Enteritidis
Hadar
Heidelberg
Infantis
Javiana
Johannesburg
Kentucky
Mbandaka
Montevideo
Muenchen
Newport
Schwarzengrund
Senftenberg
Thompson
Typhimurium
Richness
Proportion of non-classified STs
Programs (Treatments)
MLST (standard)
stringMLST (kmer_20)
stringMLST (kmer_30)
stringMLST (kmer_35)
stringMLST (kmer_45)
stringMLST (kmer_55)
stringMLST (kmer_65)
stringMLST (kmer_70)
stringMLST (kmer_80)
stringMLST (kmer_90)

D.

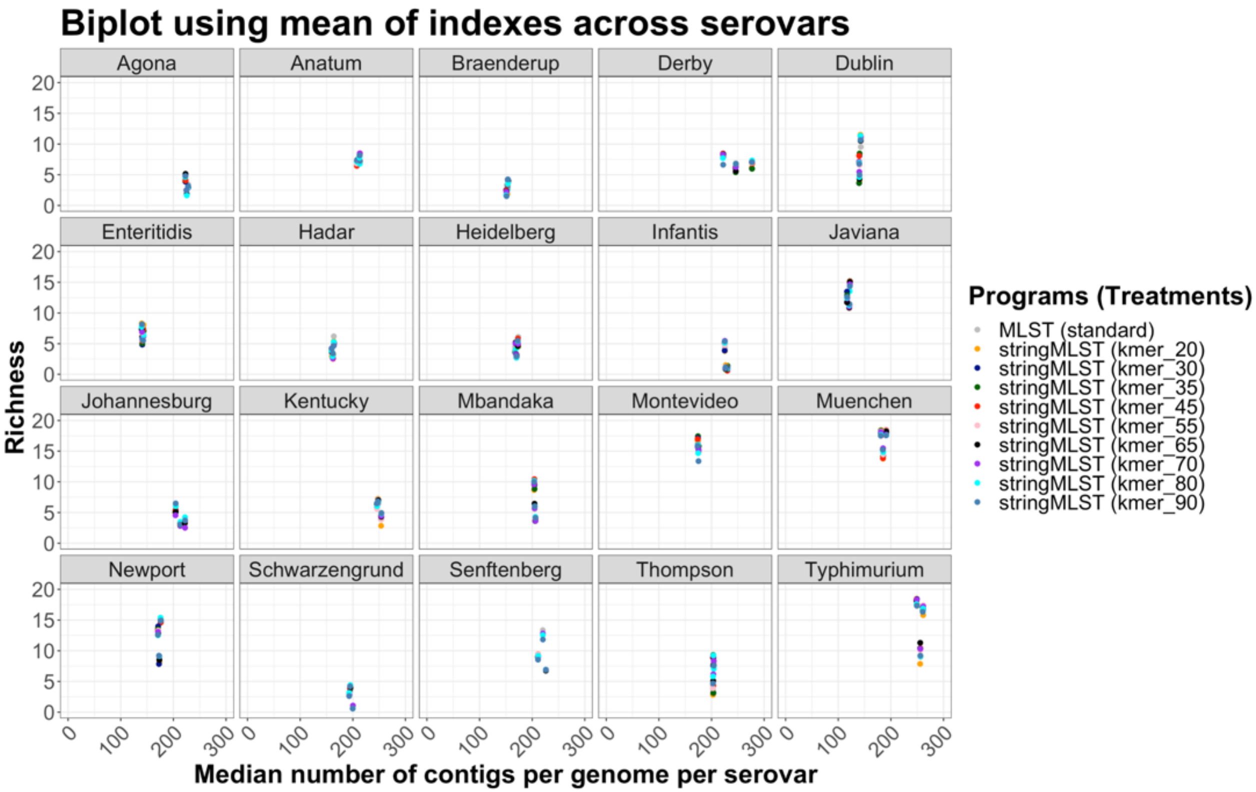

Biplot using mean of indexes across serovars
Agona
Anatum
Braenderup
Derby
Dublin
Enteritidis
Hadar
Heidelberg
Infantis
Javiana
Johannesburg
Kentucky
Mbandaka
Montevideo
Muenchen
Newport
Schwarzengrund
Senftenberg
Thompson
Typhimurium
Richness
Median number of contigs per genome per serovar
0
5
10
15
20
0
100
200
300
Programs (Treatments)
MLST (standard)
stringMLST (kmer_20)
stringMLST (kmer_30)
stringMLST (kmer_35)
stringMLST (kmer_45)
stringMLST (kmer_55)
stringMLST (kmer_65)
stringMLST (kmer_70)
stringMLST (kmer_80)
stringMLST (kmer_90)

**E.**

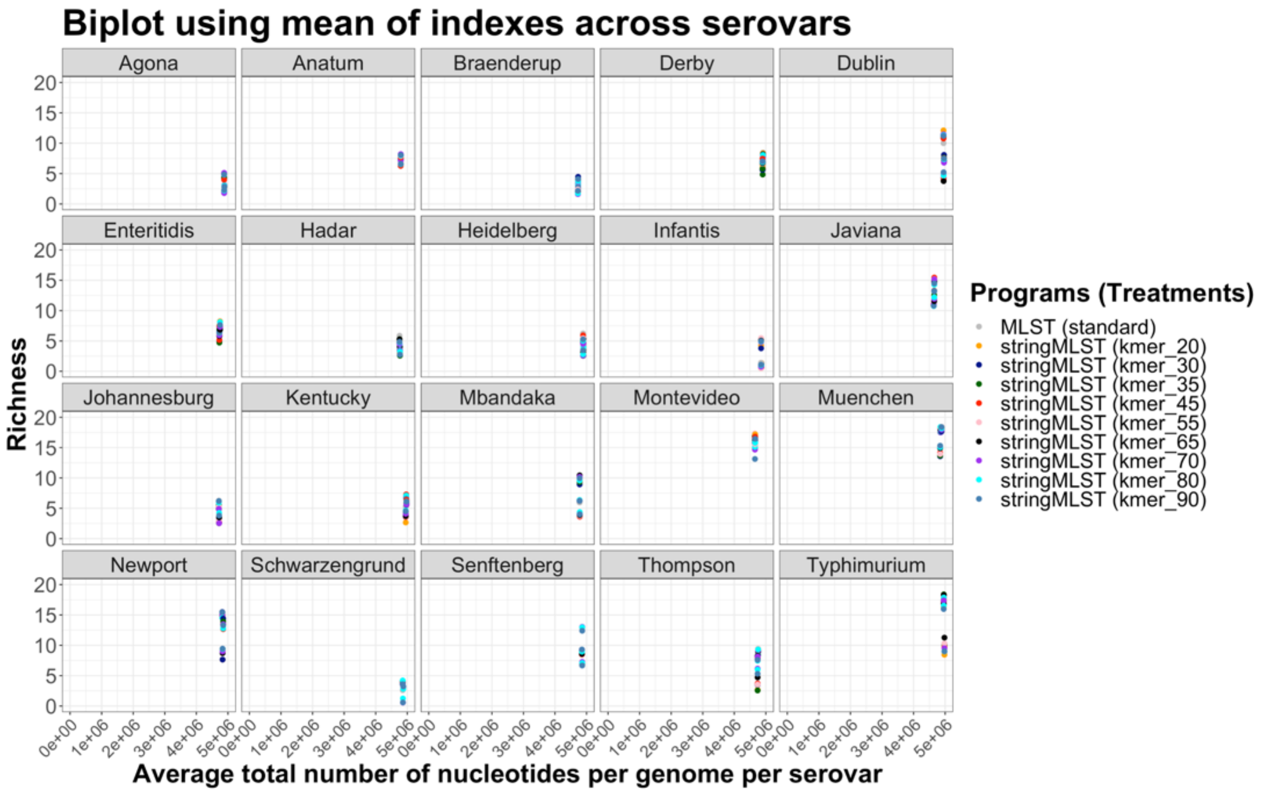

Biplot using mean of indexes across serovars
Agona
Anatum
Braenderup
Derby
Dublin
Enteritidis
Hadar
Heidelberg
Infantis
Javiana
Johannesburg
Kentucky
Mbandaka
Montevideo
Muenchen
Newport
Schwarzengrund
Senftenberg
Thompson
Typhimurium
Richness
0
5
10
15
20
0e+00
1e+06
2e+06
3e+06
4e+06
5e+06
Average total number of nucleotides per genome per serovar
Programs (Treatments)
MLST (standard)
stringMLST (kmer_20)
stringMLST (kmer_30)
stringMLST (kmer_35)
stringMLST (kmer_45)
stringMLST (kmer_55)
stringMLST (kmer_65)
stringMLST (kmer_70)
stringMLST (kmer_80)
stringMLST (kmer_90)

F.

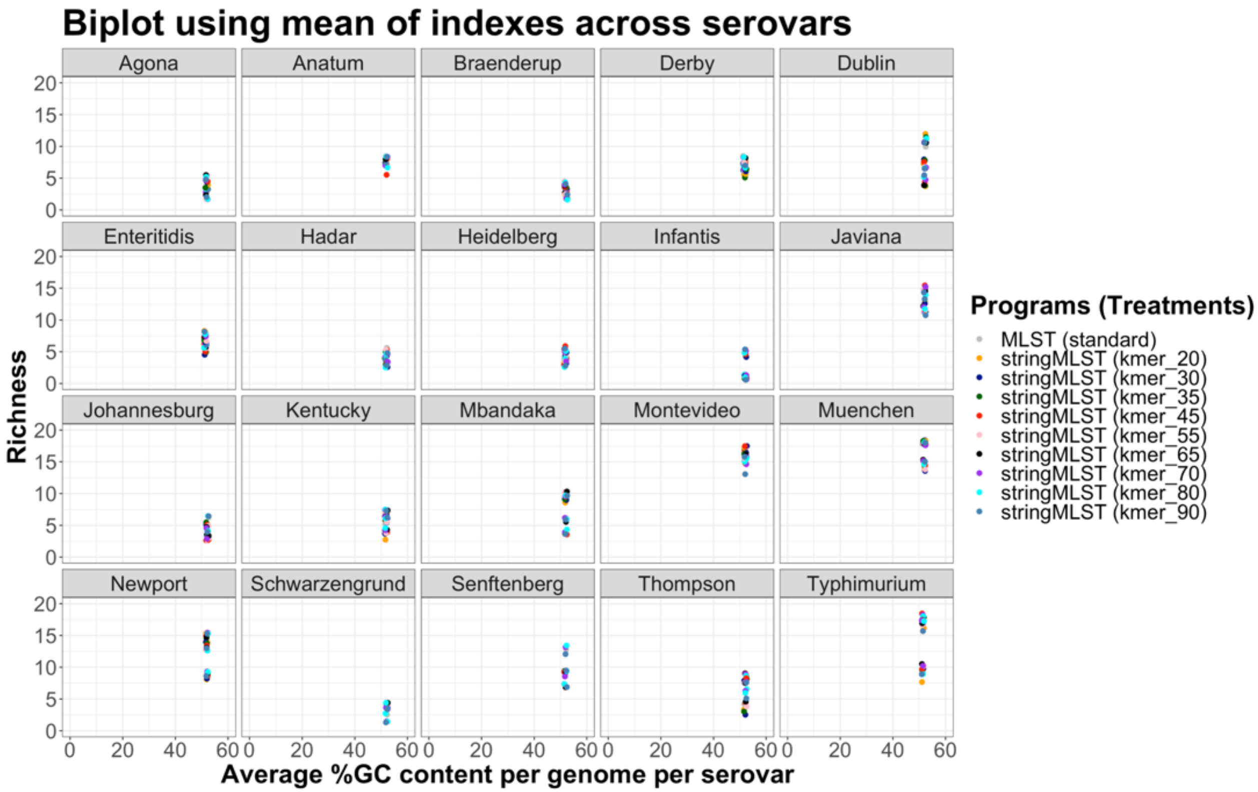
Biplot using mean of indexes across serovars
Agona
Anatum
Braenderup
Derby
Dublin
Enteritidis
Hadar
Heidelberg
Infantis
Javiana
Johannesburg
Kentucky
Mbandaka
Montevideo
Muenchen
Newport
Schwarzengrund
Senftenberg
Thompson
Typhimurium
Richness
Average %GC content per genome per serovar
Programs (Treatments)
MLST (standard)
stringMLST (kmer_20)
stringMLST (kmer_30)
stringMLST (kmer_35)
stringMLST (kmer_45)
stringMLST (kmer_55)
stringMLST (kmer_65)
stringMLST (kmer_70)
stringMLST (kmer_80)
stringMLST (kmer_90)

G.

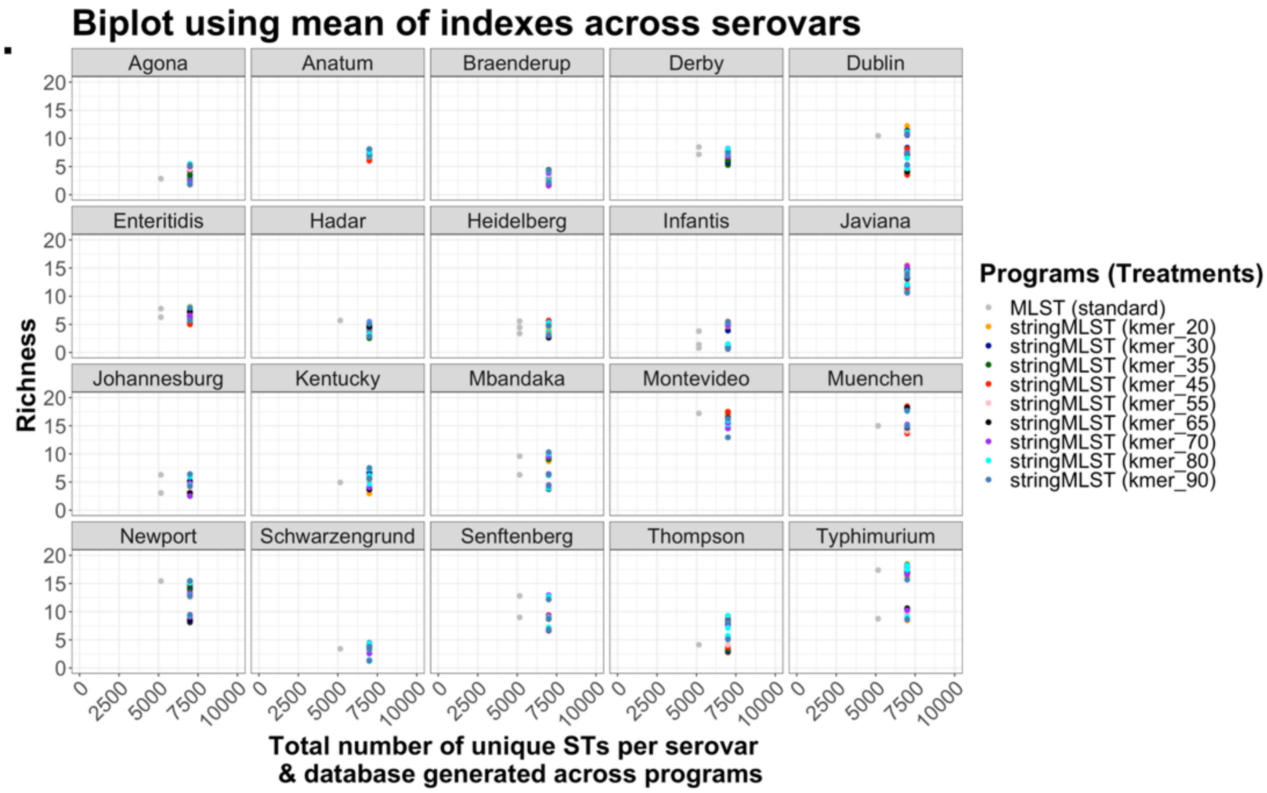
Biplot using mean of indexes across serovars
Agona
Anatum
Braenderup
Derby
Dublin
Enteritidis
Hadar
Heidelberg
Infantis
Javiana
Johannesburg
Kentucky
Mbandaka
Montevideo
Muenchen
Newport
Schwarzengrund
Senftenberg
Thompson
Typhimurium
Richness
0
5
10
15
20
0
2500
5000
7500
10000
Total number of unique STs per serovar
& database generated across programs
Programs (Treatments)
MLST (standard)
stringMLST (kmer_20)
stringMLST (kmer_30)
stringMLST (kmer_35)
stringMLST (kmer_45)
stringMLST (kmer_55)
stringMLST (kmer_65)
stringMLST (kmer_70)
stringMLST (kmer_80)
stringMLST (kmer_90)

**H.**

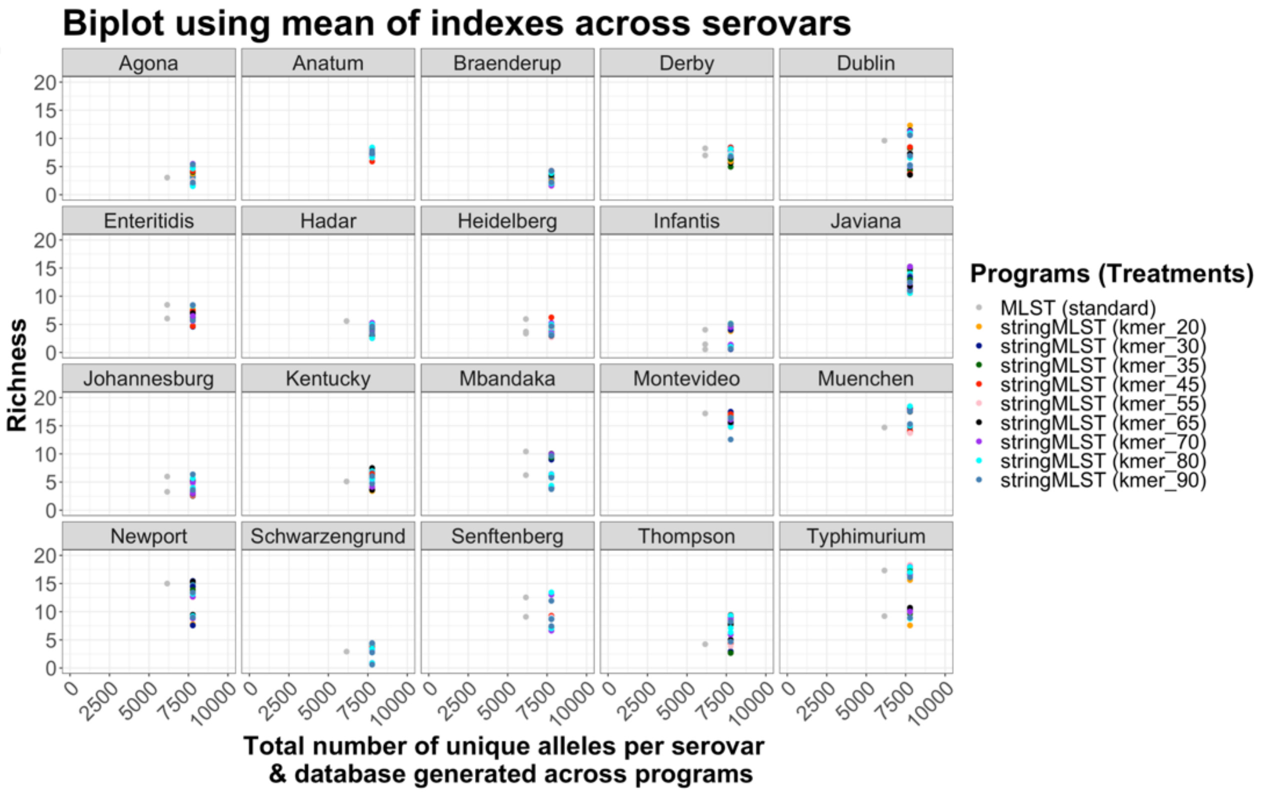
Biplot using mean of indexes across serovars
Agona
Anatum
Braenderup
Derby
Dublin
Enteritidis
Hadar
Heidelberg
Infantis
Javiana
Johannesburg
Kentucky
Mbandaka
Montevideo
Muenchen
Newport
Schwarzengrund
Senftenberg
Thompson
Typhimurium
Richness
0
5
10
15
20
0
2500
5000
7500
10000
Total number of unique alleles per serovar & database generated across programs
Programs (Treatments)
MLST (standard)
stringMLST (kmer_20)
stringMLST (kmer_30)
stringMLST (kmer_35)
stringMLST (kmer_45)
stringMLST (kmer_55)
stringMLST (kmer_65)
stringMLST (kmer_70)
stringMLST (kmer_80)
stringMLST (kmer_90)

I.

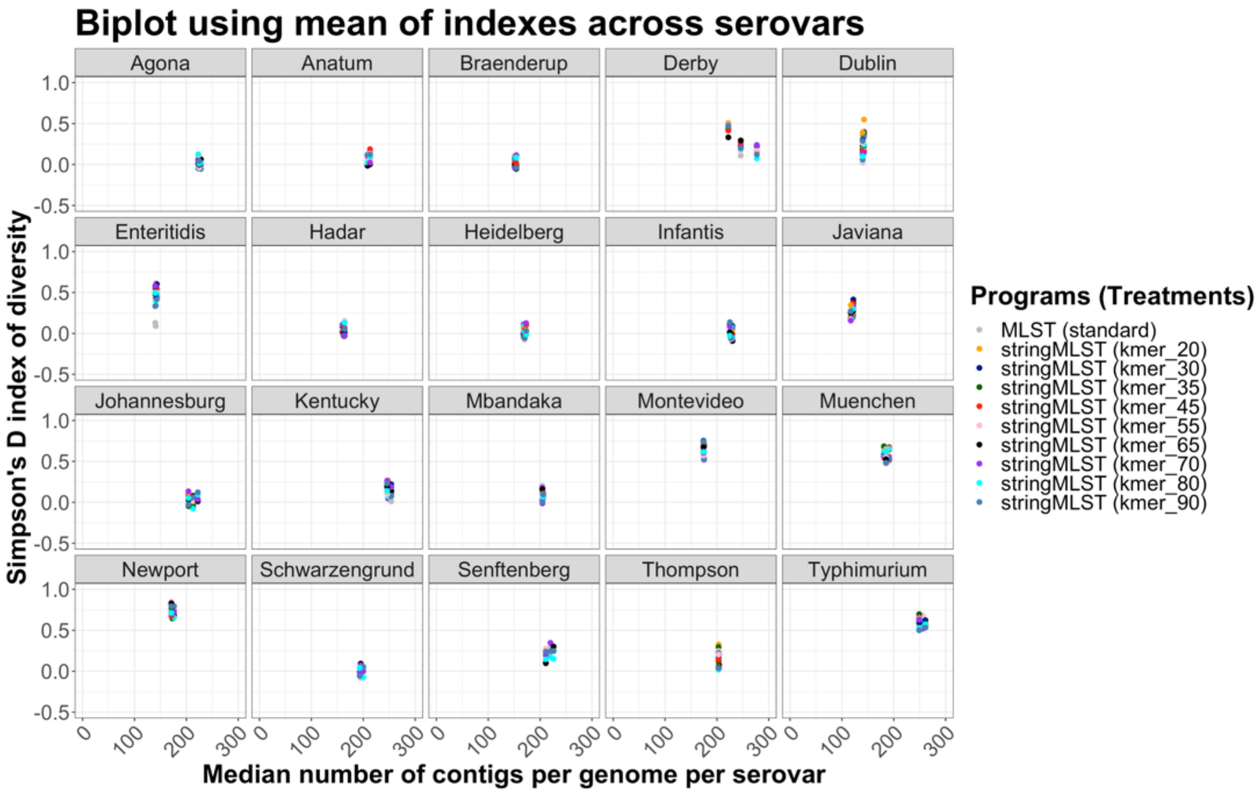
Biplot using mean of indexes across serovars
Agona
Anatum
Braenderup
Derby
Dublin
Enteritidis
Hadar
Heidelberg
Infantis
Javiana
Johannesburg
Kentucky
Mbandaka
Montevideo
Muenchen
Newport
Schwarzengrund
Senftenberg
Thompson
Typhimurium
1.0
0.5
0.0
-0.5
0
100
200
300
Simpson's D index of diversity
Median number of contigs per genome per serovar
Programs (Treatments)
MLST (standard)
stringMLST (kmer_20)
stringMLST (kmer_30)
stringMLST (kmer_35)
stringMLST (kmer_45)
stringMLST (kmer_55)
stringMLST (kmer_65)
stringMLST (kmer_70)
stringMLST (kmer_80)
stringMLST (kmer_90)

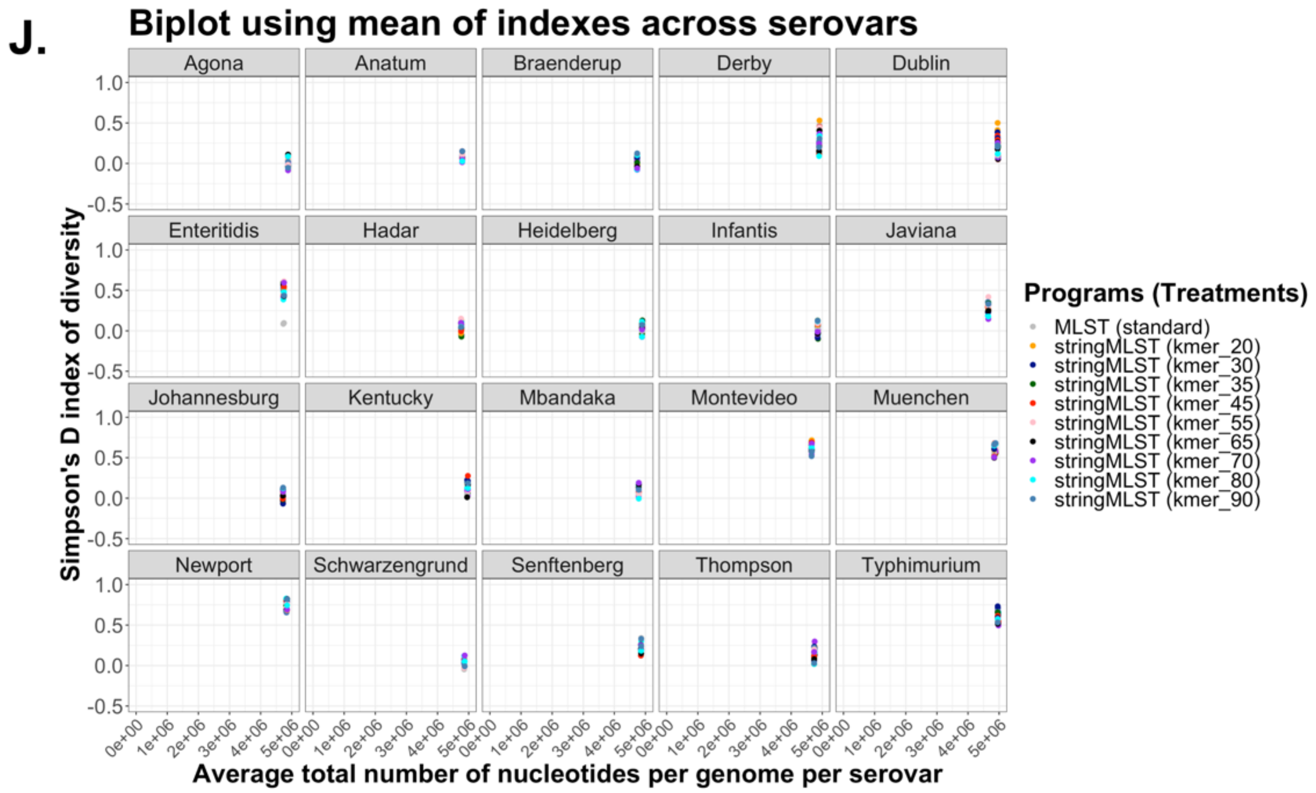
J.
Biplot using mean of indexes across serovars
Agona
Anatum
Braenderup
Derby
Dublin
Enteritidis
Hadar
Heidelberg
Infantis
Javiana
Johannesburg
Kentucky
Mbandaka
Montevideo
Muenchen
Newport
Schwarzengrund
Senftenberg
Thompson
Typhimurium
Simpson's D index of diversity
Average total number of nucleotides per genome per serovar
1.0
0.5
0.0
-0.5
0e+00
1e+06
2e+06
3e+06
4e+06
5e+06
Programs (Treatments)
MLST (standard)
stringMLST (kmer_20)
stringMLST (kmer_30)
stringMLST (kmer_35)
stringMLST (kmer_45)
stringMLST (kmer_55)
stringMLST (kmer_65)
stringMLST (kmer_70)
stringMLST (kmer_80)
stringMLST (kmer_90)

K.

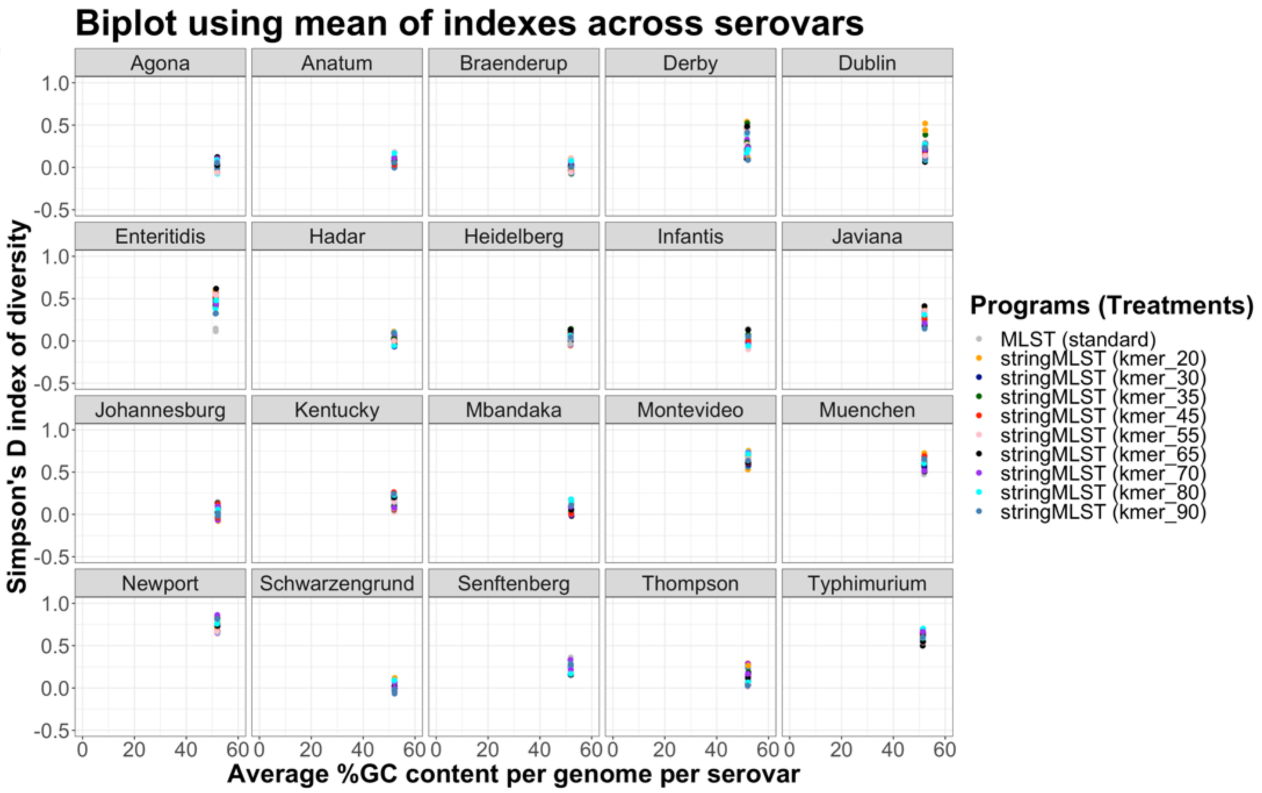
Biplot using mean of indexes across serovars
Agona
Anatum
Braenderup
Derby
Dublin
Enteritidis
Hadar
Heidelberg
Infantis
Javiana
Johannesburg
Kentucky
Mbandaka
Montevideo
Muenchen
Newport
Schwarzengrund
Senftenberg
Thompson
Typhimurium
Simpson's D index of diversity
Average %GC content per genome per serovar
1.0
0.5
0.0
-0.5
0
20
40
60
Programs (Treatments)
MLST (standard)
stringMLST (kmer_20)
stringMLST (kmer_30)
stringMLST (kmer_35)
stringMLST (kmer_45)
stringMLST (kmer_55)
stringMLST (kmer_65)
stringMLST (kmer_70)
stringMLST (kmer_80)
stringMLST (kmer_90)

L.

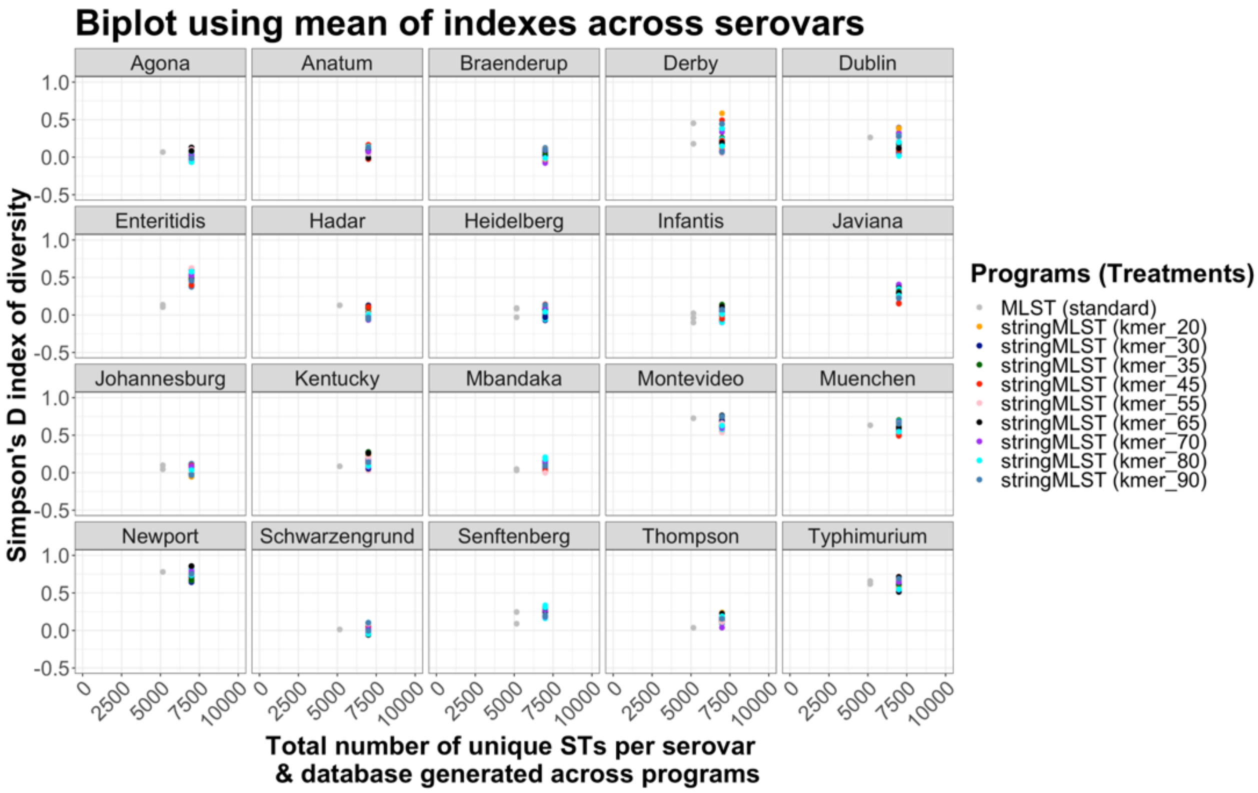
Biplot using mean of indexes across serovars
Agona
Anatum
Braenderup
Derby
Dublin
Enteritidis
Hadar
Heidelberg
Infantis
Javiana
Johannesburg
Kentucky
Mbandaka
Montevideo
Muenchen
Newport
Schwarzengrund
Senftenberg
Thompson
Typhimurium
1.0
0.5
0.0
-0.5
0
2500
5000
7500
10000
Simpson's D index of diversity
Total number of unique STs per serovar
& database generated across programs
Programs (Treatments)
MLST (standard)
stringMLST (kmer_20)
stringMLST (kmer_30)
stringMLST (kmer_35)
stringMLST (kmer_45)
stringMLST (kmer_55)
stringMLST (kmer_65)
stringMLST (kmer_70)
stringMLST (kmer_80)
stringMLST (kmer_90)

**M.**

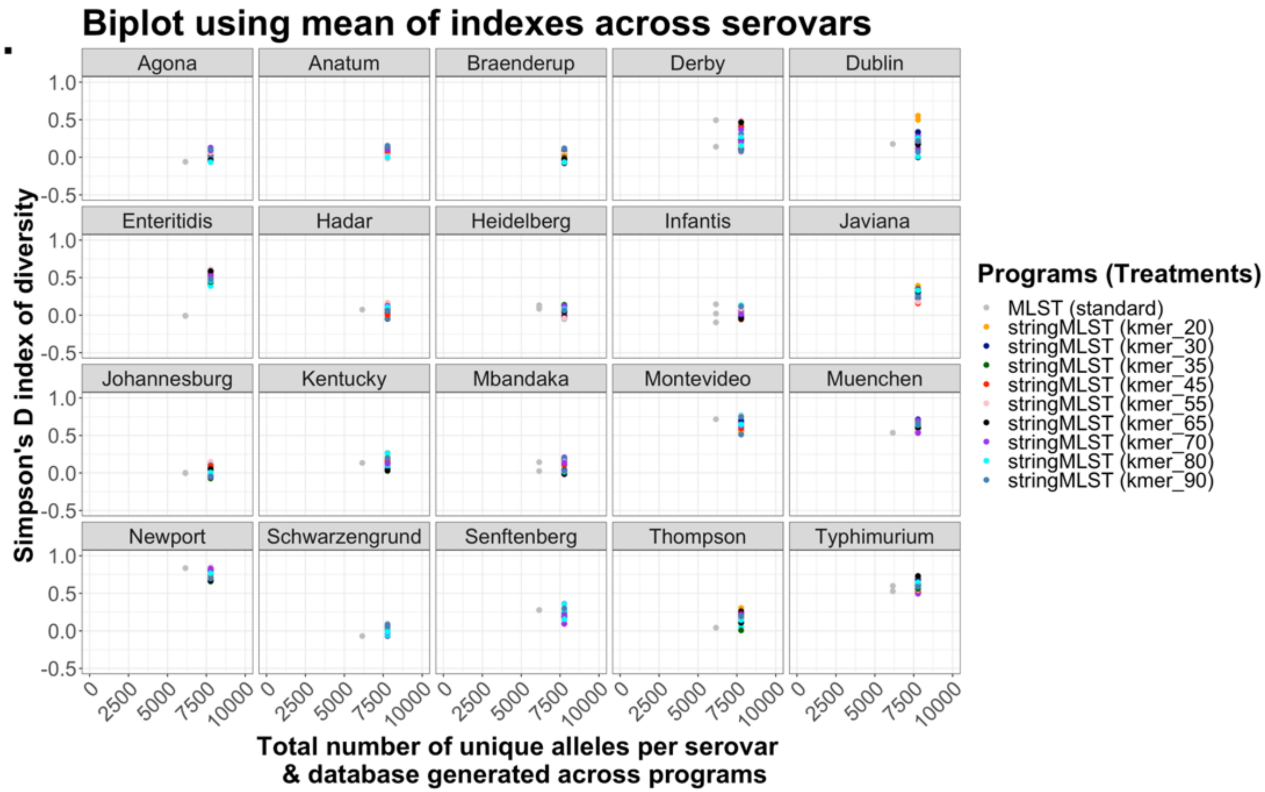
Biplot using mean of indexes across serovars
Agona
Anatum
Braenderup
Derby
Dublin
Enteritidis
Hadar
Heidelberg
Infantis
Javiana
Johannesburg
Kentucky
Mbandaka
Montevideo
Muenchen
Newport
Schwarzengrund
Senftenberg
Thompson
Typhimurium
Simpson's D index of diversity
1.0
0.5
0.0
-0.5
0
2500
5000
7500
10000
Total number of unique alleles per serovar & database generated across programs
Programs (Treatments)
MLST (standard)
stringMLST (kmer_20)
stringMLST (kmer_30)
stringMLST (kmer_35)
stringMLST (kmer_45)
stringMLST (kmer_55)
stringMLST (kmer_65)
stringMLST (kmer_70)
stringMLST (kmer_80)
stringMLST (kmer_90)

N.

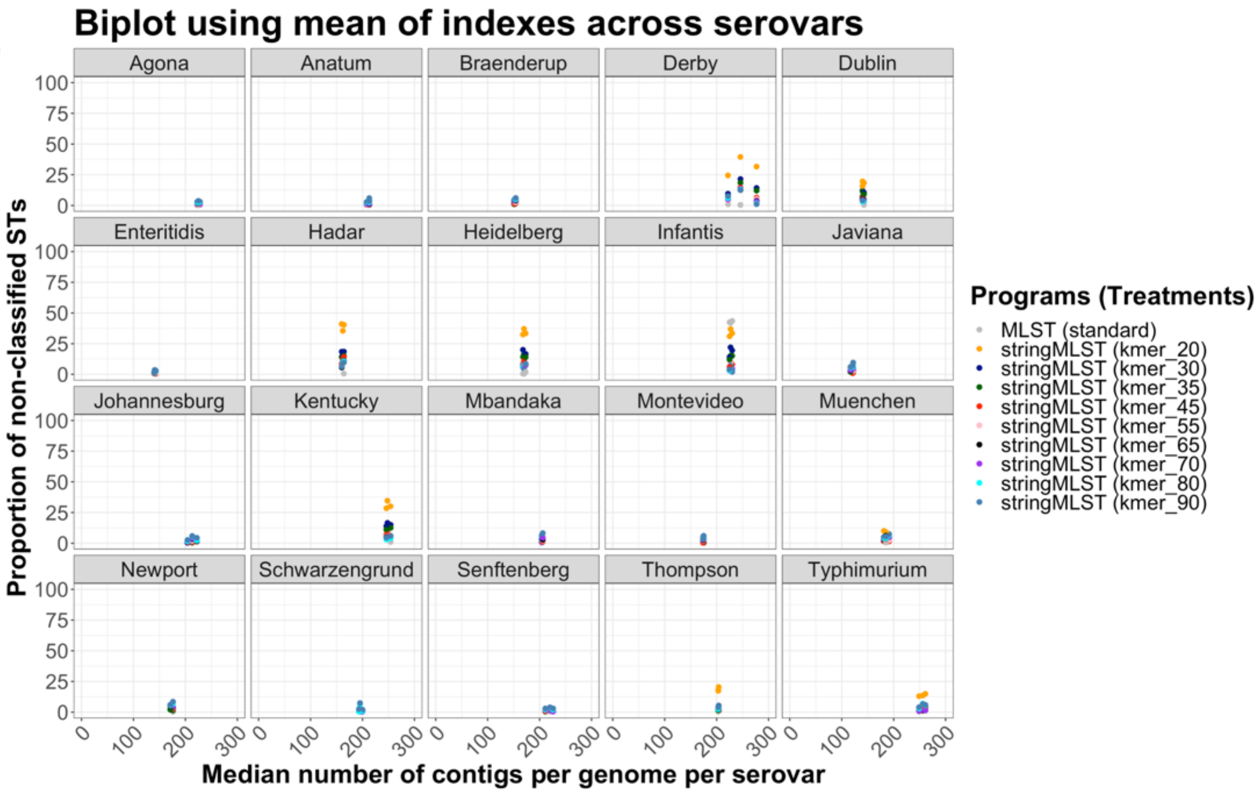
Biplot using mean of indexes across serovars
Agona
Anatum
Braenderup
Derby
Dublin
Enteritidis
Hadar
Heidelberg
Infantis
Javiana
Johannesburg
Kentucky
Mbandaka
Montevideo
Muenchen
Newport
Schwarzengrund
Senftenberg
Thompson
Typhimurium
Proportion of non-classified STs
Median number of contigs per genome per serovar
0
25
50
75
100
0
100
200
300
Programs (Treatments)
MLST (standard)
stringMLST (kmer_20)
stringMLST (kmer_30)
stringMLST (kmer_35)
stringMLST (kmer_45)
stringMLST (kmer_55)
stringMLST (kmer_65)
stringMLST (kmer_70)
stringMLST (kmer_80)
stringMLST (kmer_90)

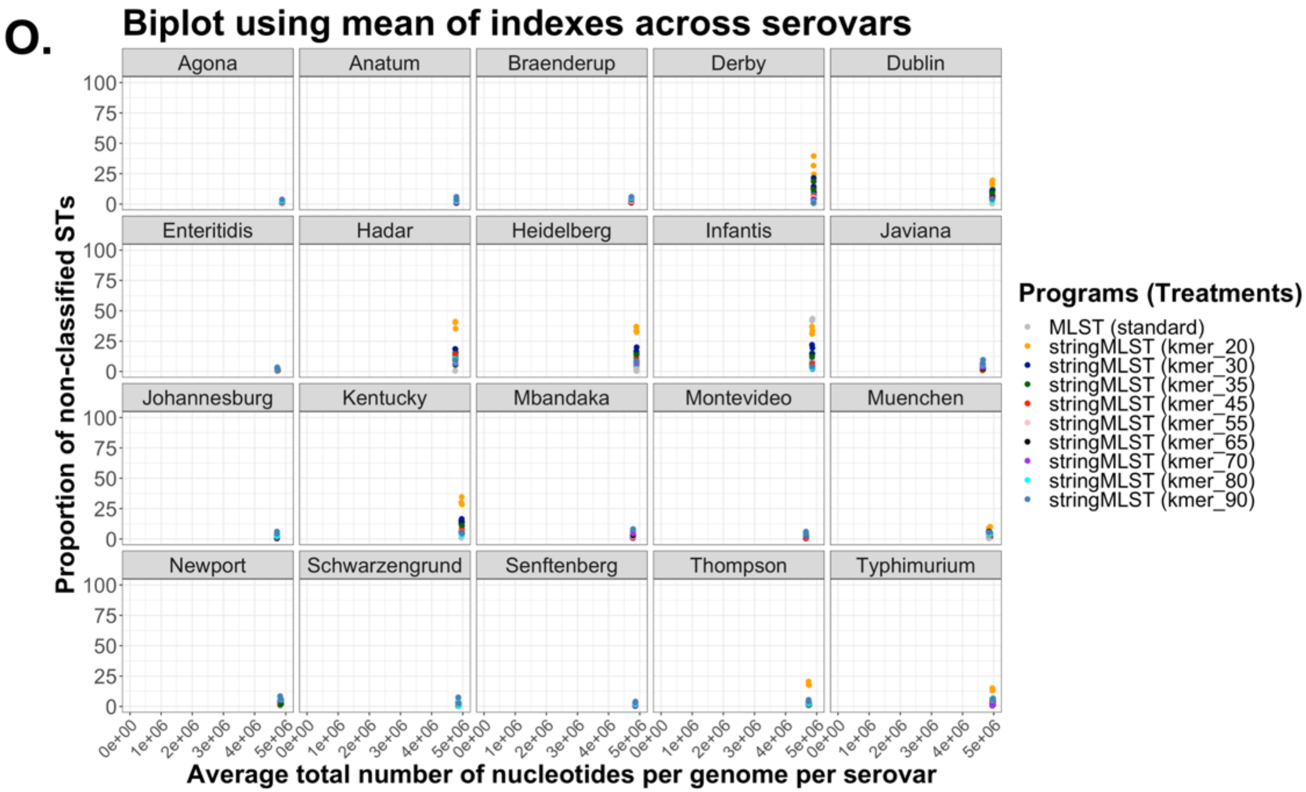
O.
Biplot using mean of indexes across serovars
Agona
Anatum
Braenderup
Derby
Dublin
Enteritidis
Hadar
Heidelberg
Infantis
Javiana
Johannesburg
Kentucky
Mbandaka
Montevideo
Muenchen
Newport
Schwarzengrund
Senftenberg
Thompson
Typhimurium
Proportion of non-classified STs
Average total number of nucleotides per genome per serovar
0e+00 1e+06 2e+06 3e+06 4e+06 5e+06
0 25 50 75 100
Programs (Treatments)
MLST (standard)
stringMLST (kmer_20)
stringMLST (kmer_30)
stringMLST (kmer_35)
stringMLST (kmer_45)
stringMLST (kmer_55)
stringMLST (kmer_65)
stringMLST (kmer_70)
stringMLST (kmer_80)
stringMLST (kmer_90)

P.

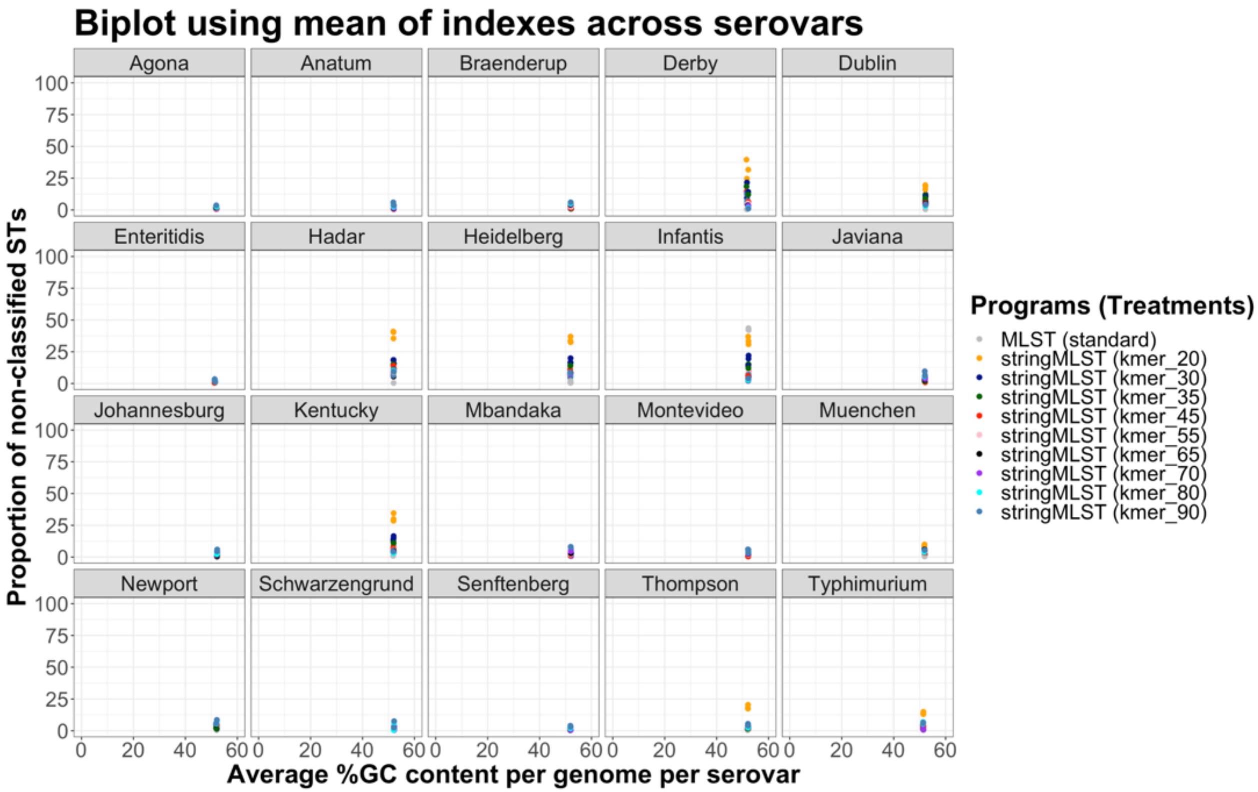
Biplot using mean of indexes across serovars
Agona
Anatum
Braenderup
Derby
Dublin
Enteritidis
Hadar
Heidelberg
Infantis
Javiana
Johannesburg
Kentucky
Mbandaka
Montevideo
Muenchen
Newport
Schwarzengrund
Senftenberg
Thompson
Typhimurium
Proportion of non-classified STs
Average %GC content per genome per serovar
0
20
40
60
25
50
75
100
Programs (Treatments)
MLST (standard)
stringMLST (kmer_20)
stringMLST (kmer_30)
stringMLST (kmer_35)
stringMLST (kmer_45)
stringMLST (kmer_55)
stringMLST (kmer_65)
stringMLST (kmer_70)
stringMLST (kmer_80)
stringMLST (kmer_90)

**Q.**

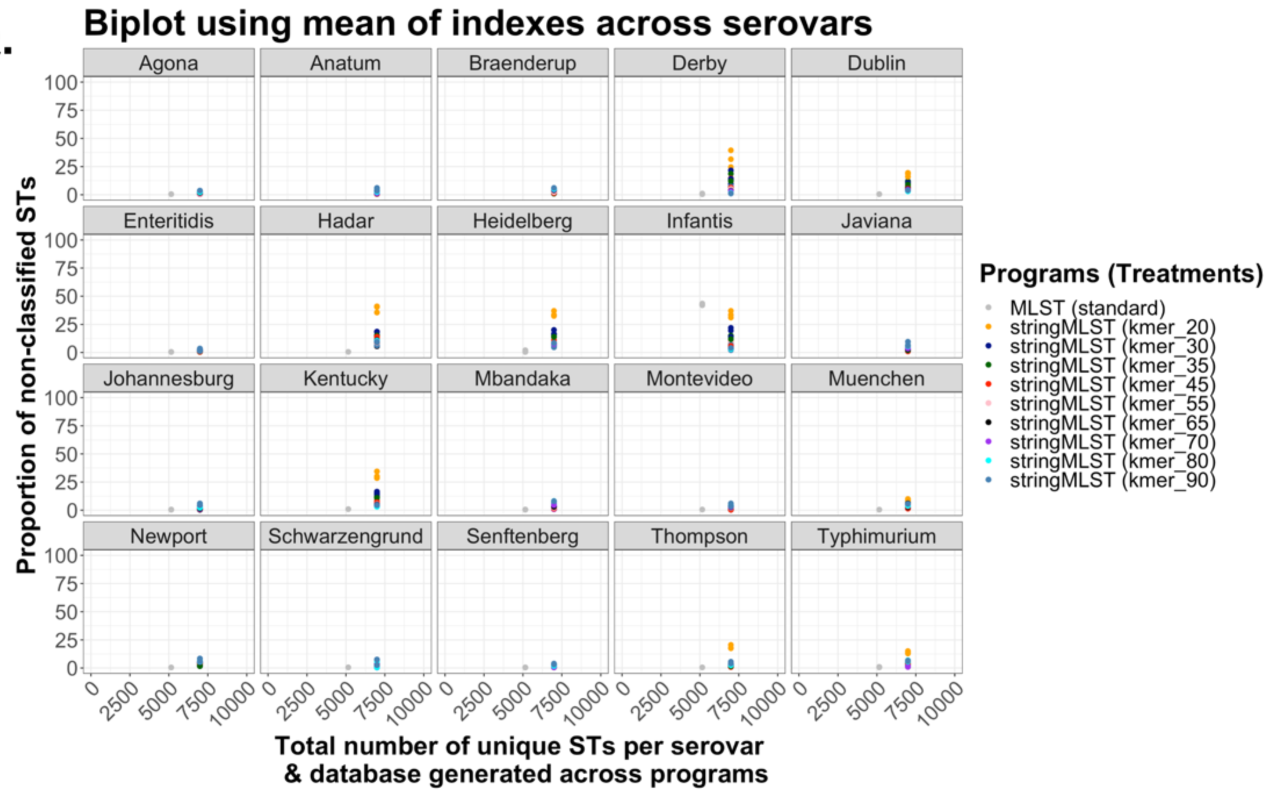
Biplot using mean of indexes across serovars
Agona
Anatum
Braenderup
Derby
Dublin
Enteritidis
Hadar
Heidelberg
Infantis
Javiana
Johannesburg
Kentucky
Mbandaka
Montevideo
Muenchen
Newport
Schwarzengrund
Senftenberg
Thompson
Typhimurium
Proportion of non-classified STs
Total number of unique STs per serovar & database generated across programs
Programs (Treatments)
MLST (standard)
stringMLST (kmer_20)
stringMLST (kmer_30)
stringMLST (kmer_35)
stringMLST (kmer_45)
stringMLST (kmer_55)
stringMLST (kmer_65)
stringMLST (kmer_70)
stringMLST (kmer_80)
stringMLST (kmer_90)

R.

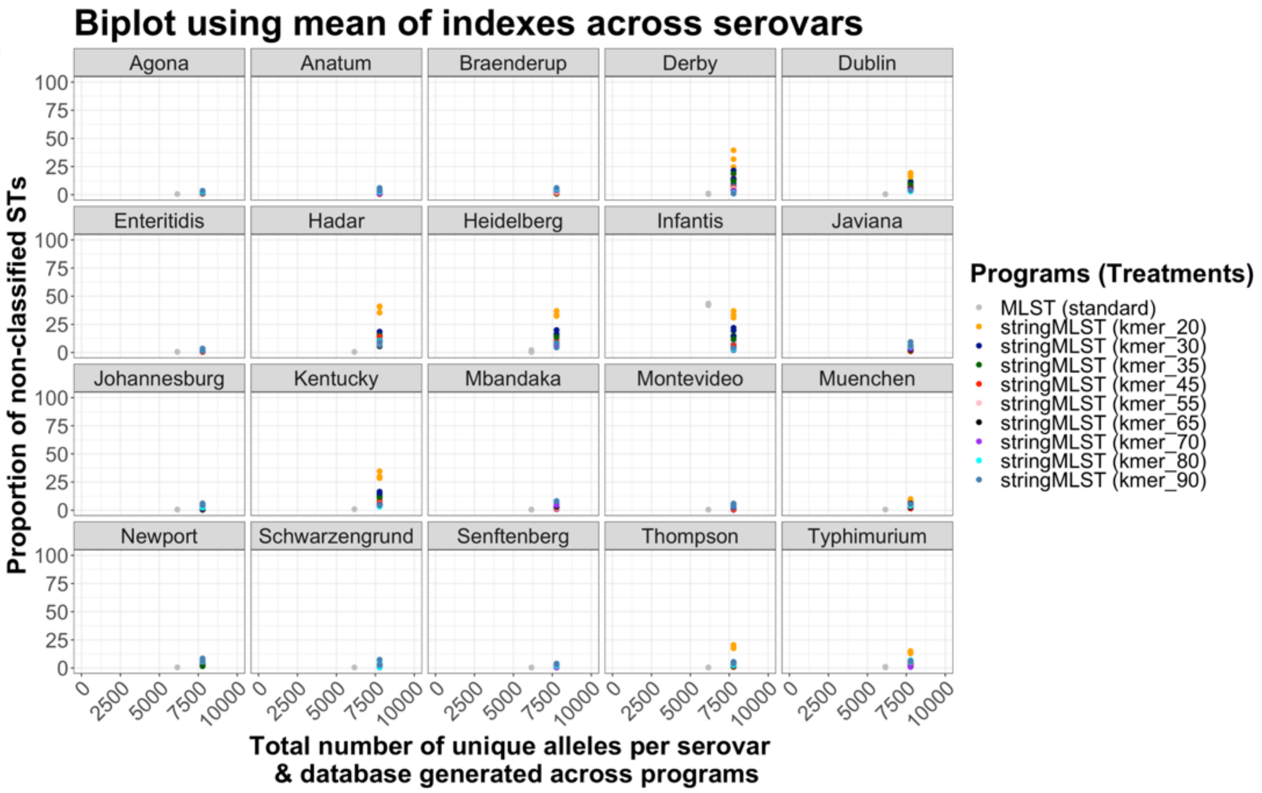

Figure S13. Bivariate association between accuracy-based classification metrics and genome-intrinsic and –extrinsic variables across *S. enterica* serovars.

(A) Scatter plot demonstrating the relationship between the Simpson's D index of diversity (1 – *D*) vs. ST richness (Richness). (B) Scatter plot demonstrating the relationship between the Simpson's D index of diversity (1 – *D*) vs. proportion of non-classified STs. (C) Scatter plot demonstrating the relationship between the ST richness (Richness) vs. proportion of non-classified STs across *S. enterica* serovars and programs. (D) Scatter plot demonstrating the relationship between the ST richness (Richness) vs. median number of contigs per genome across *S. enterica* serovars and programs. (E) Scatter plot demonstrating the relationship between the ST richness (Richness) vs. the average total number of nucleotides per genome across *S. enterica* serovars and programs. (F)

Scatter plot demonstrating the relationship between the ST richness (Richness) vs. the average GC% content per genome. (G) Scatter plot demonstrating the relationship between the ST richness (Richness) vs. the total number of unique STs in the database. (H) Scatter plot demonstrating the relationship between the ST richness (Richness) vs. the total number of unique alleles across all 7 loci in the database. (I) Scatter plot demonstrating the relationship between the Simpson's D index of diversity (1 – *D*) vs. the median number of contigs per genome. (J) Scatter plot demonstrating the relationship between the Simpson's D index of diversity (1 – *D)* vs. the average total number of nucleotides per genome. (K) Scatter plot demonstrating the relationship between the Simpson's D index of diversity (1 – *D)* vs. the average GC% content per genome. (L) Scatter plot demonstrating the relationship between the Simpson's D index of diversity (1 – *D)* vs. the total number of unique STs in the database. (M) Scatter plot demonstrating the relationship between the Simpson's D index of diversity (1 – *D)* vs. the total number of unique alleles across all 7 loci in the database. (N) Scatter plot demonstrating the relationship between the proportion of non-classified STs vs. the median number of contigs per genome. (O) Scatter plot demonstrating the relationship between the proportion of non-classified STs vs. the average total number of nucleotides per genome. (P) Scatter plot demonstrating the relationship between the proportion of non-classified STs vs. the average GC% content per genome. (Q) Scatter plot demonstrating the relationship between the proportion of non-classified STs vs. the total number of unique STs in the database. (R) Scatter plot demonstrating the relationship between the proportion of non-classified STs vs. the total number of unique alleles across all 7 loci in the database. All plots were stratified by *S. enterica* serovars and data points were colored by the program (mlst vs. stringMLST including all kmer lengths).

A.

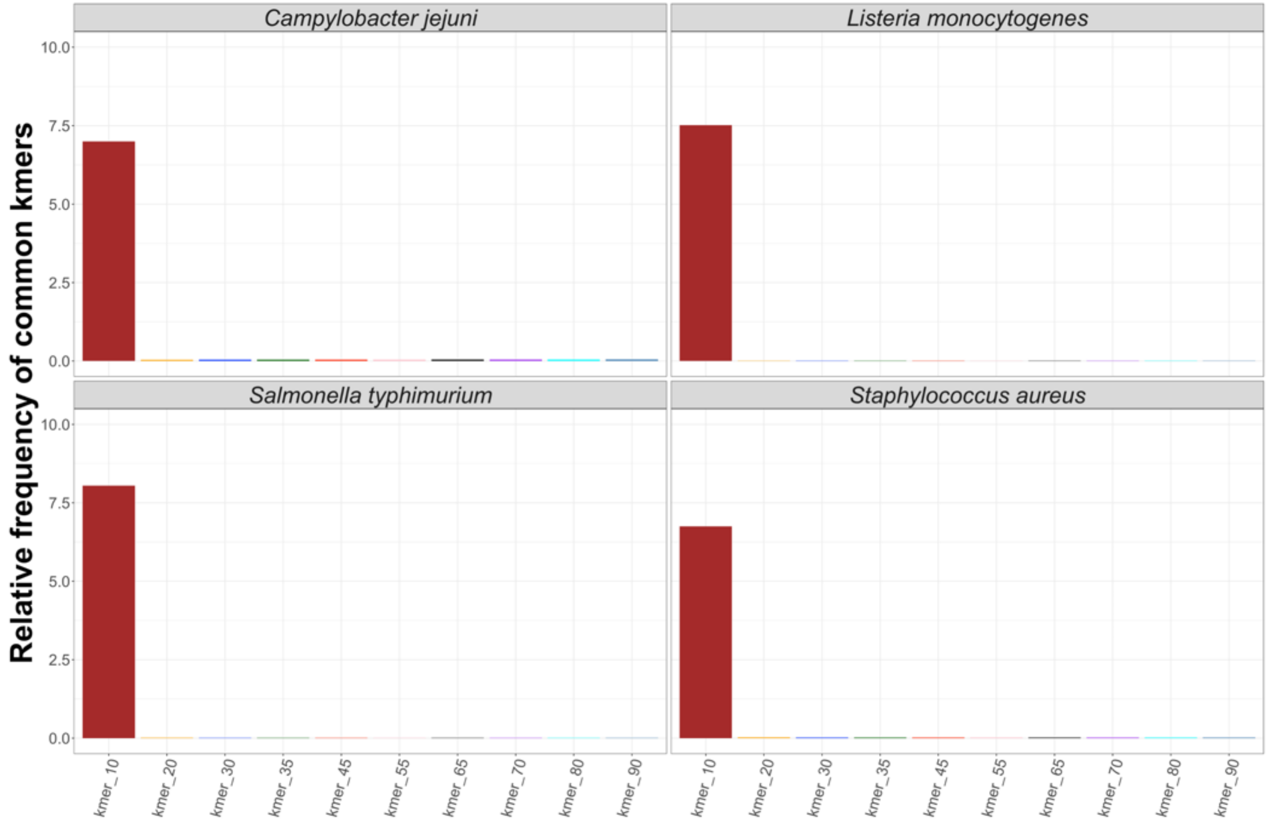

Campylobacter jejuni
Listeria monocytogenes
Salmonella typhimurium
Staphylococcus aureus
Relative frequency of common kmers
10.0
7.5
5.0
2.5
0.0
kmer_10
kmer_20
kmer_30
kmer_35
kmer_45
kmer_55
kmer_65
kmer_70
kmer_80
kmer_90

**B.**

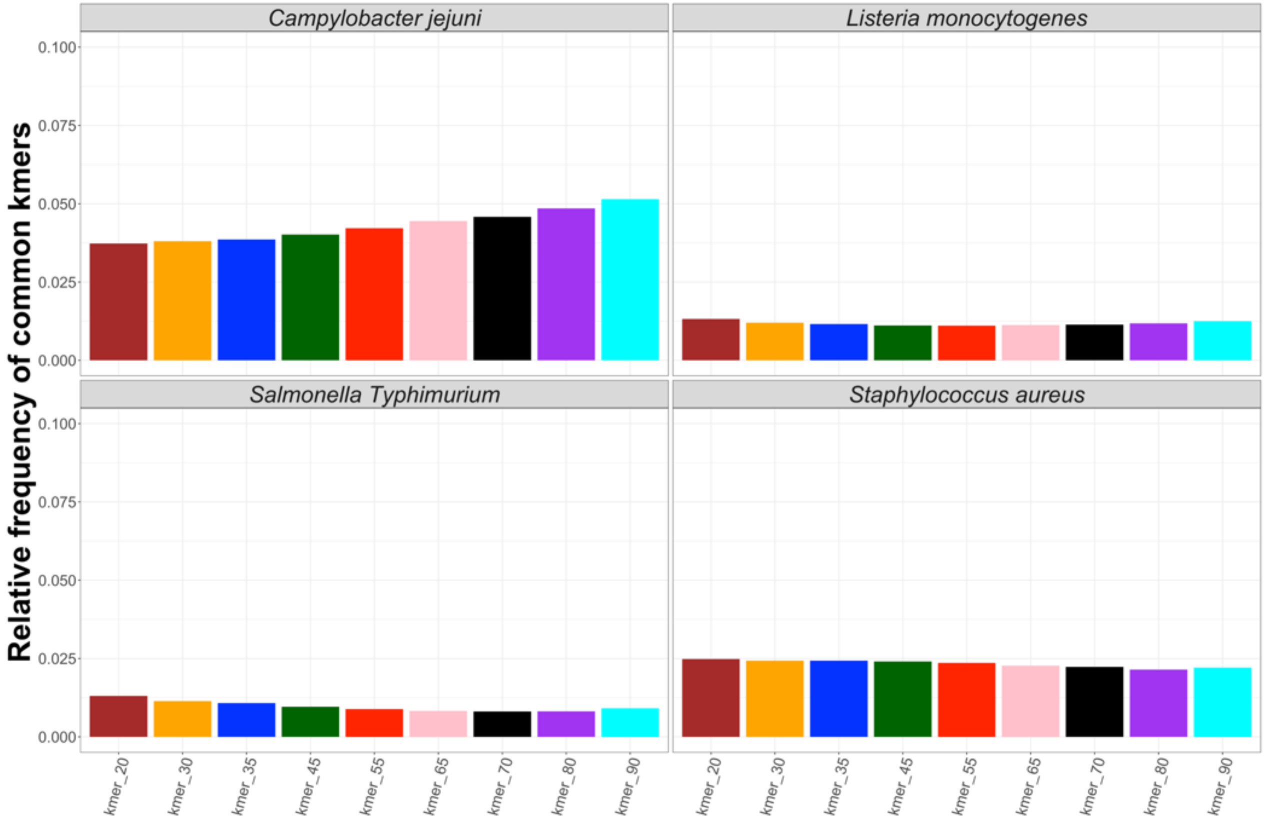

Figure S14. Relative frequency of common kmers found in the raw reads and the stringMLST database. Random 100 raw paired-end reads from the initial *C. jejuni*, *L. monocytogenes*, *S. aureus* and *S.* Typhimurium (major representative zoonotic serovar of *S. enterica*) datasets were selected, and DSK was used to count the occurrence of kmers of lengths 10, 20, 30, 35, 45, 55, 65, 70, 80 and 90 respectively in the raw reads. (A) Next, for each database created with stringMLST, a file with the kmer frequency for the used ST scheme was generated. Using the kmers generated from the raw reads and the stringMLST database, a relative frequency of the common kmers was calculated. (B) Due to the large range of frequency-based values between kmer 10 and the remaining kmer lengths, the kmer 10 data were excluded from the plot.

**A.**

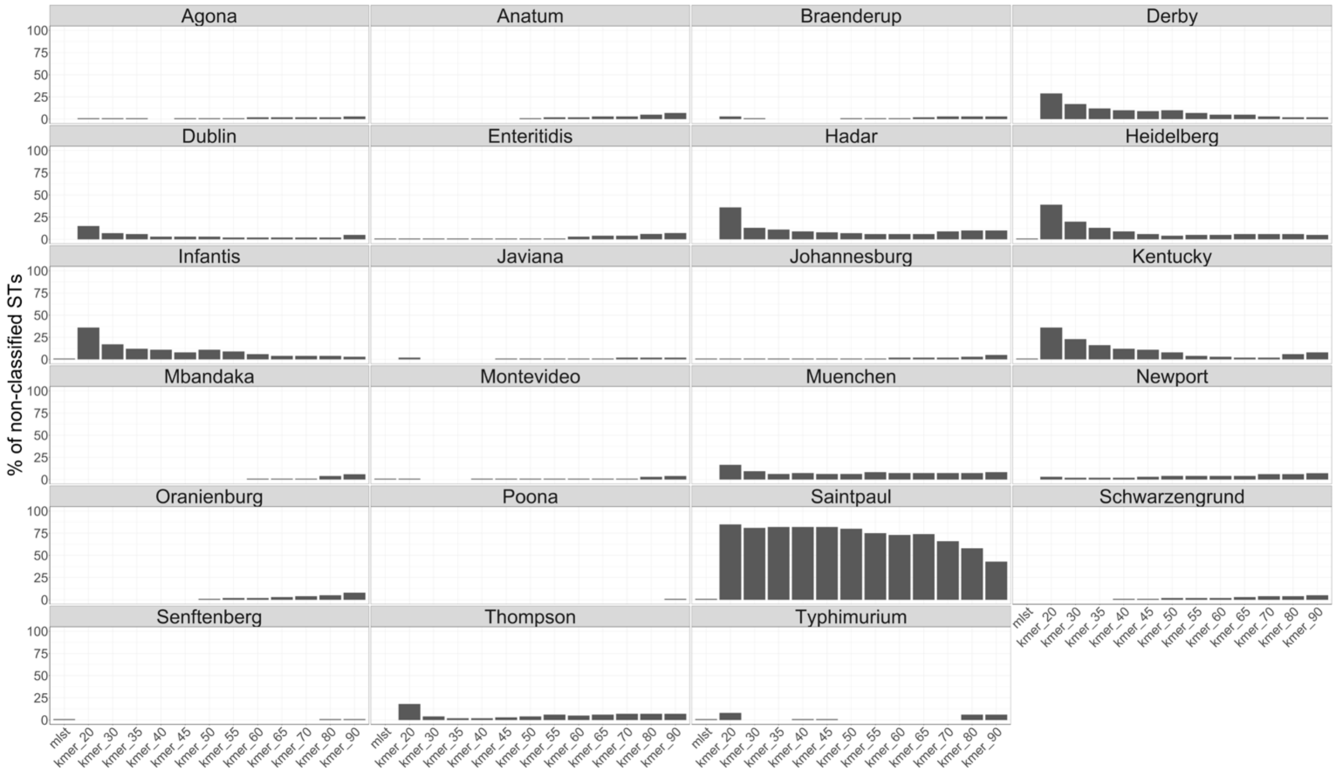

% of non-classified STs
Agona
Anatum
Braenderup
Derby
Dublin
Enteritidis
Hadar
Heidelberg
Infantis
Javiana
Johannesburg
Kentucky
Mbandaka
Montevideo
Muenchen
Newport
Oranienburg
Poona
Saintpaul
Schwarzengrund
Senftenberg
Thompson
Typhimurium
100
75
50
25
0
mlst
kmer_20
kmer_30
kmer_35
kmer_40
kmer_45
kmer_50
kmer_55
kmer_60
kmer_65
kmer_70
kmer_80
kmer_90

**B.**

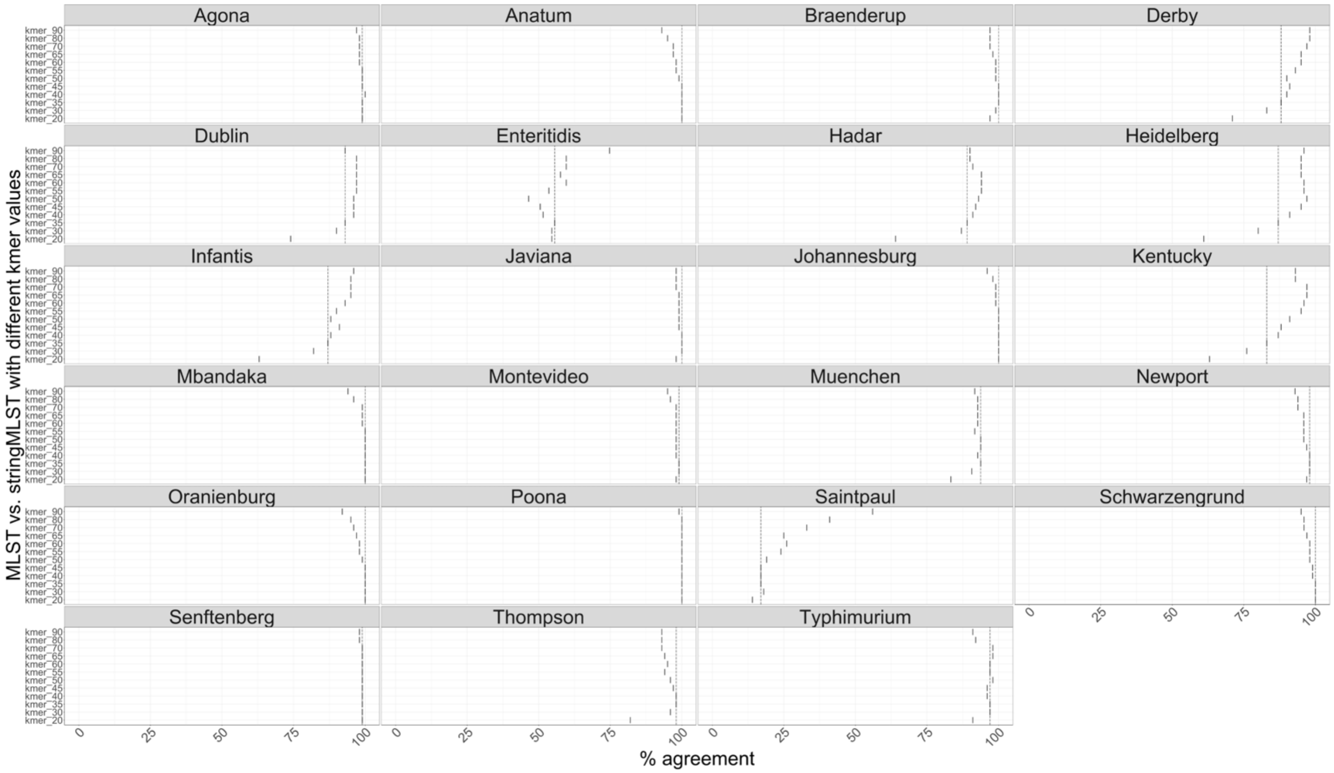

Agona
Anatum
Braenderup
Derby
Dublin
Enteritidis
Hadar
Heidelberg
Infantis
Javiana
Johannesburg
Kentucky
Mbandaka
Montevideo
Muenchen
Newport
Oranienburg
Poona
Saintpaul
Schwarzengrund
Senftenberg
Thompson
Typhimurium
kmer_90
kmer_80
kmer_70
kmer_65
kmer_60
kmer_55
kmer_50
kmer_45
kmer_40
kmer_35
kmer_30
kmer_20
0
25
50
75
100
% agreement
MLST vs. stringMLST with different kmer values

C.

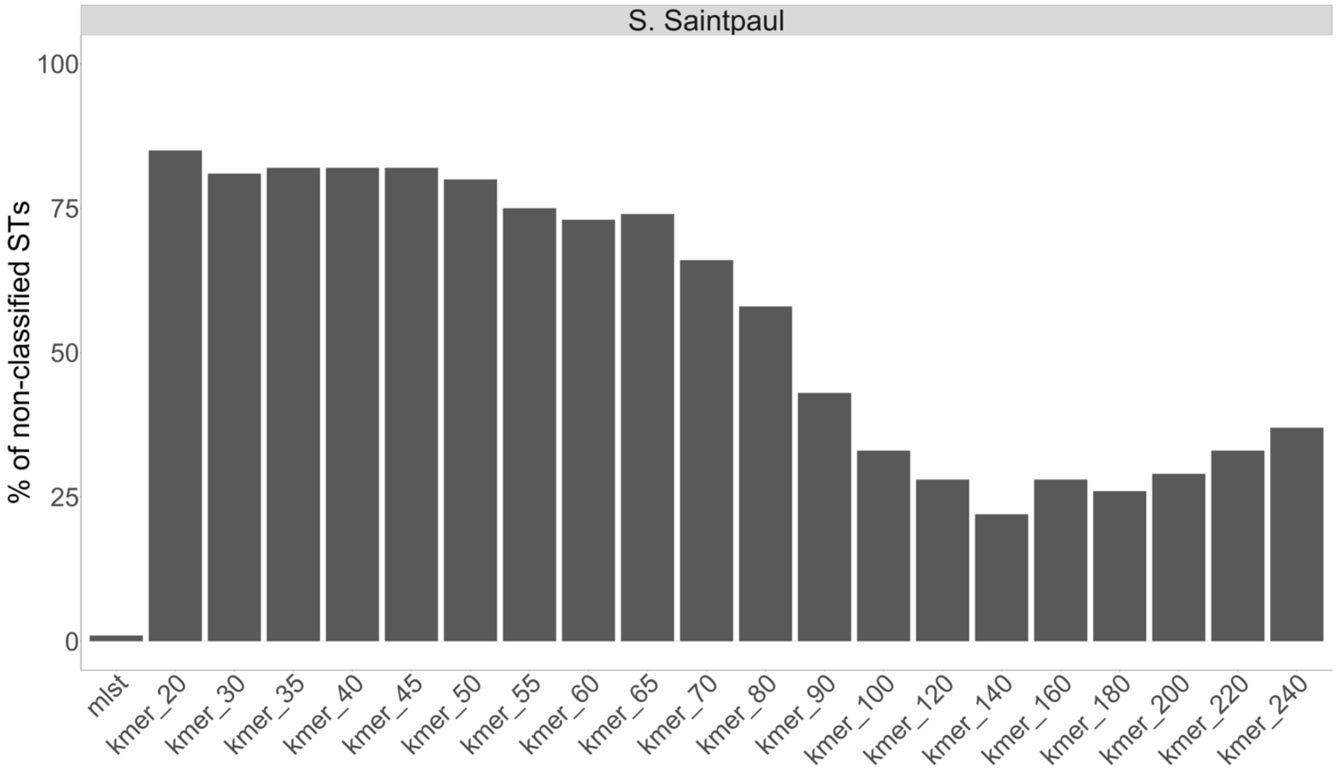
S. Saintpaul
% of non-classified STs
100
75
50
25
0
mlst
kmer_20
kmer_30
kmer_35
kmer_40
kmer_45
kmer_50
kmer_55
kmer_60
kmer_65
kmer_70
kmer_80
kmer_90
kmer_100
kmer_120
kmer_140
kmer_160
kmer_180
kmer_200
kmer_220
kmer_240

D.

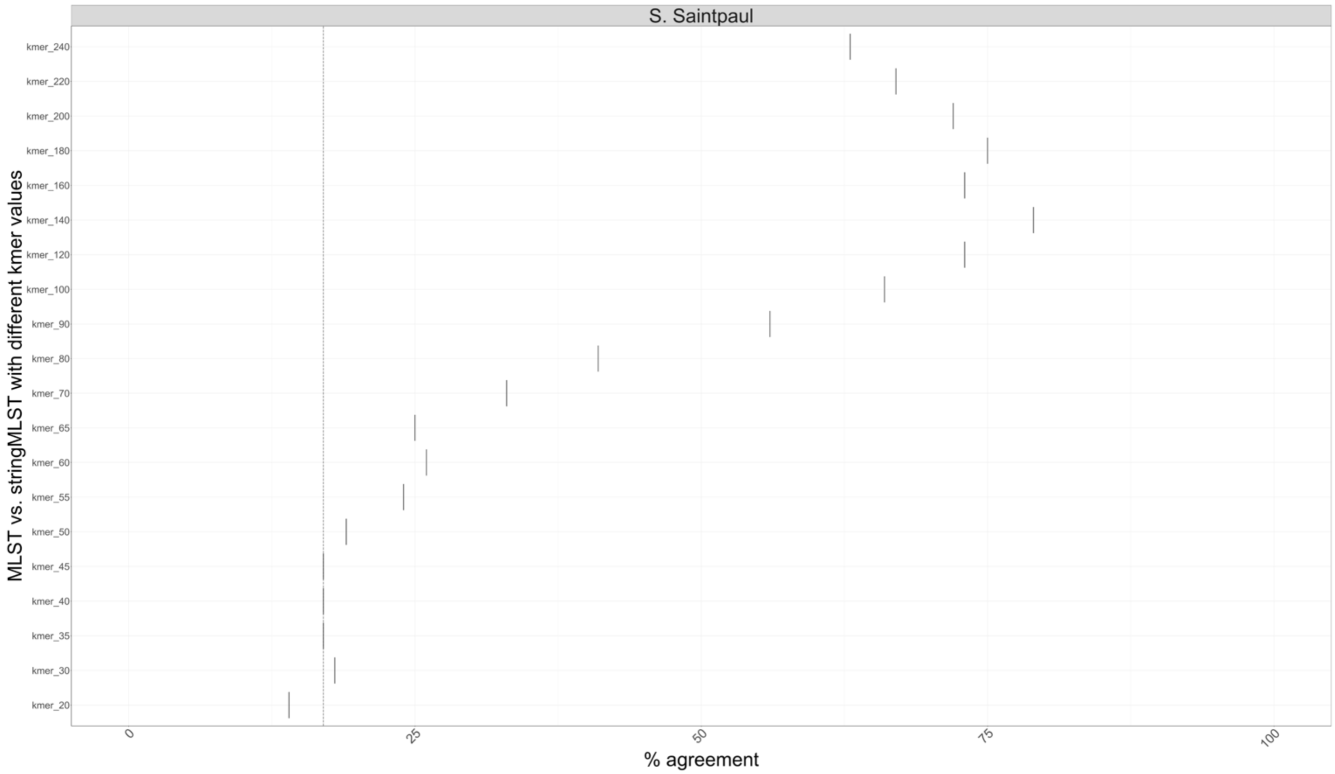
S. Saintpaul
MLST vs. stringMLST with different kmer values
kmer_240
kmer_220
kmer_200
kmer_180
kmer_160
kmer_140
kmer_120
kmer_100
kmer_90
kmer_80
kmer_70
kmer_65
kmer_60
kmer_55
kmer_50
kmer_45
kmer_40
kmer_35
kmer_30
kmer_20
0
25
50
75
100
% agreement

E.

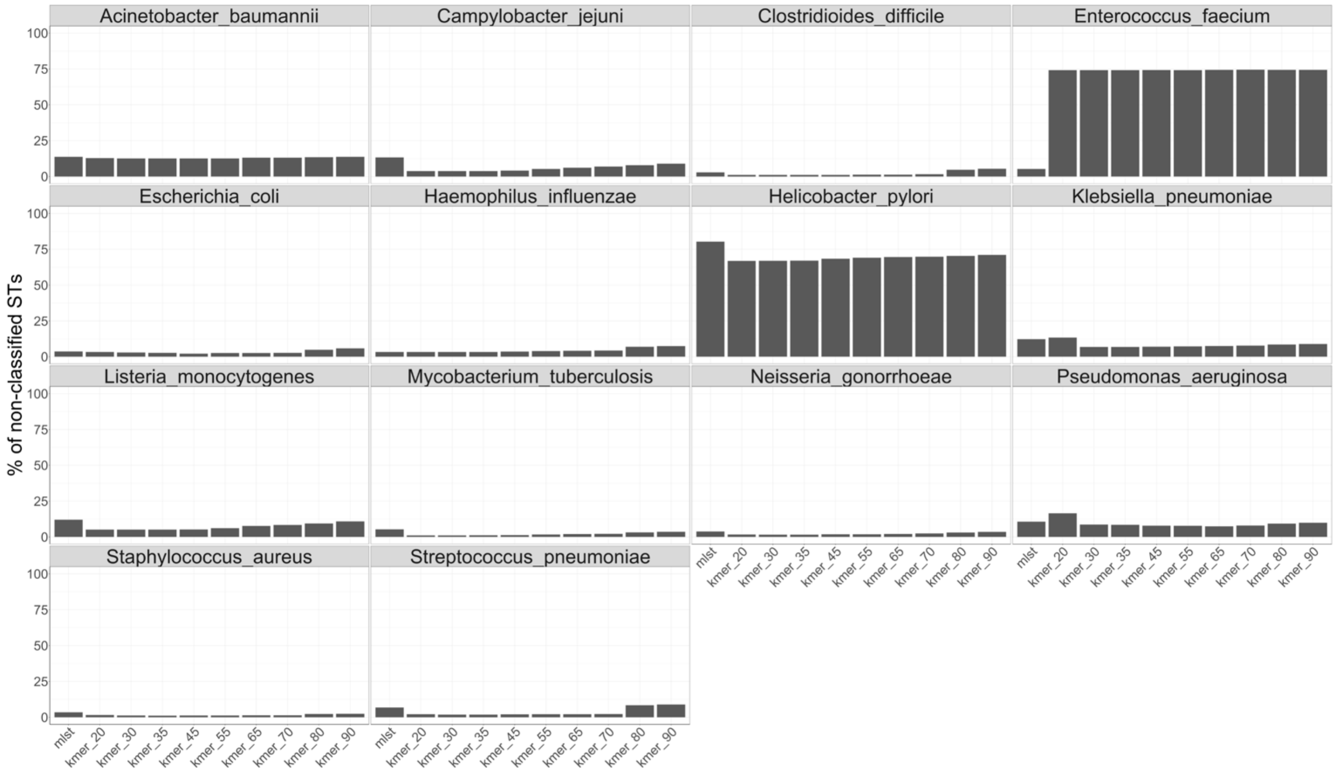
Acinetobacter_baumannii
Campylobacter_jejuni
Clostridioides_difficile
Enterococcus_faecium
Escherichia_coli
Haemophilus_influenzae
Helicobacter_pylori
Klebsiella_pneumoniae
Listeria_monocytogenes
Mycobacterium_tuberculosis
Neisseria_gonorrhoeae
Pseudomonas_aeruginosa
Staphylococcus_aureus
Streptococcus_pneumoniae
% of non-classified STs
0
25
50
75
100
mlst
kmer_20
kmer_30
kmer_35
kmer_45
kmer_55
kmer_65
kmer_70
kmer_80
kmer_90

F.

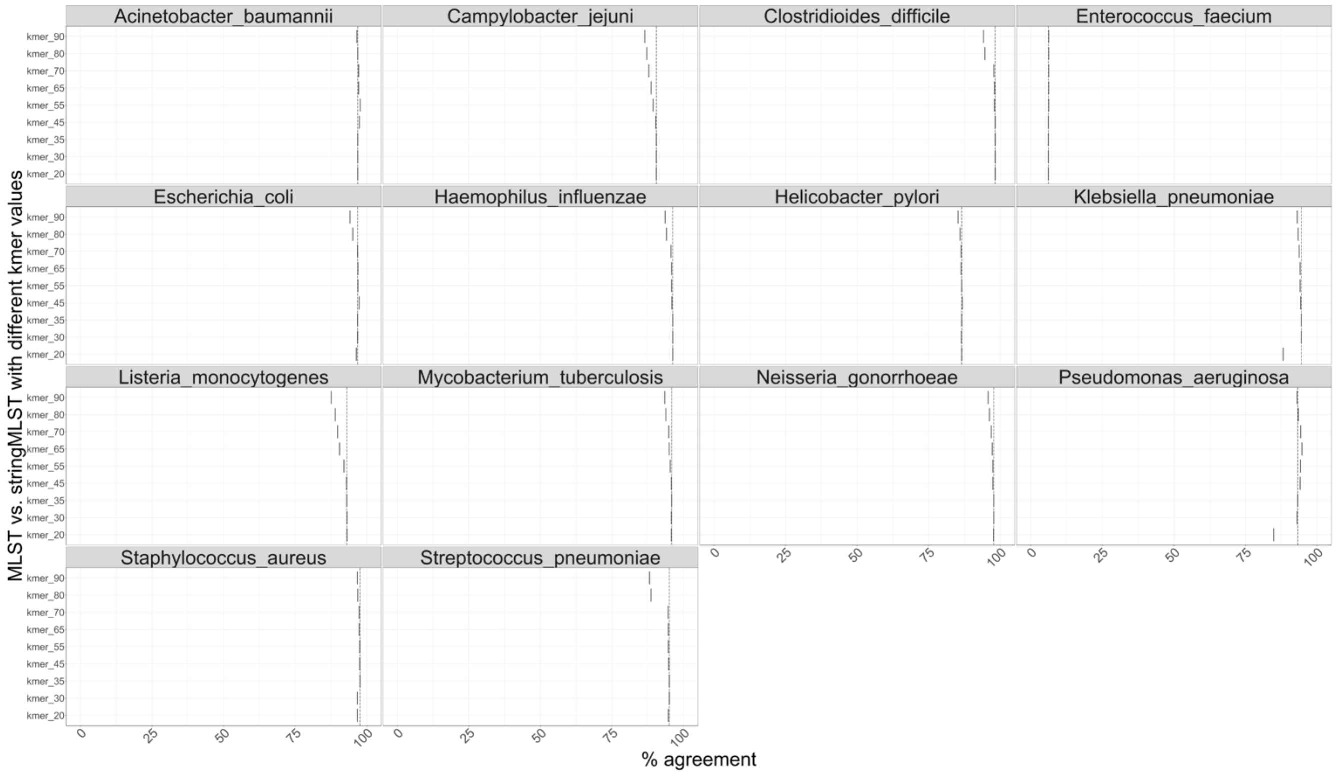
Acinetobacter_baumannii
Campylobacter_jejuni
Clostridioides_difficile
Enterococcus_faecium
Escherichia_coli
Haemophilus_influenzae
Helicobacter_pylori
Klebsiella_pneumoniae
Listeria_monocytogenes
Mycobacterium_tuberculosis
Neisseria_gonorrhoeae
Pseudomonas_aeruginosa
Staphylococcus_aureus
Streptococcus_pneumoniae
kmer_90
kmer_80
kmer_70
kmer_65
kmer_55
kmer_45
kmer_35
kmer_30
kmer_20
0
25
50
75
100
MLST vs. stringMLST with different kmer values
% agreement

G.

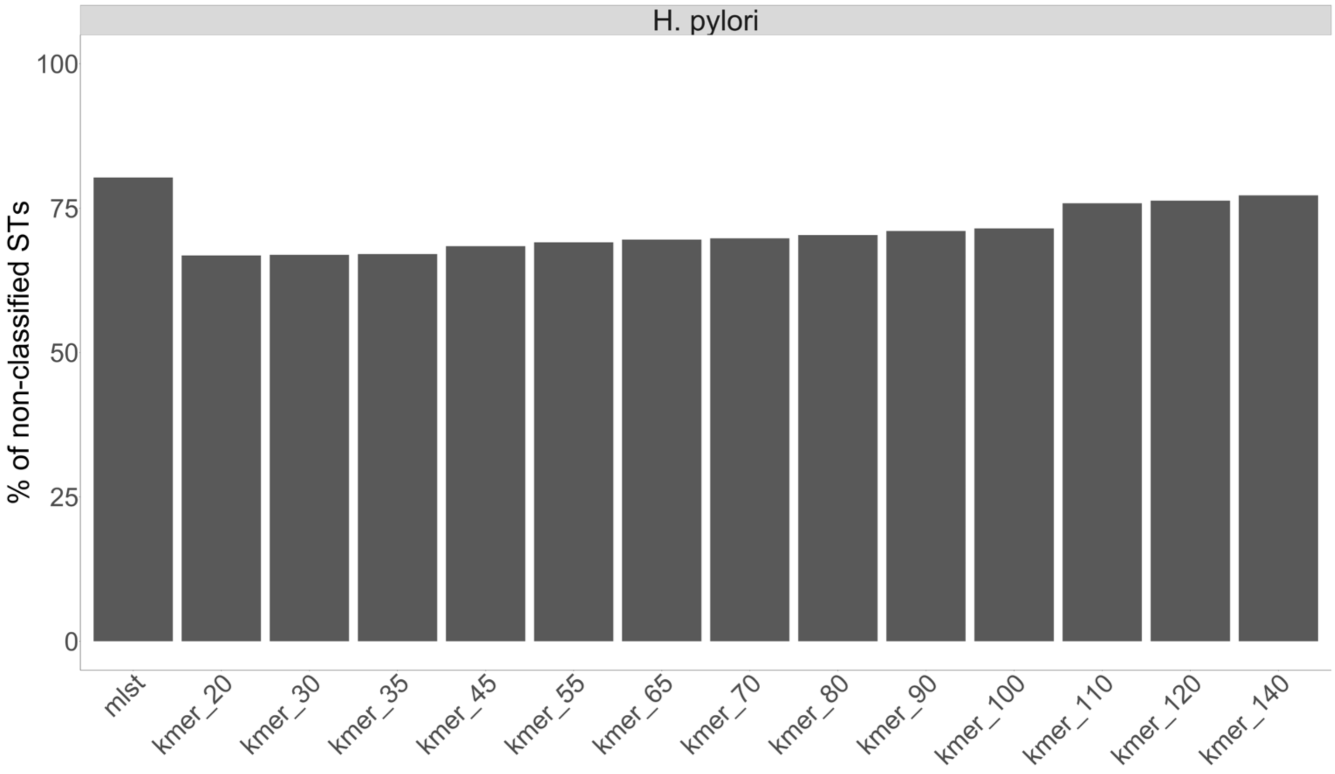

H. pylori
% of non-classified STs
100
75
50
25
0
mlst
kmer_20
kmer_30
kmer_35
kmer_45
kmer_55
kmer_65
kmer_70
kmer_80
kmer_90
kmer_100
kmer_110
kmer_120
kmer_140

H.

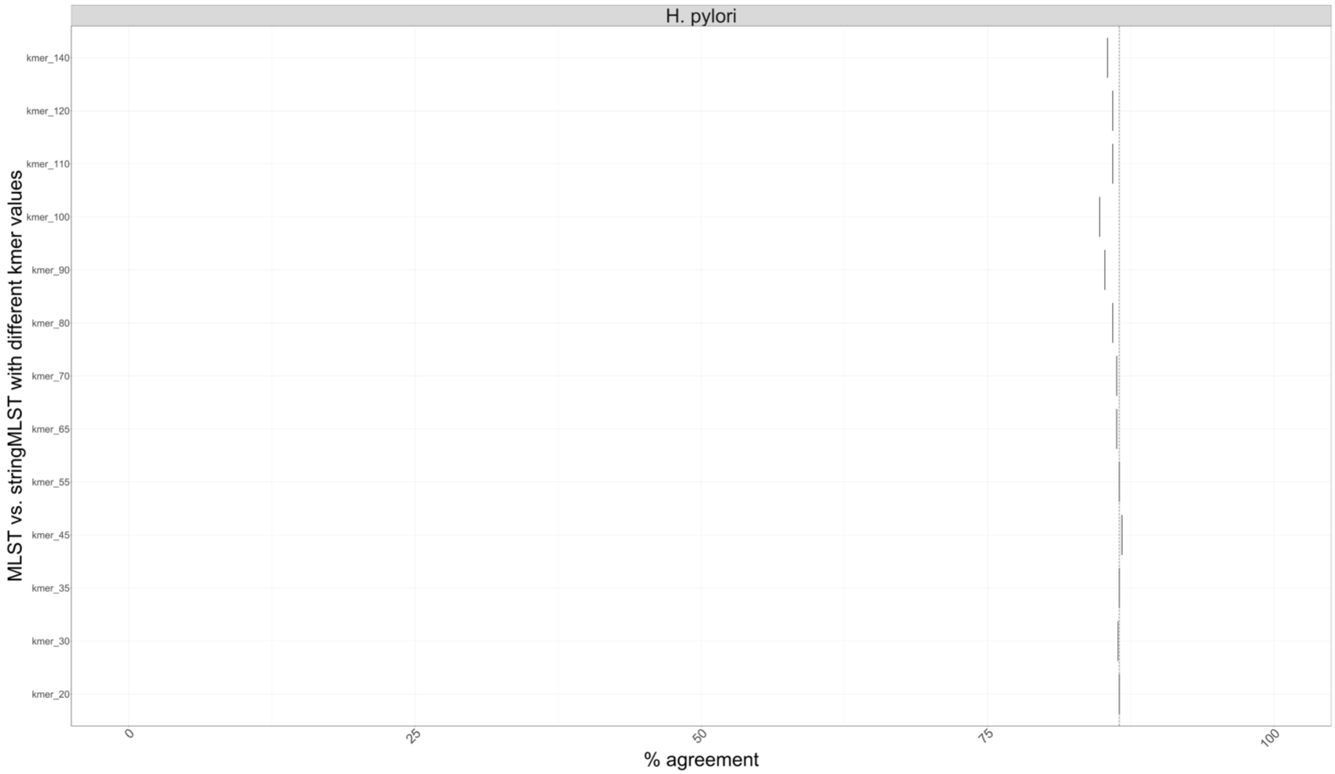

H. pylori
MLST vs. stringMLST with different kmer values
kmer_140
kmer_120
kmer_110
kmer_100
kmer_90
kmer_80
kmer_70
kmer_65
kmer_55
kmer_45
kmer_35
kmer_30
kmer_20
0
25
50
75
100
% agreement

I.

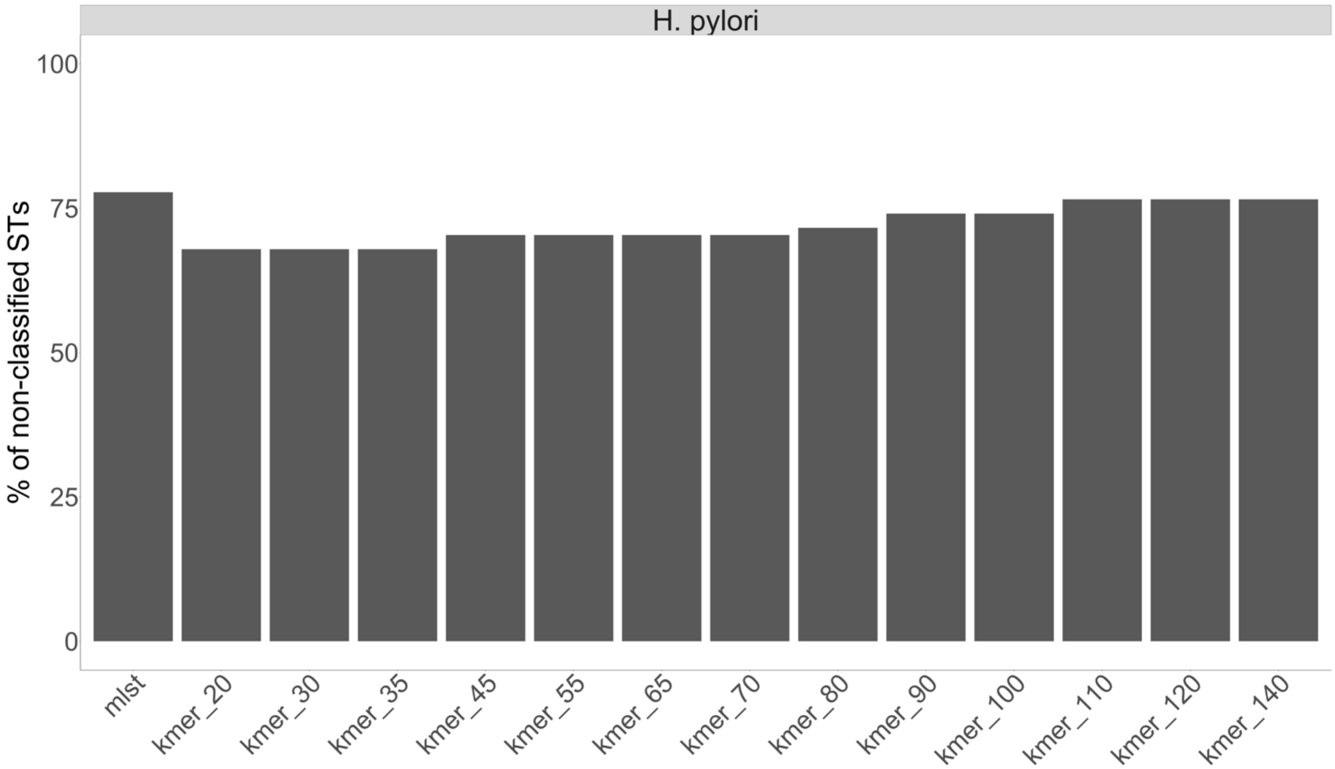

H. pylori
% of non-classified STs
100
75
50
25
0
mlst
kmer_20
kmer_30
kmer_35
kmer_45
kmer_55
kmer_65
kmer_70
kmer_80
kmer_90
kmer_100
kmer_110
kmer_120
kmer_140

J.

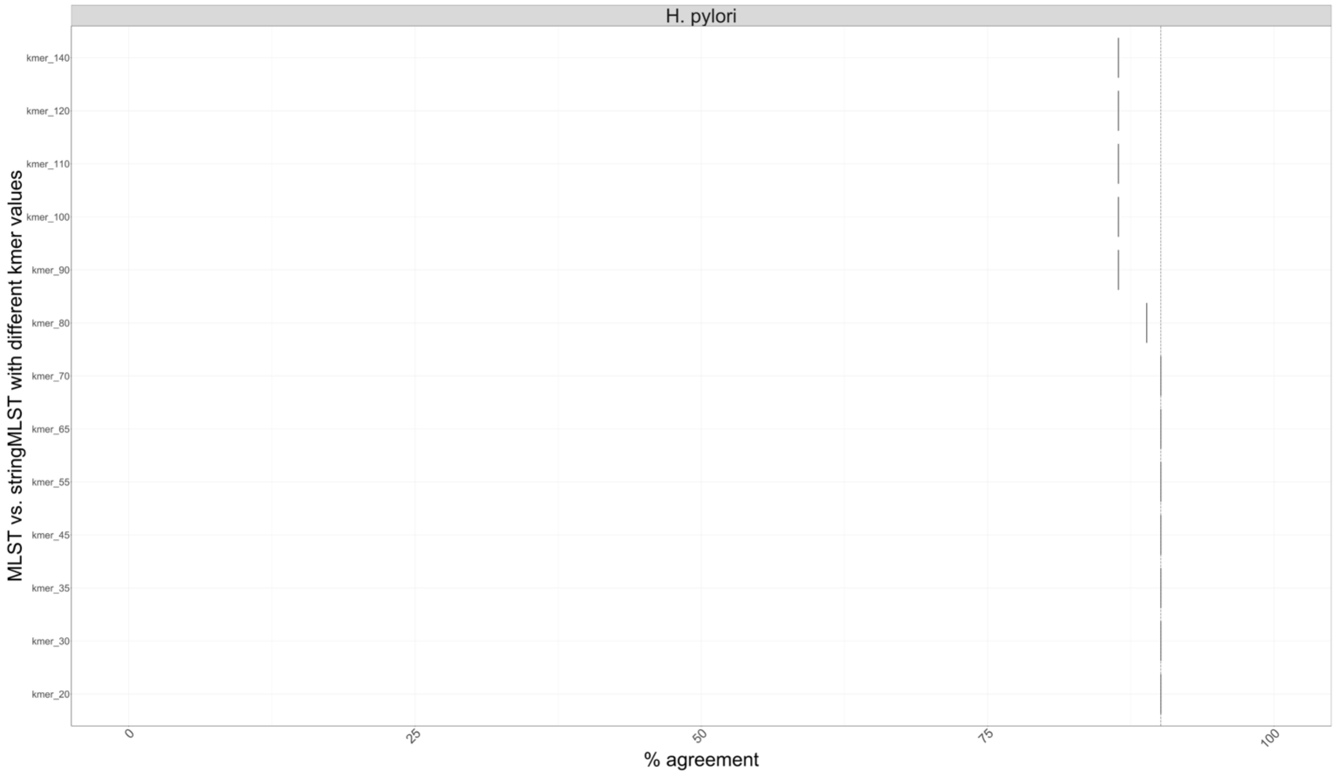
H. pylori
kmer_140
kmer_120
kmer_110
kmer_100
kmer_90
kmer_80
kmer_70
kmer_65
kmer_55
kmer_45
kmer_35
kmer_30
kmer_20
MLST vs. stringMLST with different kmer values
0
25
50
75
100
% agreement

K.

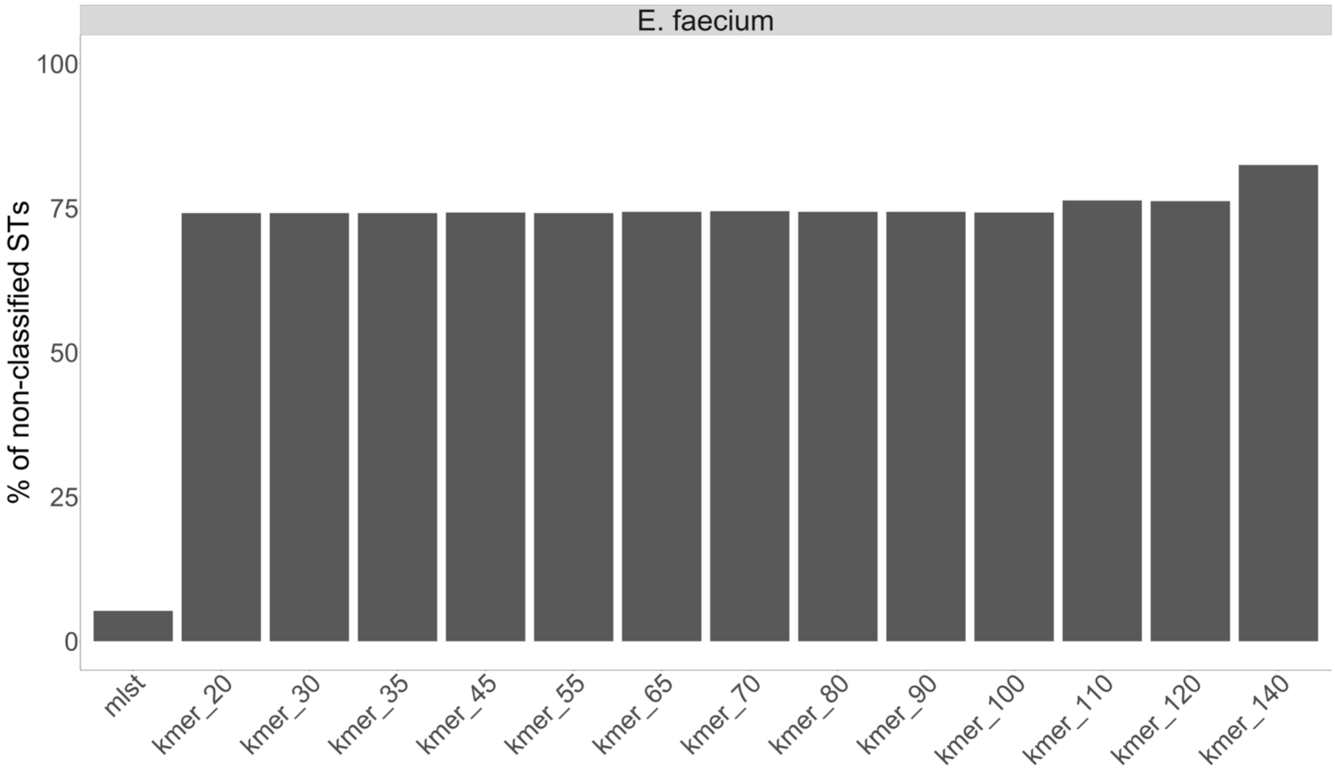
E. faecium
% of non-classified STs
100
75
50
25
0
mlst
kmer_20
kmer_30
kmer_35
kmer_45
kmer_55
kmer_65
kmer_70
kmer_80
kmer_90
kmer_100
kmer_110
kmer_120
kmer_140

L.

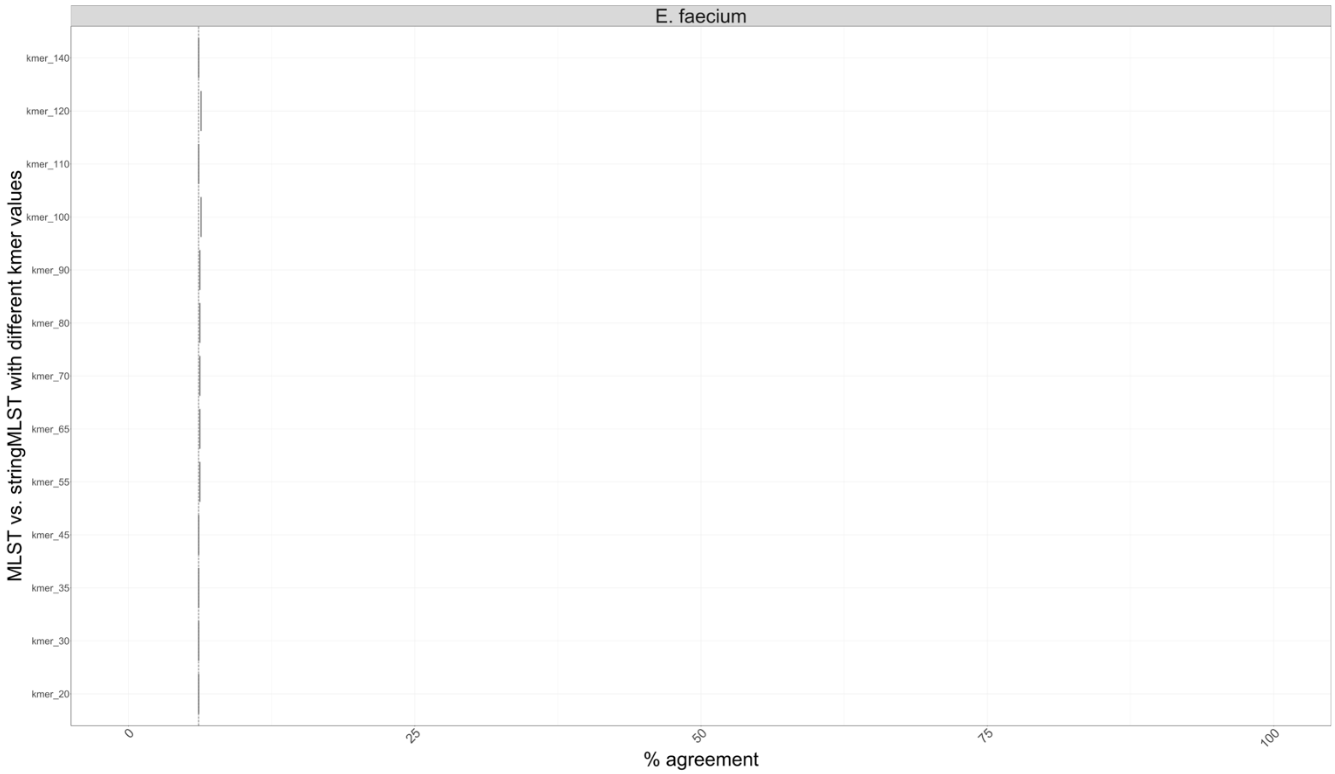

E. faecium
MLST vs. stringMLST with different kmer values
kmer_140
kmer_120
kmer_110
kmer_100
kmer_90
kmer_80
kmer_70
kmer_65
kmer_55
kmer_45
kmer_35
kmer_30
kmer_20
0
25
50
75
100
% agreement

M.

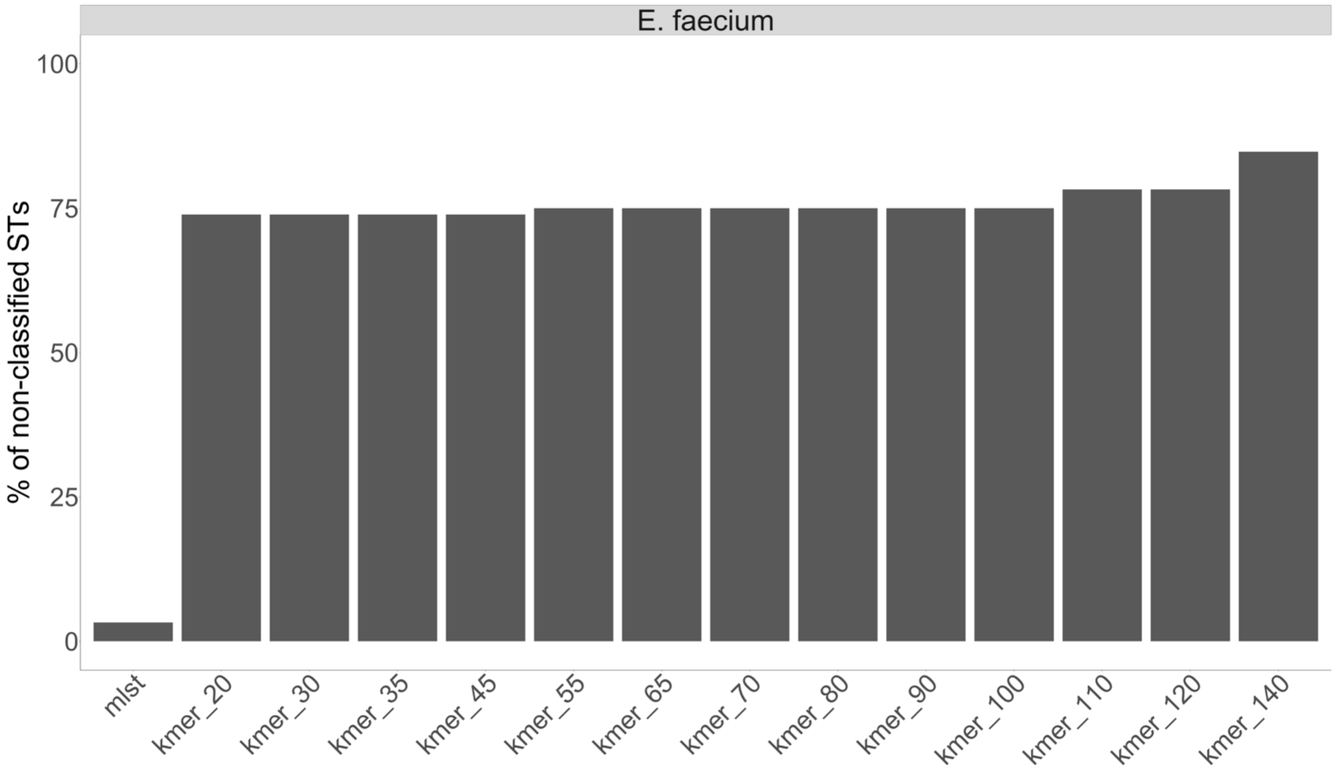
E. faecium
% of non-classified STs
100
75
50
25
0
mlst
kmer_20
kmer_30
kmer_35
kmer_45
kmer_55
kmer_65
kmer_70
kmer_80
kmer_90
kmer_100
kmer_110
kmer_120
kmer_140

N.

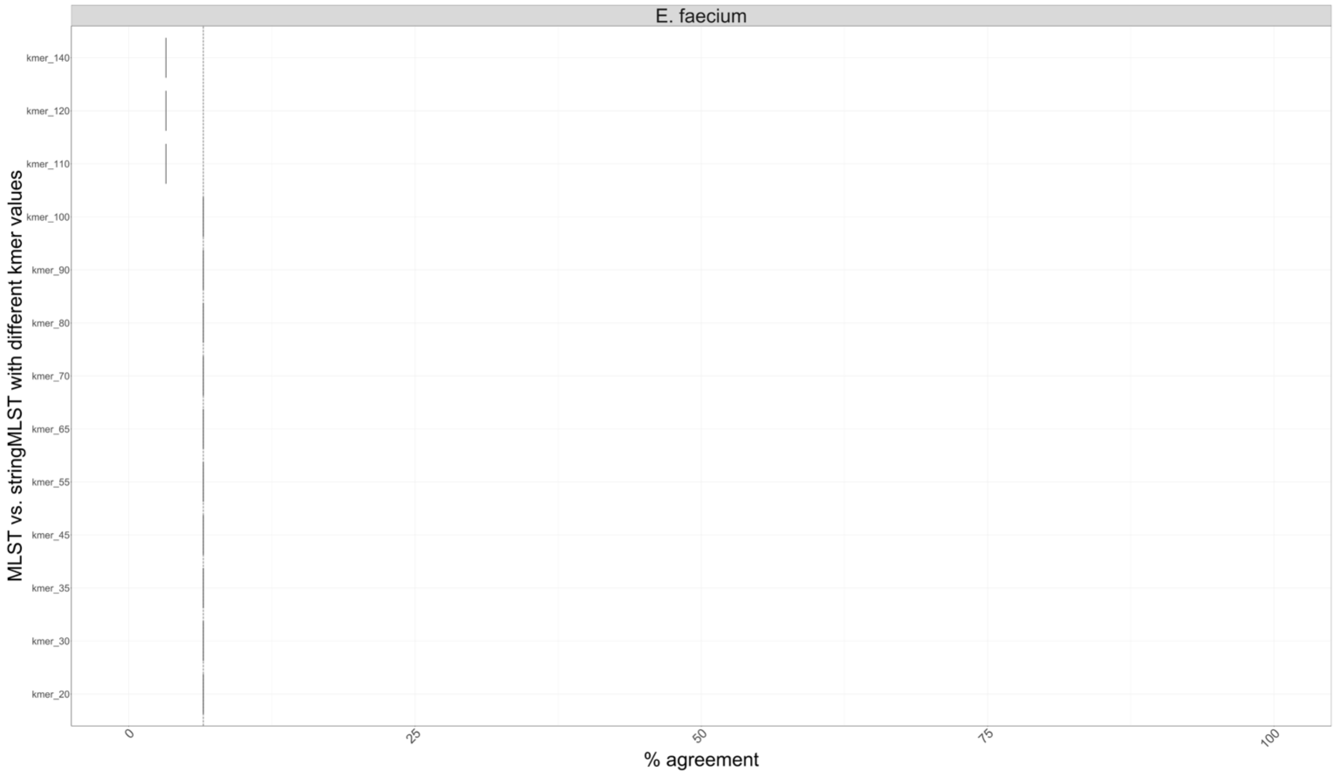

E. faecium
MLST vs. stringMLST with different kmer values
kmer_140
kmer_120
kmer_110
kmer_100
kmer_90
kmer_80
kmer_70
kmer_65
kmer_55
kmer_45
kmer_35
kmer_30
kmer_20
0
25
50
75
100
% agreement

O.

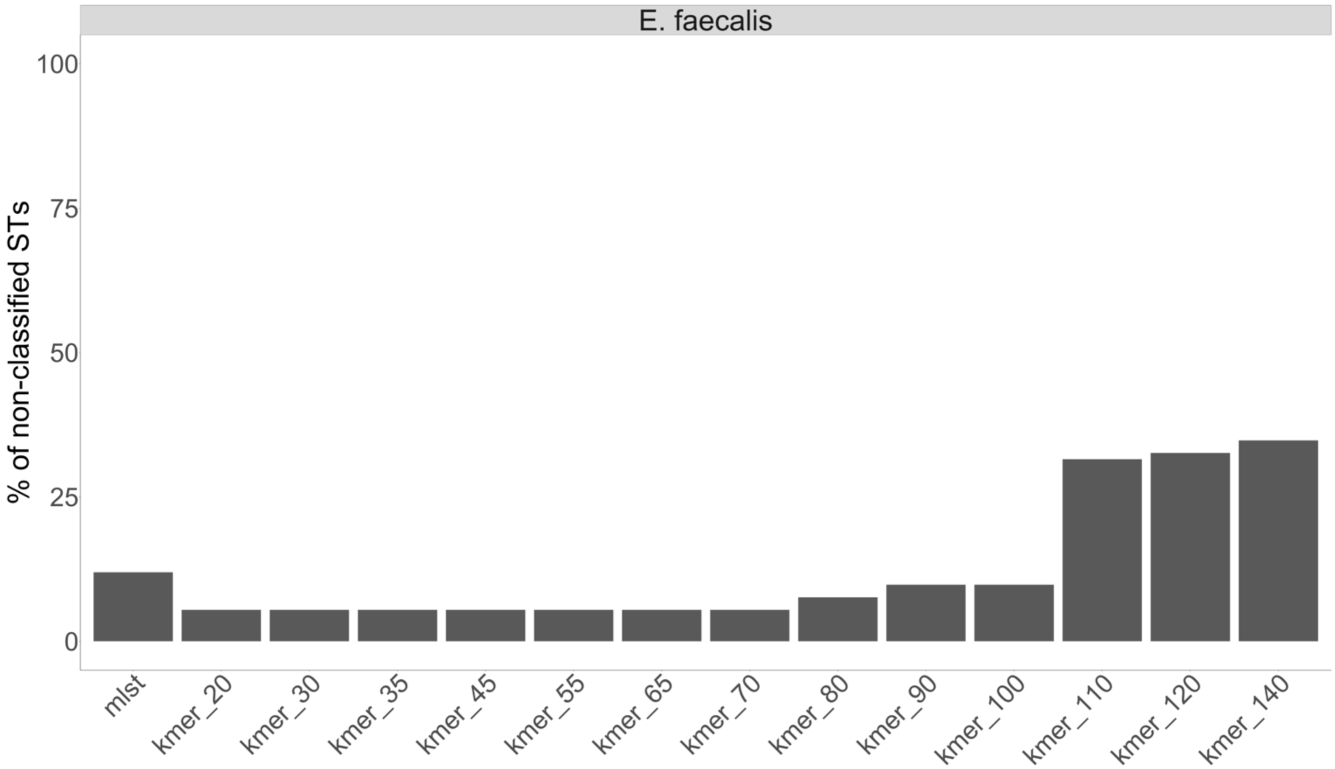
E. faecalis
% of non-classified STs
100
75
50
25
0
mlst
kmer_20
kmer_30
kmer_35
kmer_45
kmer_55
kmer_65
kmer_70
kmer_80
kmer_90
kmer_100
kmer_110
kmer_120
kmer_140

P.

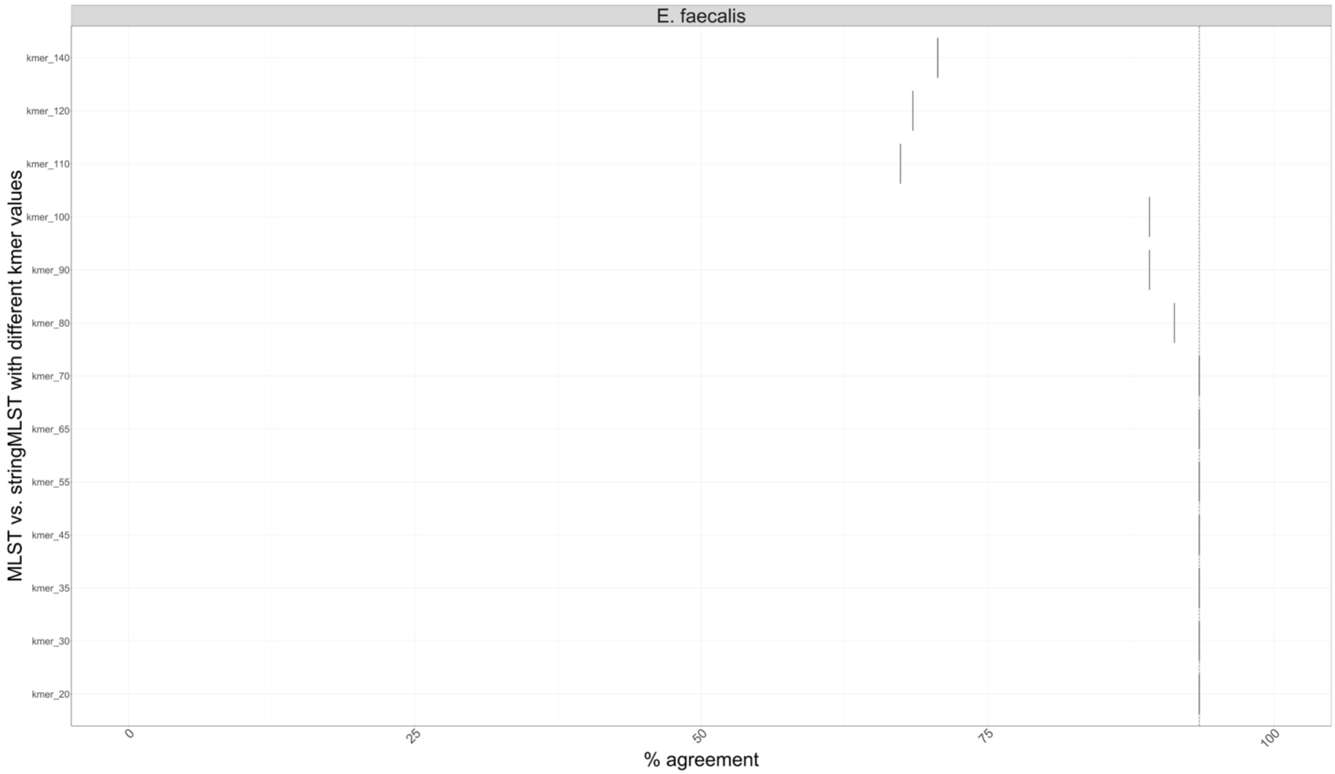

Figure S15. Comparison of the statistical performance metrics between mlst and stringMLST with different kmer lengths across *Salmonella enterica* serovars and other phylogenetic divergent bacterial pathogens.
For the twenty-three *Salmonella enterica* serovars (*S*. Agona, *S*. Anatum, *S*. Braenderup, *S*. Derby, *S*. Dublin, *S*. Enteritidis, *S*. Hadar, *S*. Heidelberg, *S*. Infantis, *S*. Javiana, *S*. Johannesburg, *S*. Kentucky, *S*. Mbandaka, *S*. Montevideo, *S*. Muenchen, *S*. Newport, *S*. Oranienburg, *S*. Poona, *S*. Saintpaul, *S*. Schwarzengrund, *S*. Senftenberg, *S*. Thompson, and *S*. Typhimurium), we randomly chose and downloaded 100 paired-end reads from NCBI-SRA. Next, for each dataset we ran mlst and stringMLST with kmer lengths ranging from 20, 30, 35, 40, 45, 50, 55, 60, 65, 70, 80, 90. For the fourteen bacterial pathogens (*Acinetobacter baumannii, Clostridioides difficile, Enterococcus faecium, Escherichia coli, Haemophilus influenzae, Helicobacter pylori, Klebsiella pneumoniae, Mycobacterium tuberculosis, Neisseria gonorrhoeae, Pseudomonas aeruginosa, Streptococcus pneumoniae,*

*Campylobacter jejuni, Listeria monocytogenes,* and *Staphylococcus aureus),* we randomly chose and downloaded 1,000 paired-end reads from NCBI-SRA. Next, for each dataset we ran mlst and stringMLST with kmer lengths ranging from 20, 30, 35, 45, 55, 65, 70, 80, 90. (A) Percentage of non-classified STs (ST calls that returned missing/blank values) across tweenty-three *S. enterica* serovars for mlst and stringMLST using an array of kmer lengths (20, 30, 35, 40, 45, 50, 55, 60, 65, 70, 80, 90); (B) Percentage of agreement (concordance) between programs ("good" or "bad" ST calls that matched between mlst and stringMLST for different kmer lengths) across all twenty-three S. *enterica* serovars; (C) Percentage of non-classified STs for *S*. Saintpaul using mlst and longer kmer lenghts with stringMLST; (D) Percentage of agreement  between programs for *S*. Saintpaul using mlst and longer kmer lengths with stringMLST; (E) Percentage of non-classified STs (ST calls that returned missing/blank values) across fourteen phylogenetic divergent bacterial pathogens for mlst and stringMLST with range of kmer lengths (20, 30, 35, 45, 55, 65, 70, 80, 90); (F) Percentage of agreement  between programs ("good" or "bad" ST calls that matched between mlst and stringMLST for different kmer lengths) across all fourteen phylogenetic divergent bacterial pathogens; (G) Percentage of non-classified STs for *H. pylori* (dataset of 1,000 genomes) using mlst and longer kmer lengths with stringMLST; (H) Percentage of agreement  between programs for *H. pylori* (dataset of 1,000 genomes) using mlst and lost kmer lengths with stringMLST; (I) Percentage of non-classified STs for *H. pylori* (dataset of 100 genomes) using mlst and longer kmer lengths with stringMLST; (J) Percentage of agreement  between programs for *H. pylori* (dataset of 100 genomes) using mlst and lost kmer lengths with stringMLST; (K) Percentage of non-classified STs for *E. faecium* (dataset of 1,000 genomes) using mlst and longer kmer lengths with stringMLST; (L) Percentage of agreement (concordance) between programs for *E. faecium* (dataset of 1,000 genomes) using mlst and longer kmer lengths with stringMLST; (M) Percentage of non-classified STs for *E. faecium* (dataset of 100 genomes) using mlst and longer kmer lengths with stringMLST; (N) Percentage of agreement (concordance) between programs for *E. faecium* (dataset of 100 genomes) using mlst and longer kmer lengths with stringMLST; (O) Percentage of non-classified STs for *Enterococcus faecalis* (dataset of 100 genomes) using mlst and longer kmer lenghts with stringMLST; (P) Percentage of agreement (concordance) between programs for *E. faecalis* (dataset of 100 genomes) using mlst and longer kmer lengths with stringMLST.

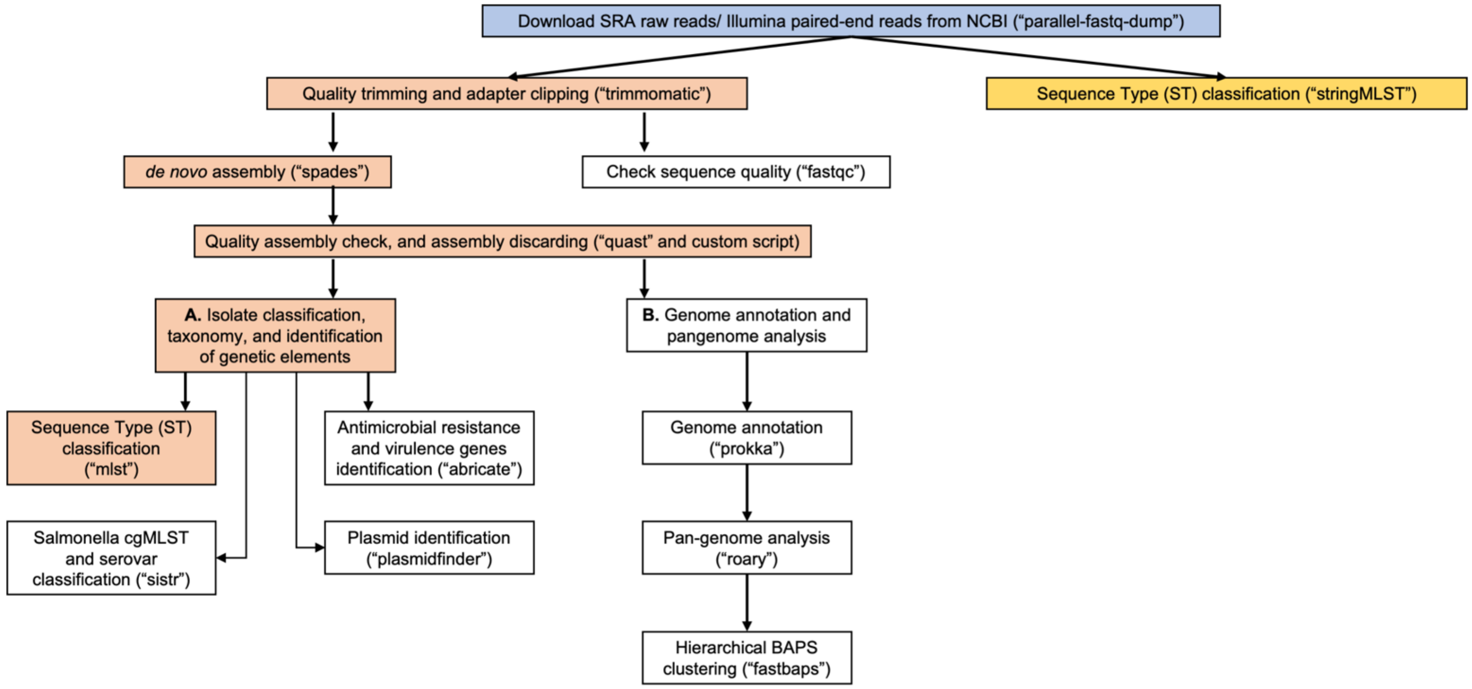

Figure S16. ProkEvo's computational workflow with both mlst and stringMLST included for ST-based classification.
Top-down flow of tasks for the ProkEvo computational pipeline. The squares represent the steps, while the bioinformatics tool used for each step is shown in brackets. The sub-workflow on the left includes all the steps of the current ProkEvo platform, for which the steps needed for obtaining ST classifications with mlst are colored in orange. The sub-workflow on the right includes all the steps of the alternative path in the ProkEvo platform, where stringMLST was integrated for ST-based classification.

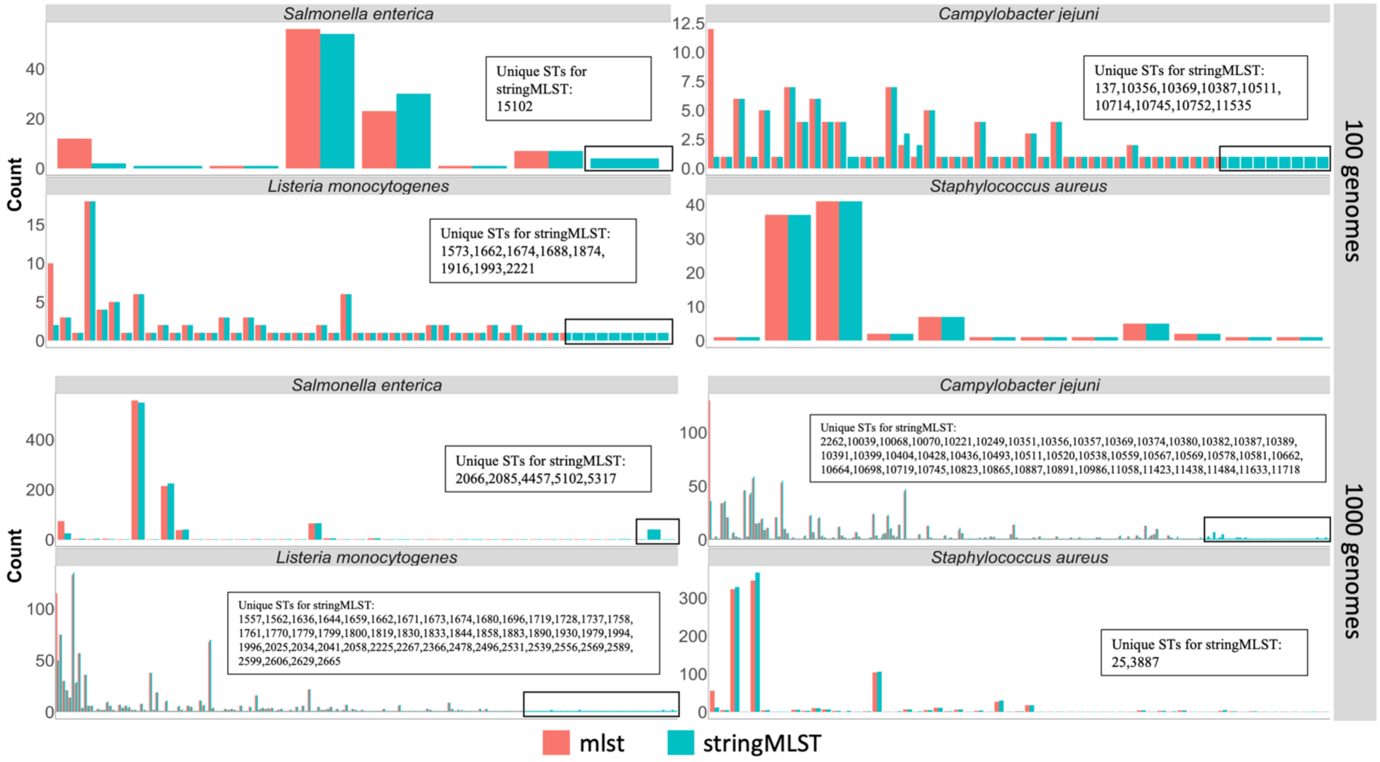

Figure S17. Frequency of ST lineages across bacterial species and programs using two random sample sizes. Pairwise frequency-based (y-axis = count) distribution of unique ST lineages identified by either mlst or stringMLST across bacterial species using a random sampling of 100 or 1,000 genomes. Each couple of pairwise bar counts (both programs) represents a unique ST number or lineage. Highlighted with black rectangles are the ST lineages uniquely found by the stringMLST program. Moreover, the STs unique for stringMLST are listed in a text box in each plot.

| **Serovar** | **Number of genomes downloaded from NCBI-SRA** |
|---|---|
| *Salmonella* Agona | 565 |
| *Salmonella* Anatum | 600 |
| *Salmonella* Braenderup | 600 |
| *Salmonella* Derby | 590 |
| *Salmonella* Dublin | 600 |
| *Salmonella* Enteritidis | 600 |
| *Salmonella* Hadar | 600 |
| *Salmonella* Heidelberg | 600 |
| *Salmonella* Infantis | 600 |
| *Salmonella* Javiana | 600 |
| *Salmonella* Johannesburg | 534 |
| *Salmonella* Kentucky | 600 |
| *Salmonella* Mbandaka | 535 |
| *Salmonella* Montevideo | 600 |
| *Salmonella* Muenchen | 600 |
| *Salmonella* Newport | 600 |
| *Salmonella* Schwarzengrund | 600 |
| *Salmonella* Senftenberg | 563 |
| *Salmonella* Thompson | 600 |
| *Salmonella* Typhimurium | 600 |

Table S1. Total counts of downloaded genomes per serovar of *S. enterica* Subspecies *enterica* Lineage I. All downloaded genomes were freely available at NCBI-SRA.

| Program | Version | Description | Databases | Link | Reference |
|---|---|---|---|---|---|
| **parallel-fastq-dump** | 0.6 | Parallel wrapper for SRA Toolkit | No | https://github.com/rvalieris/parallel-fastq-dump | [58] |
| **Trimmo-matic** | 0.38 | Trimming tool for Illumina NGS reads | No | https://github.com/timflutre/trimmomatic | [50] |
| **FastQC** | 0.11 | Tool to quality control for sequencing data | No | https://github.com/s-andrews/FastQC | [51] |
| **SPAdes** | 3.13 | Genome assembler | No | https://github.com/ablab/spades | [52] |
| **QUAST** | 5.0 | Evaluation tool for genome assembly | No | https://github.com/ablab/quast | [53] |
| **MLST** | 2.16.2 | Tool for multilocus-sequence typing | Integrated set of PubMLST databases for multiple organisms that can also be customized. | https://github.com/tseemann/mlst | [22] |
| **stringMLST** | 0.6.3 | Tool from multilocus-sequence typing of raw genome | Integrated set of PubMLST databases for multiple | https://github.com/jordanlab/stringMLST | [17] |

| | | | | | |
|---|---|---|---|---|---|
| | | sequencing reads | organisms that can also be customized. | | |
| **AMOS** | 3.1 | Collection of tools for genome assembly and related statistics | No | http://amos.sourceforge.net/wiki/index.php/AMOS | [54] |
| **EMBOSS** | 6.6 | European Molecular Biology Open Software Suite that contains multiple bioinformatics functionalities | No | http://emboss.open-bio.org/ | [55] |
| **DSK** | 2.2.0 | Tool for counting kmers from reads or genomes | No | https://github.com/GATB/dsk/ | [56] |
| **GNU Bash** | 4.4.20 | Unix shell and command tool | No | https://www.gnu.org/software/bash/ | |

Table S2. Description of bioinformatics programs used including their versions, available databases, links and references.

| *L. monocytogenes* | | | *C. jejuni* | | | *S. enterica* | | | *S. aureus* | | |
|---|---|---|---|---|---|---|---|---|---|---|---|
| **gene/locus name** | **number of genes/loci** | | **gene/locus name** | **number of genes/loci** | | **gene/locus name** | **number of genes/loci** | | **gene/locus name** | **number of genes/loci** | |
| | **string MLST** | **mlst** | | **string MLST** | **mlst** | | **string MLST** | **mlst** | | **string MLST** | **mlst** |
| **abcZ** | 423 | 255 | **aspA** | 527 | 489 | **aroC** | 1024 | 813 | **arcC** | 761 | 621 |
| **bglA** | 373 | 220 | **glnA** | 712 | 655 | **dnaN** | 1015 | 804 | **aroE** | 944 | 779 |
| **cat** | 402 | 241 | **gltA** | 608 | 563 | **hemD** | 949 | 765 | **glpF** | 845 | 705 |
| **dapE** | 535 | 308 | **glyA** | 806 | 733 | **hisD** | 1428 | 1132 | **gmk** | 506 | 399 |
| **dat** | 315 | 200 | **pgm** | 1082 | 969 | **purE** | 1115 | 848 | **pta** | 782 | 641 |
| **ldh** | 623 | 461 | **tkt** | 829 | 761 | **sucA** | 1037 | 839 | **tpi** | 721 | 579 |
| **lhkA** | 360 | 218 | **uncA** | 628 | 565 | **thrA** | 1206 | 952 | **yqiL** | 872 | 694 |
| | **Total number of STs** | **Total number of STs** | | **Total number of STs** | **Total number of STs** | | **Total number of STs** | **Total number of STs** | | **Total number of STs** | **Total number of STs** |
| **ST database** | 2323 | 1504 | | 11167 | 9745 | | 6992 | 5151 | | 6447 | 5219 |

Table S3. Distribution of counts for each allele across all seven loci per bacterial species and program (mlst vs. stringMLST), including the total count of unique STs present in the ST scheme available in the program.

| Organism | Number of raw Illumina reads | Number of Illumina reads after filtering | Number of reads after MLST |
|---|---|---|---|
| *Salmonella* Agona | 100 | 100 | 100 |
| *Salmonella* Anatum | 100 | 100 | 100 |
| *Salmonella* Braenderup | 100 | 100 | 100 |
| *Salmonella* Derby | 100 | 100 | 100 |
| *Salmonella* Dublin | 100 | 100 | 100 |
| *Salmonella* Enteritidis | 100 | 99 | 99 |
| *Salmonella* Hadar | 100 | 100 | 100 |
| *Salmonella* Heidelberg | 100 | 100 | 100 |
| *Salmonella* Infantis | 100 | 100 | 100 |
| *Salmonella* Javiana | 100 | 100 | 100 |
| *Salmonella* Johannesburg | 100 | 100 | 100 |
| *Salmonella* Kentucky | 100 | 100 | 100 |
| *Salmonella* Mbandaka | 100 | 100 | 100 |
| *Salmonella* Montevideo | 100 | 100 | 100 |
| *Salmonella* Muenchen | 100 | 96 | 96 |
| *Salmonella* Newport | 100 | 98 | 98 |
| *Salmonella* Schwarzengrund | 100 | 100 | 100 |
| *Salmonella* Senftenberg | 100 | 100 | 100 |
| *Salmonella* Thompson | 100 | 100 | 100 |
| *Salmonella* Typhimurium | 100 | 100 | 100 |
| *Acineto baumannii* | 1000 | 901 | 343 |
| *Campylobacter jejuni* | 1000 | 996 | 995 |
| *Clostridium difficile* | 1000 | 839 | 833 |
| *Enterococcus faecium* | 1000 | 915 | 913 |
| *Escherichia coli* | 1000 | 895 | 895 |
| *Haemophilus influenzae* | 1000 | 951 | 949 |
| *Helicobacter pylori* | 1000 | 874 | 874 |
| *Klebsiella pneumoniae* | 1000 | 920 | 918 |

| | | | |
|---|---|---|---|
| *Listeria monocytogenes* | 1000 | 998 | 996 |
| *Mycobacterium tuberculosis* | 1000 | 809 | 790 |
| *Neisseria gonorrhoeae* | 1000 | 915 | 915 |
| *Pseudomonas aeruginosa* | 1000 | 956 | 954 |
| *Staphylococcus aureus* | 1000 | 998 | 998 |
| *Streptococcus pneumoniae* | 1000 | 940 | 940 |
| *Enterococcus faecalis* | 100 | 92 | 92 |
| *Enterococcus faecium* | 100 | 92 | 92 |
| *Helicobacter pylori* | 100 | 81 | 81 |

Table S4. Number of reads used for calculating percentage of agreement between mlst and stringMLST for 23 *Salmonella enterica* serovars and 14 phylogenetic divergent bacterial pathogens.
The datasets used to calculate percentage of agreement contained 100 or 1,000 paired-end raw Illumina reads downloaded from NCBI-SRA (first column in the Table). Before running mlst, a few steps were required, such as quality trimming and adapter clipping, *de novo* assembly and assembly filtering. After the filtering steps were completed, a fraction of all genomes were discarded from the mlst analyses. This number is shown in the second column in the Table. Additionally, mlst infrequently adds empty columns in its results, and these records were removed before conducting the final statistical analyses to calculate the percentage of agreement between programs (third column in the Table). All downloaded genomes were freely available at NCBI-SRA.
